# *Clostridioides difficile* Toxins Unhinged: Allosterically Switchable Network Orients *β*-flap

**DOI:** 10.1101/2024.08.08.607137

**Authors:** Lauren M. Finn, Rebecca Cummer, Bastien Castagner, Bettina G. Keller

**Affiliations:** Freie Universität Berlin, Department of Biology, Chemistry, Pharmacy, Arnimallee 22, 14195 Berlin, Germany; Department of Pharmacology and Therapeutics, Faculty of Medicine and Health Sciences, McGill University, 3655 Promenade Sir-William-Osler, Montreal, Quebec, H3G 1Y6, Canada

## Abstract

Allosteric proteins exhibit a functional response upon substrate binding far from the active site. *Clostridioides difficile* toxins use allosteric binding by an endogenous co-factor to orchestrate self-cleavage from within the target cell. This binding event induces a conformational shift, primarily effecting a lever-like “*β*-flap” region, with two known orientations. We uncovered a mechanism for this allosteric transition using extensive atomistic MD simulations and computational and experimental mutagenesis. The mechanism relies on a switchable interaction network. The most prominent interaction pair is K600–E743, with K600 interactions explaining ∼70 % of the allosteric effect. Rather than gradually morphing between two end states, the interaction network adopts two mutually exclusive configurations in the active and inactive state. Similar switchable networks may explain allostery more broadly. This mechanism in particular could aid in drug development targeting the *Clostridioides difficile* toxins autoproteolysis.

## 1 Introduction

*Clostridioides difficile* infection (CDI) poses a major and growing public health threat, with symptoms ranging from diarrhea to major colon inflammation to death [1–3]. The impact is exacerbated by antibiotic resistance, more virulent strains and high recurrence of infection [4, 5]. The first line of treatment is antibiotic therapy, which paradoxically makes patients more susceptible to subsequent CDI [6]. CDI acts primarily via the cytotoxins TcdA and TcdB that enter and disrupt epithelial cells. As an alternative to antibiotics, targeting the cytotoxins could provide a route to relieve selective pressure and deliver effective treatment to patients [7–10]. Therefore, we consider these two cytotoxins [11, 12] as compelling systems for a detailed mechanistic investigation.

TcdB’s multiple domains work together in a precisely orchestrated way, enabling effective pathogenesis in the target cell [10]. The toxin’s four structural domains are coupled to functional roles [13, 14] by the following general cellular intoxication route: **1**) the receptor-binding domain binds to cell receptors and triggers endocytosis [15, 16], **2**) the low pH in the endosome results in a conformational change in the toxin [17], **3**) pore formation by the translocation domain allows the cysteine protease domain (CPD) and toxic glycosylation domains to translocate into the cell cytosol [18], **4**) the endogenous co-factor, inositol hexakisphosphate (IP6), binds and triggers an allosteric conformational change in the CPD [19, 20], **5**) this change promotes autoproteolytic cleavage of the glycosylation domain by CPD, releasing the glycosylation domain into the cell cytosol [21, 22], and **6**) the liberated domain glucosylates Rho and Rac GTPases causing actin cytoskeletal disruption, cell signaling interference, and ultimately apoptosis [23, 24].

Since autocleavage is a critical step in TcdB pathogenesis and is amenable to small molecule inhibition or premature activation, the CPD is an attractive therapeutic target [8, 28–31]. We focus on the allosteric transition in step **4)** which activates CPD. The *β*-flap substructure, depicted in *Figure 1*, straddles the allosteric pocket and active site residues in the target TcdB CPD. A *β*-flap is also found in several other toxin CPDs, and has long been suspected to play a significant role in signal modulation [32–34]. Shen *et al*. identified the “*β*-flap chain”, a chain of residues in the TcdB CPD *β*-flap, which connect the allosteric pocket to the active site [26]. The authors demonstrated the *β*-flap chain must be intact for allosteric function and concluded that the *β*-flap forms in response to IP6 binding, as previously hypothesized [26, 35, 36]. Interestingly, the *β*-flap and all *β*-flap chain interactions appear to be fully-formed in the later-published apo structure [25], offering a different perspective on the structural response to IP6 binding. Comparison of X-ray crystal structures with and without IP6 bound reveal the *β*-flap instead undergoes an ∼ 90^°^ rotation [25].

**Figure 1:**
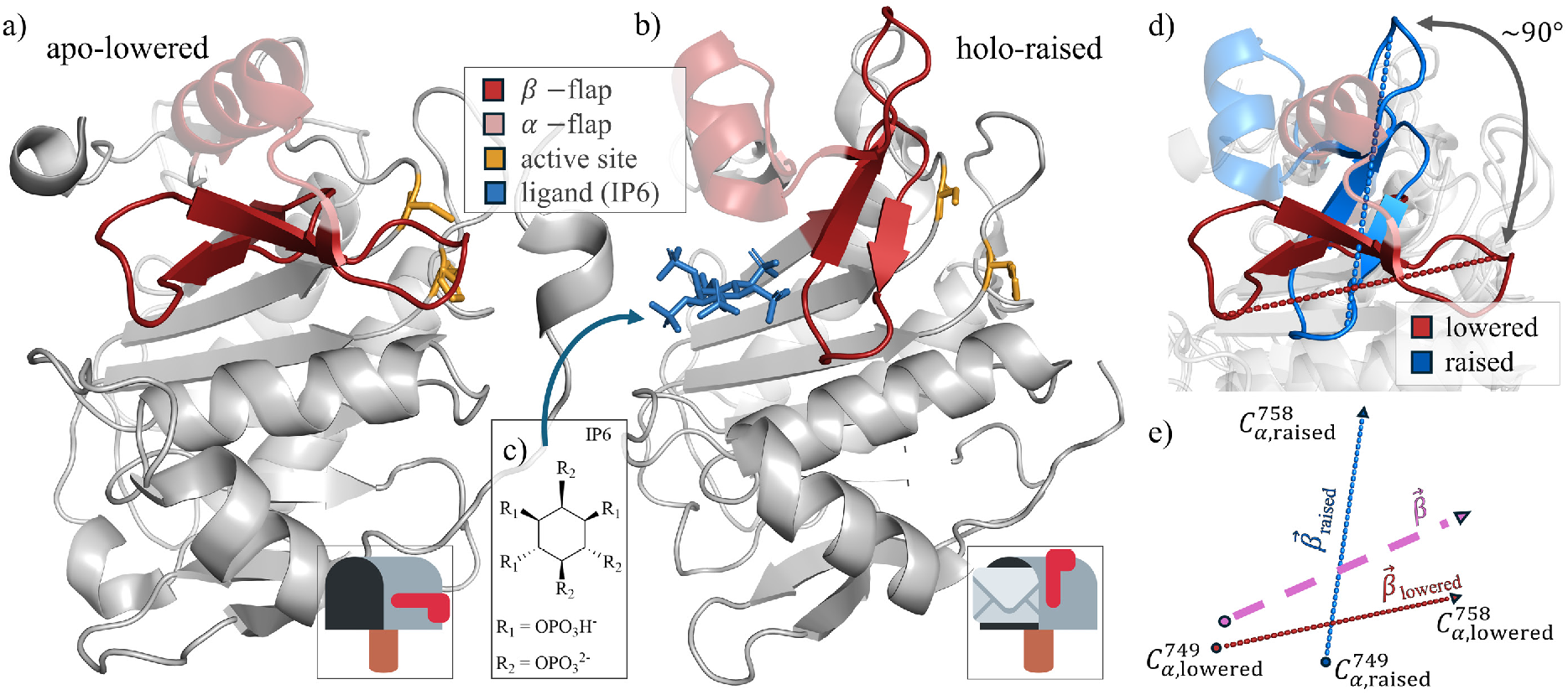
Functional organization of TcdB CPD in the a) apo-lowered (PDB entry: 6OQ5 [25]) and b) holo-raised states (PDB entry: 3PEE [26]). The mailbox icons [27] encapsulate the relation between ligand binding (no mail/mail) and *β*-flap orientation (lowered/raised lever). c) Structure of the ligand IP6 with the simulated protonation microstate indicated. d) Overlay of lowered conformation (red) and raised conformation (blue). e) Schematic of the *β*-flap orientation vectors, used to define the reaction coordinates *ξ*_1_ and *ξ*_2_. The vectors from reference structures are labeled 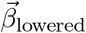 in red and 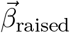 in blue. The vector in pink, 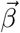, represents an arbitrary simulated structure.

It is well established that IP6 binding distal to the active site impacts the conformational distribution and vastly increases the activity of TcdB CPD [25, 26, 36]. However it remains a topic of discussion whether the change in shape is truly responsible for the change in protease activity. While the apo and holo crystal structures imply that *β*-flap rotation exposes the active site, mutagenesis studies [26, 29, 37] indicate the conformational change is just the first step in the activation of autoproteolytic catalysis and residues outside the *β*-flap may be involved in the allosteric transition. Despite this rich body of experimental data, a detailed model of the allosteric mechanism is lacking. The purpose of this study is to leverage ∼150 μs of molecular dynamics (MD) simulations to calculate the free-energy landscape of the allosteric transition [38–42] and provide an atomistic picture of the conformational changes triggered by IP6 binding.

## 2 Results

### 2.1 Overview of the system

The apo- and holo-structure [25, 26] are depicted in *Figure 1.a* and *1.b*. The lysine- and arginine-rich allosteric pocket can bind the ligand IP6 and is distal from the catalytic dyad in the active site (yellow). The *β*-flap (residues 738-761) straddles these functional regions, with the α-flap (residues 762-776) connecting to it and forming part of the allosteric pocket. The *β*-flap orientation determines active site exposure, analogous to a hinging lever. We therefore name the *β*-flap conformations in *Figure 1.a* and *Figure 1.b* the “lowered” and “raised” conformations, respectively. The largest structural deviations between lowered and raised conformations occur in the α −flap (C_*α*_ RMSD =7.9 Å) and the *β*-flap (C_*α*_ RMSD =10.4 Å), which rotates by approximately 90° (*Figure 1.d*). Otherwise the two conformations align strongly, with a low C_*α*_ RMSD of 2.1 Å for the remaining residues.

The ligand molecule IP6 has 64 conceivable microprotonation states. ^31^P NMR titration curves enable elucidation of pK_a_ values for the individual phosphates in solution [43]. The predominant net charge under physiological conditions is −8 [43]. Isothermal titration calorimetry (ITC) in buffers of various ionic strength demonstrate IP6 loses one proton upon binding TcdB CPD (see *Figure S7.d* and [31]), leading to a charge state of −9 when IP6 is bound to TcdB CPD. *Figure 1.c* shows the microprotonation state of IP6 which we use in our simulations of holo CPD.

We modelled the protonation states of all residues in the apo CPD according to the prediction by PROPKA [44]. However, IP6 losing a proton upon binding implies that simultaneously a residue in the binding pocket receives this proton and changes its protonation state. In fact, simulations of holo CPD with various protonation states showed that the IP6 only remains in the binding pocket if K775 and additionally K600 are modelled as protonated. To account for the combined effect of ligand binding and proton transfer, our biased simulations of apo and holo CPD differ in three aspects: presence of IP6 (as shown in *Figure 1.c*), protonation of K775 and protonation of K600.

### 2.2 *β***-flap is stable in both conformations**

Shen *et al*. proposed a chain of residues as a modulator of allostery, which we refer to as the *β*-flap chain: (IP6)–R751–N747–E743–R745–W761 [26]. The involved residues shown in *Figure S2.a* connect the allosteric pocket to the active site. Mutational studies demonstrate that the intact *β*-flap chain is important for protease activity, [26] which the authors interpreted as an indication that the *β*-flap forms in response to IP6 binding. This hypothesis was revisited with the apo structure [25], where the *β*-flap and its affiliated chain appeared to be merely rotated, as shown in *Figure S2.b*.

Distance histograms in *Figures S2.c-f* for the residue pairs in the *β*-flap chain are based on 10 μs MD simulations for all four combinations of ligand binding state (apo/holo) and conformation (raised/lowered). The histograms for each of the four states are remarkably indistinguishable, confirming that the *β*-flap chain remains intact and is independent of IP6 binding and conformation. This prompts the question: what other interactions drive the allosteric conformational transition?

### 2.3 Ligand Binding Perturbs the FES

Even apo TcdB CPD has a residual enzymatic activity. Evidence for this comes from partial labeling with an activity-based protein profiling probe in the absence of IP6, which occurs exclusively in the activated form [26]. This result indicates that in the apo-state both conformations, lowered and raised *β*-flap, are accessible, but ligand binding dramatically shifts the equilibrium to make the raised conformation more stable. We used umbrella sampling [45, 46] and weighted histogram analysis method (WHAM) [47–50] to characterize the change in the free-energy landscape of the protein that gives rise to this shift in equilibrium. The expected outcome [38] is a free-energy landscape with two minima, whose relative depth changes upon ligand binding (see *Figure S3*).

Defining suitable collective variables is essential for constructing the free-energy surface. The most prominent conformational change in the allosteric transition of CPD is the ∼ 90^°^ rotation of the *β*-flap. We have accordingly defined two collective variables, *ξ*_1_ and *ξ*_2_, to monitor *β*-flap rotation during the simulation. The orientation of the *β*-flap in each simulation frame is defined by the vector between the C_*α*_-atoms of residue 749 and residue 758 (*Figures 1.d,e*). The collective variables, *ξ*_1_ and *ξ*_2_, are the orientation of this vector (dashed pink in *Figure 1.e*) with respect to the corresponding vector in the lowered conformation (dotted red in *Figure 1.e*) and with respect to the corresponding vector in the raised conformation (dotted blue in *Figure 1.e*). We measure the orientations as the scalar product of each of the two vector pairs (see *Section SI S1.1*). Correspondingly, the lowered conformation occurs for small *ξ*_1_ and large *ξ*_2_, and the inverse is true for the raised conformation.

Before constructing the free-energy surface, we checked whether spontaneous transitions between the lowered and the raised state can be observed in unbiased MD simulations. *Figure S4* shows simulations of the four possible states of CPD (apo/holo + lowered/raised) projected on our two-dimensional collective variable space. Within 10 *μ*s, none of the simulations show a transition between lowered and raised state. Additionally, we enriched the conformational ensemble using AlphaFold [51, 52]. The conformations suggested by AlphaFold indicate a pathway which connects the lowered and the raised conformation (see *Section SI S4*). However, simulations started from these structures reverted back to the raised conformation and did not show spontaneous transitions. Thus, enhanced sampling simulations are needed to study the allosteric transition in CPD.

The computed free energy surfaces in the two-dimensional space spanning *ξ*_1_ and *ξ*_2_ are shown in *Figures 2.a and 2.b* for the apo and holo states, respectively. The statistical uncertainty estimates fall below 1 kJ/mol (*Figure S8.a,b*). The free-energy surfaces show the expected two minima corresponding to the lowered and raised conformation of the *β*-flap. In the free-energy surface of the apo-state, the two free-energy minima coincide with the crystal structure of the lowered and raised conformation. In the holo-state, the free-energy minimum of the lowered conformation is shifted towards larger values of *ξ*_1_, i.e. away from the reference structure of the lowered conformation.

**Figure 2:**
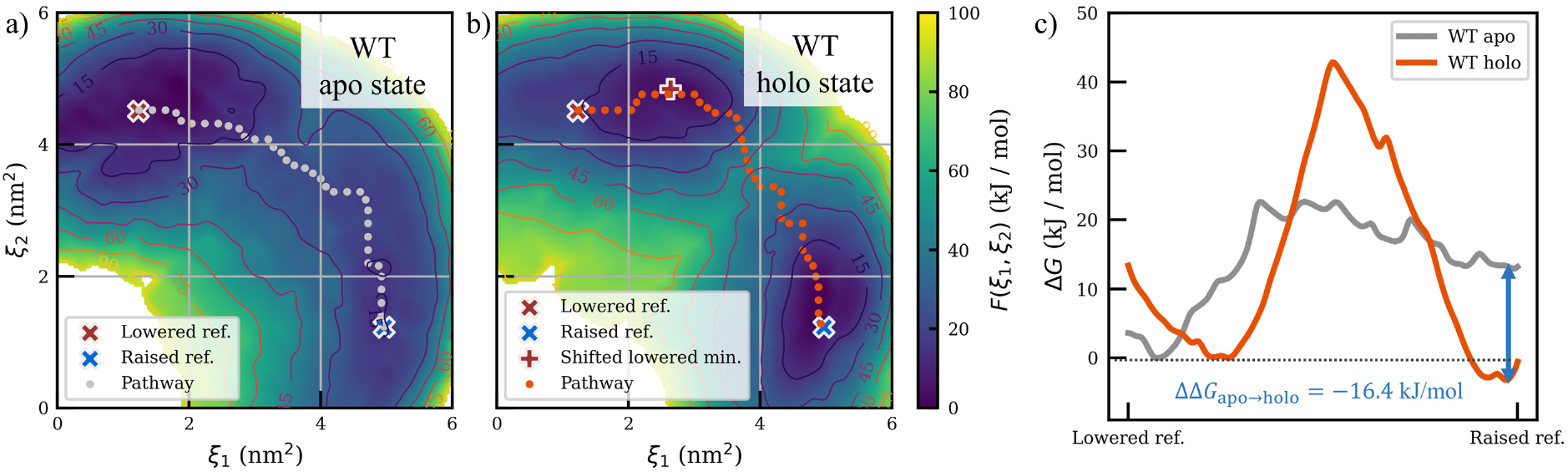
Computed 2D free-energy surface in the *ξ*_1_, *ξ*_2_ reaction coordinate space for a) apo and b) holo states. Positions of reference structures in *ξ*_1_, *ξ*_2_ space are indicated with “X”s and in b) the “+” indicates the free-energy minimum of the lowered conformation relative to the reference lowered conformation. c) The free energy profiles along the estimated lowest free energy paths as indicated in a) and b).

The influence of IP6-binding on the free-energy surface is concisely demonstrated by plotting the freeenergy along the lowest free-energy path (*Figure 2.c*). In the absence of IP6, the raised conformation is ∼ 14 kJ/mol higher in free energy than the lowered conformation, thus making the lowered conformation the by far more populated conformation. Binding of IP6 stabilizes the raised conformation, such that, in the holo state, it has ∼ 2.5 kJ/mol less free energy than the lowered conformation. Thus, the allostery in the CPD exhibits bistable switch dynamics [38], where the binding of IP6 stabilizes the raised conformation by ΔΔ*G*_apo→holo_ = −16.4 kJ/mol (see Tab. 1).

**Table 1:** Summary of free energy differences between the lowered and raised conformational states, Δ*G*_lowered→raised_, computed with 2D umbrella sampling + WHAM. The ΔΔ*G* values from the perspective of mutational state, ΔΔ*G*_WT→K600G_, and ligand binding state, ΔΔ*G*_apo→holo_, are also shown.

| $\Delta G_{\text{lowered} \rightarrow \text{raised}}$ | apo (kJ/mol) | holo (kJ/mol) | $\Delta\Delta G_{\text{apo} \rightarrow \text{holo}}$ |
| --- | --- | --- | --- |
| native | 13.9 | -2.5 | -16.4 |
| K600G | 8.7 | 4.2 | -4.5 |
| $\Delta\Delta G_{\text{WT} \rightarrow \text{K600G}}$ | -5.2 | 6.7 | |

Another notable characteristic of the free-energy surfaces in *Figure 2* is the significantly higher transition barrier in the holo-state compared to the apo-state. IP6 sterically hinders the transition in the holo-state, creating a higher transition barrier. Additionally, protonation of K775 might play a role as indicated by interactions K775–N740 and K775–E692, which are exclusively formed in the holo state, i.e. when K775 is protonated. See *Section SI S3* for a detailed discussion.

### 2.4 A switchable interaction network is the structural basis of the allosteric shift in the FES

The free-energy surface sheds light on the thermodynamics of the allosteric transition in CPD. Following this, we examined the MD data to discover the structural changes that communicate the IP6 binding signal throughout the protein, causing the *β*-flap to transition from a lowered to a raised conformation.

We analyzed residue pair distances and residue solvent accessible surface area (SASA) from the four unbiased simulations, focusing on residues in or near the α- and *β*-flaps, the allosteric pocket and the active site. Additionally, we evaluated average pair distances and relevant residue SASAs in the collective variable space spanned by *ξ*_1_ and *ξ*_2_ (*Figure 3* and *Figures S9* and *S10*). This revealed a set of residue pairs with distinctively binary distance distributions. In these residue pairs, one state was exclusively populated in the lowered conformation, while the other was exclusively populated in the raised conformation, as evidenced by minimal or no overlap of the one-dimensional distributions between simulations of the lowered conformation (red in *Figure 3*) and those of the raised conformation (blue in *Figure 3*). Furthermore, the average distances display a sharp demarcation between short and large distances in the two-dimensional collective variable space. In all of these residues pairs, short distances correspond to a stabilizing interaction, either a hydrogen bond or a salt-bridge. Thus, these residue pairs act as switches, with their interactions exclusively stabilizing one of the two *β*-flap conformations.

**Figure 3:**
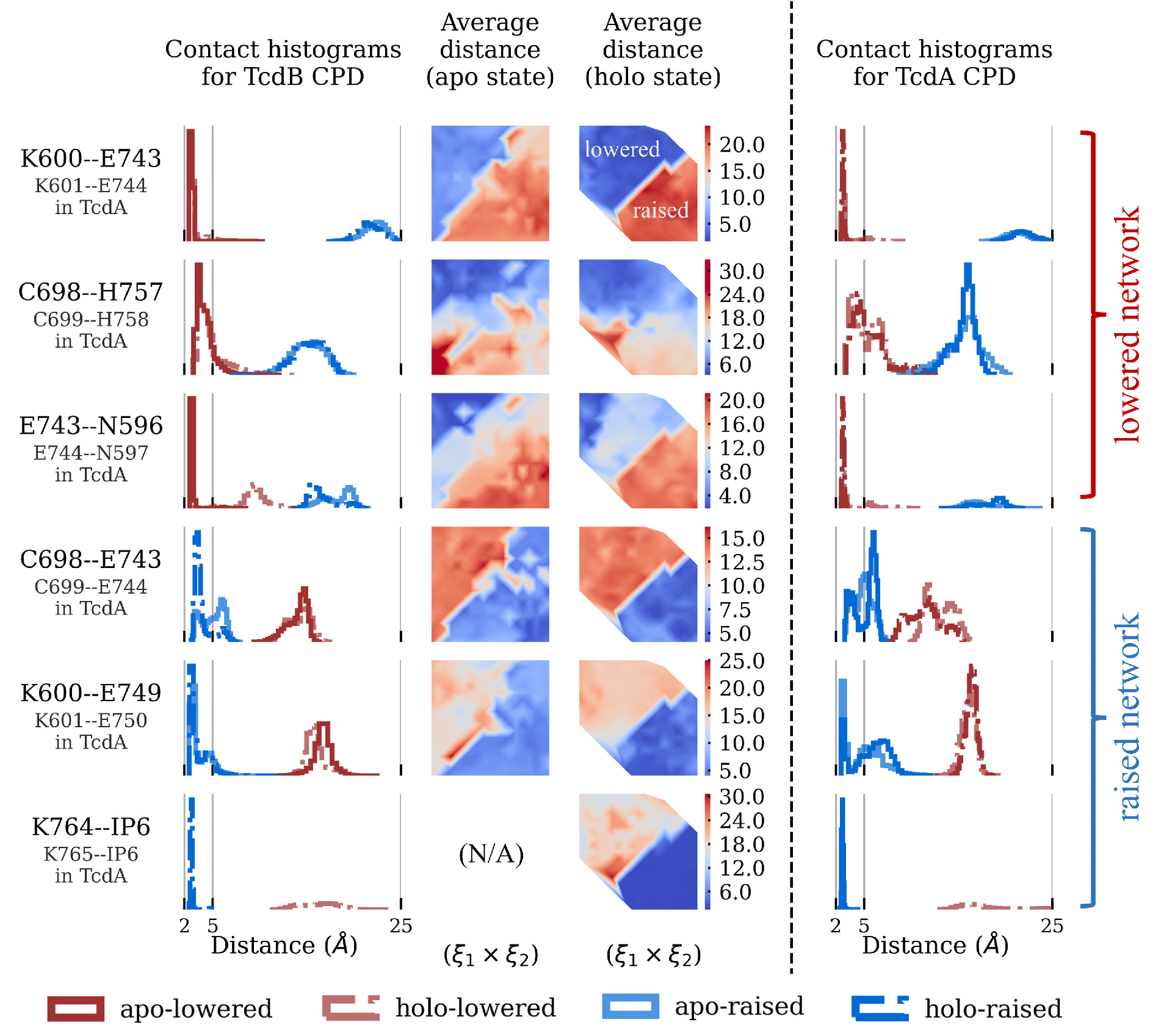
Pairwise residue interactions in the proposed interaction network. *Left column:* histograms of residue pair distances from four (apo/holo + lowered/raised) simulated states. *Two middle columns:* average residue distance in each umbrella window, for apo and holo state. These surfaces use the same reaction coordinate space (*ξ*_1_, *ξ*_2_) as in the free-energy surface (see *Figure 2*), in which the upper left and lower right correspond to the lowered and raised conformations, respectively. *Right column:* histograms of residue pair distances for each analogous interaction in TcdA CPD.

The residue pairs which exhibit switchable interactions form a network which extends from the allosteric pocket to the active site and links the *β*-flap to the core of the protein (*Figure 4*). In the lowered conformation, the *β*-flap is held in place by two crucial interactions between E743 in the *β*-flap and residues in the core of the protein: K600–E743 and N595–E743. Furthermore, the catalytic residue C698 is shielded from the solvent by a hydrogen bond to H757. In the raised conformation, these four key residues assume a complementary interaction pattern. The interactions of E743 to the core of the protein are broken. K600 instead forms an interaction to E749, and N595 forms a hydrogen bond to E592. This allows the *β*-flap to rotate away from the core, breaking the H757–C698 hydrogen bond. H757 becomes solvent-exposed. Due to the rotation of the *β*-flap, E743 moves into the vicinity of C698 and forms a hydrogen bond to this catalytic residue.

**Figure 4:**
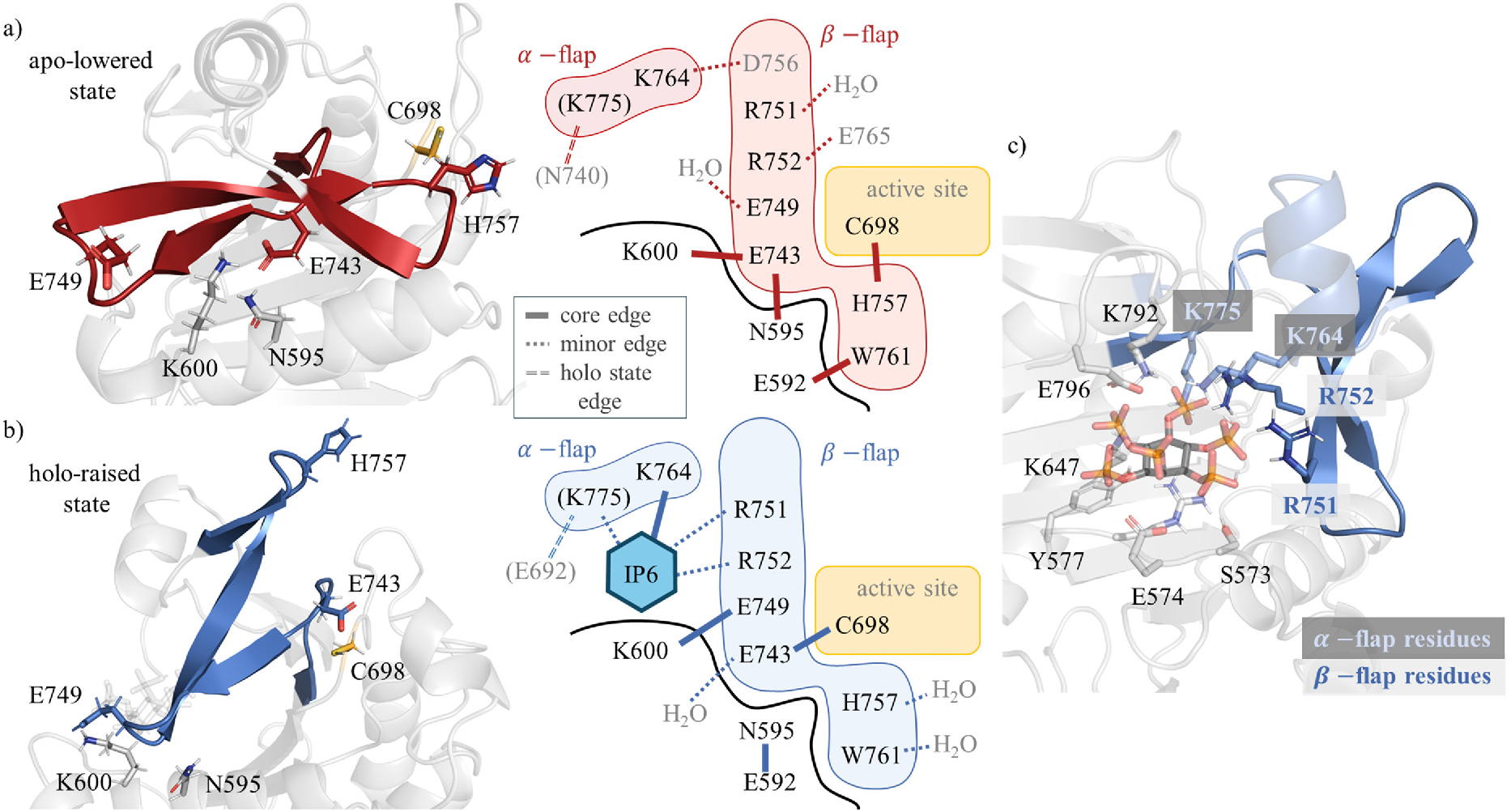
Switchable interaction network. a,b) On the left, representative structures extracted from unbiased a) apo-lowered and b) holo-raised simulations with selected residue side chains in the interaction network shown as sticks. On the right, schematics of the interaction network in the a) lowered and b) raised conformations. c) The allosteric binding pocket of TcdB CPD (simulated structure), with residues within 4 Å of IP6 shown as sticks and residues exhibiting conformation dependence in blue.

The interaction network described so far is present in both the apo and holo states of CPD, assuming two complementary configurations. However, the network extends well beyond those four key residues and offers a plausible explanation for how binding of IP6 shifts the equilibrium from lowered to raised conformation. In the allosteric pocket, residues K764 and R751 show a strong, and R752 and K775 show a weak, conformation-dependent interaction with IP6. All four contacts to IP6 stabilize the raised conformation. The contact between K764 and IP6 becomes possible because the α-flap moves in concert with the *β*-flap and thereby approaches the allosteric pocket. Similarly, R751 and R752 connect the raised *β*-flap orientation to the IP6 binding state. K775 does not move a lot during the lowered-to-raise transition, but it nonetheless forms an important interaction with IP6.

We had initially anticipated a strong conformation dependence for the salt-bridge K600–IP6, but we observe that K600 only interacts with IP6 in the holo-lowered state (see *Figure S9*). In the holo-raised state, it instead forms an interaction with E749 on the *β*-flap. K600–IP6 is likely an important interaction for destabilizing the K600–E743 interaction, which initiates *β*-flap rotation. Additionally, the K600–E743 interaction might lock the *β*-flap in the raised conformation. Once rotated, R751, R752, K764 and K775 stabilize the raised conformation via salt-bridges to IP6.

In summary, the following allosteric mechanism emerges:

- IP6 binds and forms a transient interaction with K600, thereby breaking the K600–E743 interaction that keeps the *β*-flap in the lowered conformation.
- The *β*-flap rotates into the raised conformation, which breaks the H757–C698 hydrogen bond and enables the complementary hydrogen bond C698–E743.
- The raised conformation is stabilized by salt bridges from IP6 to the *β*-flap (R751–IP6, R752–IP6) and to the α-flap (K775–IP6, K764–IP6).

The increased transition barrier in the holo-state (*Figure2.c*) might contribute to the overall allosteric effect. The barrier likely only arises after IP6 has entered the allosteric pocket, because both steric hindrance and proton transfer can only occur if IP6 is bound. Thus, the barrier might kinetically trap the system in the raised conformation. A comprehensive picture is emerging in which the allosteric transition from lowered to raised *β*-flap orientation operates by toggling the configuration of the central interaction network, which is triggered and kinetically stabilized by ligand binding and proton transfer.

### 2.5 Switchable interaction network explains experimental results

Our proposed interaction network corroborates several previous experimental findings and explains other results that were not fully understood.

For instance, Savidge *et al*. found that in the mutant E743A the cysteine protease activity already sets in at a much lower concentration of IP6 than in the wild type [29]. While the raised conformation is necessary for activation by exposing the catalytic site to the solvent, our network shows that it is likely not yet an active conformation. In the interaction network of the raised conformation, the catalytic residue C698 is engaged in a hydrogen bond to E743 (*Figure 4.b*), as previously observed [29, 37]. Thus, the E743–C698 hydrogen bond effectively serves as a safety latch which must break for the active site cysteine to perform catalysis. In the E743A mutant this hydrogen bond cannot be formed, which explains the mutant’s increased activity.

Our findings additionally clarify an apparent contradiction in the conformation-dependent fluorescence of tryptophan. The residue W761 on the *β*-flap serves as a fluorescent probe to the conformational state since tryptophan fluorescence is sensitive to its environment, and W761 is the sole tryptophan in TcdB CPD [26, 53]. Shen *et al*. observed higher fluorescence on addition of IP6, which the authors attributed to W761 entering a more hydrophobic environment [26]. This would mean that W761 had a high solvent accessible surface area (SASA) in the lowered conformation (attributed to the apo state), which quenches its fluorescence, and low SASA in the raised conformation (attributed to the holo state), which allows for unquenched high fluorescence. However, the W761 SASA sampled with MD show the exact opposite: low SASA in the lowered conformation and high SASA in the raised conformation (*Figures 5.a* and S5). Furthermore, the SASA distributions sampled by MD are consistent with the SASA-values of the experimental structures. The interaction network (*Figure 4.a,b*) and the distance histograms in *Figure 5.b* show that residue E592 interacts with W761 in the lowered conformation only. Thus, a more plausible explanation for the conformation dependence of W761 fluorescence is that in the lowered conformation, fluorescence is quenched by the interaction with the negatively charged glutamic acid [54]. Conversely, in the raised conformation this interaction is replaced by weaker quenching interactions with the surrounding solvent.

**Figure 5:**
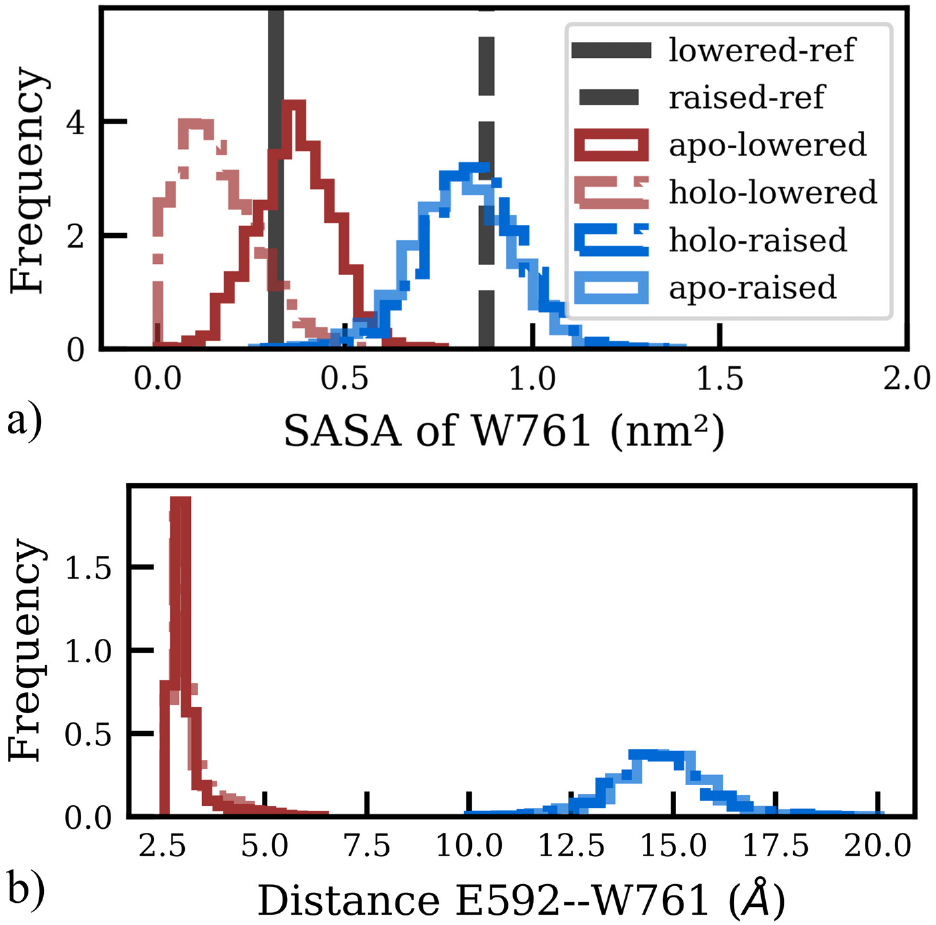
The local environment of W761 depends on conformation. a) The SASA distribution of W761 in four unbiased, simulated states (apo/holo + lowered/raised) and the reference SASA of W761 for the raised conformation (PDB entry: 3PEE [26]) and the lowered conformation (PDB entry: 6OQ5 [25]). b) The E592-W761 distance distribution for the four unbiased simulated states.

Mutations to residues in the allosteric pocket and the *β*-flap were investigated by Shen *et al*. using a specifically designed activity probe and W761 fluorescence [26, 30]. Some mutations involve residues in the interaction network, including K600N, K775N, K764N, R751Q, W761A and R752Q. These mutations resulted in a moderate to strong decrease on measured activity and/or tryptophan fluorescence, supporting the role of these residues in the allosteric interaction network. On the other hand, the mutations E753N, E753R/R745E, N747A, and R745N in the *β*-flap also decreased the measured activity/fluorescence. We believe this effect can be attributed to destabilization of the *β*-flap structure, rather than to direct participation in allosteric communication.

### 2.6 Mutant K600G Stabilizes Raised Conformation

To test the interaction network as a modulator of allostery, we consider mutants which remove the core network edge from the lowered conformation, K600–E743. Assuming information travels via the proposed interaction network, such mutations mimic IP6 binding. We therefore use both *in silico* and *in vitro* mutation experiments to test the allosteric impact of the interaction network.

We simulated mutants K600G, E743G and K600G/E743G in the apo-lowered state to check whether eliminating the interaction can stimulate *β*-flap rotation to the raised conformation. The sampled regions in the *ξ*_1_, *ξ*_2_ reaction coordinate space in *Figure 6.a* indicate these perturbations are insufficient to reach the raised conformation within a practical simulation timescale. Some of the network interactions are weakened or even dynamically broken, as indicated by residue distances for C698–H757 and E592–W761 in *Figures 6.b,d*, respectively. These histograms indicate partial breakdown of the bistable character of the network upon mutation. The double mutant K600G/E743G crosses the apparent transition barrier, but reverts back to the lowered conformation, implying the true raised conformation minimum is not sampled. Therefore, biased methods are needed to characterize the impact of mutations on conformation (de)stabilization.

**Figure 6:**
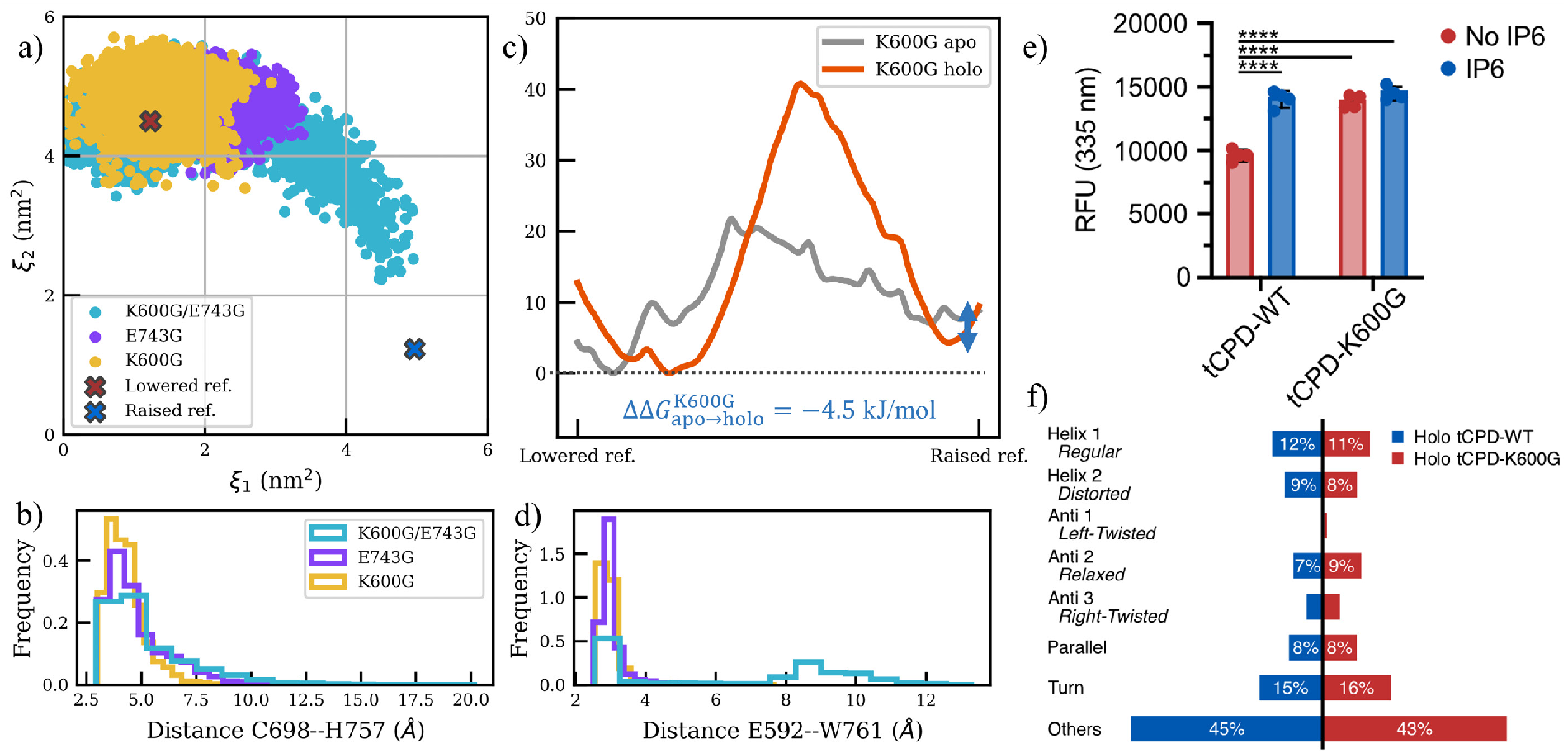
Deletion of the interaction K600–E743 by mutation a) The sampled regions in the *ξ*_1_, *ξ*_2_ reaction coordinate space for three apo state mutants initialized in the lowered conformation. b,d) Histograms of the residue pair distance, b) C698–H757 and d) E592–W761, for three mutant simulations, initialized in the lowered conformation. c) The free energy profiles for apo and holo K600G. e) Effect of the point mutation K600G on intrinsic tryptophan fluorescence of tCPD. Fluorescence emission of tCPD was measured at 335 nm after excitation at 295 nm in the absence (apo-tCPD) or presence (holo-tCPD) of IP6. Each fluorescence measurement was controlled for protein concentration (absorbance at 280 nm). Mean SD with data points, n = 4; Tukey’s Multiple Comparisons Test (MCT), **** p 0.0001. f) Secondary structure of holo-tCPD-WT and holo-tCPD-K600G as determined by circular dichroism (CD) to confirm the K600G point mutation did not modify the overall structure of tCPD. Secondary structure analysis was performed with BeStSe [55–57].

We computed the apo and holo free-energy surfaces for K600G using a two-dimensional umbrella sampling approach analogous to that applied in the native state analysis. The K600G free-energy surfaces are shown in *Figures S6* and the statistical error estimations are in the *Figure S8.c,d*. These surfaces retain the general features of the corresponding native state free-energy surfaces, including the shifted lowered conformation minima position in the holo state. Importantly, the mutation significantly alters the relative depths of the conformation minima, as summarized in *Table 1*.

The stabilization of the raised conformation due to IP6 binding (ΔΔ*G*_apo→holo_) is decreased from −16.4 kJ/mol in the wild type to just −4.5 kJ/mol in the mutant K600G. The free energy profiles along the corresponding low free energy pathways in *Figure 6.e*, illustrate the impact this mutation has on the ligand binding effect. Thus, the mutant K600 should show a much weaker allosteric response to the presence of IP6. In terms of free energy, the interactions of K600 with E743 in the lowered conformation and with E749 in the raised conformation account for about 70% 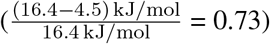 of the allosteric effect. Loss of the central network edge destabilizes the lowered conformation and partially mimics IP6 binding. These results strongly implicate the interaction K600-E743 in allosteric communication, and support the proposed switchable interaction network as a significant allosteric modulator.

We also tested the point mutation K600G for IP6 sensitivity experimentally, using truncated CPD (tCPD, TcdB_543−799_) in an intrinsic tryptophan fluorescence assay. The fluorescence of W761 differentiates the lowered and raised conformation [26]. The relative fluorescence of apo-lowered tCPD-WT was significantly less than holo-raised tCPD-WT, confirming the change in tryptophan local environment depends on IP6 binding (*Figure 6.e* and *S7.a,b*). In contrast, the relative fluorescence of tCPD-K600G is indistinguishable for apo and holo states, matching the fluorescence of holo tCPD-WT and presumably adopting the raised conformation (*Figure 6.e and S7.a-c*). In addition, to confirm the point mutation did not alter the overall structure of the protein, we determined the secondary structure of the protein variant via circular dichroism (CD). The secondary structural content was then calculated from this data. The CD analysis indicated no structural perturbations as both tCPD-WT and tCPD-K600G had a 3-layer (aba) sandwich fold-recognition with a nearly identical secondary structure (*Figure 6.f*).

### 2.7 The Interaction Network Generalizes to TcdA CPD

As a close homolog (56% identity), [26] TcdA CPD exhibits faithful alignment, and structural and functional similarity to TcdB CPD [34]. We expect to find a similar network of conformation-dependent switches in the TcdA CPD system, and therefore simulated TcdA CPD in four states (apo/holo + lowered/raised) Analysis of residue interactions reveal an analogous conformation-dependent network in TcdA CPD to the salt-bridge network of TcdB CPD (*Figures 3, S9* and *S10*). All residues in the proposed salt-bridge network have a corresponding residue in TcdA CPD, except for the conservative replacement of R752_TcdB_ with K753_TcdA_ and substitution of D756_TcdB_ with A757_TcdA_. While the interactions R751_TcdA_–IP6 and K765_TcdA_–A757_TcdA_ do not exhibit conformation dependence in TcdA CPD, the remaining network indicates the switch-based network mechanism of allosteric communication is very similar to that of TcdB CPD. These findings support the physiological role of the switchable interaction network, which we identified in TcdB CPD.

## 3 Online Methods

See supplementary material.

## 4 Discussion

Here we used extensive atomistic MD simulations and (computational and experimental) mutagenesis to derive an atomistic model for the allosteric transition in TcdB CPD triggered by IP6 binding. We showed that the crucial *β*-flap structure does not form in response to IP6 binding, as was previously thought [26, 35, 36]. Rather, IP6 initiates and stabilizes the ∼ 90^°^ rotation of the *β*-flap in the raised orientation, consistent with the previous finding that an intact *β*-flap chain is a prerequisite to CPD activity [26]. Free-energy surfaces of the apo and the holo state reproduce the expected shift in the free-energy surface and show that the allosteric stabilization of the raised conformation is about ΔΔ*G*_apo→holo_ = −16.4 kJ/mol.

Rather than gradual morphing between two end states, our MD simulations show that an interaction network with two mutually exclusive configurations distinguishes the lowered from the raised conformation. In this network, pairwise distances have strongly bimodal distributions, thus effectively switching between the lowered and raised conformations. The switchable interaction network is in place whether or not IP6 is bound. Binding of IP6 establishes additional interactions which transiently destabilize the lowered conformation and stabilize the raised conformation.

The switchable interaction network additionally allows for an understanding of the allosteric effect. The most prominent interaction, K600-E743, stabilizes the *β*-flap in the lowered conformation. IP6 destabilizes this interaction by forming a transient interaction to K600. With the K600–E743 interaction broken, the *β*-flap can rotate into the raised conformation and is stabilized by multiple interactions between IP6 and the *β*- and α-flaps. Mutational studies verify the crucial role of K600–E743. MD simulations show that the allosteric effect is reduced to ΔΔ*G*_apo→holo_ = −4.5 kJ/mol in the K600G mutant. Additionally, our fluorescence assays and circular dichorism experiments indicate that the conformational ensemble in K600G exhibits significantly reduced sensitivity to the presence of IP6 compared to the wild type, corroborating that the allosteric network is disrupted.

Beyond rotation of the *β*-flap, TcdB CPD activation requires a second step. The K600–E743 interaction of the *β*-flap residue E743 to K600 in the lowered conformation is replaced by a hydrogen bond to the catalytic residue C698 in the raised conformation. This obstructs the nucleophilic attack of the cysteine thiol group on a substrate. Thus, the hydrogen bond E743–C698 functions as a safety latch on the irreversible autoproteolytic reaction, discouraging premature release of the toxic domain.

An intriguing drug design strategy against CDI is to inactivate the toxin by IP6 mimetics which allosterically activate CPD and preemptively trigger autoproteolysis before the toxin enters the cell [8]. The K600–E743 interaction perfectly explains structure activity relationship results with IP6 analogs that show the importance of a negatively charged group in position 5 of the inositol core that interacts with K600 [31]. Furthermore, Puri *et al*. designed covalent CPD inhibitors based on the TcdB substrate and found that inhibitors with a lysine or arginine in position P2 were “unexpectedly” more potent than a serine that mimics the natural substrate [30]. Position P2 is in close proximity to E743. Replacing the serine hydrogen bond to E743 by a salt bridge additionally stabilizes the raised conformation. Thus, disrupting K600–E743 and/or stabilizing E743 in the raised conformation emerges as an important design consideration for (allosteric) drugs towards TcdB CPD. Other key interactions include those between ligand and R751, R752, K764 and K775, and E743–C698 to severe the catalytic safety latch.

We demonstrated that TcdA contains an interaction network similar to TcdB, reinforcing the hypothesis that this network serves a physiological purpose. In fact, the significance of the K600–E743 interaction within the allosteric network identified here may extend to other bacterial toxins and effectors that use IP6 as a co-factor to achieve a stable active conformation after translocation through a pore or type III secretion system. Indeed TcdB and TcdA have 76% and 77% sequence identity with the TcsL and TcsH of *Paeniclostridium sordellii* and the key E743 and K600 residues are conserved [24].

Switchable interaction networks also occur in allosteric proteins unrelated to TcdB. In C-type lectin Langerin, Ca^2+^-release from the binding pocket is allosterically triggered by the protonation of a histidine residue [41, 58]. This protonation breaks a crucial hydrogen bond in the Ca^2+^-bound conformation, and thereby gives access to a interaction network configuration in which Ca^2+^ can unbind (*Figure 5* in [41]). Switchable interaction networks which separate a protein’s active and inactive conformation might in fact be at the heart of allostery. An allosteric ligand then stabilizes the active state by additional interactions, as in TcdB CPD, or to destabilize the inactive state by breaking a crucial interaction, as in Langerin and in TcdB CPD. Understanding and controlling allostery requires a detailed understanding of the switchable interaction network and a ligand design strategy that targets critical interactions in this network.

## Supporting information

Scripts, Structures etc.

## 5 Acknowledgements

Funded by Deutsche Forschungsgemeinschaft (DFG) through grant IRTG 2662, Project Number 434130070, project B3, Canadian Institutes of Health Research (CIHR) project grant (PJT-173262) to BC Natural Science and Engineering Research council of Canada (NSERC) through discovery grant (RGPIN-2020-04908) to BC. Computational resources were provided by FUB-IT at Free University Berlin and the Paderborn Center for Parallel Computing (PC2) at Paderborn University. We acknowledge Dr. Matthew Bogyo and Dr. Aimee Shen for providing the tCPD544-797 plasmid, and Dr. Kim Munro and the Centre for Structural Biology Research (CRBS) for use of the core facilities.

