## Supplementary Information for "*Clostridioides difficile* Toxins Unhinged: Allosterically Switchable Network Orients *β*-flap"

August 8, 2024

### 1 Methods

#### 1.1 Collective Variable

We constructed a reaction coordinate system to capture the rotation of the $\beta-$flap for any simulated structure, relative to the lowered and raised endstate conformations (see *Section SI 1.2* for the pdb identifiers of the reference structures). We first introduce $\vec{\beta}$ to quantify the orientation of the $\beta-$flap as,

$$\vec{\beta} \equiv \vec{r}_{\mathrm{C}_\alpha^{749}} - \vec{r}_{\mathrm{C}_\alpha^{758}} \tag{1}$$

where $\vec{r}_{\mathrm{C}_\alpha^{749}}$ and $\vec{r}_{\mathrm{C}_\alpha^{758}}$ are the positions of two $\mathrm{C}_\alpha$'s of residues 749 and 758 in the $\beta-$flap. We arbitrarily chose a single reference structure from an unbiased simulation and used it throughout the project. The pdb structures of the lowered and raised conformation, as well as each simulated structure were aligned to this reference structure using a set of residues with low $\mathrm{C}_\alpha$ RMSF (see *Section SI 1.6*), to determine the quantities $\vec{\beta}_{\text{lowered}}$, $\vec{\beta}_{\text{raised}}$, and $\vec{\beta}$ using *eq. 1*. The two reaction coordinates, $\xi_1$ and $\xi_2$, were then computed to quantify the strength of $\vec{\beta}$ alignment to the lowered and raised endstates as the dot product of $\vec{\beta}$ with the respective $\vec{\beta}_{\text{ref.}}$, or,

$$\xi_2 = \vec{\beta} \cdot \vec{\beta}_{\text{lowered}}. \tag{2}$$

$$\xi_1 = \vec{\beta} \cdot \vec{\beta}_{\text{raised}} \tag{3}$$

By this formalism, a larger, positive dot product between two vectors indicates better alignment. The reference quantities $\vec{\beta}_{\text{raised}}$ and $\vec{\beta}_{\text{lowered}}$ are overlayed on the reference conformations in *Figure 1d* and the $\vec{\beta}$ system is summarized in *Figure 1e*. Both of these reaction coordinates in conjunction are needed to monitor progress, since rotation away from one reference state does not necessarily imply movement towards the other reference.

*

#### 1.2 System Preparation

Initial apo and holo state structures of the CPD subdomain of TcdA and TcdB were retrieved from the Protein Data Bank entries 3HO6 (TcdA CPD holo), 4R04 (TcdA full-toxin apo), 3PEE (TcdB CPD holo) and 6OQ5 (TcdB full-toxin apo) [1–4]. Co-solutes and ions and were removed, as well as chain B for the dimer crystallizations (3HO6 and 3PEE). The 3PEE structure has 8 N-terminal and 3 C-terminal missing residues and the 3HO6 structure has 4 N-terminal missing residues in the original PDB files, which were modeled using PyMOL's builder function [5]. The 4R04 and 6OQ5 structures contained additional domains not relevant to our study and were truncated to retain only the CPD subdomain. For the several differences in the amino acid sequences between 3PEE and 6OQ5, MODELLER was used to adjust the sequence on structure 6OQ5 to match 3PEE [6]. The missing loop [29-31] in 3HO6 and missing side chains in all structures were modeled using OpenMM's PDBFixer tool [7].

The experimental structures in *Figure 1* provide initial structures for MD simulations of the apo-lowered and holo-raised states of TcdB CPD. The possibly nonphysical apo-raised and holo-lowered states are additionally constructed by removing or adding the IP6 ligand from the two known state structures.

PROPKA predictions for ionizable side chains for the 3PEE structure matched the expectation at physiological pH 7.4, except for residues K600 and K775, which PROPKA predict as neutral [8]. However, IP6 tends to dissociate in holo state simulations for neutral K600 and K775, so standard protonations at physiological pH 7.4 for all side chains were used for all systems, other than the two exceptions listed in *Table 1*. Hydrogens were added on topology creation with the GROMACS utility `pdb2gmx` and flag `-his` to assign all histidine side chains as neutral, with a protonation at $\mathrm{N}(\epsilon)$. Mutated K600G structures were obtained by removing side chain atoms and renaming the residue to glycine. The eight AlphaFold2 structures with lowered probability conformations, af 1-8, were obtained by reducing the depth of the input multiple sequence alignment parameter [9], according to the approach of del Alamo *et al.* [10] (see *Section SI 4*). Structures modelling TcdA CPD (TcdA 541-798) and TcdB CPD (TcdB 544-797) consist of 257 and 254 residues, respectively. For reproducibility, all starting structures, scripts for initialization, simulation and post processing, force field files and simulation parameter files are available for download in a zip file.

#### 1.3 Unbiased MD Simulations

In this study, we performed all molecular dynamics simulations with GROMACS (version 2021.5) [11, 12], interfaced with CUDA (version 11.4.1) and PLUMED (version 2.8.0) [13, 14] plugins. Each initial structure was solvated with TIP3P [15] in a dodecahedron box with at least 1.2 nm to the nearest box edge. Sodium and chloride ions were added for charge neutrality and to replicate a physiological ionic strength of 150 mM, while accounting for screening effects using the SLTCAP method [16]. Temperature control was maintained at 310 K with coupling time 0.1 ps using the V-rescale thermostat [17] and pressure was regulated at 1 bar using the Parrinello-Rahman barostat with coupling time 2 ps [18]. Protein residues used the Amber14sb force field [19], while the tri-protonated IP6 ligand (see *Figure 1c*) was parameterized with GAFF2 for the bonded and Lennard Jones parameters [20]. The atom-centered charges were determined using a two stage RESP fitting protocol according to Vila Verde *et al.* [21–24]: briefly, the RESP fitted charges were determined using a HF/6-31G* optimized geometry in a preliminary 100 ns simulation in a water box. Simulated conformers were extracted at 1 ns intervals, submitted for RESP fitting and the average charge on each atom was used in the final parameter set. A schematic explaining the topology and a plot of atom-charge distributions are shown in *Figure S1*. All runs used a 2 fs time step with the leap-frog integrator, a short-range cutoff of 1 nm, PME for long-range electrostatics and periodic boundary conditions. Bonds with hydrogen atoms were constrained using LINCS [25]. All apo state simulations were subjected to energy minimization using the steepest descent algorithm followed by equilibration with the protein restrained in both NVT and NPT ensembles for 100 ps and 200 ps, respectively. Holo state simulations then

| System | Simulation time | Protonation | System | Simulation time | Protonation |
|---|---|---|---|---|---|
| apo-raised | 10 µs | $K600^{+}/K775^{+}$ | apo-lowered | 10 µs | $K600^{+}/K775^{+}$ |
| holo-raised | 10 µs | $K600^{+}/K775^{+}$ | holo-lowered | 10 µs | $K600^{+}/K775^{+}$ |
| apo-af 1-8 | 500 ns $\times$ 8 | $K600^{+}/K775^{+}$ | apo-K600G | 10 µs | $K600G^{n}/K775^{+}$ |
| apo-E743G | 10 µs | $K600^{+}/K775^{+}$ | apo-K600G/E743G | 16 µs | $K600G^{n}/K775^{+}$ |
| apo<br>2D umbrella | 25 ns $\times$ 4 runs<br>$\times$ 181 windows | $K600^{n}/K775^{n}$ | holo<br>2D umbrella | 25 ns $\times$ 4 runs<br>$\times$ 170 windows | $K600^{+}/K775^{+}$ |
| apo K600G<br>2D umbrella | 25 ns $\times$ 4 runs<br>$\times$ 120 windows | $K600G^{n}/K775^{n}$ | holo K600G<br>2D umbrella | 25 ns $\times$ 4 runs<br>$\times$ 120 windows | $K600G^{n}/K775^{+}$ |
| TcdA apo-raised | 2 µs | $K601^{+}/K776^{+}$ | TcdA apo-lowered | 2 µs | $K601^{+}/K776^{+}$ |
| TcdA holo-raised | 2 µs | $K601^{+}/K776^{+}$ | TcdA holo-lowered | 2 µs | $K601^{+}/K776^{+}$ |

Table 1: Summary of simulated systems and the production run times. Systems were initialized with (holo) or without (apo) IP6 in the allosteric pocket and named systems all consist of the CPD subdomain of TcdB, unless TcdA is specified. $K600^{+}$ and $K775^{+}$ indicate positively protonated lysines. $K600^{n}$, $K775^{n}$ and $K600G^{n}$ indicate a neutral lysine or glycine.

used additional NVT and NPT steps of 100 and 200 ps respectively, with only the ligand restrained to allow the protein side chains to accommodate the ligand inside the allosteric pocket. Production run times are summarized in *Table 1*. For long production runs, the system was re-equilibrated and continued every 500 ns following group internal protocol.

#### 1.4 Umbrella sampling

Umbrella sampling was used to explore the free-energy surface as a function of the $\beta-$flap orientation. This enabled sampling in the conformation transition region and construction of two-dimensional free energy profiles. The reaction coordinate system is based on two collective variables $\xi_1$ and $\xi_2$ which quantify $\beta-$flap orientation, as described in *Section SI 1.1*. Each umbrella window is associated with a restraint point $P_i$ on a grid, where $P$ is a set of 2D points such that, $P = \{(\xi_1, \xi_2)|\xi_1 \in [0, 6], \xi_2 \in [0, 6]\}$. The number of windows differed for each of the constructed FESs, with 181 windows for the native apo state, 170 for the native holo state and 120 for both apo and holo K600G states. The number of windows differed because we in the first FES (apo wt-TcdB CPD) covered the entire grid for $\xi_1 \in [0, 6]$ and $\xi_2 \in [0, 6]$, and later truncated the grid to the accessible regions at 310 K. We introduced additional windows in the transition regions, depending on the distribution overlap. Initial structures for each window in the native states were extracted from the unbiased and the alpha fold simulations, based on the nearest sampled structure to $P_i$. In the holo WT system this approach left a large gap in the transition region (see *Figure 4.b*). Instead we took an iterative approach where we simulated umbrella windows with initial structures from unbiased holo simulations and used these simulations to extract structures for yet unexplored windows. The initial structures for mutant states were extracted from the nearest sample in the corresponding native FES, and then mutated as K600G. All the initial structures and the plumed input files, including the associated $P_i$'s, are available for each window within the downloadable zip file.

For each umbrella window, we conducted 4 simulations, equilibrated by the previously mentioned protocol and simulated in a production run for 25 ns with a harmonic biasing potential using the PLUMED plugin [13, 14], yielding a total 100 ns of sampling per window. The force constant of the umbrella potential was set to $250\ \mathrm{kJ/mol} \cdot \mathrm{nm}^2$ for both dimensions and for all windows. The ensembles were reweighted for the computed FESs, using binless WHAM [26, 27]. The statistical error was estimated using bootstrapped averages. Window data was blocked using 1 ns intervals, from which $N = 200$ bootstraps were constructed, reweighted with WHAM and used to estimate the average and variance ($\sigma$) on

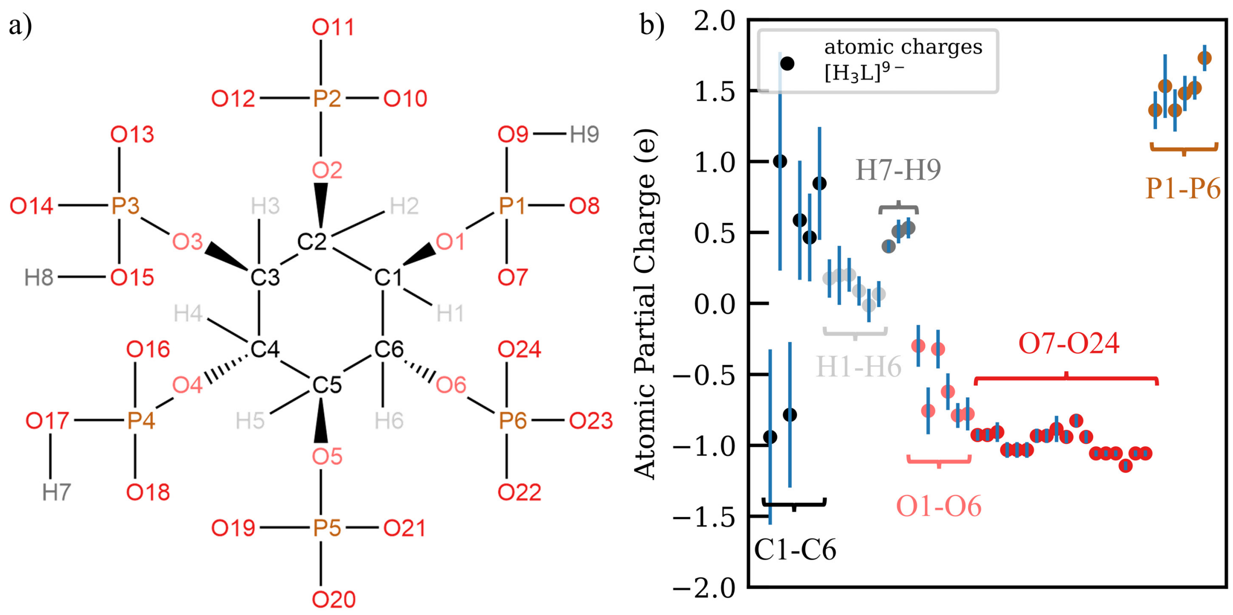


Figure 1: IP6 topology a) Schematic of the IP6 topology and atom naming convention. b) Plot of the atom-centered charges, averaged over 101 structures. The error bar is given as the standard deviation of the charge distribution.

the FES. The lowest free energy paths for each surface was estimated using the A* algorithm (`https://en.wikipedia.org/wiki/A*_search_algorithm`), from which the corresponding simplified 1D free energy profiles were constructed (see *Figures 2.c* and *6.c*). The error was computed as $\epsilon = \sqrt{\frac{\sigma^2}{N}}$.

#### 1.5 IP6 Parameterization

Ligand parameterization prioritized compatibility with the amber14sb protein force field [19]. Therefore, the bonded parameters were derived from GAFF2 (Generalized Amber Force Field) [22] using the `acpype` tool [28] and atomic partial charges were determined with the RESP (Restrained Electrostatic Potential) fitting approach [23, 29]. Due to the sensitivity of electrostatic potentials to molecular conformation, we used a protocol which accounts for multiple conformations [24]. The standard RESP protocol was applied to 101 IP6 structures, extracted from 100 ns of simulation in a water box, and the average value per atom was used to assign atom-centered charges. The topology in *Figure S1a* explains the atom naming convention for re-use of parameters. The plot in *Figure S1b* shows the charge of the atom-centered charges with the standard deviation as the error bar. Atoms in similar environments have similar, but non-identical charges, except for carbon. The standard deviation of RESP charges on all carbons is large since they are buried relative to calculated points on the electrostatic potential, making the calculated charges sensitive to small variations in conformation. This is typical in the RESP charge-fitting approach [29].

The IP6 ligand parameters are available in a downloadable zip file as the residue “IPL” (IP6 in ligand form) under `amber14sb_ip6.ff` as a modified Amber14sb force field [19], adapted for simulations with GROMACS [11].

#### 1.6 Analysis methods

Analysis relied on several python packages including numpy (1.26.4) [30], pandas (2.0.3) [31], MDAnalysis (2.5.0) [32, 33], Biopython's PDBParser (1.76) [34, 35], MDTraj (1.9.9) [36] and plumed (2.8.2.2) [13]. Both PyMOL (2.5.8) [5] and VMD (1.9.4a51) [37] were used for visualization and matplotlib (3.7.2) [38] was used for plotting. In post-processing, trajectories were concatenated and the protein was centered to remove periodic boundary effects using the GROMACS utility `gmx trjconv`. The solvent accessible surface area (SASA) was computed with the GROMACS implementation (`gmx sasa`) of the double cubic lattice method [39]. The set of 114 residues from 254 with $C_\alpha$ root-mean-square fluctuation (RMSF) below 1.5 Å in both the apo-lowered and apo-raised simulations was used for structure alignment throughout the analysis. Residue distances were calculated with the MDAnalysis function `distance_array`, using selection strings based on the anticipated heavy atom distance (either side chain or backbone interactions). The configuration files, helper functions, analysis scripts and conda environments for this project are available for download in a zip file.

#### 1.7 Expression of the Truncated Cysteine Protease Domain

To generate the truncated cysteine protease domain (tCPD) from TcdB (TcdB 543-799 His6), the nucleotide sequence coding for amino acids 543—799 of TcdB were used. This protein excludes the active site substrate, so binding of the allosteric binding site will not induce auto-proteolysis. The wild-type (WT) pET22b-$TcdB_{543-799}$ plasmid was kindly donated by Dr. Matthew Bogyo and Dr. Aimee Shen, Stanford University. The mutant pET22b-$TcdB_{543-799}$-K600G was ordered from GenScript. The plasmids were transformed into Escherichia coli BL21(DE3) by standard techniques. Overnight cultures of transformed BL21(DE3) were diluted 1:100 in 2 L Terrific Broth with ampicillin and grown at 37ºC until an $OD_{600}$ of 0.8—0.9 was reached. IPTG was added and the cultures were grown for 3.5 hr at 30ºC. The cultures were pelleted by centrifugation at 5,000 x g for 30 min at 4ºC (Beckman J2-21, JS5.3). The cell pellets were resuspended in sonication buffer (20 mM phosphate, 100 mM NaCl, 1 mM MgCl2, pH 8). The cell lysates were shaken for 30 min at 4ºC and then sonicated with a probe sonicator (Misonix Sonicator 3000). The cells were centrifuged at 5,000 x g for 30 min at 4ºC, then the supernatant was collected. tCPD variants were purified from the cleared lysate by metal-ion affinity chromatography using Co-NTA resin (ThermoFisher Scientific) at 4ºC. Eluted fractions containing protein were placed on a size exclusion gel filtration column (Superdex 75 HiLoad Prep column) at 4ºC and eluted into the desired buffer.

#### 1.8 Intrinsic Fluorescence Assay

The intrinsic fluorescence assay was performed as described previously with some modifications [3]. This assay was performed to track the position of W761 to indicate whether the tCPD was in the lowered or raised conformation. tCPD variants (tCPD-WT, -K600G) were diluted to 20 M in combination with varied concentrations of IP6 (0.5, 1, 2, 4, 8, 16, 32, 64, 128 M) in 25 mM tris, 100 mM NaCl, 1 mM TCEP, pH 7.5 in a 96 well clear bottom plate (Thermo Scientific, Nunclon Delta Surface). Conditions were repeated four times, and the conditions protein alone (0 M IP6) and buffer alone were included on the plate. The plate was rocked at room temperature for 10 min. The absorbance of each well was measured at 280 nm using a plate reader (Tecan Spark 10M Multimode Plate Reader) to determine the concentration of protein in each well. Next, the fluorescence emission of each sample was then determined by exciting the samples at 295 nm, and the emission spectrum was measured at 1 nm increments from 325 nm to 375 nm. Maximal fluorescence was observed at 335 nm and the correspondent value was used to determine fluorescence. The fluorescence at 335 nm was averaged and the tris buffer background was subtracted. The resultant relative fluorescence units (RFU) were then divided by the average A280 value. First, the RFU was plotted against IP6 concentration (M) in Prism 10 (GraphPad Prism) to see how the tryptophan fluorescence changed in the presence of IP6. Next, the RFU of apo- (0 M IP6) and holo- (128 M IP6) tCPD-WT and tCPD-K600G were plotted in a bar graph with Prism 10 to compare the fluorescence of the proteins in the apo- and holo-forms. A Shapiro-Wilk test of normality was performed, then the groups were compared using Tukey's MCT, ****p 0.0001, n = 4.

#### 1.9 Circular Dichroism

Circular dichroism (CD) data were collected on a Chirascan spectrophotometer (Applied Photophysics). All protein variants were expressed the day of the run to ensure optimal fold and functionality of the proteins. tCPD variants were concentrated to 25 M in 25 mM tris, 100 mM NaF, 1 mM TCEP, pH 7.5 in the absence or presence of 50 M IP6. 50 M IP6 in 25 mM tris, 100 mM NaF, 1 mM TCEP, pH 7.5 and buffer alone were also prepared to ensure the samples had no signal. A blank run was performed with a 0.1 mm quartz cuvette. All runs were set against the blank run. For data collection, 200 L of the samples were placed in the 0.1 mm quartz cuvette. A constant stream of nitrogen gas at 25 psi was applied to the lamp. The bandwidth was set to 0.5 nm with a 0.5 s collection time per data point. Scanning was performed from 180 – 260 nm. Fifteen scans were collected and averaged followed by a Savitsky-Golay smoothing function with a window size of 3. The secondary structure analysis was performed using the BeStSel software [40–42].

#### 2 Additional figures

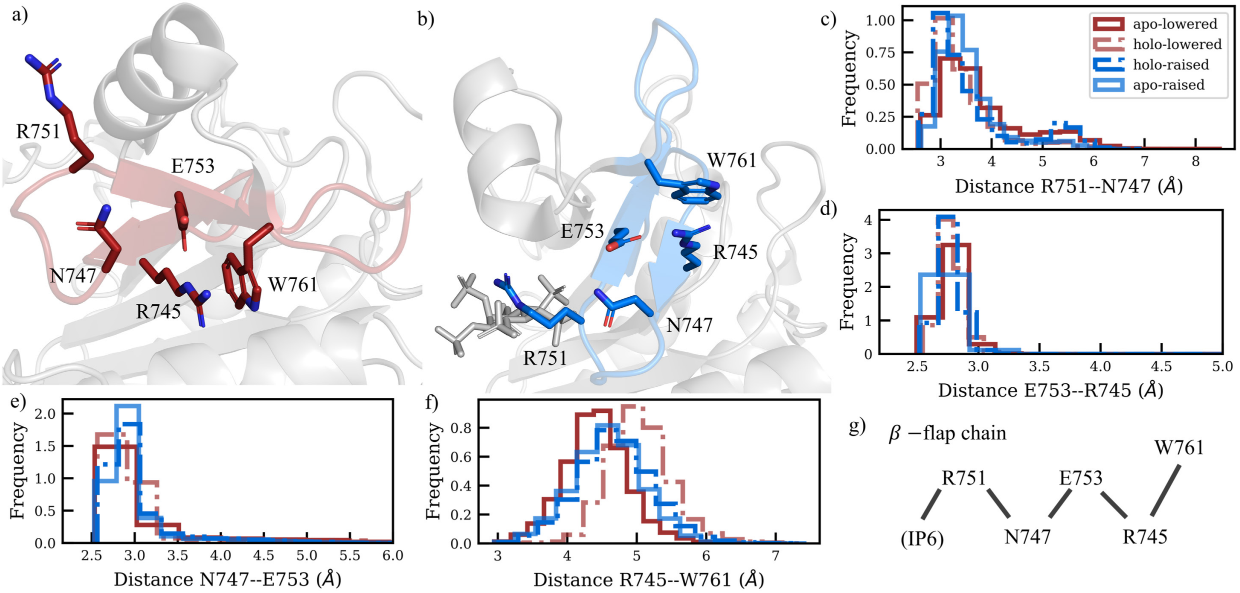


Figure 2: The $\beta-$flap chain proposed by Shen *et al.* is independent of the conformation [3]. a) The $\beta-$flap chain in the reference lowered conformation (PDB entry: 6OQ5 [4]). b) The $\beta-$flap chain in the reference raised conformation (PDB entry: 3PEE [3]). Side chains in the $\beta-$flap chain are shown as colored sticks in a) and b) and IP6 is shown as gray sticks in b). c-f) Distance histograms for residue pairs in the $\beta-$flap chain. g) Summary of $\beta-$flap chain interactions

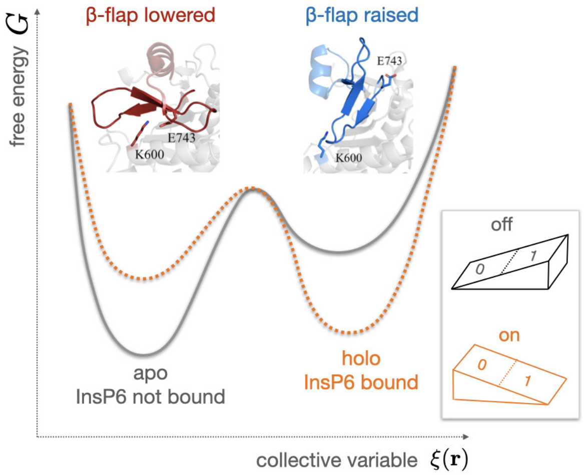


Figure 3: Allosteric regulation modelled as a bistable switch. The TcdB CPD exhibits a bistable free-energy profile with two distinct minima — one for the $\beta$-flap lowered conformation and one for the $\beta$-flap raised conformation. Binding of the allosteric modulator IP6 switches the relative stability of these minima: the apo state stabilizes the lowered conformation, while the holo state stabilizes the raised conformation. In this sense the free-energy profile functions like a rocker switch (inset).

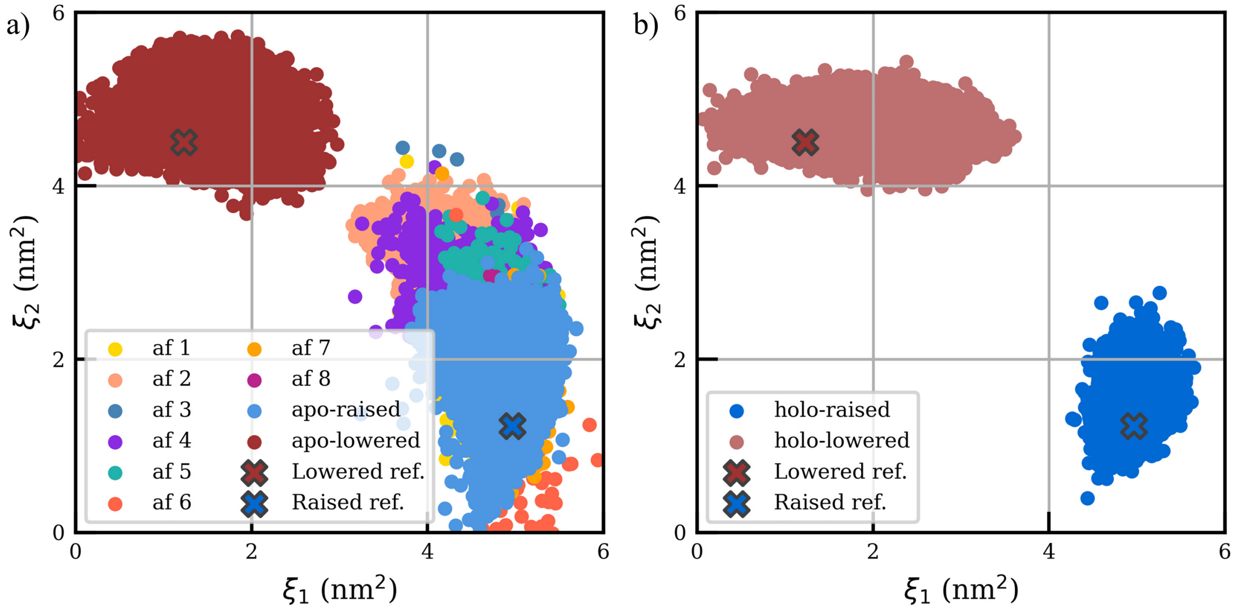


Figure 4: Unbiased simulations do not transition between conformational states on a microsecond timescale. The sampled region in the $\xi_1$, $\xi_2$ reaction coordinate space for a) unbiased apo states and b) unbiased holo states. Sampling from apo-state AlphaFold2 conformations are shown in a), where ”af” abbreviates numbered AlphaFold2 structures [9, 10]. Points are plotted at 1 ns intervals. Positions of references structures in $\xi_1$, $\xi_2$ space are indicated with ”X”s.

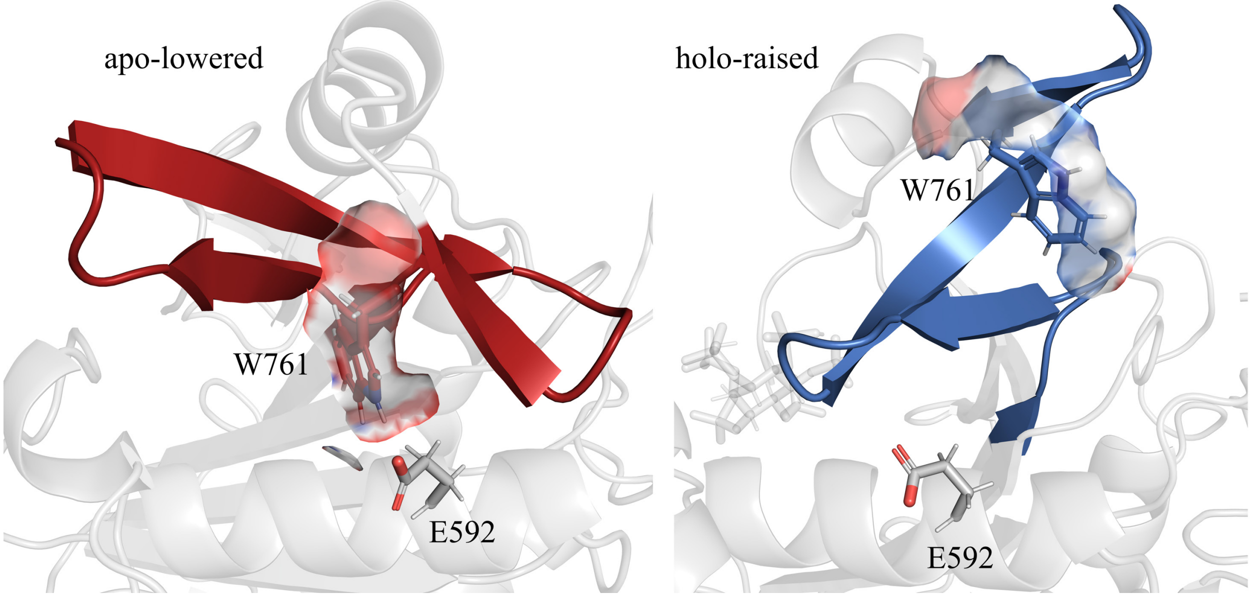


Figure 5: Solvent exposure of W761. The SASA of W761 and proximity to E592 in the a) lowered conformation (extracted from unbiased apo-lowered MD simulation) and b) raised conformation (extracted from unbiased holo-raised MD simulation).

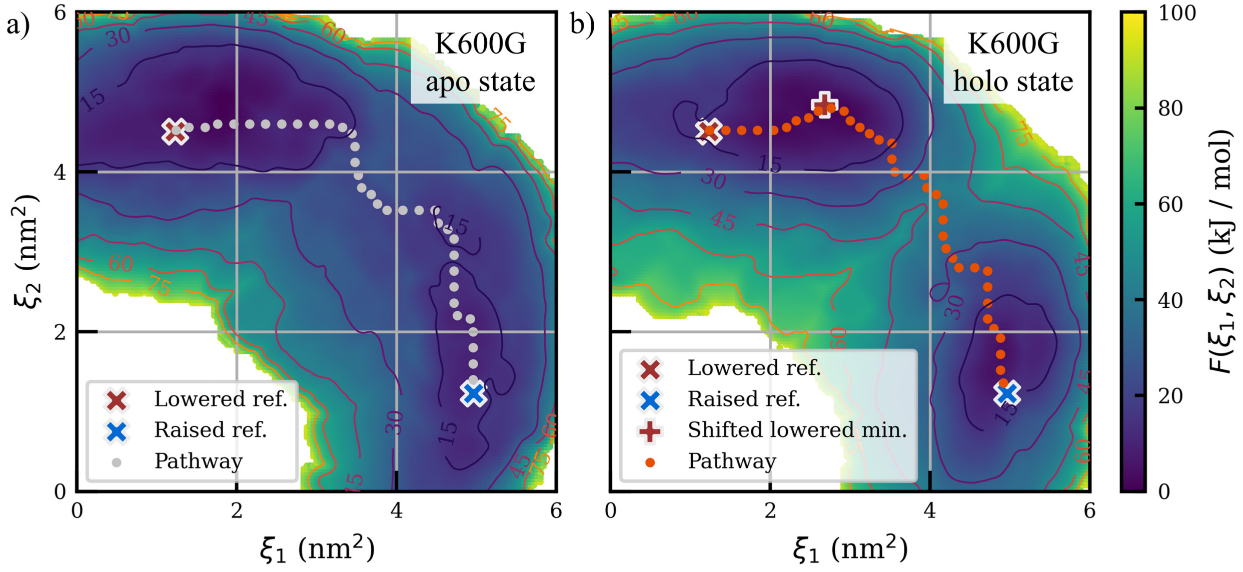


Figure 6: Computed 2D free-energy surfaces for K600G in the $\xi_1$, $\xi_2$ reaction coordinate space for a) apo and b) holo states. Positions of reference structures in $\xi_1$, $\xi_2$ space are indicated with "X"s and in b) the "+" indicates the free-energy minimum of the lowered conformation relative to the reference lowered conformation.

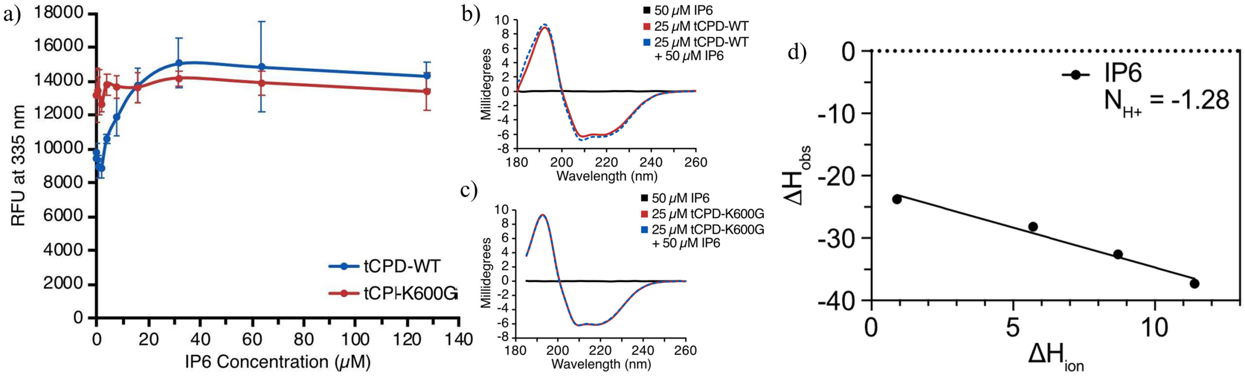


Figure 7: Experimental mutagenesis and net proton change upon IP6 binding to tCPD a) Effect of the point mutation K600G on intrinsic tryptophan fluorescence of tCPD. Fluorescence emission of tCPD was measured at 335 nm after excitation at 295 nm in the absence (apo-tCPD, 0 $\mu$M IP6) or presence (holo-tCPD) of varied concentrations of IP6. Each fluorescence measurement was controlled for protein concentration (absorbance at 280 nm). Mean ± SD, n = 4. b,c) Secondary structure of b) holo-tCPD-WT and c) holo-tCPD-K600G as determined by circular dichroism (CD) to confirm the K600G point mutation did not modify the overall structure of tCPD. d) Enthalpy change of binding, $\Delta H_{obs}$, of tCPD to IP6 as a function of ionization enthalpy, $\Delta H_{ion}$, of the respective buffers at pH 7.5. The solid lines represent least-square fitting of these points. The change in number of bound protons per binding interaction ($N_{H+}$) is given by the slope. Figure d) is reproduced here with permission from Cummer *et al.* [43].

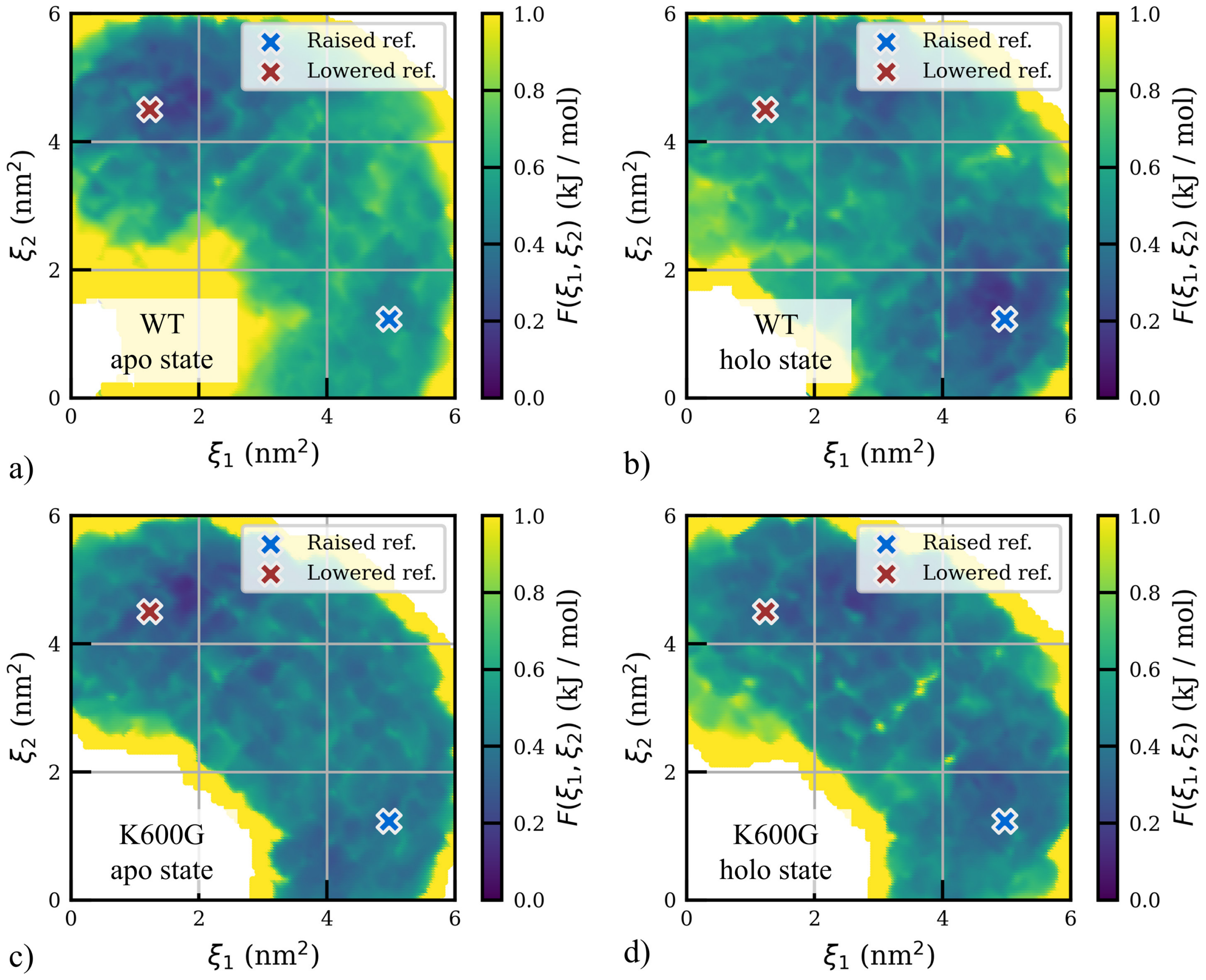


Figure 8: Statistical uncertainty estimates in the free energy surfaces. Estimates are based on bootstrapping for reweighted umbrella sampling data. Positions of references structures in $\xi_1$, $\xi_2$ space are indicated with "X"s.

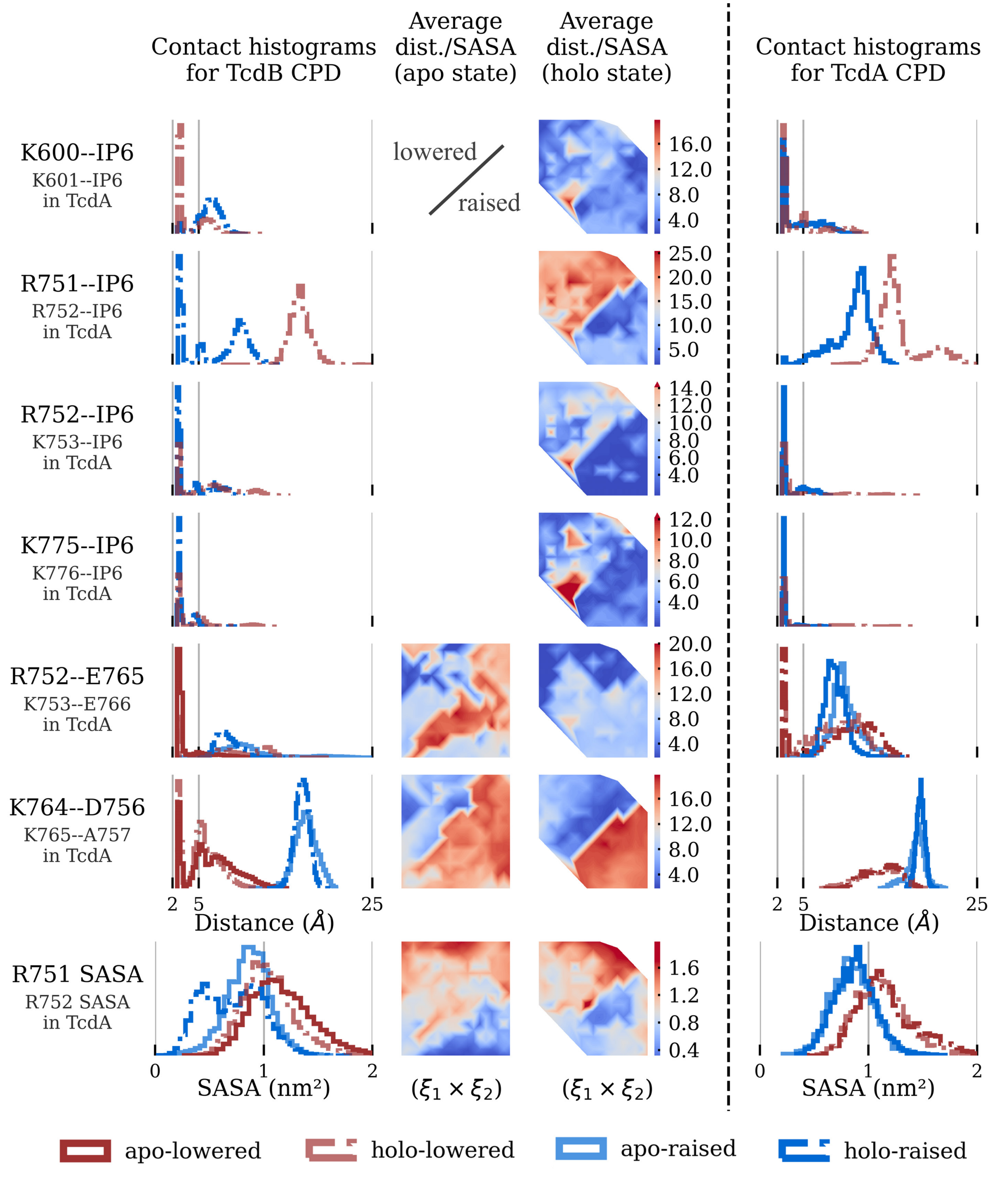


Figure 9: Additional pairwise residue interactions in the proposed interaction network. *Left column:* histograms of residue pair distances from four (apo/holo + lowered/raised) simulated states. *Two middle columns:* average residue distance in each umbrella window, for apo and holo state. These surfaces use the same reaction coordinate space ($\xi_1, \xi_2$) as in the free-energy surface (see *Figure 2*), in which the upper left and lower right correspond to the lowered and raised conformations, respectively. *Right column:* histograms of residue pair distances for each analogous interaction in TcdA CPD.

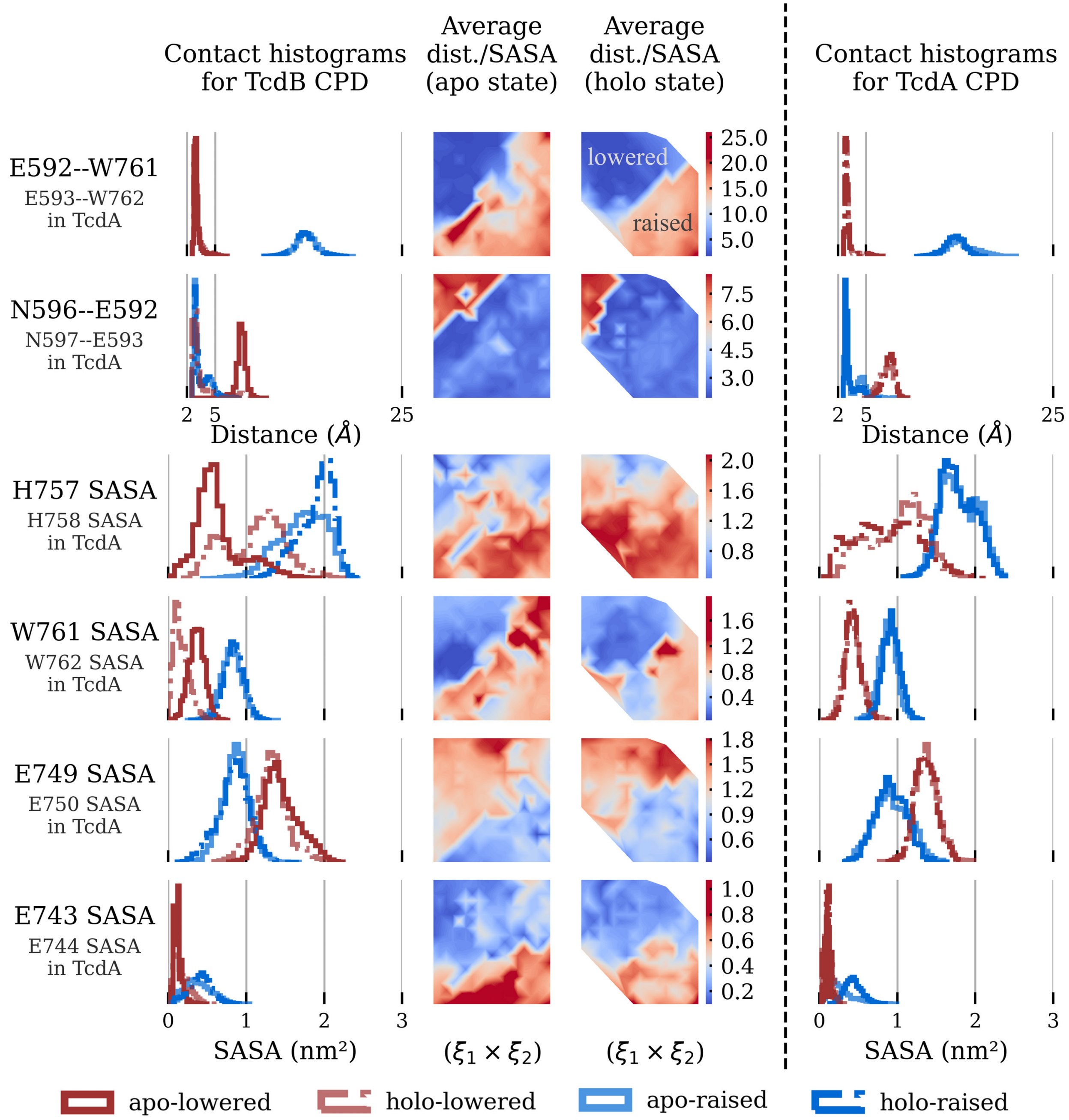


Figure 10: Pairwise residue interactions in the proposed interaction network. *Left column:* histograms of residue pair distances from four (apo/holo + lowered/raised) simulated states. *Two middle columns:* average residue distance in each umbrella window, for apo and holo state. These surfaces use the same reaction coordinate space ($\xi_1, \xi_2$) as in the free-energy surface (see *Figure 2*), in which the upper left and lower right correspond to the lowered and raised conformations, respectively. *Right column:* histograms of residue pair distances for each analogous interaction in TcdA CPD.

### 3 Interpreting the Ligand-dependence of the Barrier Height

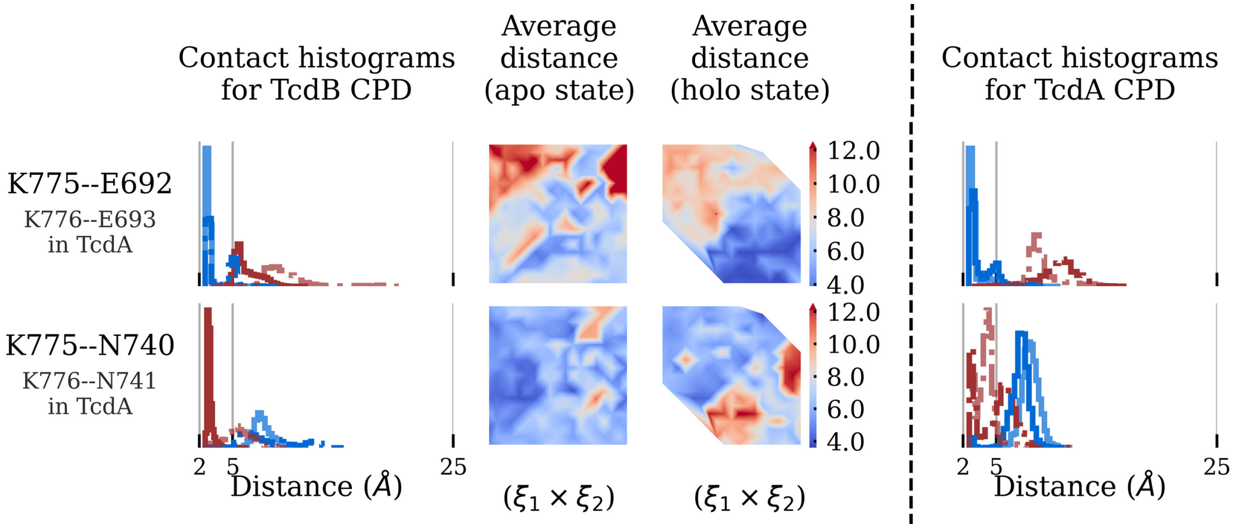


Figure 11: Additional pairwise residue interactions impacted by K775 protonation. *Left column:* histograms of residue pair distances from four (apo/holo + lowered/raised) simulated states. *Two middle columns:* average residue distance in each umbrella window, for apo and holo state. These surfaces use the same reaction coordinate space ($\xi_1, \xi_2$) as in the free-energy surface (see *Figure 2*), in which the upper left and lower right correspond to the lowered and raised conformations, respectively. *Right column:* histograms of residue pair distances for each analogous interaction in TcdA CPD.

A noticeable feature of the free energy surfaces is a much higher barrier for the holo states, than the apo states. This feature persists in both the native state and K600G (*Figures 2, 6.c, S6*). In our simulations, apo- and holo-states differ in three aspects: presence of IP6, protonation of K775 and protonation of K600. While the biased apo structures have neutral lysines 600 and 775 for the wild type and neutral lysine 775 in K600G, the holo structures are protonated at these lysines. The most straightforward explanation for the higher transition barrier in the holo state is that IP6 sterically hinders the transition between lowered and raised conformation. We nonetheless explored the influence of the protonation states on the height of this barrier.

The simulations of the K600G mutant show that even though K600 is critical in stabilizing the lowered conformation, it has little influence on the barrier height. Because glycine does not act as a base, the protonation of residue 600 in the K600G mutant is the same in the apo and the holo state, yet we observe the same increased barrier in its holo state as in the wild type. Additionally, residue pairs involving K600 in the wild type show binary distance distributions and a sharp demarcation between short and large distances in the collective variable space regardless of whether we simulated the apo or the holo state. Thus, protonation of K600 likely does not cause the increase in barrier height.

By contrast residue pairs involving K775 exhibit conformation dependence exclusively in the holo state, i.e. when K775 is protonated. In our interaction network these are the hydrogen bond K775–N740, which is formed in the lowered conformation and the salt bridge K775–E692, which is formed in the raised conformation (*Figure S11*). This implies that only when K775 is protonated there is a sharp transition barrier for breaking or forming these two interactions. Thus, protonation of K775 likely contributes to the increase in barrier height of the holo state relative to the apo state.

An increased transition barrier in the holo-state might contribute to the overall allosteric effect. If the barrier increases after $\beta$-flap rotation, e.g. possibly because only then the proton can be transferred from IP6 to K775, then the system is kinetically trapped in the raised conformation. A comprehensive picture is emerging in which CPD allostery operates through a concerted mechanism of ligand binding, proton transfer, and structural rearrangements with corresponding changes in the interaction network. To fully characterize the transition path at the barrier, one should utilize constant-pH simulations [44] in combination with transition path finding algorithms [45].

### 4 AlphaFold2 structures

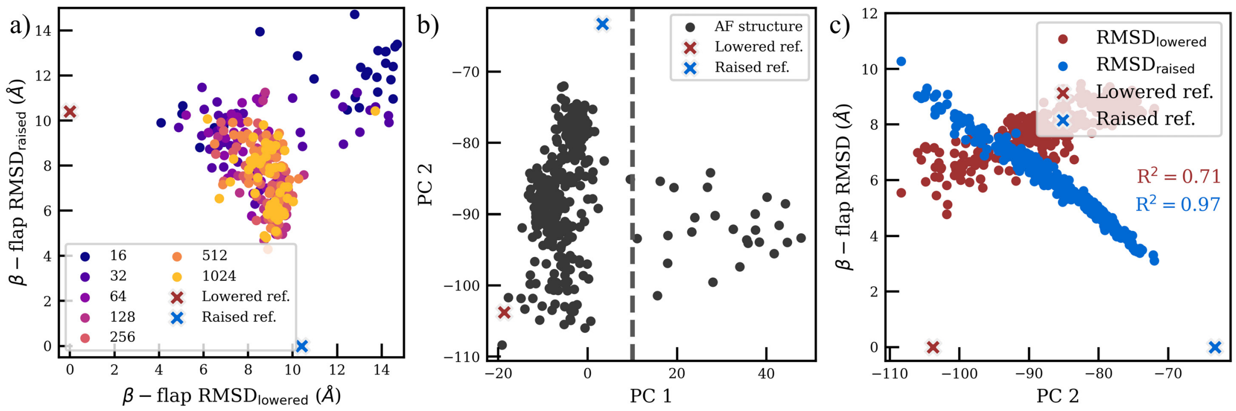


Figure 12: AlphaFold2 structures. a) The $\beta-$flap RMSD for both reference conformations, with MSA depth indicated by color. The default MSA depth is 5012. b) The first two principal components (PC) based on PC analysis (PCA) of the $C_\alpha$'s in the $\beta-$flap from 350 AlphaFold2 structures with variable MSA depth. c) The correlation between PC2 and the $\beta-$flap RMSD against both reference structures. AlphaFold2 structures are indicated with "o"s and reference structures are indicated with "X"s in all subplots.

AlphaFold2 makes highly accurate protein structure predictions directly from amino acid sequence and, in the case of the TcdB CPD, predicts a structure with high fidelity to the raised conformation [9]. To encourage conformational variety, the standard AlphaFold2 model was modified according to the approach of del Alamo *et al.* [10], primarily by lowering the multiple sequence alignment (MSA) depth from the default value of 5012. Fifty structures were predicted for each of seven MSA depths, yielding 350 potential initial conformations for MD simulations. Some structures may have an intermediate $\beta-$flap conformation, based on the $\beta-$flap RSMDs shown in *Figure S12a*. Structures which are closer to the closed conformation tend to have a shallow MSA depth, in contrast to the standard prediction. This supports an alternative use case for AlphaFold2, to produce high energy, intermediate conformations [10]. Importantly, a low MSA depth is also more likely to produce an outlier structure, where the $\beta-$flap RMSD is high regardless of the reference structure, so it is important to screen structures.

A principal component analysis (PCA) was used to sort out misfolded structures and to check the correlation with the allosteric transition. The PCA is based on the $C_\alpha$ positions of the $\beta-$flap residues. *Figure S12b* shows the first principal component (PC1) sorts out the outliers (PC1 $> 10$), while the second principal component (PC2) effectively captures the transition between the two reference structures. The correlation between PC2 and the $\beta-$flap RMSDs in *Figure S12c* verifies that structures interpolate from the lowered conformation to the raised conformation as PC2 increases. Ten non-outlier structures were randomly selected along PC2, checked for obvious abnormalities and used as initial structures in unbiased MD simulations for 500 ns each. *Figure S4a* shows that eight of the simulated AlphaFold2 structures sample reasonable spaces in the reaction coordinate space and even improve the sampling relative to experimental structures. However, no structure was able to surpass the barrier and enter the lowered conformation minimum.

It's worth noting that AlphaFold2 is expected to predict the lowered conformation since it is associated with the apo state, i.e. the model *should* predict the ligand-free conformation. Instead, it predicts structures closer to the raised conformation minimum, associated with the holo state crystallization, than the lowered conformation minimum. We explain this behavior as an artefact of the data set included in the training set of AlphaFold2, which includes the raised conformation and excludes the lowered conformation.
