## Supplementary material for "*Clostridioides difficile* Toxins Unhinged: Allosterically Switchable Network Orients *β*-flap": Scripts, Structures etc.: AF_PCA.py

```
import sys
import os
import numpy as np
import matplotlib.pyplot as plt
from Bio.PDB import *
# Hamelryck, T., Manderick, B. (2003) PDB parser and structure class implemented
# in Python. Bioinformatics 19: 2308–2310
import MDAnalysis as mda
import shutil
sys.path.insert(0, "/home/lf1071fu/project_b3/ProjectB3")
from tools import utils, traj_funcs
import config.settings as cf

def main(argv):

    # Set up specifications for the system and initialize some general variables
    template = False
    path = f"{ cf.path_head }/alphafold/MSA_depths"

    global bound, apo, msa_depths, alignment, beta_align, central
    bound, apo = "3pee", "6oq5"
    msa_depths = [32, 64, 128, 256, 512, 1024, 5120] # excluding msa depth of 16
    alignment = {bound : range(8,252), apo : range(552, 796)}
    beta_align = {bound : range(198,232), apo : range(743, 777)}
    central = np.array(list(range(8,195)) + list(range(234,252)))

    files = {bound : f"{ path }/../structures/3pee_A.pdb",
             apo : f"{ path }/../structures/6oq5_cpd.pdb"}
    af_keys = [bound, apo]

    files, af_keys = add_models(files, af_keys, path, template)

    np_files = ["coord_data.npy", "eigenvals.npy", "eigenvecs.npy",
                "rmsd_holo.npy", "rmsd_apo.npy",]
    np_files = [path + "/" + f for f in np_files]

    if all(list(map(lambda x : os.path.exists(x), np_files))):

        centered_strs = np.load(np_files[0], allow_pickle=True)
        eigvals = np.load(np_files[1], allow_pickle=True)
        eigvecs = np.load(np_files[2], allow_pickle=True)
        rmsd_holo = np.load(np_files[3], allow_pickle=True)
        rmsd_apo = np.load(np_files[4], allow_pickle=True)

    else:

        strs, rmsd_holo, rmsd_apo = get_coord_data(files)

        # Apply row centering for a centered data matrix
        # TO DO: include centering for the most stable residue positions
        centered_strs = strs[198*3:232*3, :] - strs[198*3:232*3, :].mean()

        covar = np.cov(centered_strs)
        eigvals, eigvecs = np.linalg.eig(covar)

        np.save(np_files[0], centered_strs)
        np.save(np_files[1], eigvals)
        np.save(np_files[2], eigvecs)
        np.save(np_files[3], rmsd_holo)
        np.save(np_files[4], rmsd_apo)

    # plot_eigvals(eigvals, f"{ path }/figures_holo_algn")
    # plot_pca_correl(centered_strs, eigvecs, [1,2],
    #                 f"{ path }/figures_holo_algn")
    # selection = plot_pca_correl(centered_strs, eigvecs, [1,2],
    #                             f"{ path }/af_subset", select=True)

    # Get the first and second PC of each structure
    pc1 = np.dot(centered_strs.T, eigvecs[:,0])
    pc2 = np.dot(centered_strs.T, eigvecs[:,1])
    plot_pca12(centered_strs, eigvecs, [1,2], path)

    # non_outliers = []
    # print(pc1)
    # for n, p in zip(af_keys[2:], pc1[2:]):
    #     if p > -12:
    #         non_outliers.append(n)
    # print(non_outliers)
    # np.save(f"{ path }/non_outliers.npy", non_outliers)

    # Filter out outlier structures using PC 1
    plot_rmsd_pc(pc1, pc2, rmsd_holo, rmsd_apo)
    # # print(len(pc2_r))
    # # af_keys = np.array(af_keys)
    # # min_pc1, max_pc1 = af_keys[np.argmin(pc1_r)], af_keys[np.argmax(pc1_r)]
    # # min_pc2, max_pc2 = af_keys[np.argmin(pc2_r)], af_keys[np.argmax(pc2_r)]
    # # print(f"PC1 min and max: { min_pc1 }, { max_pc1 }")
    # # print(f"PC2 min and max: { min_pc2 }, { max_pc2 }")
    # # get_strs_pc2(pc1, pc2, af_keys)
    # select_files = np.array(list(files.values()))[selection]
    # for f in select_files:
    #     shutil.copy(f, f"{ path }/af_subset")

    return None

def add_models(files, af_keys, path, template):
    # Add model data from .pdb files
    prefix = f"{ path }/no_template_structures"

    for depth in msa_depths:
        for i in range(50):

            if template:
                file = f"{ path }/apo_template_structures/{ i }_{ depth }.pdb"
            else:
                file = f"{ path }/no_template_structures/{ i }_{ depth }.pdb"

            key = f"m{ i }d{ depth }"
            files[key] = file
            af_keys.append(key)

    return files, af_keys

def get_coord_data(files):

    str_count = len(files)
    # Atoms per res is 1 for alpha carbon based analysis
    atoms_per_res = 1
    dof_count = len(alignment[bound]) * atoms_per_res * 3
    strs = np.zeros((str_count, dof_count))
    str = 0
    rmsd_holo, rmsd_apo = [], []

    for key, file in files.items():

        # Parse the pdb file and store as a Bio.PDB.Chain object
        parser = PDBParser(PERMISSIVE=1, QUIET=True)
        struct = parser.get_structure(id, file)[0]["A"]

        # Align to the central alignment residues
        if key != bound:

            if key == apo:
                ref_atoms = get_backbone_atoms(holo_struct, alignment[bound])
                moving_atoms = get_backbone_atoms(struct, alignment[apo])
            else:
                align = central
                ref_atoms = get_backbone_atoms(holo_struct, alignment[bound])
                moving_atoms = get_backbone_atoms(struct, alignment[bound])

            super_imposer = Superimposer()
            super_imposer.set_atoms(ref_atoms, moving_atoms)
            super_imposer.apply(struct.get_atoms())

        if key == apo:

            apo_struct = struct
            apo_beta_ca = get_backbone_atoms(struct, beta_align[apo])

        elif key == bound:

            holo_struct = struct
            holo_beta_ca = get_backbone_atoms(struct, beta_align[bound])

        # Get the RMSD of the beta flap to the experimental structures
        if key not in [bound, apo]:
            af_beta_ca = get_backbone_atoms(struct, beta_align[bound])
            rmsd_holo.append(calc_rmsd(af_beta_ca, holo_beta_ca))
            rmsd_apo.append(calc_rmsd(af_beta_ca, apo_beta_ca))

        # Get the selected residues, get the alpha-carbon atoms and get the atom
        # coordinates, then add to a np array.
        atom_count = 0

        for res in struct.get_residues():

            res_id = res.get_id()[1]
            if key == apo:
                select_atoms = alignment[apo]
            else:
                select_atoms = alignment[bound]

            if res_id in select_atoms:

                # Adapt if all the backbone atoms are desired
                for atom in [res["CA"]]:
                    ind = 3 * atom_count
                    strs[str, ind:(ind + 3)] = atom.get_coord()
                    atom_count += 1

        str += 1

    # Exlcude the crystal data and msa depth 16 from analysis and use the
    # transpose so that conformations are the columns and DoFs are the rows.
    strs = strs.T

    return strs, np.array(rmsd_holo), np.array(rmsd_apo)

def get_backbone_atoms(struct, atoms_to_be_aligned, full_backbone=False):
    """Get a list of the backbone atoms from a subset of residues.

    Parameters
    ----------
    struct : Bio.PDB.Chain
        A protein structure from a single chain, loaded from a pdb file.
    atoms_to_be_aligned : (int) iter
        An iterable to select the subset of residues.

    Returns
    -------
    atoms : (Bio.PDB.Atom) list
        An ordered list of backbone atoms from the selected residues.

    """
    atoms = []

    for res in struct.get_residues():
        res_id = res.get_id()[1]
        if res_id in atoms_to_be_aligned:
            if full_backbone:
                atoms += [res["N"], res["CA"], res["C"], res["O"]]
            else:
                atoms += [res["CA"]]

    return atoms

def get_strs_pc2(pc1, pc2, af_keys):
    """
    """
    pc2_r = pc2[(pc1 < 5)]
    s = []
    min = -110
    for i in [-105, -100, -95, -90, -85, -80, -75, -70]:
        max = i
        filter = pc2_r[(min < pc2_r) & (pc2_r < max)]
        val = np.random.choice(filter)
        s.append(af_keys[np.where(val == pc2)[0][0]])

    print(s)

    return None

def calc_rmsd(atoms, ref_atoms):
    """Calculate the RMSD from the list of backbone atoms.

    Parameters
    ----------
    atoms : (Bio.PDB.Atom) list
        The backbone CA atoms of the included residues.
    ref_atoms : (Bio.PDB.Atom) list
        The backbone CA atoms of the included residues.

    Returns
    -------
    rmsd : float
        The RMSD of the residue.

    """
    dists_sq = []
    n_atoms = len(atoms)

    for atom, ref_atom in zip(atoms, ref_atoms):
        dists_sq.append( np.linalg.norm((atom - ref_atom)) ** 2 )

    rmsd = np.sqrt( 1/n_atoms * sum(dists_sq) )

    return rmsd

def plot_eigvals(eigvals, path):
    """Make a plot of the first eigenvals from PC analysis.

    The relative contribution to the overall variance by each principle
    component is shown for the first 15 PCs. This should simply demonstrate the
    number of relevant PCs in the dimensionality reduction.

    Parameters
    ----------
    eigvals : np.ndarray
        A 1D array of the eigenvalues from PCA of the covariance matrix.
    path : str
        The path to the working directory. The plot will be saved to the
        figures sub directory.

    Returns
    -------
    None.

    """
    fig, ax = plt.subplots(constrained_layout=True)
    filled_marker_style = dict(marker='o', markersize=10, linestyle="-", lw=3,
                               markeredgecolor='#A31130')
    normalize = lambda x : x / sum(eigvals) * 100
    ax.plot(np.arange(1,16), normalize(eigvals[:15]), color="#FF6666",
            **filled_marker_style)

    # Plot settings
    ax.tick_params(axis='y', labelsize=16, direction='in', width=2, \
                    length=5, pad=10)
    ax.tick_params(axis='x', labelsize=16, direction='in', width=2, \
                    length=5, pad=10)
    for i in ["top","bottom","left","right"]:
        ax.spines[i].set_linewidth(2)
    ax.grid(True)
    ax.set_xlabel(r"Principle component", labelpad=5, fontsize=16)
    ax.set_ylabel(r"Contribution to overall variance (%)", \
                    labelpad=5, fontsize=16)

    # Needs a "figures" subdirectory within the path
    plt.savefig(f"{path}/pca_eigenvals.png")
    plt.close()

    return None

def plot_pca12(centered_strs, eigvecs, pcs, path, select=False):
    """Make a plot of the first two principle components.

    The experimental structures are projected onto the first 2 PCs
    for comparison.

    """
    fig, ax = plt.subplots(constrained_layout=True, figsize=(2.5,2.5))
    # Format relevant data for the first 2 PCs
    pc_a, pc_b = pcs
    pca1 = np.dot(centered_strs.T, eigvecs[:,pc_a - 1])
    pca2 = np.dot(centered_strs.T, eigvecs[:,pc_b - 1])
    holo_pca1, holo_pca2 = np.dot(centered_strs[:,0], eigvecs[:,pc_a - 1]), \
                           np.dot(centered_strs[:,0], eigvecs[:,pc_b - 1])
    apo_pca1, apo_pca2 = np.dot(centered_strs[:,1], eigvecs[:,pc_a - 1]), \
                         np.dot(centered_strs[:,1], eigvecs[:,pc_b - 1])

    # Plot the PC1 & 2 values of all AF structures and also the experimental
    # values.
    ax.axvline(10, ls="--", lw=2, color="#585858")
    ax.scatter(pca1, pca2, label="AF structure", color="#383838", s=10)
    ax.scatter(apo_pca1, apo_pca2, label="Lowered ref.", marker="X", 
            color=cf.closed_color, edgecolors="#ededed", s=50, lw=1)
    ax.scatter(holo_pca1, holo_pca2, label="Raised ref.", marker="X", 
            color=cf.open_color, edgecolors="#ededed", s=50, lw=1)

    # Plot settings
    ax.tick_params(axis='both', which='major', pad=3)
    ax.grid(False)
    ax.set_xlabel(f"PC { pc_a }", labelpad=3, fontsize=8)
    ax.set_ylabel(f"PC { pc_b }", labelpad=3, fontsize=8)
    plt.legend(fontsize=6)

    utils.save_figure(fig, 
        f"{ cf.figure_head }/alphafold/pc_1_2.png")
    plt.close()

def plot_pca_correl(centered_strs, eigvecs, pcs, path, select=False):
    """Make a plot of the first two principle components.

    The experimental structures are projected onto the first 2 PCs
    for comparison.

    """
    fig, ax = plt.subplots(constrained_layout=True)
    filled_marker_style = dict(marker='o', linestyle="")
    # Format relevant data for the first 2 PCs
    pc_a, pc_b = pcs
    pca1 = np.dot(centered_strs.T, eigvecs[:,pc_a - 1])
    pca2 = np.dot(centered_strs.T, eigvecs[:,pc_b - 1])
    holo_pca1, holo_pca2 = np.dot(centered_strs[:,0], eigvecs[:,pc_a - 1]), \
                           np.dot(centered_strs[:,0], eigvecs[:,pc_b - 1])
    apo_pca1, apo_pca2 = np.dot(centered_strs[:,1], eigvecs[:,pc_a - 1]), \
                         np.dot(centered_strs[:,1], eigvecs[:,pc_b - 1])
    selection = None

    # Plot the PC1 & 2 values of all AF structures and also the experimental
    # values.
    if not select:
        ax.axvline(-5, ls="--", lw=2, color="#585858")
    ax.plot(pca1, pca2, label="AF structure", color="#FF6666",
            markeredgecolor="#A31130", markersize=7.5, **filled_marker_style)
    ax.plot(holo_pca1, holo_pca2, label="holo state", color="#BA59EE",
            markeredgecolor="#4D2563", markersize=10, **filled_marker_style)
    ax.plot(apo_pca1, apo_pca2, label="apo state", color="#5BE49F",
            markeredgecolor="#36875E", markersize=10, **filled_marker_style)

    if select:

        # Set axis lims
        plt.xlim(ax.get_xlim())
        plt.ylim(ax.get_ylim())

        # Plot the line connecting the experimental points
        slope = (apo_pca2 - holo_pca2) / (apo_pca1 - holo_pca1)
        y_int = apo_pca2 - slope * apo_pca1
        x_vals = np.array(ax.get_xlim())
        #y_vals = np.array(ax.get_ylim())
        #x_vals = (y_vals - y_int) / slope
        y_vals = slope * x_vals + y_int
        plt.plot(x_vals, y_vals + 3, '--')
        plt.plot(x_vals, y_vals - 3, '--')

        cond1 = pca2 < slope * pca1 + y_int + 3
        cond2 = pca2 > slope * pca1 + y_int - 3
        inds = np.arange(len(pca1))
        selection = np.random.choice(inds[(cond1) & (cond2)][2:], size=10)
        ax.plot(pca1[selection], pca2[selection], label="AF selection",
                markeredgecolor="#A31130", markersize=7.5, **filled_marker_style)


    # Plot settings
    ax.tick_params(axis='y', labelsize=16, direction='in', width=2, \
                    length=5, pad=10)
    ax.tick_params(axis='x', labelsize=16, direction='in', width=2, \
                    length=5, pad=10)
    for i in ["top","bottom","left","right"]:
        ax.spines[i].set_linewidth(2)
    ax.grid(True)
    ax.set_xlabel(f"PC { pc_a }", labelpad=5, fontsize=16)
    ax.set_ylabel(f"PC { pc_b }", labelpad=5, fontsize=16)
    plt.legend(fontsize=12)

    # Needs a "figures" subdirectory within the path
    plt.savefig(f"{ path }/structure_select.png")
    plt.close()

    return selection

def plot_rmsd_pc(pc1, pc2, rmsd_holo, rmsd_apo):
    """
    """
    pc2_f = pc2[(pc1 < 10)]
    rmsd_holo_f = rmsd_holo[(pc1[2:] < 10)]
    rmsd_apo_f = rmsd_apo[(pc1[2:] < 10)]

    fig, ax1 = plt.subplots(constrained_layout=True, figsize=(2.5,2.5))

    ax1.scatter(pc2_f[2:], rmsd_apo_f, label=r"RMSD$_{\mathrm{lowered}}$", 
                color=cf.closed_color, s=10, marker='o')
    r_1 = np.round(np.corrcoef(rmsd_apo_f, pc2_f[2:])[0,1] ** 2, 2)

    ax1.scatter(pc2_f[2:], rmsd_holo_f, label=r"RMSD$_{\mathrm{raised}}$", 
                color=cf.open_color, s=10, marker='o')
    r_2 = np.round(np.corrcoef(rmsd_holo_f, pc2_f[2:])[0,1] ** 2, 2)

    # Add in reference positions
    ref1 = ax1.scatter(pc2[1], 0, edgecolors="#ededed", s=50, lw=1,
                label="Lowered ref.", marker="X", color=cf.closed_color)
    ref2 = ax1.scatter(pc2[0], 0, edgecolors="#ededed", s=50, lw=1,
                label="Raised ref.", marker="X", color=cf.open_color)
              
    # Add in the coefficients of determination
    plt.text(.97, .5, r"R$^2 = $"+str(r_1), ha="right", va="bottom",
             color=cf.closed_color, transform=ax1.transAxes, fontsize=8)
    plt.text(.97, .42, r"R$^2 = $"+str(r_2), ha="right", va="bottom",
             color=cf.open_color, transform=ax1.transAxes, fontsize=8)

    # Plot settings
    ax1.tick_params(axis='both', which='major', pad=3)
    ax1.grid(False)
    ax1.set_xlabel(f"PC 2", labelpad=3, fontsize=8)
    ax1.set_ylabel(r"$\beta-$flap RMSD ($\AA$)", labelpad=3, fontsize=8)
    ax1.set_ylim([-.5,12])

    ax1.legend(loc="upper right", fontsize=8)

    # Needs a "figures" subdirectory within the path
    utils.save_figure(fig, 
        f"{ cf.figure_head }/alphafold/pc2_vs_rmsd.png")
    plt.show()
    plt.close()

    return None

if __name__ == '__main__':
    main(sys.argv)
```
