## Supplementary material for "*Clostridioides difficile* Toxins Unhinged: Allosterically Switchable Network Orients *β*-flap": Scripts, Structures etc.: alphafold.py

```
# -*- coding: utf-8 -*-
"""AlphaFold.ipynb

Automatically generated by Colaboratory.

Original file is located at
    https://colab.research.google.com/github/deepmind/alphafold/blob/main/notebooks/AlphaFold.ipynb

# AlphaFold Colab

This Colab notebook allows you to easily predict the structure of a protein using a slightly simplified version of [AlphaFold v2.1.0](https://doi.org/10.1038/s41586-021-03819-2). 

**Differences to AlphaFold v2.1.0**

In comparison to AlphaFold v2.1.0, this Colab notebook uses **no templates (homologous structures)** and a selected portion of the [BFD database](https://bfd.mmseqs.com/). We have validated these changes on several thousand recent PDB structures. While accuracy will be near-identical to the full AlphaFold system on many targets, a small fraction have a large drop in accuracy due to the smaller MSA and lack of templates. For best reliability, we recommend instead using the [full open source AlphaFold](https://github.com/deepmind/alphafold/), or the [AlphaFold Protein Structure Database](https://alphafold.ebi.ac.uk/).

**This Colab has a small drop in average accuracy for multimers compared to local AlphaFold installation, for full multimer accuracy it is highly recommended to run [AlphaFold locally](https://github.com/deepmind/alphafold#running-alphafold).** Moreover, the AlphaFold-Multimer requires searching for MSA for every unique sequence in the complex, hence it is substantially slower. If your notebook times-out due to slow multimer MSA search, we recommend either using Colab Pro or running AlphaFold locally.

Please note that this Colab notebook is provided as an early-access prototype and is not a finished product. It is provided for theoretical modelling only and caution should be exercised in its use. 

The **PAE file format** has been updated to match AFDB. Please see the [AFDB FAQ](https://alphafold.ebi.ac.uk/faq/#faq-7) for a description of the new format.

**Citing this work**

Any publication that discloses findings arising from using this notebook should [cite](https://github.com/deepmind/alphafold/#citing-this-work) the [AlphaFold paper](https://doi.org/10.1038/s41586-021-03819-2).

**Licenses**

This Colab uses the [AlphaFold model parameters](https://github.com/deepmind/alphafold/#model-parameters-license) which are subject to the Creative Commons Attribution 4.0 International ([CC BY 4.0](https://creativecommons.org/licenses/by/4.0/legalcode)) license. The Colab itself is provided under the [Apache 2.0 license](https://www.apache.org/licenses/LICENSE-2.0). See the full license statement below.

**More information**

You can find more information about how AlphaFold works in the following papers:

*   [AlphaFold methods paper](https://www.nature.com/articles/s41586-021-03819-2)
*   [AlphaFold predictions of the human proteome paper](https://www.nature.com/articles/s41586-021-03828-1)
*   [AlphaFold-Multimer paper](https://www.biorxiv.org/content/10.1101/2021.10.04.463034v1)

FAQ on how to interpret AlphaFold predictions are [here](https://alphafold.ebi.ac.uk/faq).

If you have any questions not covered in the FAQ, please contact the AlphaFold team at.

**Get in touch**

We would love to hear your feedback and understand how AlphaFold has been useful in your research. Share your stories with us at.

## Setup

Start by running the 2 cells below to set up AlphaFold and all required software.
"""

# Commented out IPython magic to ensure Python compatibility.
#@title 1. Install third-party software

#@markdown Please execute this cell by pressing the _Play_ button 
#@markdown on the left to download and import third-party software 
#@markdown in this Colab notebook. (See the [acknowledgements](https://github.com/deepmind/alphafold/#acknowledgements) in our readme.)

#@markdown **Note**: This installs the software on the Colab 
#@markdown notebook in the cloud and not on your computer.

from IPython.utils import io
import os
import subprocess
import tqdm.notebook

TQDM_BAR_FORMAT = '{l_bar}{bar}| {n_fmt}/{total_fmt} [elapsed: {elapsed} remaining: {remaining}]'

try:
  with tqdm.notebook.tqdm(total=100, bar_format=TQDM_BAR_FORMAT) as pbar:
    with io.capture_output() as captured:
      # Uninstall default Colab version of TF.
#       %shell pip uninstall -y tensorflow

#       %shell sudo apt install --quiet --yes hmmer
      pbar.update(6)

      # Install py3dmol.
#       %shell pip install py3dmol
      pbar.update(2)

      # Install OpenMM and pdbfixer.
#       %shell rm -rf /opt/conda
#       %shell wget -q -P /tmp \
        https://repo.anaconda.com/miniconda/Miniconda3-latest-Linux-x86_64.sh \
          && bash /tmp/Miniconda3-latest-Linux-x86_64.sh -b -p /opt/conda \
          && rm /tmp/Miniconda3-latest-Linux-x86_64.sh
      pbar.update(9)

      PATH=%env PATH
#       %env PATH=/opt/conda/bin:{PATH}
#       %shell conda install -qy conda==4.13.0 \
          && conda install -qy -c conda-forge \
            python=3.7 \
            openmm=7.5.1 \
            pdbfixer
      pbar.update(80)

      # Create a ramdisk to store a database chunk to make Jackhmmer run fast.
#       %shell sudo mkdir -m 777 --parents /tmp/ramdisk
#       %shell sudo mount -t tmpfs -o size=9G ramdisk /tmp/ramdisk
      pbar.update(2)

#       %shell wget -q -P /content \
        https://git.scicore.unibas.ch/schwede/openstructure/-/raw/7102c63615b64735c4941278d92b554ec94415f8/modules/mol/alg/src/stereo_chemical_props.txt
      pbar.update(1)
except subprocess.CalledProcessError:
  print(captured)
  raise

# Commented out IPython magic to ensure Python compatibility.
#@title 2. Download AlphaFold

#@markdown Please execute this cell by pressing the *Play* button on 
#@markdown the left.

GIT_REPO = 'https://github.com/deepmind/alphafold'

SOURCE_URL = 'https://storage.googleapis.com/alphafold/alphafold_params_colab_2022-03-02.tar'
PARAMS_DIR = './alphafold/data/params'
PARAMS_PATH = os.path.join(PARAMS_DIR, os.path.basename(SOURCE_URL))

try:
  with tqdm.notebook.tqdm(total=100, bar_format=TQDM_BAR_FORMAT) as pbar:
    with io.capture_output() as captured:
#       %shell rm -rf alphafold
#       %shell git clone --branch main {GIT_REPO} alphafold
      pbar.update(8)
      # Install the required versions of all dependencies.
#       %shell pip3 install -r ./alphafold/requirements.txt
      # Run setup.py to install only AlphaFold.
#       %shell pip3 install --no-dependencies ./alphafold
      pbar.update(10)

      # Apply OpenMM patch.
#       %shell pushd /opt/conda/lib/python3.7/site-packages/ && \
          patch -p0 < /content/alphafold/docker/openmm.patch && \
          popd

      # Make sure stereo_chemical_props.txt is in all locations where it could be searched for.
#       %shell mkdir -p /content/alphafold/alphafold/common
#       %shell cp -f /content/stereo_chemical_props.txt /content/alphafold/alphafold/common
#       %shell mkdir -p /opt/conda/lib/python3.7/site-packages/alphafold/common/
#       %shell cp -f /content/stereo_chemical_props.txt /opt/conda/lib/python3.7/site-packages/alphafold/common/

#       %shell mkdir --parents "{PARAMS_DIR}"
#       %shell wget -O "{PARAMS_PATH}" "{SOURCE_URL}"
      pbar.update(27)

#       %shell tar --extract --verbose --file="{PARAMS_PATH}" \
        --directory="{PARAMS_DIR}" --preserve-permissions
#       %shell rm "{PARAMS_PATH}"
      pbar.update(55)
except subprocess.CalledProcessError:
  print(captured)
  raise

import jax
if jax.local_devices()[0].platform == 'tpu':
  raise RuntimeError('Colab TPU runtime not supported. Change it to GPU via Runtime -> Change Runtime Type -> Hardware accelerator -> GPU.')
elif jax.local_devices()[0].platform == 'cpu':
  raise RuntimeError('Colab CPU runtime not supported. Change it to GPU via Runtime -> Change Runtime Type -> Hardware accelerator -> GPU.')
else:
  print(f'Running with {jax.local_devices()[0].device_kind} GPU')

# Make sure everything we need is on the path.
import sys
sys.path.append('/opt/conda/lib/python3.7/site-packages')
sys.path.append('/content/alphafold')

# Make sure all necessary environment variables are set.
import os
os.environ['TF_FORCE_UNIFIED_MEMORY'] = '1'
os.environ['XLA_PYTHON_CLIENT_MEM_FRACTION'] = '2.0'

"""## Making a prediction

Please paste the sequence of your protein in the text box below, then run the remaining cells via _Runtime_ > _Run after_. You can also run the cells individually by pressing the _Play_ button on the left.

Note that the search against databases and the actual prediction can take some time, from minutes to hours, depending on the length of the protein and what type of GPU you are allocated by Colab (see FAQ below).
"""

#@title 3. Enter the amino acid sequence(s) to fold ⬇️
#@markdown Enter the amino acid sequence(s) to fold:
#@markdown * If you enter only a single sequence, the monomer model will be used.
#@markdown * If you enter multiple sequences, the multimer model will be used.

from alphafold.notebooks import notebook_utils

sequence_1 = 'MAAHKGAEHHHKAAEHHEQAAKHHHAAAEHHEKGEHEQAAHHADTAYAHHKHAEEHAAQAAKHDAEHHAPKPH'  #@param {type:"string"}
sequence_2 = ''  #@param {type:"string"}
sequence_3 = ''  #@param {type:"string"}
sequence_4 = ''  #@param {type:"string"}
sequence_5 = ''  #@param {type:"string"}
sequence_6 = ''  #@param {type:"string"}
sequence_7 = ''  #@param {type:"string"}
sequence_8 = ''  #@param {type:"string"}

input_sequences = (sequence_1, sequence_2, sequence_3, sequence_4,
                   sequence_5, sequence_6, sequence_7, sequence_8)

MIN_SINGLE_SEQUENCE_LENGTH = 16
MAX_SINGLE_SEQUENCE_LENGTH = 2500
MAX_MULTIMER_LENGTH = 2500

# Validate the input.
sequences, model_type_to_use = notebook_utils.validate_input(
    input_sequences=input_sequences,
    min_length=MIN_SINGLE_SEQUENCE_LENGTH,
    max_length=MAX_SINGLE_SEQUENCE_LENGTH,
    max_multimer_length=MAX_MULTIMER_LENGTH)

#@title 4. Search against genetic databases

#@markdown Once this cell has been executed, you will see
#@markdown statistics about the multiple sequence alignment 
#@markdown (MSA) that will be used by AlphaFold. In particular, 
#@markdown you’ll see how well each residue is covered by similar 
#@markdown sequences in the MSA.

# --- Python imports ---
import collections
import copy
from concurrent import futures
import json
import random

from urllib import request
from google.colab import files
from matplotlib import gridspec
import matplotlib.pyplot as plt
import numpy as np
import py3Dmol

from alphafold.model import model
from alphafold.model import config
from alphafold.model import data

from alphafold.data import feature_processing
from alphafold.data import msa_pairing
from alphafold.data import pipeline
from alphafold.data import pipeline_multimer
from alphafold.data.tools import jackhmmer

from alphafold.common import protein

from alphafold.relax import relax
from alphafold.relax import utils

from IPython import display
from ipywidgets import GridspecLayout
from ipywidgets import Output

# Color bands for visualizing plddt
PLDDT_BANDS = [(0, 50, '#FF7D45'),
               (50, 70, '#FFDB13'),
               (70, 90, '#65CBF3'),
               (90, 100, '#0053D6')]

# --- Find the closest source ---
test_url_pattern = 'https://storage.googleapis.com/alphafold-colab{:s}/latest/uniref90_2021_03.fasta.1'
ex = futures.ThreadPoolExecutor(3)
def fetch(source):
  request.urlretrieve(test_url_pattern.format(source))
  return source
fs = [ex.submit(fetch, source) for source in ['', '-europe', '-asia']]
source = None
for f in futures.as_completed(fs):
  source = f.result()
  ex.shutdown()
  break

JACKHMMER_BINARY_PATH = '/usr/bin/jackhmmer'
DB_ROOT_PATH = f'https://storage.googleapis.com/alphafold-colab{source}/latest/'
# The z_value is the number of sequences in a database.
MSA_DATABASES = [
    {'db_name': 'uniref90',
     'db_path': f'{DB_ROOT_PATH}uniref90_2021_03.fasta',
     'num_streamed_chunks': 59,
     'z_value': 135_301_051},
    {'db_name': 'smallbfd',
     'db_path': f'{DB_ROOT_PATH}bfd-first_non_consensus_sequences.fasta',
     'num_streamed_chunks': 17,
     'z_value': 65_984_053},
    {'db_name': 'mgnify',
     'db_path': f'{DB_ROOT_PATH}mgy_clusters_2019_05.fasta',
     'num_streamed_chunks': 71,
     'z_value': 304_820_129},
]

# Search UniProt and construct the all_seq features only for heteromers, not homomers.
if model_type_to_use == notebook_utils.ModelType.MULTIMER and len(set(sequences)) > 1:
  MSA_DATABASES.extend([
      # Swiss-Prot and TrEMBL are concatenated together as UniProt.
      {'db_name': 'uniprot',
       'db_path': f'{DB_ROOT_PATH}uniprot_2021_03.fasta',
       'num_streamed_chunks': 98,
       'z_value': 219_174_961 + 565_254},
  ])

TOTAL_JACKHMMER_CHUNKS = sum([cfg['num_streamed_chunks'] for cfg in MSA_DATABASES])

MAX_HITS = {
    'uniref90': 10_000,
    'smallbfd': 5_000,
    'mgnify': 501,
    'uniprot': 50_000,
}


def get_msa(fasta_path):
  """Searches for MSA for the given sequence using chunked Jackhmmer search."""

  # Run the search against chunks of genetic databases (since the genetic
  # databases don't fit in Colab disk).
  raw_msa_results = collections.defaultdict(list)
  with tqdm.notebook.tqdm(total=TOTAL_JACKHMMER_CHUNKS, bar_format=TQDM_BAR_FORMAT) as pbar:
    def jackhmmer_chunk_callback(i):
      pbar.update(n=1)

    for db_config in MSA_DATABASES:
      db_name = db_config['db_name']
      pbar.set_description(f'Searching {db_name}')
      jackhmmer_runner = jackhmmer.Jackhmmer(
          binary_path=JACKHMMER_BINARY_PATH,
          database_path=db_config['db_path'],
          get_tblout=True,
          num_streamed_chunks=db_config['num_streamed_chunks'],
          streaming_callback=jackhmmer_chunk_callback,
          z_value=db_config['z_value'])
      # Group the results by database name.
      raw_msa_results[db_name].extend(jackhmmer_runner.query(fasta_path))

  return raw_msa_results


features_for_chain = {}
raw_msa_results_for_sequence = {}
for sequence_index, sequence in enumerate(sequences, start=1):
  print(f'\nGetting MSA for sequence {sequence_index}')

  fasta_path = f'target_{sequence_index}.fasta'
  with open(fasta_path, 'wt') as f:
    f.write(f'>query\n{sequence}')

  # Don't do redundant work for multiple copies of the same chain in the multimer.
  if sequence not in raw_msa_results_for_sequence:
    raw_msa_results = get_msa(fasta_path=fasta_path)
    raw_msa_results_for_sequence[sequence] = raw_msa_results
  else:
    raw_msa_results = copy.deepcopy(raw_msa_results_for_sequence[sequence])

  # Extract the MSAs from the Stockholm files.
  # NB: deduplication happens later in pipeline.make_msa_features.
  single_chain_msas = []
  uniprot_msa = None
  for db_name, db_results in raw_msa_results.items():
    merged_msa = notebook_utils.merge_chunked_msa(
        results=db_results, max_hits=MAX_HITS.get(db_name))
    if merged_msa.sequences and db_name != 'uniprot':
      single_chain_msas.append(merged_msa)
      msa_size = len(set(merged_msa.sequences))
      print(f'{msa_size} unique sequences found in {db_name} for sequence {sequence_index}')
    elif merged_msa.sequences and db_name == 'uniprot':
      uniprot_msa = merged_msa

  notebook_utils.show_msa_info(single_chain_msas=single_chain_msas, sequence_index=sequence_index)

  # Turn the raw data into model features.
  feature_dict = {}
  feature_dict.update(pipeline.make_sequence_features(
      sequence=sequence, description='query', num_res=len(sequence)))
  feature_dict.update(pipeline.make_msa_features(msas=single_chain_msas))
  # We don't use templates in AlphaFold Colab notebook, add only empty placeholder features.
  feature_dict.update(notebook_utils.empty_placeholder_template_features(
      num_templates=0, num_res=len(sequence)))

  # Construct the all_seq features only for heteromers, not homomers.
  if model_type_to_use == notebook_utils.ModelType.MULTIMER and len(set(sequences)) > 1:
    valid_feats = msa_pairing.MSA_FEATURES + (
        'msa_species_identifiers',
    )
    all_seq_features = {
        f'{k}_all_seq': v for k, v in pipeline.make_msa_features([uniprot_msa]).items()
        if k in valid_feats}
    feature_dict.update(all_seq_features)

  features_for_chain[protein.PDB_CHAIN_IDS[sequence_index - 1]] = feature_dict


# Do further feature post-processing depending on the model type.
if model_type_to_use == notebook_utils.ModelType.MONOMER:
  np_example = features_for_chain[protein.PDB_CHAIN_IDS[0]]

elif model_type_to_use == notebook_utils.ModelType.MULTIMER:
  all_chain_features = {}
  for chain_id, chain_features in features_for_chain.items():
    all_chain_features[chain_id] = pipeline_multimer.convert_monomer_features(
        chain_features, chain_id)

  all_chain_features = pipeline_multimer.add_assembly_features(all_chain_features)

  np_example = feature_processing.pair_and_merge(
      all_chain_features=all_chain_features)

  # Pad MSA to avoid zero-sized extra_msa.
  np_example = pipeline_multimer.pad_msa(np_example, min_num_seq=512)

#@title 5. Run AlphaFold and download prediction

#@markdown Once this cell has been executed, a zip-archive with
#@markdown the obtained prediction will be automatically downloaded
#@markdown to your computer.

#@markdown In case you are having issues with the relaxation stage, you can disable it below.
#@markdown Warning: This means that the prediction might have distracting
#@markdown small stereochemical violations.

run_relax = True  #@param {type:"boolean"}

#@markdown Relaxation is faster with a GPU, but we have found it to be less stable.
#@markdown You may wish to enable GPU for higher performance, but if it doesn't
#@markdown converge we suggested reverting to using without GPU.

relax_use_gpu = False  #@param {type:"boolean"}

# --- Run the model ---
if model_type_to_use == notebook_utils.ModelType.MONOMER:
  model_names = config.MODEL_PRESETS['monomer'] + ('model_2_ptm',)
elif model_type_to_use == notebook_utils.ModelType.MULTIMER:
  model_names = config.MODEL_PRESETS['multimer']

output_dir = 'prediction'
os.makedirs(output_dir, exist_ok=True)

plddts = {}
ranking_confidences = {}
pae_outputs = {}
unrelaxed_proteins = {}

with tqdm.notebook.tqdm(total=len(model_names) + 1, bar_format=TQDM_BAR_FORMAT) as pbar:
  for model_name in model_names:
    pbar.set_description(f'Running {model_name}')

    cfg = config.model_config(model_name)
    if model_type_to_use == notebook_utils.ModelType.MONOMER:
      cfg.data.eval.num_ensemble = 1
    elif model_type_to_use == notebook_utils.ModelType.MULTIMER:
      cfg.model.num_ensemble_eval = 1
    params = data.get_model_haiku_params(model_name, './alphafold/data')
    model_runner = model.RunModel(cfg, params)
    processed_feature_dict = model_runner.process_features(np_example, random_seed=0)
    prediction = model_runner.predict(processed_feature_dict, random_seed=random.randrange(sys.maxsize))

    mean_plddt = prediction['plddt'].mean()

    if model_type_to_use == notebook_utils.ModelType.MONOMER:
      if 'predicted_aligned_error' in prediction:
        pae_outputs[model_name] = (prediction['predicted_aligned_error'],
                                   prediction['max_predicted_aligned_error'])
      else:
        # Monomer models are sorted by mean pLDDT. Do not put monomer pTM models here as they
        # should never get selected.
        ranking_confidences[model_name] = prediction['ranking_confidence']
        plddts[model_name] = prediction['plddt']
    elif model_type_to_use == notebook_utils.ModelType.MULTIMER:
      # Multimer models are sorted by pTM+ipTM.
      ranking_confidences[model_name] = prediction['ranking_confidence']
      plddts[model_name] = prediction['plddt']
      pae_outputs[model_name] = (prediction['predicted_aligned_error'],
                                 prediction['max_predicted_aligned_error'])

    # Set the b-factors to the per-residue plddt.
    final_atom_mask = prediction['structure_module']['final_atom_mask']
    b_factors = prediction['plddt'][:, None] * final_atom_mask
    unrelaxed_protein = protein.from_prediction(
        processed_feature_dict,
        prediction,
        b_factors=b_factors,
        remove_leading_feature_dimension=(
            model_type_to_use == notebook_utils.ModelType.MONOMER))
    unrelaxed_proteins[model_name] = unrelaxed_protein

    # Delete unused outputs to save memory.
    del model_runner
    del params
    del prediction
    pbar.update(n=1)

  # --- AMBER relax the best model ---

  # Find the best model according to the mean pLDDT.
  best_model_name = max(ranking_confidences.keys(), key=lambda x: ranking_confidences[x])

  if run_relax:
    pbar.set_description(f'AMBER relaxation')
    amber_relaxer = relax.AmberRelaxation(
        max_iterations=0,
        tolerance=2.39,
        stiffness=10.0,
        exclude_residues=[],
        max_outer_iterations=3,
        use_gpu=relax_use_gpu)
    relaxed_pdb, _, _ = amber_relaxer.process(prot=unrelaxed_proteins[best_model_name])
  else:
    print('Warning: Running without the relaxation stage.')
    relaxed_pdb = protein.to_pdb(unrelaxed_proteins[best_model_name])
  pbar.update(n=1)  # Finished AMBER relax.

# Construct multiclass b-factors to indicate confidence bands
# 0=very low, 1=low, 2=confident, 3=very high
banded_b_factors = []
for plddt in plddts[best_model_name]:
  for idx, (min_val, max_val, _) in enumerate(PLDDT_BANDS):
    if plddt >= min_val and plddt <= max_val:
      banded_b_factors.append(idx)
      break
banded_b_factors = np.array(banded_b_factors)[:, None] * final_atom_mask
to_visualize_pdb = utils.overwrite_b_factors(relaxed_pdb, banded_b_factors)


# Write out the prediction
pred_output_path = os.path.join(output_dir, 'selected_prediction.pdb')
with open(pred_output_path, 'w') as f:
  f.write(relaxed_pdb)


# --- Visualise the prediction & confidence ---
show_sidechains = True
def plot_plddt_legend():
  """Plots the legend for pLDDT."""
  thresh = ['Very low (pLDDT < 50)',
            'Low (70 > pLDDT > 50)',
            'Confident (90 > pLDDT > 70)',
            'Very high (pLDDT > 90)']

  colors = [x[2] for x in PLDDT_BANDS]

  plt.figure(figsize=(2, 2))
  for c in colors:
    plt.bar(0, 0, color=c)
  plt.legend(thresh, frameon=False, loc='center', fontsize=20)
  plt.xticks([])
  plt.yticks([])
  ax = plt.gca()
  ax.spines['right'].set_visible(False)
  ax.spines['top'].set_visible(False)
  ax.spines['left'].set_visible(False)
  ax.spines['bottom'].set_visible(False)
  plt.title('Model Confidence', fontsize=20, pad=20)
  return plt

# Show the structure coloured by chain if the multimer model has been used.
if model_type_to_use == notebook_utils.ModelType.MULTIMER:
  multichain_view = py3Dmol.view(width=800, height=600)
  multichain_view.addModelsAsFrames(to_visualize_pdb)
  multichain_style = {'cartoon': {'colorscheme': 'chain'}}
  multichain_view.setStyle({'model': -1}, multichain_style)
  multichain_view.zoomTo()
  multichain_view.show()

# Color the structure by per-residue pLDDT
color_map = {i: bands[2] for i, bands in enumerate(PLDDT_BANDS)}
view = py3Dmol.view(width=800, height=600)
view.addModelsAsFrames(to_visualize_pdb)
style = {'cartoon': {'colorscheme': {'prop': 'b', 'map': color_map}}}
if show_sidechains:
  style['stick'] = {}
view.setStyle({'model': -1}, style)
view.zoomTo()

grid = GridspecLayout(1, 2)
out = Output()
with out:
  view.show()
grid[0, 0] = out

out = Output()
with out:
  plot_plddt_legend().show()
grid[0, 1] = out

display.display(grid)

# Display pLDDT and predicted aligned error (if output by the model).
if pae_outputs:
  num_plots = 2
else:
  num_plots = 1

plt.figure(figsize=[8 * num_plots, 6])
plt.subplot(1, num_plots, 1)
plt.plot(plddts[best_model_name])
plt.title('Predicted LDDT')
plt.xlabel('Residue')
plt.ylabel('pLDDT')

if num_plots == 2:
  plt.subplot(1, 2, 2)
  pae, max_pae = list(pae_outputs.values())[0]
  plt.imshow(pae, vmin=0., vmax=max_pae, cmap='Greens_r')
  plt.colorbar(fraction=0.046, pad=0.04)

  # Display lines at chain boundaries.
  best_unrelaxed_prot = unrelaxed_proteins[best_model_name]
  total_num_res = best_unrelaxed_prot.residue_index.shape[-1]
  chain_ids = best_unrelaxed_prot.chain_index
  for chain_boundary in np.nonzero(chain_ids[:-1] - chain_ids[1:]):
    if chain_boundary.size:
      plt.plot([0, total_num_res], [chain_boundary, chain_boundary], color='red')
      plt.plot([chain_boundary, chain_boundary], [0, total_num_res], color='red')

  plt.title('Predicted Aligned Error')
  plt.xlabel('Scored residue')
  plt.ylabel('Aligned residue')

# Save the predicted aligned error (if it exists).
pae_output_path = os.path.join(output_dir, 'predicted_aligned_error.json')
if pae_outputs:
  # Save predicted aligned error in the same format as the AF EMBL DB.
  pae_data = notebook_utils.get_pae_json(pae=pae, max_pae=max_pae.item())
  with open(pae_output_path, 'w') as f:
    f.write(pae_data)

# --- Download the predictions ---
!zip -q -r {output_dir}.zip {output_dir}
files.download(f'{output_dir}.zip')

"""### Interpreting the prediction

In general predicted LDDT (pLDDT) is best used for intra-domain confidence, whereas Predicted Aligned Error (PAE) is best used for determining between domain or between chain confidence.

Please see the [AlphaFold methods paper](https://www.nature.com/articles/s41586-021-03819-2), the [AlphaFold predictions of the human proteome paper](https://www.nature.com/articles/s41586-021-03828-1), and the [AlphaFold-Multimer paper](https://www.biorxiv.org/content/10.1101/2021.10.04.463034v1) as well as [our FAQ](https://alphafold.ebi.ac.uk/faq) on how to interpret AlphaFold predictions.

## FAQ & Troubleshooting


*   How do I get a predicted protein structure for my protein?
    *   Click on the _Connect_ button on the top right to get started.
    *   Paste the amino acid sequence of your protein (without any headers) into the “Enter the amino acid sequence to fold”.
    *   Run all cells in the Colab, either by running them individually (with the play button on the left side) or via _Runtime_ > _Run all._ Make sure you run all 5 cells in order.
    *   The predicted protein structure will be downloaded once all cells have been executed. Note: This can take minutes to hours - see below.
*   How long will this take?
    *   Downloading the AlphaFold source code can take up to a few minutes.
    *   Downloading and installing the third-party software can take up to a few minutes.
    *   The search against genetic databases can take minutes to hours.
    *   Running AlphaFold and generating the prediction can take minutes to hours, depending on the length of your protein and on which GPU-type Colab has assigned you.
*   My Colab no longer seems to be doing anything, what should I do?
    *   Some steps may take minutes to hours to complete.
    *   If nothing happens or if you receive an error message, try restarting your Colab runtime via _Runtime_ > _Restart runtime_.
    *   If this doesn’t help, try resetting your Colab runtime via _Runtime_ > _Factory reset runtime_.
*   How does this compare to the open-source version of AlphaFold?
    *   This Colab version of AlphaFold searches a selected portion of the BFD dataset and currently doesn’t use templates, so its accuracy is reduced in comparison to the full version of AlphaFold that is described in the [AlphaFold paper](https://doi.org/10.1038/s41586-021-03819-2) and [Github repo](https://github.com/deepmind/alphafold/) (the full version is available via the inference script).
*   What is a Colab?
    *   See the [Colab FAQ](https://research.google.com/colaboratory/faq.html).
*   I received a warning “Notebook requires high RAM”, what do I do?
    *   The resources allocated to your Colab vary. See the [Colab FAQ](https://research.google.com/colaboratory/faq.html) for more details.
    *   You can execute the Colab nonetheless.
*   I received an error “Colab CPU runtime not supported” or “No GPU/TPU found”, what do I do?
    *   Colab CPU runtime is not supported. Try changing your runtime via _Runtime_ > _Change runtime type_ > _Hardware accelerator_ > _GPU_.
    *   The type of GPU allocated to your Colab varies. See the [Colab FAQ](https://research.google.com/colaboratory/faq.html) for more details.
    *   If you receive “Cannot connect to GPU backend”, you can try again later to see if Colab allocates you a GPU.
    *   [Colab Pro](https://colab.research.google.com/signup) offers priority access to GPUs.
*   I received an error “ModuleNotFoundError: No module named ...”, even though I ran the cell that imports it, what do I do?
    *   Colab notebooks on the free tier time out after a certain amount of time. See the [Colab FAQ](https://research.google.com/colaboratory/faq.html#idle-timeouts). Try rerunning the whole notebook from the beginning.
*   Does this tool install anything on my computer?
    *   No, everything happens in the cloud on Google Colab.
    *   At the end of the Colab execution a zip-archive with the obtained prediction will be automatically downloaded to your computer.
*   How should I share feedback and bug reports?
    *   Please share any feedback and bug reports as an [issue](https://github.com/deepmind/alphafold/issues) on Github.


## Related work

Take a look at these Colab notebooks provided by the community (please note that these notebooks may vary from our validated AlphaFold system and we cannot guarantee their accuracy):

*   The [ColabFold AlphaFold2 notebook](https://colab.research.google.com/github/sokrypton/ColabFold/blob/main/AlphaFold2.ipynb) by Sergey Ovchinnikov, Milot Mirdita and Martin Steinegger, which uses an API hosted at the Södinglab based on the MMseqs2 server ([Mirdita et al. 2019, Bioinformatics](https://academic.oup.com/bioinformatics/article/35/16/2856/5280135)) for the multiple sequence alignment creation.
"""


"""# License and Disclaimer

This is not an officially-supported Google product.

This Colab notebook and other information provided is for theoretical modelling only, caution should be exercised in its use. It is provided ‘as-is’ without any warranty of any kind, whether expressed or implied. Information is not intended to be a substitute for professional medical advice, diagnosis, or treatment, and does not constitute medical or other professional advice.

Copyright 2021 DeepMind Technologies Limited.


## AlphaFold Code License

Licensed under the Apache License, Version 2.0 (the "License"); you may not use this file except in compliance with the License. You may obtain a copy of the License at https://www.apache.org/licenses/LICENSE-2.0.

Unless required by applicable law or agreed to in writing, software distributed under the License is distributed on an "AS IS" BASIS, WITHOUT WARRANTIES OR CONDITIONS OF ANY KIND, either express or implied. See the License for the specific language governing permissions and limitations under the License.

## Model Parameters License

The AlphaFold parameters are made available under the terms of the Creative Commons Attribution 4.0 International (CC BY 4.0) license. You can find details at: https://creativecommons.org/licenses/by/4.0/legalcode


## Third-party software

Use of the third-party software, libraries or code referred to in the [Acknowledgements section](https://github.com/deepmind/alphafold/#acknowledgements) in the AlphaFold README may be governed by separate terms and conditions or license provisions. Your use of the third-party software, libraries or code is subject to any such terms and you should check that you can comply with any applicable restrictions or terms and conditions before use.


## Mirrored Databases

The following databases have been mirrored by DeepMind, and are available with reference to the following:
* UniProt: v2021\_03 (unmodified), by The UniProt Consortium, available under a [Creative Commons Attribution-NoDerivatives 4.0 International License](http://creativecommons.org/licenses/by-nd/4.0/).
* UniRef90: v2021\_03 (unmodified), by The UniProt Consortium, available under a [Creative Commons Attribution-NoDerivatives 4.0 International License](http://creativecommons.org/licenses/by-nd/4.0/).
* MGnify: v2019\_05 (unmodified), by Mitchell AL et al., available free of all copyright restrictions and made fully and freely available for both non-commercial and commercial use under [CC0 1.0 Universal (CC0 1.0) Public Domain Dedication](https://creativecommons.org/publicdomain/zero/1.0/).
* BFD: (modified), by Steinegger M. and Söding J., modified by DeepMind, available under a [Creative Commons Attribution-ShareAlike 4.0 International License](https://creativecommons.org/licenses/by/4.0/). See the Methods section of the [AlphaFold proteome paper](https://www.nature.com/articles/s41586-021-03828-1) for details.
"""
```
