## Supplementary material for "*Clostridioides difficile* Toxins Unhinged: Allosterically Switchable Network Orients *β*-flap": Scripts, Structures etc.: analyze_AF_confs.py

```
import sys
import os
import numpy as np
import matplotlib.pyplot as plt
from Bio.PDB import *
# Hamelryck, T., Manderick, B. (2003) PDB parser and structure class implemented
# in Python. Bioinformatics 19: 2308–2310
sys.path.insert(0, "/home/lf1071fu/project_b3/ProjectB3")
from tools import utils, traj_funcs
import config.settings as cf

def main(argv):

    # Set up specifications for the system and initialize some general variables
    template = False
    path = f"{ cf.path_head }/alphafold"
    global holo, apo, msa_depths
    holo, apo = "3pee", "6oq5"
    msa_depths = [16, 32, 64, 128, 256, 512, 1024] #, 5120]

    central = np.array(list(range(8,195)) + list(range(234,252)))
    pdb_files = [file for file in os.listdir(f"{ path }/MSA_depths/no_template_structures") if file.endswith('.pdb')]
    print(pdb_files)

    residues = {holo : range(8, 252), apo : range(552, 796),
                "full-cpd" : range(1, 255), "apo-cpd" : range(545, 799),
                "central" : central, "central_apo" : central + 545,
                "beta_flap" : range(198,232), "beta_flap_apo" : range(743,777)}

    # Create kSeys and collect paths to pdbs for all reference and structure
    ids = [holo, apo]
    files = [f"{ path }/structures/3pee_A.pdb",
             f"{ path }/structures/6oq5_cpd.pdb"]
    for n in pdb_files:
        i = n.split("_")[0]
        d = n.split("_")[1].split(".")[0]
        ids.append(f"m{i}d{d}")
        if template:
            files.append(f"{ path }/apo_template_structures/{i}_{d}")
        else:
            files.append(f"{ path }/MSA_depths/no_template_structures/{ n }")


    # Make a dictionary of the key data on each structure
    models = get_models(ids, files, residues, holo)

    # Determine TM scores and RMSD to the beta flap region
    for key, mod in models.items():

        mod.holo_tm_score = get_tm_score(models[holo].alphaCs, mod.alphaCs)
        mod.holo_rmsd = calc_rmsd(models[holo].beta_flap, mod.beta_flap)

        mod.apo_tm_score = get_tm_score(models[apo].alphaCs, mod.alphaCs)
        mod.apo_rmsd = calc_rmsd(models[apo].beta_flap, mod.beta_flap)

    # plot_tm_scores(models, path, template)
    plot_rmsd_beta_flap(models)

    # The rmsf is calculated for a particular msa depth
    # depth = 32
    # samples = len([x for x in non_outliers if str(depth) in x])
    # rmsfs = get_rmsfs(models, residues, depth=depth, strs=samples)
    # exp_dists = [a - b for a, b in zip(models[apo].alphaCs,
    #              models[holo].alphaCs)]

    # plot_res_comparison(exp_dists, rmsfs, depth, path, template)
    # plot_rmsf(residues[holo], rmsfs, path, template)

    # # Identify the stable residues and use these for structural alignment
    # align_res = []
    # for i, r in zip(residues[holo], rmsfs):
    #     if r < 0.5:
    #         align_res.append(i)
    # np.save(f"{ path }/alignment_residues.npy", np.array(align_res))

    return None

class Model:
    def __init__(self, id, file, msa_depth=None):
        self.id = id
        self.file = file
        self.msa_depth = msa_depth

        self.alphaCs = None
        self.beta_flap = None

        self.holo_tm_score = 0
        self.apo_tm_score = 0

        self.holo_rmsd = 0
        self.apo_rmsd = 0

    def __str__(self):
        return f"Model object for { self.id } with { self.msa_depth }"

def get_models(ids, files, residues, ref):
    """Returns a dictionary of Model objects based on pdb data.

    Note that "residues" should reflect the most stable residues of the protein,
    i.e. those in the central beta-sheet region. This will be the region of
    alignment for the structures.

    Parameters
    ----------
    ids : (str) list
        A list of keys used to access the Model objects.
    files : (str) list
        A list of the paths to pdb structures.
    residues : ((int) list) dict
        Dictionary related to various groups of residues for alignment,
        selecting all alpha carbons and selecting the beta flap region.
    ref : str
        The key for the reference structure for alignment must also match the
        initial value in ids and files.

    Returns
    -------
    models : (Model) dict
        A dictionary to easily access Model objects, containing structural and
        general data.
    """
    models = dict()

    # Add structures to "models" dict. and align against the reference (3pee)
    for i, file in zip(ids, files):

        print(i, file)
        # Make a Model object of each structure to store and access general and
        # structural data
        if i not in ids[:2]:
            depth = int(i.split("d")[1])
            model = Model(i, file, msa_depth=depth)
        else:
            model = Model(i, file)

        # Parse the pdb file and store as a Bio.PDB.Chain object
        parser = PDBParser(PERMISSIVE=1, QUIET=True)
        struct = parser.get_structure(id, file)[0]["A"]

        # Apply rotation-translation transformations onto the structure to
        # minimize the RMSD wrt the selected residues of the reference
        # structure.
        if i != ref:

            ref_atoms = get_backbone_atoms(models[ref].struct,
                                           residues["central"])
            if i == apo:
                res_num = residues["central_apo"]
            else:
                res_num = residues["central"]
            moving_atoms = get_backbone_atoms(struct, res_num)

            super_imposer = Superimposer()
            super_imposer.set_atoms(ref_atoms, moving_atoms)
            super_imposer.apply(struct.get_atoms())

        model.struct = struct

        if i == apo:
            model.alphaCs = get_backbone_atoms(struct, residues[apo],
                                               alpha_only=True)
            model.beta_flap = get_backbone_atoms(struct,
                                                 residues["beta_flap_apo"])
        else:
            model.alphaCs = get_backbone_atoms(struct, residues[holo],
                                               alpha_only=True)
            model.beta_flap = get_backbone_atoms(struct, residues["beta_flap"])

        models[i] = model

    return models

def get_backbone_atoms(struct, atoms_to_be_aligned, alpha_only=False):
    """Get a list of the backbone atoms from a subset of residues.

    Parameters
    ----------
    struct : Bio.PDB.Chain
        A protein structure from a single chain, loaded from a pdb file.
    atoms_to_be_aligned : (int) iter
        An iterable to select the subset of residues.

    Returns
    -------
    atoms : (Bio.PDB.Atom) list
        An ordered list of backbone atoms from the selected residues.

    """
    atoms = []

    for res in struct.get_residues():
        res_id = res.get_id()[1]
        if res_id in atoms_to_be_aligned:
            if alpha_only:
                atoms += [res["CA"]]
            else:
                atoms += [res["N"], res["CA"], res["C"]]

    return atoms

def get_tm_score(alphaCs_input, alphaCs_ref):
    """Determine the template model scores against an experimental reference.

    The TM-score is calculated according to the reference using the carbon atoms
    of the reference and the input structures.

    Parameters
    ----------
    alphaCs_input : (Bio.PDB.Atom) list
        An ordered list of alpha carbon atoms from the input structure.
    alphaCs_ref : (Bio.PDB.Atom) list
        An ordered list of alpha carbon atoms from the reference structure.

    Returns
    -------
    tm_score : float
        Some number between 0 and 1, inclusive, where numbers closer to 1
        indicate strong similarity to the reference structure.

    """
    d0_param = 1.24 * (len(alphaCs_ref) - 15) ** (1. / 3) - 1.8
    tm_score = 0

    for i, j in zip(alphaCs_ref, alphaCs_input):
        dist = np.linalg.norm(i.get_coord() - j.get_coord())
        tm_score += 1 / ( 1 + ( dist / d0_param ) ** 2 )

    tm_score = tm_score / len(alphaCs_ref)

    return tm_score

def calc_rmsd(atoms, ref_atoms):
    """Calculate the RMSD of the backbone atoms wrt the reference atoms.

    Parameters
    ----------
    atoms : (Bio.PDB.Atom) list
        The backbone N, CA and C atoms of the included residues.
    ref_atoms : (Bio.PDB.Atom) list
        The backbone N, CA and C atoms of the included residues.

    Returns
    -------
    rmsd : float
        The RMSD of the residue group wrt the reference structure.

    """
    dists_sq = []
    n_atoms = len(atoms)

    for atom, ref_atom in zip(atoms, ref_atoms):
        dists_sq.append( np.linalg.norm((atom - ref_atom)) ** 2 )

    rmsd = np.sqrt( 1/n_atoms * sum(dists_sq) )

    return rmsd

def get_rmsfs(models, residues, depth=128, strs=50):
    """Calculate the RMSF of each residue over selected structure.

    Parameters
    ----------
    models : (Model) dict
        A dictionary to easily access Model objects, containing structural and
        general data.
    residues : ((int) list) dict
        Dictionary related to various groups of residues for alignment,
        selecting all alpha carbons and selecting the beta flap region.
    depth : int
        The MSA depth(s) used for the RMSF determination.
    strs : int
        The number of structures used in the calculation must match strs.

    Returns
    -------
    rmsfs : (float) list
        The RMSF of each residue is included in the ordered list.

    """
    # Iterate over the individual models (MSA depth = 128) to get alpha carbon
    # coords and make a nested list of the coordinates for each residue
    model_alphas = np.zeros((244, strs, 3))
    i = 0

    for mod in models.values():

        if mod.msa_depth != depth:
            continue

        c_alpha_coords = list(map(lambda x : x.get_coord(), mod.alphaCs))

        for e, c in enumerate(c_alpha_coords):
            model_alphas[e, i, :] = c

        i += 1

    rmsfs = []
    for Catom in model_alphas:
        rmsfs.append(calc_rmsf(Catom))

    return rmsfs

def calc_rmsf(Catoms):
    """Calculate the RMSF from the list of alpha carbon atom coordinates.

    Parameters
    ----------
    Catoms : (float, float, float) list
        The alpha carbon atom coordinate of the residue is listed for each
        model.

    Returns
    -------
    rmsf : float
        The calculated RMSF of the residue.

    """
    ave_coord = np.mean(Catoms, axis=0)
    n_atoms = len(Catoms)
    dists_sq = []

    for atom in Catoms:
        dists_sq.append( np.linalg.norm((atom - ave_coord)) ** 2 )

    rmsf = np.sqrt( 1/n_atoms * sum(dists_sq) )

    return rmsf

def plot_tm_scores(models, path, template):
    fig, ax = plt.subplots(constrained_layout=True, figsize=(12,8))

    from palettable.cmocean.sequential import Deep_5
    filled_marker_style = dict(marker='o', markersize=15, linestyle="", \
                                markeredgecolor='black')
    colormap = list(Deep_5.mpl_colormap(range(256)))

    for i, d in enumerate(msa_depths):
        holo_scores = []
        apo_scores = []
        for mod in models.values():
            if mod.msa_depth == d:
                holo_scores.append(mod.holo_tm_score)
                apo_scores.append(mod.apo_tm_score)
        ax.plot(holo_scores, apo_scores, label=str(d),
                markerfacecolor=colormap[int(i/(len(msa_depths)) * 256)],
                **filled_marker_style)

    # Plot settings
    ax.tick_params(axis='y', labelsize=14, direction='in', width=2, \
                    length=5, pad=10)
    ax.tick_params(axis='x', labelsize=14, direction='in', width=2, \
                    length=5, pad=10)
    for i in ["top","bottom","left","right"]:
        ax.spines[i].set_linewidth(2)
    ax.grid(True)
    ax.set_xlabel("Similarity to holo state (TM-score)", labelpad=5, \
                    fontsize=22)
    ax.set_ylabel("Similartiy to apo state (TM-score)", labelpad=5, fontsize=22)
    plt.xlim([0.75,1.01])
    plt.ylim([0.75,1.01])
    plt.legend(fontsize=16)

    if template:
        plt.savefig(f"{path}/figures/similarity_TM-score_template.png")
    else:
        plt.savefig(f"{path}/figures/similarity_TM-score_no_template.png")
    plt.close()

    return None

def plot_rmsd_beta_flap(models):
    fig, ax = plt.subplots(constrained_layout=True, figsize=(2.5,2.5))

    cmap = plt.get_cmap("plasma")
    colormap = [cmap(i/255) for i in range(256)]

    for i, d in enumerate(msa_depths):
        holo_rmsd = []
        apo_rmsd = []
        for mod in models.values():
            if mod.msa_depth == d:
                holo_rmsd.append(mod.holo_rmsd)
                apo_rmsd.append(mod.apo_rmsd)
        ax.scatter(apo_rmsd, holo_rmsd, label=str(d), s=10,
                color=colormap[int(i/(len(msa_depths)) * 256)],
                marker='o')
    
    # Add in reference positions
    ax.scatter(0, 10.4, 
                label="Lowered ref.", marker="X", color=cf.closed_color, 
                edgecolors="#ededed", s=50, lw=1)
    ax.scatter(10.4, 0, 
                label="Raised ref.", marker="X", color=cf.open_color, 
                edgecolors="#ededed", s=50, lw=1)

    # Plot settings
    ax.tick_params(axis='both', which='major', pad=3)
    ax.grid(False)
    ax.set_xlabel(r"$\beta-$flap RMSD$_{\mathrm{lowered}}$ ($\AA$)", labelpad=3, fontsize=8)
    ax.set_ylabel(r"$\beta-$flap RMSD$_{\mathrm{raised}}$ ($\AA$)", labelpad=3, \
                    fontsize=8)
    plt.legend(fontsize=6, loc=3, ncols=2)
    plt.xlim([-.5,15])
    plt.ylim([-.5,15])

    utils.save_figure(fig, 
        f"{ cf.figure_head }/alphafold/rmsd_beta-flap_no_template.png")
    plt.show()
    plt.close()

    return None

def plot_res_comparison(exp_dists, rmsfs, depth, path, template):

    fig, ax = plt.subplots(constrained_layout=True, figsize=(12,8))

    filled_marker_style = dict(marker='o', markersize=15, linestyle="", \
                                markeredgecolor='black')
    r_all = np.round(np.corrcoef(exp_dists, rmsfs)[0,1] ** 2, 2)

    r_subset = np.round(np.corrcoef(exp_dists[198:232],
                        rmsfs[198:232])[0,1] ** 2, 2)

    ax.plot(exp_dists, rmsfs, label= "all residues", color="#FF6666", \
            **filled_marker_style)
    ax.plot(exp_dists[198:232], rmsfs[198:232], label="beta-flap residues", \
            color="#1cbbc7", **filled_marker_style)

    # Plot settings
    ax.tick_params(axis='y', labelsize=18, direction='in', width=2, \
                    length=5, pad=10)
    ax.tick_params(axis='x', labelsize=18, direction='in', width=2, \
                    length=5, pad=10)
    for i in ["top","bottom","left","right"]:
        ax.spines[i].set_linewidth(2)
    ax.grid(True)
    ax.set_xlabel(r"Distance between experimental structures ($\AA$)", \
                    labelpad=5, fontsize=28)
    ax.set_ylabel(r"RMSF among AF models ($\AA$)", labelpad=5, fontsize=28)
    plt.legend(fontsize=20)

    # Add in the Pearson correlation coeff
    plt.text(.97, .12, r"R$^2 = $"+str(r_all), ha="right", va="bottom",
             color="#FF6666", transform=ax.transAxes, fontsize=24)
    plt.text(.97, .07, r"R$^2 = $"+str(r_subset), ha="right",
             va="bottom", color="#1cbbc7", transform=ax.transAxes, fontsize=24)

    if template:
        plt.savefig(f"{path}/figures/res_comparison_template_{depth}.png")
    else:
        plt.savefig(f"{path}/figures/res_comparison_no_template_{depth}.png")
    plt.close()

    return None

def plot_rmsf(resnums, rmsf, path, template):
    """Makes an RMSF plot.

    Parameters
    ----------
    resnums : (int) list
        The residue numbers used for the RMSF calculation. This should consist
        of all the alpha carbons in the protein.
    rmsf : np.ndarray
        The rmsf of the selected atom groups.
    path : str
        Path to the primary working directory.
    template : bool
        AF models are generated with or without a template.

    Returns
    -------
    None.

    """
    fig, ax = plt.subplots(constrained_layout=True, figsize=(12,4))
    resids = list(map(lambda x : x + 543, resnums))
    plt.plot(resids, rmsf, lw=3, color="#b21856")

    # Plot settings
    ax.tick_params(axis='y', labelsize=18, direction='in', width=2, \
                    length=5, pad=10)
    ax.tick_params(axis='x', labelsize=18, direction='in', width=2, \
                    length=5, pad=10)
    for i in ["top","bottom","left","right"]:
        ax.spines[i].set_linewidth(2)
    ax.grid(True)
    ax.set_xlabel(r"Residue number", labelpad=5, \
                    fontsize=28)
    ax.set_ylabel(r"RMSF ($\AA$)", labelpad=5, fontsize=28)
    bottom, top = ax.get_ylim()
    ax.vlines([743,776], bottom, top, linestyles="dashed", alpha=0.6, lw=3,
              colors="r")

    if template:
        plt.savefig(f"{path}/figures/rmsf_template.png")
    else:
        plt.savefig(f"{path}/figures/rmsf_no_template.png")
    plt.close()

    return None

if __name__ == '__main__':
    main(sys.argv)
```
