## Supplementary material for "*Clostridioides difficile* Toxins Unhinged: Allosterically Switchable Network Orients *β*-flap": Scripts, Structures etc.: colab_af_confs.py

```
from af2_conformations.scripts import predict
from af2_conformations.scripts import mmseqs2

import random
import os

from absl import logging
logging.set_verbosity(logging.DEBUG)

jobname = 'TcdB_CPD'
sequence = ("GEDDNLDFSQNIVVDKEYLLEKISSLARSSERGYIHYIVQLQGDKISYEAACNLFAKTPYDSVLFQKNIEDSEIAYYYNPGDGEIQEIDKYKIPSIISDRPKIKLTFIGHGKDEFNTDIFAGFDVDSLSTEIEAAIDLAKEDISPKSIEINLLGCNMFSYSINVEETYPGKLLLKVKDKISELMPSISQDSIIVSANQYEVRINSEGRRELLDHSGEWINKEESIIKDISSKEYISFNPKENKITVKSKNLPEL")

# PDB IDs, written uppercase with chain ID specified
# pdbs = ["3PA8_A", "3PEE_A", "6OQ5_A", "7POG_A", "7NY9_A"]

pdbs = ["6OQ5_A", "7POG_A", "7NY9_A"]

path = os.getcwd()

# The MMSeqs2Runner object submits the amino acid sequence to
# the MMSeqs2 server, generates a directory, and populates it with
# data retrieved from the server. Templates may be specified by the user.
# All templates are fetched if none are provided or the list is empty.
mmseqs2_runner = mmseqs2.MMSeqs2Runner( jobname, sequence )

# Fetch sequences and download data
a3m_lines, template_path = mmseqs2_runner.run_job( templates = pdbs )

# A nested loop in which 5 models are generated per MSA depth value
# In our manuscript we use three MSA depths: 32 sequences, 128, and 5120
for nseq in [16,32]:
  for n_model in range(400):

    # Randomly choose one of the two AlphaFold neural
    # networks capable of using templates.
    # In our experience, model 1 is more sensitive to input templates.
    # However, this observation is purely anecdotal and not backed up by
    # hard numbers.
    model_id = random.choice( ( 1, 2 ) )

    # Specify the name of the output PDB
    outname = f"{ path }/no_template_structures/{ n_model }_{ nseq }.pdb"

    # Run the job and save as a PDB
    # file = predict.predict_structure_from_templates(
    #     mmseqs2_runner.seq, # NOTE mmseqs2_runner removes whitespace from seq
    #     outname,
    #     a3m_lines,
    #     template_path = template_path,
    #     model_id = model_id,
    #     max_msa_clusters = nseq // 2,
    #     max_extra_msa = nseq,
    #     max_recycles = 1
    # )

    # Alternatively, users can run a template-free prediction by uncommenting
    # the line below:
    predict.predict_structure_no_templates( sequence, outname,
         a3m_lines, model_id = model_id, max_msa_clusters = nseq // 2,
         max_extra_msa = nseq, max_recycles = 1, n_struct_module_repeats = 8,
         ptm = True)
```
