## Supplementary material for "*Clostridioides difficile* Toxins Unhinged: Allosterically Switchable Network Orients *β*-flap": Scripts, Structures etc.: CombinedRMSD.py

```
import sys
import numpy as np
import os
import matplotlib.pyplot as plt
import MDAnalysis as mda
import TrajFunctions as tf
sys.path.insert(0, "/home/lf1071fu/project_b3/ProjectB3")
from tools import utils, traj_funcs
import config.settings as cf

global home_path, struct_path, holo_state, apo_state

def main(argv):

    # Load in some reference structures
    open_state = mda.Universe(f"{ cf.struct_head }/open_ref_state.pdb")
    closed_state = mda.Universe(f"{ cf.struct_head }/closed_ref_state.pdb")
    ref_state = mda.Universe(f"{ cf.struct_head }/alignment_struct.pdb")

    beta_flap_group = cf.beta_flap_group

    # Make a dictionary containing the RMSD data
    open_RMSDs = dict()
    closed_RMSDs = dict()

    for i in np.arange(1,11):

        # Skip replica 6, no data
        if i == 6:
            continue

        af_path = f"{ home_path }/af_{ i }/md_run"

        # Load in universe objects for the simulation and the reference structures
        u = mda.Universe(f"{ af_path }/topol.top", f"{ af_path }/fitted_traj.xtc",
                         topology_format="ITP")

        open_RMSDs[i], closed_RMSDs[i] = get_rmsds(u, af_path)

    # Make a combined plot
    fig, ax = plt.subplots(constrained_layout=True, figsize=(12,8))

    for i in np.arange(1,11):

        if i == 6:
            continue

        d = ax.scatter(open_RMSDs[i][:,3], closed_RMSDs[i][:,3], label=f"Replica {i}")

    # Plot settings
    ax.tick_params(axis='y', labelsize=18, direction='in', width=2, \
                    length=5, pad=10)
    ax.tick_params(axis='x', labelsize=18, direction='in', width=2, \
                    length=5, pad=10)
    for i in ["top","bottom","left","right"]:
        ax.spines[i].set_linewidth(2)
    ax.grid(True)
    ax.set_xlabel(r"RMSD to holo state ($\AA$)", labelpad=5, \
                    fontsize=28)
    ax.set_ylabel(r"RMSD to apo state ($\AA$)", labelpad=5, fontsize=28)
    _, top = ax.get_ylim()
    _, right = ax.get_xlim()
    ax.set_xlim(0, right)
    ax.set_ylim(0, top)
    plt.legend(fontsize=16)

    plt.savefig(f"{ home_path }/../../figures/unbiased_sims/af_replicas/combined_rmsd.png")
    plt.close()

def get_rmsds(u, path):
    """Get the RMSD to the reference holo & apo structures.

    The RMSD is to the beta flap region--the selected residues can be controlled
    from within the function body.

    Parameters
    ----------
    u : MDAnalysis.core.universe
        The universe object, where the trajectory does not need to be loaded in.
    path : str
        Path to particular af replica.
    Returns
    -------
    R_holo : np.ndarray
        A timeseries of the RMSD against the given reference, for the given
        atom group.
    R_apo : np.ndarray
        A timeseries of the RMSD against the given reference, for the given
        atom group.

    """
    np_files = [f"{ path }/rmsd_holo.npy", f"{ path }/rmsd_apo.npy"]

    if all(list(map(lambda x : os.path.exists(x), np_files))):

        R_holo = np.load(np_files[0], allow_pickle=True)
        R_apo = np.load(np_files[1], allow_pickle=True)

    else:

        u.transfer_to_memory()

        # Determine RMSD to ref structures
        R_holo = tf.get_rmsd(u, holo_state, "backbone and resid 8-251",
                          beta_flap_group, "holo", path)
        R_apo = tf.get_rmsd(u, apo_state,
                         "backbone and (resid 1-253 or resid 545-797)",
                         beta_flap_group, "apo", path)

        for f, v in zip(np_files, [R_holo, R_apo]):
            np.save(f, v)

    return R_holo, R_apo

if __name__ == '__main__':
    main(sys.argv)
```
