## Supplementary material for "*Clostridioides difficile* Toxins Unhinged: Allosterically Switchable Network Orients *β*-flap": Scripts, Structures etc.: gen_af_mutants.py

```
import sys
sys.path.insert(0, "/home/lf1071fu/project_b3/alphafold")
from af2_conformations.scripts import predict
from af2_conformations.scripts import mmseqs2
import json
import random
import os

from absl import logging
logging.set_verbosity(logging.DEBUG)

af_mut_path = "/home/lf1071fu/project_b3/alphafold/af_mutants_betaflap"
str_path = f"{ af_mut_path }/sturctures"
os.chdir(af_mut_path)

with open(f"{ af_mut_path }/sequences.json", "r") as f:
    sequences = json.load(f)

pdbs = ["6OQ5_A", "7POG_A", "7NY9_A"]

# The MMSeqs2Runner object submits the amino acid sequence to
# the MMSeqs2 server, generates a directory, and populates it with
# data retrieved from the server. Templates may be specified by the user.
# All templates are fetched if none are provided or the list is empty.
for mutant, sequence in sequences.items():

  jobname = mutant
  mmseqs2_runner = mmseqs2.MMSeqs2Runner( jobname, sequence )

  # Fetch sequences and download data
  a3m_lines, template_path = mmseqs2_runner.run_job( templates = pdbs )

  # A nested loop in which 10 models are generated per MSA depth value
  for nseq in [16,32,64]:
    for n_model in range(10):

      # Randomly choose one of the two AlphaFold neural
      # networks capable of using templates.
      # In our experience, model 1 is more sensitive to input templates.
      # However, this observation is purely anecdotal and not backed up by
      # hard numbers.
      model_id = random.choice( ( 1, 2 ) )

      # Specify the name of the output PDB
      outname = f"{ str_path }/{ mutant }_{ nseq }_{ n_model }.pdb"

      # Template-free prediction
      predict.predict_structure_no_templates( sequence, outname,
          a3m_lines, model_id = model_id, max_msa_clusters = nseq // 2,
          max_extra_msa = nseq, max_recycles = 1, n_struct_module_repeats = 8,
          ptm = True)
```
