## Supplementary material for "*Clostridioides difficile* Toxins Unhinged: Allosterically Switchable Network Orients *β*-flap": Scripts, Structures etc.: BasicMD.py

```
import sys
import numpy as np
import os
import argparse
import config.settings as c
from tools import utils, traj_funcs
import matplotlib.pyplot as plt
import matplotlib.patheffects as pe
import MDAnalysis as mda
from MDAnalysis.analysis import rms, align, dihedrals, contacts, \
                                distances, hydrogenbonds
from MDAnalysis.analysis.distances import distance_array
import pandas as pd

def main(argv):
    """Perform analysis for individual simulations.

    Calculations are generally stored as npy arrays which can be combined 
    with other simulations data in combination type analysis. The command
    line arguements are used to select a particular simulation from the 
    given options. 
    """

    try:
        parser = argparse.ArgumentParser()

        parser.add_argument("-p", "--path",
                            action = "store",
                            dest = "path",
                            default = "unbiased_sims/apo_open/nobackup",
                            help = "Set path to the data directory.")
        parser.add_argument("-r", "--recalc",
                            action = "store_true",
                            dest = "recalc",
                            default = False,
                            help = ("Chose whether the trajectory "
                                "arrays should be recomputed."))
        parser.add_argument("-s", "--state",
                            action = "store",
                            dest = "state",
                            default = "apo",
                            help = ("Chose the type of simulation i.e." 
                                "'holo' or 'apo' or in the case of "
                                "mutational, 'K57G' etc."))
        parser.add_argument("-f", "--figpath",
                            action = "store",
                            dest = "figpath",
                            default = None,
                            help = ("Set a path destination for the "
                                "figure."))    
        parser.add_argument("-t", "--topol",
                            action = "store",
                            dest = "topol",
                            default = "topol_protein.top",
                            help = ("File name for topology, inside the " 
                                "path directory."))   
        parser.add_argument("-x", "--xtc",
                            action = "store",
                            dest = "xtc",
                            default = "fitted_traj_100.xtc",
                            help = ("File name for trajectory, inside "
                                "the path directory."))
        args = parser.parse_args()

    except argparse.ArgumentError:
        print("Command line arguments are ill-defined, please check the " 
            "arguments")
        raise

    global conform, recalc, state, fig_path, data_path, np_files, analysis_path

    # Assign group selection from argparse
    recalc = args.recalc
    state = args.state
    data_path = f"{ c.data_head }/{ args.path }"
    fig_path = f"{ c.figure_head }/{ args.figpath }"
    topol = f"{ data_path }/{ args.topol }"
    xtc = f"{ data_path }/{ args.xtc }"

    # Check if topology file exists
    utils.process_topol(data_path, args.topol)

    # Load in universe objects for the simulation and the reference
    # structures
    u = mda.Universe(topol, xtc, topology_format='ITP')
    open_state = mda.Universe(f"{ c.struct_head }/open_ref_state.pdb")
    closed_state = mda.Universe(f"{ c.struct_head }/closed_ref_state.pdb")
    
    # For TcdA sims
    if "tcda" in state:
        tcda = True

    # Starting structure as the ref state
    ref_state = mda.Universe(f"{ c.struct_head }/alignment_struct.pdb")

    # Indicies of the inflexible residues
    core_res, core = traj_funcs.get_core_res()

    # Store calculated outputs as numpy arrays
    analysis_path = f"{ os.path.dirname(data_path) }/analysis"
    if not os.path.exists(analysis_path):
        os.makedirs(analysis_path)
    print(f"data path : { data_path }\n"
        f"fig path : { fig_path }\n"
        f"analysis path : { analysis_path }")

    # Analysis files stored as numpy arrays
    np_files = {"R_open" : f"{ analysis_path }/rmsd_open.npy", 
                "R_closed" : f"{ analysis_path }/rmsd_closed.npy",
                "rmsf" : f"{ analysis_path }/rmsf.npy", 
                "calphas" : f"{ analysis_path }/calphas.npy",
                "r_gyr" : f"{ analysis_path }/rad_gyration.npy",
                "rama" : f"{ analysis_path }/rama.npy", 
                "chi_1s" : f"{ analysis_path }/chi_1s.npy",
                "sc" : f"{ analysis_path }/salt.npy", 
                "salt_dist" : f"{ analysis_path}/salt_dist.npy",
                "hpairs" : f"{ analysis_path }/hpairs.npy", 
                # "hbond_count" : f"{ analysis_path }/hbond_count.npy",
                "time_ser" : f"{ analysis_path }/timeseries.npy"}

    # Store numpy arrays from analysis in the dictionary
    arrs = {}

    if all(list(map(lambda x : os.path.exists(x), np_files.values()))) and not recalc:

        print(
            "LOADING NUMPY ARRAYS"
        )

        for key, file in np_files.items(): 
            arrs[key] = np.load(file, allow_pickle=True)

    else:
    
        print(
            "EVALUATING WITH MDANALYSIS"
        )

        # Load in the trajectory and do alignment
        u.transfer_to_memory()
        u = traj_funcs.do_alignment(u)

        # Write out snapshots every 1µs to a pdb file
        protein = u.select_atoms("protein")
        with mda.Writer(f"{ analysis_path }/snap_shots.pdb", 
                        protein.n_atoms) as W:
            for ts in u.trajectory:
                if ((ts.time % 1000000) == 0):
                    W.write(protein)

        # Determine RMSD to ref structures
        arrs["R_open"] = get_rmsd(u, open_state, core, 
                            ["backbone and resid 8:251", 
                            c.beta_flap_group, 
                            c.alpha_flap_group],
                            "open")
        arrs["R_closed"] = get_rmsd(u, closed_state, core, 
                            ["backbone and resid 8:251",
                            c.beta_flap_group, 
                            c.alpha_flap_group], 
                            "closed")

        # Determine RMSF by first finding and aligning to the average 
        # structure
        arrs["calphas"], arrs["rmsf"] = get_rmsf(u, 
                                            f"{ data_path }/topol_protein.top",
                                            core_res)

        # Determine the radius of gyration and extract a time series of
        # the simulation
        arrs["r_gyr"], arrs["time_ser"] = get_rgyr(u)

        # # Phi and Psi backbone dihedrals
        resids = u.residues[195-1:232]
        rama = dihedrals.Ramachandran(resids).run()
        arrs["rama"] = rama.results.angles
        utils.save_array(np_files["rama"], arrs["rama"])
        
        # Chi1 dihedrals
        arrs["chi_1s"] = np.zeros((len(arrs["time_ser"]), len(resids)))
        for r, res in enumerate(resids):
            group = res.chi1_selection()
            if group is not None:
                dihs = dihedrals.Dihedral([group]).run()
                arrs["chi_1s"][:,r] = dihs.results.angles[:,0]
        utils.save_array(np_files["chi_1s"], arrs["chi_1s"])
        
        if tcda:

            print("Skipping salt-bridge and hpair time series")

        else: 

            # Salt-bridge contacts
            arrs["sc"] = get_salt_contacts(u, ref_state)
            arrs["salt_dist"] = get_bridges(u)

            # Hydrogen bonds analysis (intramolecular, not with solvent)
            arrs["hpairs"] = get_hbond_pairs(u)
            # arrs["hbond_count"] = get_hbonds(u)

    # Make plots for simulation analysis
    plot_rmsd(arrs["R_open"], arrs["R_closed"], arrs["time_ser"])

    plot_rmsf(arrs["calphas"], arrs["rmsf"])
    plot_rmsd_time(arrs["R_open"], arrs["time_ser"], ref_state="open")
    plot_rmsd_time(arrs["R_closed"], arrs["time_ser"], ref_state="closed")
    plot_rgyr(arrs["r_gyr"], arrs["time_ser"])

    # Plots for individual residues: Ramachandran and Chi_1
    for i, id in enumerate(np.arange(195-1,232)):
        res = u.residues[id]
        print(res)
        plot_ramachandran(arrs["rama"][:,i,0], arrs["rama"][:,i,1], res)
        plot_chi1(res, arrs["chi_1s"][:,i])

    if not tcda:
        plot_salt_frac(arrs["sc"], arrs["time_ser"])
        plot_bridges(arrs["salt_dist"], arrs["time_ser"])
        plot_hbond_pairs(arrs["hpairs"], arrs["time_ser"])
        # plot_hbonds_count(arrs["hbond_count"], arrs["time_ser"])

    return None

def get_rmsd(system, reference, alignment, group, ref_state):
    """Determines the rmsd over the trajectory against a reference structure.

    The MDAnalysis.analysis.rms.results array is saved as a numpy array file,
    which can be loaded if it has alreay been determined.

    Parameters
    ----------
    system : MDAnalysis.core.universe
        The universe object, with the trajectory loaded in.
    reference : MDAnalysis.core.universe
        The topology of the reference structure.
    alignment : MDAnalysis.core.groups.AtomGroup
        The group for the alignment, i.e. all the alpha carbons.
    group : MDAnalysis.core.groups.AtomGroup
        The group for the RMSD, e.g. the beta flap residues.
    ref_state : str
        The reference state as a conformational description, i.e. "open" or "closed".

    Returns
    -------
    rmsd_arr : np.ndarray
        A timeseries of the RMSD against the given reference, for the given
        atom group.
    """
    if type(group) != list:
        group = [group]
    R = rms.RMSD(system,
                 reference,  # reference universe or atomgroup
                 select=alignment,  # group to superimpose and calculate RMSD
                 groupselections=group)  # groups for RMSD
    R.run()

    rmsd_arr = R.results.rmsd

    if ref_state == "open":
        utils.save_array(np_files["R_open"], rmsd_arr)
    elif ref_state == "closed":
        utils.save_array(np_files["R_closed"], rmsd_arr)

    return rmsd_arr

def get_rmsf(u, top, core_res, get_core=False):
    """Determines the RMSF per residue.

    The average structure is calculated and then used as the reference for
    structure alignment, before calculating the rmsf per residue.

    Parameters
    ----------
    u : MDAnalysis.core.universe
        The universe object, with the trajectory loaded in.
    top : str
        The topology file, e.g. xxx.gro.
    ref_group : str
        Atom group selection string of the core residues for the alignment.

    Returns
    -------
    resnums : MDAnalysis.core.groups.AtomGroup
        The atom group used for the RMSF calculation consists of all the alpha
        carbons in the protein.
    rmsf : np.ndarray
        The rmsf of the selected atom groups.

    """
    # Perform alignment to the average structure and write to a new traj file.
    protein = u.select_atoms("protein")
    prealigner = align.AlignTraj(u, u, select="protein and name CA",
                                 in_memory=True).run()
    ref_coords = u.trajectory.timeseries(asel=protein).mean(axis=1)
    ref = mda.Merge(protein).load_new(ref_coords[:, None, :],
                              order="afc")
    aligner = align.AlignTraj(u, ref, select="protein and name CA", in_memory=True).run()
    with mda.Writer(f"{ data_path }/rmsfit.xtc", n_atoms=u.atoms.n_atoms) as W:
        for ts in u.trajectory:
            W.write(u.atoms)

    from MDAnalysis.analysis.rms import RMSF

    u = mda.Universe(top, f"{ data_path }/rmsfit.xtc", topology_format='ITP')
    calphas = protein.select_atoms("protein and name CA")

    rmsfer = RMSF(calphas).run()

    if not get_core:

        utils.save_array(np_files["calphas"], calphas.resnums)
        utils.save_array(np_files["rmsf"], rmsfer.results.rmsf)

    return calphas.resnums, rmsfer.results.rmsf

def get_salt_contacts(u, ref_state):
    """Runs the Contacts module to get the number of salt contacts.

    Arginine and lysine residues are considered as the basic residues, while
    aspargine and glutamine are classified as acidic residues. A soft cut off
    is used for determining contacts within a radius of 4.5 A.

    Parameters
    ----------
    u : MDAnalysis.core.universe.Universe
        The universe object, with the trajectory loaded in.
    ref_state : MDAnalysis.core.universe.Universe
        The universe object for the reference state.

    Returns
    -------
    sc : np.ndarray
        A timeseries of the fraction of retained salt bridges, with respect to
        the reference state.
    """
    sel_basic = "(resname ARG LYS) and (name NH* NZ*)"
    sel_acidic = "(resname ASP GLU) and (name OE* OD*)"
    acidic = ref_state.select_atoms(sel_acidic)
    basic = ref_state.select_atoms(sel_basic)
    salt_contacts = contacts.Contacts(u, select=(sel_acidic, sel_basic),
                    refgroup=(acidic, basic), radius=4.5, method="soft_cut")

    salt_contacts.run()
    sc = salt_contacts.results.timeseries

    utils.save_array(np_files["sc"], sc)

    return sc

def get_bridges(u):
    """Calculates the distance of special salt contacts in the alpha + beta flaps.

    Arginine and lysine residues are considered as the basic residues, while
    aspargine and glutamine are classified as acidic residues.

    Parameters
    ----------
    u : MDAnalysis.core.universe.Universe
        The universe object, with the trajectory loaded in.

    Returns
    -------
    distances : np.ndarray
        Trajectory of the four distance arrays. 

    """
    acid_base_pairs = [
        (("resid 208 and name NH*"),("resid 222 and name OE*")), # R208 -- E222
        (("resid 227 and name NZ*"),("resid 213 and name OD*")), # K227 -- D213
        #(("resid 232 and name NZ*"),("resid 149 and name OE*")), # K232 -- E149
        (("resid 202 and name NH*"),("resid 210 and name OE*")), # R202 -- E210
        (("resid 57 and (name NZ* or (resname GLY and name CA))"),
         ("resid 200 and (name OE* or (resname GLY and name CA))"))] # K57 -- E200

    pairs = []

    for b, a in acid_base_pairs:
        sel_basic = u.select_atoms(b)
        sel_acidic = u.select_atoms(a)
        pairs.append((sel_basic, sel_acidic))

        #dist_pair1 = distance_array(u.coord[ca_pair1_1], u.coord[ca_pair1_2])
    sig_bridges = 4
    distances = np.zeros((u.trajectory.n_frames, sig_bridges))

    # Loop over all frames in the trajectory
    for ts in u.trajectory:
        # Calculate the distances between the four acid-base pairs for this frame
        d1 = distance_array(pairs[0][0].positions, pairs[0][1].positions)
        d2 = distance_array(pairs[1][0].positions, pairs[1][1].positions)
        d3 = distance_array(pairs[2][0].positions, pairs[2][1].positions)
        d4 = distance_array(pairs[3][0].positions, pairs[3][1].positions)
        distances[ts.frame] = [np.min(d1), np.min(d2), np.min(d3), np.min(d4)]

    utils.save_array(np_files["salt_dist"], distances)

    return distances

def get_hbonds(u):
    """Find the intramolecular hydrogen bonds in the simulation.

    The donor-acceptor distance cutoff is 3 A and the donor-hydrogen-acceptor
    angle cutoff is minimum 150 deg. The function generates two files, one to
    track the types of hbonds formed and one to identify significant and
    persistent bonds.

    Parameters
    ----------
    u : MDAnalysis.core.universe
        The universe object, with the trajectory loaded in.

    Returns
    -------
    hbonds : MDAnalysis...HydrogenBondAnalysis
        Each row consists of a particular h-bond observation: [frame,
        donor_index, hydrogen_index, acceptor_index, DA_distance, DHA_angle].
    hbond_count : np.ndarray
        The hbond count as a trajectory.

    """
    hbonds = hydrogenbonds.hbond_analysis.HydrogenBondAnalysis(universe=u)
    hbonds.hydrogens_sel = hbonds.guess_hydrogens("protein")
    hbonds.acceptors_sel = hbonds.guess_acceptors("protein")
    hbonds.run()
    tau_timeseries, timeseries = hbonds.lifetime()

    # these get their own np arrays
    times = hbonds.times
    hbond_count = hbonds.count_by_time()

    # Analyze the bond types
    f = open(f"{ analysis_path }/hbond_types.txt", "w")
    f.write("The average number of each type of hydrogen bond formed at each"
            " frame:\n")

    for donor, acceptor, count in hbonds.count_by_type():

        donor_resname, donor_type = donor.split(":")
        n_donors = u.select_atoms(f"resname { donor_resname } " \
                                  f"and type { donor_type }").n_atoms

        # average number of hbonds per donor molecule per frame
        # multiply by two as each hydrogen bond involves two water molecules
        mean_count = 2 * int(count) / (hbonds.n_frames * n_donors)
        if mean_count > 0.1:
            f.write(f"{donor} to {acceptor}: {mean_count:.2f}\n")
    f.close()

    f = open(f"{ analysis_path }/significant_hbonds.txt", "w")
    f.write("A descending list of h-bonds formed between atom pairs.\n"\
            "Donor\tHydrogen\tAcceptor\tCount\tFlaps?\n")

    def describe_at(atom):
        a = u.atoms[atom-1]
        return f"{ a.resname }{ a.resid }--{ a.type }"

    # List the most important h-bonds by order of occurence
    c = hbonds.count_by_ids()
    for d, h, a, count in c:
        if count < 10:
            continue
        if d.resid or a.resid in np.arange(195,232 + 1):
            in_flap = True
        else:
            in_flap = False
        f.write(f"{ describe_at(d) }\t{ describe_at(h) }\t{ describe_at(a) } "\
                f"\t{ count }\t{ in_flap }\n")
    f.close()

    utils.save_array(np_files["hbond_count"], hbond_count)

    return hbond_count

def get_hbond_pairs(u):
    """Calculates the distance of special hbond contacts in the alpha + beta flaps.

    Parameters
    ----------
    u : MDAnalysis.core.universe.Universe
        The universe object, with the trajectory loaded in.

    Returns
    -------
    distances : np.ndarray
        Trajectory of the four distance arrays. 

    """
    hbond_pairs = [
        (("resid 204 and name OD1"),("(resid 206 or resid 208) and name H")),
        (("resid 231 and name O*"),("resid 209 and name NH*")),
        (("resid 197 and name OD1"),("resid 209 and name N*")),
        (("resid 53 and name H*"),("resid 200 and name O*"))]

    pairs = []

    for a, d in hbond_pairs:
        sel_accept = u.select_atoms(a)
        sel_donor = u.select_atoms(d)
        pairs.append((sel_accept, sel_donor))

    distances = np.zeros((u.trajectory.n_frames, len(pairs)))

    # Loop over all frames in the trajectory
    for ts in u.trajectory:
        # Calculate the distances between the 3 hbond pairs for this frame
        d1 = distance_array(pairs[0][0].positions, pairs[0][1].positions)
        d2 = distance_array(pairs[1][0].positions, pairs[1][1].positions)
        d3 = distance_array(pairs[2][0].positions, pairs[2][1].positions)
        d4 = distance_array(pairs[3][0].positions, pairs[3][1].positions)

        # Store the distances in the distances array
        distances[ts.frame] = [np.min(d1), np.min(d2), np.min(d3), np.min(d4)]

    utils.save_array(np_files["hpairs"], distances)

    return distances

def get_rgyr(u):
    """Determine the radius of gyration as a timeseries.

    The radius of gyration is a measure of how compact the 
    structure is, such that an increase may indicate unfolding 
    or opening.

    Parameters
    ----------
    u : MDAnalysis.core.universe.Universe
        The universe object, with the trajectory loaded in.

    Returns
    -------
    r_gyr : nd.nparray
        The radius of gyration in units of Angstrom.
    time_ser : nd.nparray
        A time series of the trajectory in ps. 
    
    """
    r_gyr = []
    time_ser = []
    protein = u.select_atoms("protein")
    for ts in u.trajectory:
       r_gyr.append(protein.radius_of_gyration())
       time_ser.append(ts.time)
    r_gyr = np.array(r_gyr)
    time_ser = np.array(time_ser)

    utils.save_array(np_files["r_gyr"], r_gyr)
    utils.save_array(np_files["time_ser"], time_ser)

    return r_gyr, time_ser

def plot_rmsd(r_open, r_closed, time_ser): 
    """Makes a plot of the rmsd against two ref structures with a color bar.

    Parameters
    ----------
    r_open : MDAnalysis.analysis.rms.RMSD or nparray
        A time series of the rmsd with the open crystal structure as reference.
    r_closed : MDAnalysis.analysis.rms.RMSD or nparray
        A time series of the rmsd with the closed crystal structure as reference.
    time_ser : np.ndarray
        A time series of the simulation time in ps.

    Returns
    -------
    None.

    """
    fig, ax = plt.subplots(constrained_layout=True, figsize=(12,8))

    ax.scatter(r_open[:,3], r_closed[:,3], c=time_ser,
                cmap="cividis", label=r"Full backbone", marker="s", s=300, 
                edgecolors="#404040")
    ax.scatter(r_open[:,5], r_closed[:,5], c=time_ser,
                cmap="cividis", label=r"$\alpha$-flap", marker="D", s=300,
                edgecolors="#404040")
    d = ax.scatter(r_open[:,4], r_closed[:,4], c=time_ser,
                cmap="cividis", label=r"$\beta$-flap", marker="o", s=300,
                edgecolors="#404040")

    print(len(time_ser))

    # Colormap settings
    cbar = plt.colorbar(d)
    num_ticks = 10
    cticks = time_ser[::(len(time_ser) // num_ticks)]
    cbar.ax.yaxis.set_ticks(cticks)
    if time_ser[-1] > 1e6:
        # Use microseconds for time labels
        cbar.set_label(r'Time ($\mu$s)', fontsize=32, labelpad=10)
        ctick_lab = list(map(lambda x: str(x/1e6).split(".")[0], cticks))
    else:
        # Use nanoseconds for time labels
        cbar.set_label(r'Time (ns)', fontsize=32, labelpad=10)
        ctick_lab = list(map(lambda x: str(x/1e3).split(".")[0], cticks))
    cbar.ax.yaxis.set_ticklabels(ctick_lab)
    cbar.ax.tick_params(labelsize=24, direction='out', width=2, length=5)
    cbar.outline.set_linewidth(2)

    # Plot settings
    ax.tick_params(axis='y', labelsize=24, direction='in', width=2, \
                    length=5, pad=10)
    ax.tick_params(axis='x', labelsize=24, direction='in', width=2, \
                    length=5, pad=10)
    for i in ["top","bottom","left","right"]:
        ax.spines[i].set_linewidth(2)
    ax.grid(True)
    ax.set_xlabel(r"RMSD to open conformation ($\AA$)", labelpad=5, \
                    fontsize=32)
    ax.set_ylabel(r"RMSD to closed conformation ($\AA$)", labelpad=5, fontsize=32)
    _, xmax = ax.get_xlim()
    _, ymax = ax.get_ylim()
    ax.set_xlim(0,15)
    ax.set_ylim(0,15)

    plt.legend(fontsize=28)

    #plt.savefig(f"{ fig_path }/rmsd_2D.png", dpi=300)
    utils.save_figure(fig, f"{ fig_path }/rmsd_2D.png")
    plt.close()

    return None

def plot_rmsf(calphas, rmsf):
    """Makes an RMSF plot.

    RMSF helps to understand which residues in the protein tend to undergo
    large fluctuations and which residues may be part of a more stable core 
    structure.

    Parameters
    ----------
    resnums : MDAnalysis.core.groups.AtomGroup
        The atom group used for the RMSF calculation consists of all the 
        alpha carbons in the protein.
    rmsf : np.ndarray
        The rmsf of the selected atom groups.

    Returns
    -------
    None.

    """
    fig, ax = plt.subplots(constrained_layout=True, figsize=(12,4))
    #resids = list(map(lambda x : x + 544, calphas))
    plt.plot(calphas, rmsf, lw=3, color="#47abc4")

    # Plot settings
    ax.tick_params(axis='y', labelsize=18, direction='in', width=2, \
                    length=5, pad=10)
    ax.tick_params(axis='x', labelsize=18, direction='in', width=2, \
                    length=5, pad=10)
    for i in ["top","bottom","left","right"]:
        ax.spines[i].set_linewidth(2)
    ax.grid(True)
    ax.set_xlabel(r"Residue number", labelpad=5, fontsize=28)
    ax.set_ylabel(r"RMSF ($\AA$)", labelpad=5, fontsize=28)
    bottom, top = ax.get_ylim()
    ax.vlines([195,217.5], bottom, top, linestyles="dashed", lw=3,
              colors="#FF1990")
    ax.vlines([219.5,231], bottom, top, linestyles="dashed", lw=3,
              colors="#dba61f")
    ax.set_ylim(-1,10)

    utils.save_figure(fig, f"{ fig_path }/rmsf.png")
    plt.close()

    return None

def plot_rmsd_time(rmsd, time_ser, ref_state, stride=10):
    """Makes a time series of the RMSD against a reference.

    The RMSD is shown for the protein backbone as well as the backbone of the
    alpha and beta flaps.

    Parameters
    ----------
    r_open : MDAnalysis.analysis.rms.RMSD or nparray
        A time series of the rmsd with the open crystal structure as reference.
    time_ser : np.ndarray
        A time series of the simulation time in ps.
    ref_state : str
        Name of the reference structure used in the RMSD computation, i.e. "open" 
        or "closed".

    Returns
    -------
    None.

    """
    fig, ax = plt.subplots(constrained_layout=True, figsize=(24,8))
    #plt.plot(time, rmsd[:,2], lw=2, color="#02ab64", alpha=0.8,
    #         label="core")
    plt.plot(time_ser[::stride], rmsd[:,5][::stride], lw=3, color="#02ab64", alpha=0.8,
             label=r"$\alpha-$flap")
    plt.plot(time_ser[::stride], rmsd[:,4][::stride], lw=3, color="#dba61f", alpha=0.8,
             label=r"$\beta-$flap")
    plt.plot(time_ser[::stride], rmsd[:,3][::stride], lw=3, color="#47abc4", alpha=0.8,
             label="full backbone")

    # Plot settings
    ax.tick_params(axis='y', labelsize=32, direction='in', width=2, \
                    length=5, pad=10)
    ax.tick_params(axis='x', labelsize=32, direction='in', width=2, \
                    length=5, pad=10)
    num_ticks = 10
    xticks = time_ser[::(len(time_ser) // num_ticks)]
    ax.set_xticks(xticks)
    if time_ser[-1] > 1e6:
        # Use microseconds for time labels
        ax.set_xlabel(r'Time ($\mu$s)', fontsize=38, labelpad=5)
        ax.set_xticklabels(list(map(lambda x : str(x/1e6).split(".")[0], xticks)))
    else:
        # Use nanoseconds for time labels
        ax.set_xlabel(r'Time (ns)', fontsize=38, labelpad=5)
        ax.set_xticklabels(list(map(lambda x : str(x/1e3).split(".")[0], xticks)))
    for i in ["top","bottom","left","right"]:
        ax.spines[i].set_linewidth(2)
    ax.grid(True)
    ax.set_ylabel(f"RMSD$_{ {ref_state} }$ ($\AA$)", labelpad=5, fontsize=38)
    _, ymax = ax.get_ylim()
    ax.set_ylim(0,ymax)
    plt.legend(fontsize=28)

    utils.save_figure(fig, f"{ fig_path }/rmsd_time_{ ref_state }.png")
    plt.close()

    return None

def plot_ramachandran(phis, psis, res):
    """Makes a Ramachandran plot for the given residue.

    A separate Ramachandran plot should be used for each of the 
    residues under consideration. 

    Parameters
    ----------
    dihs_phi : np.ndarray
        A timeseries of the phi dihedrals for the residue group.
    dihs_psi : np.ndarray
        A timeseries of the psi dihedrals for the residue group.
    resnum : MDAnalysis.core.groups.Residue
        The residue object for the ramachandran plot.

    Returns
    -------
    None.

    """
    fig, ax = plt.subplots(constrained_layout=True, figsize=(12,8))

    d = ax.scatter(phis, psis, c=np.arange(0,len(phis)), cmap="cividis",
                   label=f"{ res.resname } { res.resid }")

    # Colormap settings
    cbar = plt.colorbar(d)
    cbar.set_label(r'Time [$\mu$s]', fontsize=28, labelpad=10)
    cbar.ax.yaxis.set_ticks(np.arange(0,len(psis),1000))
    cticks = list(map(lambda x: str(x/1000).split(".")[0],
                      np.arange(0,len(psis),1000)))
    cbar.ax.yaxis.set_ticklabels(cticks)
    cbar.ax.tick_params(labelsize=16, direction='out', width=2, length=5)
    cbar.outline.set_linewidth(2)

    # Plot settings
    ax.tick_params(axis='y', labelsize=18, direction='in', width=2, \
                    length=5, pad=10)
    ax.tick_params(axis='x', labelsize=18, direction='in', width=2, \
                    length=5, pad=10)
    for i in ["top","bottom","left","right"]:
        ax.spines[i].set_linewidth(2)
    ax.grid(True)
    ax.set_xlabel(r"$\Phi$", labelpad=5, \
                    fontsize=28)
    ax.set_ylabel(r"$\Psi$", labelpad=5, fontsize=28)
    plt.xlim([-180, 180])
    plt.ylim([-180, 180])
    plt.legend(fontsize=20)

    # Save plot to file
    plot_path = f"{ fig_path }/ramachandran"
    if not os.path.exists(plot_path):
       os.makedirs(plot_path)
    utils.save_figure(fig, f"{ plot_path }/res_{ res.resid }.png")
    plt.close()

    return None

def plot_chi1(res, chi_1):
    """Makes a chi_1 plot for the given residue.

    A separate Ramachandran plot should be used for each of the 
    residues under consideration. 

    Parameters
    ----------
    res : MDAnalysis.core.groups.Residue
        The residue object for the chi_1 plot.
    chi_1 : np.ndarray
        A timeseries of the chi 1 dihedrals for the residue group.

    Returns
    -------
    None.

    """
    fig, ax = plt.subplots(constrained_layout=True, figsize=(12,8))

    d = ax.hist(chi_1, bins=180, density=True, color="#b21856",
                label=f"{ res.resname } { res.resid }")

    # Plot settings
    ax.tick_params(axis='y', labelsize=18, direction='in', width=2, \
                    length=5, pad=10)
    ax.tick_params(axis='x', labelsize=18, direction='in', width=2, \
                    length=5, pad=10)
    for i in ["top","bottom","left","right"]:
        ax.spines[i].set_linewidth(2)
    ax.grid(True)
    ax.set_xlabel(r"$\chi_1$ (°)", labelpad=5, \
                    fontsize=28)
    ax.set_ylabel(r"Density", labelpad=5, fontsize=28)
    plt.xlim([-180, 180])
    plt.legend(fontsize=20)

    plot_path = f"{ fig_path }/chi_1s"
    if not os.path.exists(plot_path):
       os.makedirs(plot_path)
    utils.save_figure(fig, f"{ plot_path }/res_{ res.resid }.png")
    plt.close()

    return None

def plot_salt_frac(sc, time_ser):
    """Makes a plot for the fraction of salt contacts compared to the reference.

    A soft cut-off from 4 A was used to determine the fraction of salt bridges
    retained from the reference structure. This includes bridges between the
    charged residues, i.e. N in ARG and LYS with O in ASP and GLU.

    Parameters
    ----------
    sc : np.ndarray
        A timeseries of the fraction of salt bridge contacts, relative to the
        reference structure.
    time_ser : np.ndarray
        A time series of the simulation time in ps.

    Returns
    -------
    None.

    """
    fig, ax = plt.subplots(constrained_layout=True, figsize=(12,8))
    ave_contacts = np.mean(sc[:, 1])
    print(f"average contacts = { ave_contacts }")
    stride = 1

    ax.scatter(time_ser[::stride], sc[::stride, 1], s=150, color="#00C27B")

    # Plot settings
    ax.tick_params(axis='y', labelsize=18, direction='in', width=2, \
                    length=5, pad=10)
    ax.tick_params(axis='x', labelsize=18, direction='in', width=2, \
                    length=5, pad=10)
    num_ticks = 10
    xticks = time_ser[::(len(time_ser) // num_ticks)]
    ax.set_xticks(xticks)
    if time_ser[-1] > 1e6:
        # Use microseconds for time labels
        ax.set_xlabel(r'Time ($\mu$s)', fontsize=24, labelpad=5)
        ax.set_xticklabels(list(map(lambda x : str(x/1e6).split(".")[0], xticks)))
    else:
        # Use nanoseconds for time labels
        ax.set_xlabel(r'Time (ns)', fontsize=24, labelpad=5)
        ax.set_xticklabels(list(map(lambda x : str(x/1e3).split(".")[0], xticks)))
    for i in ["top","bottom","left","right"]:
        ax.spines[i].set_linewidth(2)
    ax.grid(True)
    ax.set_ylabel(r"Fraction of Native Contacts", labelpad=5, fontsize=24)
    plt.ylim([0, 1])

    utils.save_figure(fig, f"{ fig_path }/salt_contacts.png")
    plt.close()

    return None

def plot_hbonds_count(hbond_count, time_ser):
    """Makes a plot for the number of hbonds at each time step.

    Parameters
    ----------
    hbond_count : np.ndarray
        A trajectory of the hydrogen bond count.
    time_ser : np.ndarray
        A time series of the simulation time in ps.

    Returns
    -------
    None.

    """
    fig, ax = plt.subplots(constrained_layout=True, figsize=(12,8))
    stride = 1

    plt.scatter(time_ser[::stride], hbond_count[::stride], s=150, color="#00C27B")

    # Plot settings
    ax.tick_params(axis='y', labelsize=18, direction='in', width=2, \
                    length=5, pad=10)
    ax.tick_params(axis='x', labelsize=18, direction='in', width=2, \
                    length=5, pad=10)
    num_ticks = 10
    xticks = time_ser[::(len(time_ser) // num_ticks)]
    ax.set_xticks(xticks)
    if time_ser[-1] > 1e6:
        # Use microseconds for time labels
        ax.set_xlabel(r'Time ($\mu$s)', fontsize=24, labelpad=5)
        ax.set_xticklabels(list(map(lambda x : str(x/1e6).split(".")[0], xticks)))
    else:
        # Use nanoseconds for time labels
        ax.set_xlabel(r'Time (ns)', fontsize=24, labelpad=5)
        ax.set_xticklabels(list(map(lambda x : str(x/1e3).split(".")[0], xticks)))
    for i in ["top","bottom","left","right"]:
        ax.spines[i].set_linewidth(2)
    ax.grid(True)
    ax.set_ylabel(r"$N_{HB}$", labelpad=5, fontsize=24)
    plt.ylim([0,100])

    utils.save_figure(fig, f"{ fig_path }/hbonds_count.png")
    plt.close()

    return None

def plot_rgyr(r_gyr, time_ser, stride=10):
    """Makes a timeseries of the radius of gyration.

    The radius of gyration is a measure of how compact the 
    structure is, such that an increase my indicate unfolding 
    or opening.

    Parameters
    ----------
    r_gyr : np.ndarray
        The radius of gyration in units of Angstrom.
    time_ser : np.ndarray
        A time series of the simulation time in ps.
    """
    fig, ax = plt.subplots(constrained_layout=True, figsize=(24,8))

    plt.plot(time_ser[::stride], r_gyr[::stride], "--", lw=3, color="#02ab64", label=r"$R_G$")

    # Plot settings
    ax.tick_params(axis='y', labelsize=32, direction='in', width=2, \
                    length=5, pad=10)
    ax.tick_params(axis='x', labelsize=32, direction='in', width=2, \
                    length=5, pad=10)
    
    num_ticks = 10
    xticks = time_ser[::(len(time_ser) // num_ticks)]
    ax.set_xticks(xticks)
    if time_ser[-1] > 1e6:
        # Use microseconds for time labels
        ax.set_xlabel(r'Time ($\mu$s)', fontsize=24, labelpad=5)
        ax.set_xticklabels(list(map(lambda x : str(x/1e6).split(".")[0], xticks)))
    else:
        # Use nanoseconds for time labels
        ax.set_xlabel(r'Time (ns)', fontsize=24, labelpad=5)
        ax.set_xticklabels(list(map(lambda x : str(x/1e3).split(".")[0], xticks)))
    for i in ["top","bottom","left","right"]:
        ax.spines[i].set_linewidth(2)
    ax.grid(True)
    ax.set_xlabel(r"time ($\mu s$)", labelpad=5, fontsize=38)
    ax.set_ylabel(r"Radius of Gyration $R_G$ ($\AA$)", labelpad=5, fontsize=38)
    plt.legend(fontsize=28)

    utils.save_figure(fig, f"{ fig_path }/rad_gyration.png")
    plt.close()

    return None

def plot_bridges(salt_dist, time_ser, stride=10):
    """Makes a time series plot of the distances of notable bridges.

    The plot includes a dotted line to indicate the upper limit of a typical 
    salt bridge interaction.

    Parameters
    ----------
    hpairs : np.ndarray
        A time series of the four distance arrays for notable salt bridges in the beta
        flap, alpha flap and the IP6 binding pocket.
    time_ser : np.ndarray
        A time series of the simulation time in ps.

    Returns
    -------
    None.

    """
    fig, ax = plt.subplots(constrained_layout=True, figsize=(12,4))

    colors = ["#1f77b4", "#ff7f0e", "#2ca02c", "#9467bd"]
    labels = [r"$\alpha - \beta \:$ R208$-$E222",
              r"$\alpha - \beta \:$ K227$-$D213",
              #r"$\alpha - pocket \:$ K232-E149",
              r"$\beta - \beta \:$ R202$-$E210",
              r"Pocket$-\beta$ K57$-$E200"]

    for i in range(len(salt_dist[0,:])):

        plt.plot(time_ser[::stride], salt_dist[::stride,i], "-", lw=3, color=colors[i], 
            label=labels[i], alpha=0.8,
            path_effects=[pe.Stroke(linewidth=5, foreground='#595959'), pe.Normal()])

    ax.axhline(y=4.5, linestyle='--', lw=3, color='red', alpha=0.5,
        path_effects=[pe.Stroke(linewidth=5, foreground='#595959'), pe.Normal()])

    # Plot settings
    ax.tick_params(axis='y', labelsize=18, direction='in', width=2, \
                    length=5, pad=10)
    ax.tick_params(axis='x', labelsize=18, direction='in', width=2, \
                    length=5, pad=10)
    num_ticks = 10
    xticks = time_ser[::(len(time_ser) // num_ticks)]
    ax.set_xticks(xticks)
    if time_ser[-1] > 1e6:
        # Use microseconds for time labels
        ax.set_xlabel(r'Time ($\mu$s)', fontsize=24, labelpad=5)
        ax.set_xticklabels(list(map(lambda x : str(x/1e6).split(".")[0], xticks)))
    else:
        # Use nanoseconds for time labels
        ax.set_xlabel(r'Time (ns)', fontsize=24, labelpad=5)
        ax.set_xticklabels(list(map(lambda x : str(x/1e3).split(".")[0], xticks)))
    for i in ["top","bottom","left","right"]:
        ax.spines[i].set_linewidth(2)
    ax.grid(True)
    ax.set_ylabel(r"Distance ($\AA$)", labelpad=5, fontsize=24)
    plt.ylim([0, 40])
    _, xmax = ax.get_xlim()
    ax.set_xlim(0,xmax)
    plt.legend(fontsize=24, ncol=2)

    utils.save_figure(fig, f"{ fig_path }/salt_bridges.png")
    plt.close()

    return None

def plot_hbond_pairs(hpairs, time_ser, stride=10):
    """Makes a time series plot of the distances of notable H-bonding pairs.

    The plot includes a dotted line to indicate the upper limit of a typical 
    hydrogen bond.

    Parameters
    ----------
    hpairs : np.ndarray
        A time series of the four distance arrays for notable H-bonds in the beta
        flap, alpha flap and the IP6 binding pocket.
    time_ser : np.ndarray
        A time series of the simulation time in ps.

    Returns
    -------
    None.

    """
    fig, ax = plt.subplots(constrained_layout=True, figsize=(12,4))

    colors = ["#1f77b4", "#ff7f0e", "#2ca02c", "#9467bd"]
    labels = [r"N204$-$R208", r"S231$-$R209",
              r"N197$-$R209", r"N53$-$E200"]

    for i in range(4):

        plt.plot(time_ser[::stride], hpairs[::stride,i], "-", lw=3, color=colors[i], 
            label=labels[i], alpha=0.8,
            path_effects=[pe.Stroke(linewidth=5, foreground='#595959'), pe.Normal()])

    ax.axhline(y=4.5, linestyle='--', lw=3, color='red', alpha=0.5,
        path_effects=[pe.Stroke(linewidth=5, foreground='#595959'), pe.Normal()])

    # Plot settings
    ax.tick_params(axis='y', labelsize=18, direction='in', width=2, \
                    length=5, pad=10)
    ax.tick_params(axis='x', labelsize=18, direction='in', width=2, \
                    length=5, pad=10)
    num_ticks = 10
    xticks = time_ser[::(len(time_ser) // num_ticks)]
    ax.set_xticks(xticks)
    if time_ser[-1] > 1e6:
        # Use microseconds for time labels
        ax.set_xlabel(r'Time ($\mu$s)', fontsize=24, labelpad=5)
        ax.set_xticklabels(list(map(lambda x : str(x/1e6).split(".")[0], xticks)))
    else:
        # Use nanoseconds for time labels
        ax.set_xlabel(r'Time (ns)', fontsize=24, labelpad=5)
        ax.set_xticklabels(list(map(lambda x : str(x/1e3).split(".")[0], xticks)))
    for i in ["top","bottom","left","right"]:
        ax.spines[i].set_linewidth(2)
    ax.grid(True)
    ax.set_ylabel("Distance ($\AA$)", labelpad=5, fontsize=24)
    plt.ylim([0, 30])
    _, xmax = ax.get_xlim()
    ax.set_xlim(0,xmax)
    plt.legend(fontsize=24, ncol=2)

    utils.save_figure(fig, f"{ fig_path }/hpairs.png")
    plt.close()

    return None

if __name__ == '__main__':
    main(sys.argv)
```
