## Supplementary material for "*Clostridioides difficile* Toxins Unhinged: Allosterically Switchable Network Orients *β*-flap": Scripts, Structures etc.: ExtractConforms.py

```
import sys
import os
import pandas as pd
import subprocess
import time
import argparse
import MDAnalysis as mda
import numpy as np
import matplotlib.pyplot as plt
sys.path.insert(0, "/home/lf1071fu/project_b3/ProjectB3")
import config.settings as config
from tools import utils, traj_funcs

def main(argv):

    try:
        parser = argparse.ArgumentParser()

        parser.add_argument("-p", "--path",
                            action = "store",
                            dest = "paths",
                            nargs='+',
                            default = ("unbiased_sims/holo_open/nobackup"
                                " unbiased_sims/holo_closed/nobackup"),
                            help = "Set path to the data directory.")
        parser.add_argument("-r", "--recalc",
                            action = "store_true",
                            dest = "recalc",
                            default = False,
                            help = ("Chose whether the trajectory arrays "
                                "should  be recomputed."))
        parser.add_argument("-f", "--figpath",
                            action = "store",
                            dest = "fig_path",
                            default = "",
                            help = """Set a path destination for the 
                                figure.""")
        parser.add_argument("-t", "--topols",
                            action = "store",
                            dest = "topols",
                            nargs = "+",
                            default = "holo_open.tpr holo_closed.tpr",
                            help = """File name for topologies, inside the 
                                path directory.""")   
        parser.add_argument("-x", "--xtc",
                            action = "store",
                            dest = "xtc",
                            default = "fitted_traj_100.xtc",
                            help = """File name for trajectory, inside 
                                the path directory.""")
        parser.add_argument("-w", "--workdir",
                            action = "store",
                            dest = "workdir",
                            default = "umbrella/holo_state",
                            help = "Main directory head for the FES.")      
        parser.add_argument("-d", "--dataframe",
                            action = "store",
                            dest = "df",
                            default = "restraint_pts.csv",
                            help = ("File name for table of restraint "
                                "points."))   
        parser.add_argument("-a", "--alphafold",
                            action = "store_true",
                            dest = "alphafold",
                            default = False,
                            help = "Include alpha fold trajectories.")   
        parser.add_argument("-c", "--cat_trajs",
                            action = "store",
                            dest = "cat_traj_path",
                            default = "dataframe_beta_vec_apo-biased.csv",
                            help = ("Name of the .csv file with reaction"
                                " coordinate data."))   
        parser.add_argument("-i", "--initial_conforms",
                            action = "store_true",
                            dest = "initial_conforms",
                            default = False,
                            help = ("Use script to find initial conforms"
                                "for umbrella sampling?"))      
        parser.add_argument("-s", "--state",
                            action = "store",
                            dest = "state",
                            default = "holo",
                            help = ("Select ligand binding state as, apo"
                                " or holo"))                 
        args = parser.parse_args()

    except argparse.ArgumentError:
        print("""Command line arguments are ill-defined, please check the
              arguments""")
        raise

    global traj_paths, top_paths, state, index, wrkdir

    # Assign group selection from argparse 
    wrkdir = f"{ config.data_head }/{ args.workdir }"
    data_paths = [f"{ config.data_head }/{ p }" for p in args.paths]
    fig_path = f"{ config.figure_head }/{ args.fig_path }"
    recalc = args.recalc
    topols = args.topols
    xtc = args.xtc
    df_path = f"{ wrkdir }/{ args.df }"
    cat_traj_path = args.cat_traj_path
    alphafold = args.alphafold
    initial_conforms = args.initial_conforms
    state = args.state

    index = (f"{ config.data_head }/unbiased_sims/holo_closed/"
            "nobackup/index.ndx")

    # Check for valid paths
    for p in (data_paths + [fig_path] + [df_path]):
        utils.validate_path(p)

    # Alignment and reference structures
    refs = {"open" : f"{ config.struct_head }/open_ref_state.pdb",
            "closed" : f"{ config.struct_head }/closed_ref_state.pdb"}
    align = False
    if align:
        plumed_align_ref(wrkdir)
        for ref, ref_path in refs.items():
            traj_funcs.align_refs(ref, ref_path)

    # Get a table of all the sampled reaction coordinates
    df_cat_path = f"{ config.data_head }/cat_trajs/{ cat_traj_path }"
    df_cat = pd.read_csv(df_cat_path)

    # Determine the desired points in the reaction coordinate space
    df_pts = pd.read_csv(df_path)
    
    # Collect paths for trajectories which may be extracted from
    traj_paths, top_paths = sim_data_paths(data_paths, topols, xtc, 
                                           alphafold=alphafold)

    print("traj-paths", traj_paths)

    # Find the closest sampled structure for each point in df_pts
    if state == "salt_bridge":
        df_pts = find_nearest_sb(df_pts, df_cat)

    else:

        df_pts = find_nearest_sample(df_pts, df_cat)
        print(f"DataFrame of the desired points and the nearby sampled "
            f"value\n{ df_pts }")

        # Plot the restraint points
        plot_restraint_positions(df_pts, f"{ wrkdir }/restraint_points.png")

    if initial_conforms:
        with open(f"{ wrkdir }/plumed.dat", "r") as f:
            # Get the plumed template 
            plumed_lines = f.readlines() 
        extract_window_conforms(df_pts, plumed_lines)

    else: 
        # Extract the nearby structure for each point in df_pts
        print(df_pts)
        for i, row in df_pts.iterrows():
            print(row)
            out_struct = f"{ wrkdir }/conform_{ i }.pdb"
            extract_conform(row, out_struct)

def extract_window_conforms(df_pts, plumed_lines):
    """Extracts conforms, prepares plumed files, prepares batch script.

    Parameters
    ----------
    df_pts : pd.DataFrame
        Contains the positions of the desired points in the reaction 
        coordinate space. 
    plumed_lines : (str) list
        Lines from the plumed template file. 

    Returns
    -------
    None. 

    """
    # Keeps a record of new windows ready for sampling
    batch_arr = []

    # OpenPoint,ClosedPoint 
    for i, row in df_pts.iterrows():

        w = i + 1

        # Set up some paths
        destination = (f"{ wrkdir }/windows_setup/window{ w }")
        os.makedirs(destination, exist_ok=True)
        plumed_file = f"{ destination }/plumed_{ w }.dat"
        initial_out = f"{ destination }/initial_conform.pdb"

        # Extract the nearby structure for each point in df_pts
        extract_conform(row, initial_out)

        if row["NearestConformName"] is not None:

            batch_ids = [4 * (w - 1) + r for r in range(1,5)]
            batch_arr.extend(batch_ids)

        plumed_window(row, w, plumed_lines, plumed_file)

    # Get the array sbatch script template file
    with open(f"{ wrkdir }/us_array_template.sh", "r") as f:
        sbatch_lines = f.readlines()

    # Modify the template to include the relevant windows array
    # Recall, 4 runs per window, each with a separate batch array
    batch_arr = list(map(lambda x : str(int(x)), batch_arr))
    batch_str = ",".join(batch_arr)
    new_batch_lines = []
    for line in sbatch_lines:
        if "--array=test" in line:
            line = line.replace("--array=test",
                                f"--array={ batch_str }")
        new_batch_lines.append(line)

    # Write out the new batch script
    with open(f"{ wrkdir }/windows_setup2/us_array_initial.sh", "w") as f:
        f.writelines(new_batch_lines)

    return None

def cv_min_dist(grid_pt, data):
    """How far is the grid point to the nearest sampled point?

    Finds the minimum distance and and the index of the sampled point 
    closest to the grid point.
    
    Parameters
    ----------
    grid_pt : np.ndarray
        The 2D grid point.
    data : np.ndarray
        The 2D array of sampled points.

    Returns
    -------
    min_d : float
        The distance of the closest point, which helps determine whether
        the sample is sufficiently close.
    min_ind : int
        The index of the relevant sample. 

    """
    d = np.sqrt(np.sum(np.square(np.transpose(data) 
                                 - np.array(grid_pt)), axis=1))
    min_d = np.min(d)
    min_ind = np.argmin(d)

    return min_d, min_ind 

def plumed_align_ref(wrkdir):
    """Writes to pdb the subset of atoms for alignment and biasing.

    The original atom indicies must be preservered since this is what 
    plumed is using to identify atoms from the reference file.

    Parameters
    ----------
    wrkdir : str
        Path to the working directory.

    Returns
    -------
    None.

    """
    ref_file = f"{ wrkdir }/ref.pdb"

    # Define the atom selection based on alignment residues and the 
    # Beta vector c-alpha atoms
    vec_select = (" or (resid 206 and name CA) or "
                  "(resid 215 and name CA))")
    core_res, core = traj_funcs.get_core_res() 

    # Grab the atom selection
    u = mda.Universe(f"{ config.struct_head }/ref_all_atoms.pdb")
    core_and_vec = u.select_atoms(core[:-1] + vec_select)

    # Write the relevant lines to a pdb file
    with open(ref_file, 'w') as pdb_file:
        for atom in core_and_vec:
            pdb_line = (f"ATOM  {(atom.ix + 1):5} {atom.name:<4}"
                        f"{atom.resname:<3} X {atom.resid:>4}    "
                        f"{atom.position[0]:8.3f}{atom.position[1]:8.3f}"
                        f"{atom.position[2]:8.3f}{atom.occupancy:6.2f}"
                        f"{atom.bfactor:6.2f}          \n")
            pdb_file.write(pdb_line)

    print(f"Wrote reference alignment structure for plumed to "
          f"{ ref_file }.")

    return None

def sim_data_paths(data_paths, topols, xtc, alphafold):
    """Collects paths related to simulation data. 

    Parameters
    ----------
    data_paths : (str) list
        A list of paths to simulation data.
    topols : (str) list
        A list of names of topol files.
    xtc : str
        Name for all the xtc files.
    alphafold : bool
        If true, will include alphafold simulation data when searching 
        for suitable conformations.

    Returns
    -------
    traj_paths : dict
        Dictionary of the paths for trajectory files, assuming 'nobackup'
        as first level dirname.
    top_paths : dict
        Dictionary of the paths for topology files, assuming 'nobackup'
        as first level dirname.
    """
    traj_paths = {}
    top_paths = {}
    af_xtc = "fitted_traj_100.xtc"
    af_top = "topol_protein.top"
    if alphafold:
        af_path = f"{ data_head }/unbiased_sims/af_replicas"
        for i in range(1,10):
            trajs[f"af {i}"] = f"{ af_path }/af{i}/nobackup/{ af_xtc }"
            tops[f"af {i}"] = f"{ af_path }/af{i}/nobackup/{ af_top }"
    for p, top in zip(data_paths, topols):
        print("\n", p, "\n")
        n = p.split("/")[-3]
        traj_paths[n] = f"{ p }/{ xtc }"
        top_paths[n] = f"{ p }/{ top }"
    
    return traj_paths, top_paths

def find_nearest_sample(df_pts, df_cat):
    """Finds the closest sampled structure for each point in df_pts.

    Parameters
    ----------
    df_pts : pd.DataFrame
        Contains the positions of the desired points in the reaction 
        coordinate space. 
    df_cat : pd.DataFrame
        Contains trajectory data for the reaction coordinates, produced
        using the VectorCoordCombo.py script. 

    Returns
    -------
    df_pts : pd.DataFrame
        Contains the positions of the desired points in the reaction 
        coordinate space and the trajectory name and timestep for points
        with a nearby sample. 

    """
    # Make an array of the desired points
    restraint_grid = np.array((df_pts.OpenPoint, df_pts.ClosedPoint))

    # Add columns to the df to record the trajectory and timestep of the
    # matching conformation
    traj_names = [None] * len(df_pts.OpenPoint)
    traj_frames = [None] * len(df_pts.OpenPoint)

    # Determine which sampled conform is closest
    ds = []
    for i, g in enumerate(restraint_grid.T):
        d, d_ind = cv_min_dist(g, (df_cat["dot-open"], df_cat["dot-closed"]))
        ds.append(d)
        
        # Use a minimum distance so no conform is extracted for poor 
        # sampling in the vicinity
        if d < 0.5:
            traj_names[i] = df_cat.loc[d_ind, "traj"]
            traj_frames[i] = df_cat.loc[d_ind, "ts"]

    # Collect data in the DataFrame
    df_pts["NearestDist"] = ds
    df_pts["NearestConformName"] = traj_names
    df_pts["NearestConformFrame"] = traj_frames

    return df_pts

def find_nearest_sb(df_pts, df_cat):
    """Finds the closest sampled structure for each point in df_pts.

    Parameters
    ----------
    df_pts : pd.DataFrame
        Contains the positions of the desired points in the reaction 
        coordinate space. 
    df_cat : pd.DataFrame
        Contains trajectory data for the reaction coordinates, produced
        using the VectorCoordCombo.py script. 

    Returns
    -------
    df_pts : pd.DataFrame
        Contains the positions of the desired points in the reaction 
        coordinate space and the trajectory name and timestep for points
        with a nearby sample. 

    """
    # Make an array of the desired points
    restraint_pts = np.array(df_pts.SaltBridge)

    # Add columns to the df to record the trajectory and timestep of the
    # matching conformation
    traj_names = [None] * len(df_pts.OpenPoints)
    traj_frames = [None] * len(df_pts.OpenPoints)

    # Determine which sampled conform is closest
    ds = []
    for i, g in enumerate(restraint_pts):
        d = np.abs(g-df_cat["salt-bridge"])
        min_d = np.min(d)
        d_ind = np.argmin(d)
        ds.append(min_d)
        
        # Use a minimum distance so no conform is extracted for poor 
        # sampling in the vicinity
        if min_d < 0.5:
            traj_names[i] = df_cat.loc[d_ind, "traj"]
            traj_frames[i] = df_cat.loc[d_ind, "ts"]

    # Collect data in the DataFrame
    df_pts["NearestDist"] = ds
    df_pts["NearestConformName"] = traj_names
    df_pts["NearestConformFrame"] = traj_frames

    return df_pts

def plot_restraint_positions(df_pts, fig_path):
    """Plots and labels the restraint points.

    Parameters
    ----------
    df_pts : pd.DataFrame
        Contains the positions of the desired points in the reaction 
        coordinate space. 
    fig_path : str
        The path and name for the figure. 

    Returns
    -------
    None.

    """
    fig, ax = plt.subplots()

    # Scatter plot for all data, and data subset
    ax.scatter(df_pts["OpenPoint"], df_pts["ClosedPoint"], s=100, 
        label="All restraint points")
    masked_df = df_pts.mask(df_pts["NearestConformName"].notna())
    ax.scatter(masked_df["OpenPoint"], masked_df["ClosedPoint"], s=50, 
        label="Conforms available\nfrom sampling")

    # Plot settings
    plt.legend(fontsize=20)
    ax.set_xlabel(r"$\xi_1$ (nm$^2$)", labelpad=5, fontsize=24)
    ax.set_ylabel(r"$\xi_2$ (nm$^2$)", labelpad=5, fontsize=24)

    # Save figure and close
    utils.save_figure(fig, fig_path)
    plt.close()

def plumed_window(row, w, plumed_lines, plumed_file):
    """Writes a plumed.dat file for the window using a template.

    Parameters
    ----------
    row : pd.Series
        The row data for a particular restraint point.
    w : int
        The numerical label for the window.
    plumed_lines : (str) list
        Lines from the plumed template file. 
    plumed_file : str
        The path and file name for the plumed file for the window. 

    Returns
    -------
    None. 

    """
    plumed_restraints = []

    # Substitute variable names into template
    for line in plumed_lines:
        if "RESTRAINT_OPEN" in line:
            line = line.replace("RESTRAINT_OPEN", 
                str(row["OpenPoint"]))
        if "RESTRAINT_CLOSED" in line:
            line = line.replace("RESTRAINT_CLOSED", 
                str(row["ClosedPoint"]))
        if "COLVAR_WINDOW" in line:
            line = line.replace("COLVAR_WINDOW", "COLVAR_" + str(w))
        plumed_restraints.append(line)

    # Write the plumed file for the specific window
    with open(plumed_file, "w") as f:
        f.writelines(plumed_restraints)

    return None

def extract_conform(row, out_struct):
    """Extracts a conformation for the restraint point with gromacs. 

    Parameters
    ----------
    row : pd.Series
        The row data for a particular restraint point.
    out_struct : str
        Path to the extracted structure. 

    Returns
    -------
    None. 

    """
    if row["NearestConformName"] is not None:
        traj = traj_paths[row["NearestConformName"]]
        top = top_paths[row["NearestConformName"]]
        time_frame = row["NearestConformFrame"]

        if state == "holo":
            ind_code = "24"
        else:
            ind_code = "1"

        gmx = ["echo", ind_code, "|", "gmx22", "trjconv", "-f", traj, 
            "-s", top, "-o", out_struct, "-b", str(time_frame - 1000), 
            "-dump", str(time_frame), "-nobackup"]

        if state == "holo":
            gmx.extend(["-n", index])
        
        # Uses a subprocess to run gromacs
        output = subprocess.Popen(" ".join(gmx), 
                                    stdout=subprocess.PIPE, 
                                    shell=True)

        print(out_struct)
        time.sleep(1)

    return None

if __name__ == '__main__':
    main(sys.argv)
```
