## Supplementary material for "*Clostridioides difficile* Toxins Unhinged: Allosterically Switchable Network Orients *β*-flap": Scripts, Structures etc.: FES2D.py

```
import numpy as np
import matplotlib.pyplot as plt
import sys
import os
import plumed
import wham
import MDAnalysis as mda
from MDAnalysis.analysis import align
import subprocess    
from mpl_toolkits.mplot3d import Axes3D
from matplotlib.tri import Triangulation
from scipy.signal import find_peaks
import argparse
import pandas as pd
sys.path.insert(0, "/home/lf1071fu/project_b3")
sys.path.insert(0, "/home/lf1071fu/project_b3/ProjectB3")
import config.settings as config
from tools import utils, traj_funcs
import matplotlib.colors as mcolors

def main(argv):

    try:
        parser = argparse.ArgumentParser()

        parser.add_argument("-p", "--path",
                            action = "store",
                            dest = "path",
                            default = ("umbrella/holo_state/nobackup"),
                            help = ("Set relative path to the data "
                                "directory."))        
        parser.add_argument("-a", "--angle",
                            action = "store_true",
                            dest = "angle_coord",
                            default = False,
                            help = ("Chose whether the reaction "
                                "coordinate should be converted into an "
                                "angle."))
        parser.add_argument("-s", "--state",
                            action = "store",
                            dest = "state",
                            default = "holo",
                            help = ("Chose the state for the FES, i.e. "
                                "apo, holo, mutant"))
        parser.add_argument("-f", "--figpath",
                            action = "store",
                            dest = "fig_path",
                            default = ("umbrella/holo_state"),
                            help = ("Set a relative path destination for"
                                " the figure."))  
        parser.add_argument("-b", "--blocks",
                            action = "store",
                            dest = "nblocks",
                            default = 100,
                            help = ("Number of blocks to use in "
                                "bootstrap analysis; 1 ns each by "
                                "default."))            
        parser.add_argument("--nbins",
                            action = "store",
                            dest = "nbins",
                            default = 150,
                            help = ("Number of bins for histograms."))   
        parser.add_argument("-r", "--recalc",
                            action = "store_true",
                            dest = "recalc",
                            default = False,
                            help = ("Chose whether the reweighting "
                                "should be recomputed."))
        parser.add_argument("-n", "--nobootstrap",
                            action = "store_false",
                            dest = "no_bootstrap",
                            default = True,
                            help = ("Compute the free energy surface "
                                "without error analysis."))
        parser.add_argument("-w", "--num_windows",
                            action = "store",
                            dest = "num_windows",
                            default = 170,
                            help = ("Number of windows used in the "))
        args = parser.parse_args()

    except argparse.ArgumentError:
        print("Command line arguments are ill-defined, please check the "
            "arguments.")
        raise

    global nb, cv, nw, kBT, angle_coord, recalc, home, colvar_columns
    global no_bs, nbins, state, bs

    # Set key path variables
    home = f"{ config.data_head }/{ args.path }"
    fig_path = f"{ config.figure_head }/{ args.fig_path }"
    struct_path = config.struct_head

    # Set other variables
    angle_coord = args.angle_coord
    recalc = args.recalc
    no_bs = args.no_bootstrap # don't do error analysis
    nb = int(args.nblocks) # number of blocks for bootstrapping
    nw = int(args.num_windows) # number of windows 
    bs = 200 # number of bootstraps
    kBT = 310 * 8.314462618 * 0.001 # use kJ/mol here
    nbins = int(args.nbins)
    state = args.state

    # Helpful objects for structure alignment
    core_res, core = traj_funcs.get_core_res()
    ref_state = mda.Universe(f"{ struct_path }/ref_all_atoms.pdb", 
                             length_unit="nm")

    vec_open, vec_closed = get_ref_vecs(struct_path, core, ref_state)
    if state == "holo":
        colvar_columns = ["time", "opendot", "closeddot", "theta1", 
            "theta2", "d4", "d5", "d6", "bias", "force"]
    elif state == "apo":
        colvar_columns = ["time", "opendot", "closeddot", "theta1", 
            "theta2", "bias", "force"]
    else: 
        colvar_columns = ["time", "opendot", "closeddot", "theta1", 
            "theta2", "bias", "force"]
            
    bs_path = f"{ home }/bootstrap_files"
    if not os.path.exists(bs_path):
        os.makedirs(bs_path)
    np_ave_files = [f"{ home }/hist_ave.npy", 
                    f"{ home }/hist_sq_ave.npy", 
                    f"{ home }/bin_counter.npy"]

    # Make the FES without error estimates 
    if no_bs:

        fes = get_no_bs_fes(home, bs_path)
        print(fes)

        # Make the 2D FES plot
        plot_2dfes(fes, vec_open, vec_closed, fig_path)
        
        # Also make the 1D FES plots
        for rxn_coord in ["open", "closed"]:
            plot_1dfes(bs_path, rxn_coord, fig_path)

    elif not all(list(map(lambda x : os.path.exists(x), 
                                np_ave_files))) or recalc: 

        pfes, fsum, fsq_sum, count = get_bs_fes(home, bs_path, 
                                                np_ave_files)

        # Make the 2D FES plot
        plot_2dfes(pfes, vec_open, vec_closed, fig_path, fsum=fsum, 
                   fsq_sum=fsq_sum, count=count)

        # Error quantities
        var = (1 / (count - 1)) * ( fsq_sum / count - fsum * fsum ) 
        error = np.sqrt( var )
        ferr = error / (fsum / count)

        # Make the 2D error estimates plot
        plot_2derror(ferr, pfes, vec_open, vec_closed, fig_path)

    # Load calculated values from pandas tables and numpy arrays
    else:

        # Get bin positions for the CVs
        pfes = plumed.read_as_pandas(f"{ bs_path }/"
                    "fes_catr_0.dat").replace([np.inf, -np.inf], np.nan)

        # Load the sum of the free energy esimates from each bin
        fsum = np.load(np_ave_files[0], allow_pickle=True)
        fsq_sum = np.load(np_ave_files[1], allow_pickle=True)
        count = np.load(np_ave_files[2], allow_pickle=True)

        # Make the 2D FES plot
        plot_2dfes(pfes, vec_open, vec_closed, fig_path, fsum=fsum, 
                   fsq_sum=fsq_sum, count=count)

        # Error quantities
        count[(count == 0) | (count == 1)] = np.nan
        ave = fsum / count
        ave2 = fsq_sum / count
        var = (count / (count - 1)) * ( ave2 - ave * ave ) 
        error = np.sqrt( var )
        ferr = error / (ave)

        # Make the 2D error estimates plot
        plot_2derror(error, pfes, vec_open, vec_closed, fig_path)

    # Calculate free energy surface from histogram averages and 
    # free energy errors from the histograms, see expressions in ex-5 
    # https://www.plumed.org/doc-v2.8/user-doc/html/masterclass-21-2.html
    #fes = ave / count
    # # fes = convert_fes(hist.odot, hist.cdot, ave)

    # fes = - kBT * np.log(ave)
    #error = np.sqrt( var )
    #ferr = error / ave

    # Make the error estimate plot for the 2D FES

    return None

def plot_2dfes(fes, vec_open, vec_closed, fig_path, fsum=None, 
        fsq_sum=None, count=None):
    """Plots the 2D FES as a colorbar + contour map.

    fes : pd.DataFrame
        The table containing discretized free energy surface data. 
    vec_open : float
        The reference beta-vector for the open conformation.
    vec_closed : float
        The reference beta-vector for the closed conformation.
    fig_path : str
        Path for storing the figure. 
    ave : np.ndarray

    ave_sq : np.ndarray

    """
    fig, ax = plt.subplots(constrained_layout=True, figsize=(2.5,2))

    # Get the relevant discretized arrays from table columns
    open_bins, closed_bins = fes.odot, fes.cdot 
    if no_bs:
        fes = fes.ffr
    else:
        fes = np.divide(fsum, count, where=count != 0)

    mask = ((-1e2 < fes) & (fes < 1e2) & (count != 0))
    mask[0] = False
    # mask = (fes < 1e6)

    x, y = open_bins[mask], closed_bins[mask]
    z = fes[mask] - min(fes[mask])
    #x, y = open_bins, closed_bins
    #z = fes
    # z, err = fes[mask], ferr[mask]

    # Save plotted data as np array
    combined_array = np.stack((x, y, z), axis=0)
    print(combined_array)
    print(combined_array.shape)
    np.save(f"{ home }/fes_data_{ state }.npy", combined_array)

    closed_mask = z[y>3]
    open_mask = z[y<=3]
    print(f"MIN-closed: { np.min(closed_mask)}\nMIN-open: { np.min(open_mask)}")
    print(f"diff : {np.min(open_mask) - np.min(closed_mask)}")

    d = ax.scatter(x, y, c=z, cmap=plt.cm.viridis, s=1,
            norm=mcolors.Normalize(vmin=0, vmax=100))

    # Get Free energy values of the reference states
    o_ref_x, o_ref_y = (np.dot(vec_open, vec_open), np.dot(vec_open, vec_closed))
    distances_open = np.sqrt((x - o_ref_x)**2 + (y - o_ref_y)**2)
    nearest_open_index = np.argmin(distances_open)
    print(f"\nF(open_ref) = {z[nearest_open_index]}")

    c_ref_x, c_ref_y = (np.dot(vec_open, vec_closed), np.dot(vec_closed, vec_closed))
    distances_closed = np.sqrt((x - c_ref_x)**2 + (y - c_ref_y)**2)
    nearest_closed_index = np.argmin(distances_closed)
    print(f"\nF(closed_ref) = {z[nearest_closed_index]}")

    # if angle_coord:
    #     fes_r=plumed.read_as_pandas("fes_theta_catr.dat").replace([np.inf, -np.inf], np.nan).dropna()
    #     x = [ x for x, z in zip(fes_r.theta1, fes_r.ffthetar) if -80 < z < 80 ]
    #     y = [ y for y, z in zip(fes_r.theta2, fes_r.ffthetar) if -80 < z < 80 ]
    #     zmin_g = np.min(np.array([f for f in fes_r.ffthetar if f > -80]))
    #     zmin_l = np.min(np.array([f for x, f in zip(fes_r.theta1, fes_r.ffthetar) if ((f > -80) and (x < 0.75))]))
    #     z = [ z - zmin_g for z in fes_r.ffthetar if -80 < z < 80 ]

    # else:
    # fes_r=plumed.read_as_pandas("fes_catr.dat").replace([np.inf, -np.inf], np.nan).dropna()
    # x = [ x for x, z in zip(fes_r.opendot, fes_r.ffr) if -80 < z < 80 ]
    # y = [ y for y, z in zip(fes_r.closeddot, fes_r.ffr) if -80 < z < 80 ]
    # zmin_g = np.min(np.array([f for f in fes_r.ffr if f > -80]))
    # zmin_l = np.min(np.array([f for x, f in zip(fes_r.opendot, fes_r.ffr) if ((f > -80) and (x > 4))]))
    # z = [ z - zmin_g for z in fes_r.ffr if -80 < z < 80 ]
    # print(f"Global min: { zmin_g } kJ/mol\tLocal min: { zmin_l } kJ/mol")

    tri = Triangulation(x, y)

    # Create contour lines on the XY plane using tricontour
    if "K57G" in state:
        contours = ax.tricontour(tri, z, cmap="inferno", levels=[0,15,30,45,60,75,101,102])
    else:
        contours = ax.tricontour(tri, z, cmap="inferno", levels=6)

    # Colormap settings
    cbar = plt.colorbar(d, ticks=np.arange(0, 101, 20))
    # if angle_coord:
    #     cbar.set_label(r'$F(\theta_{open}, \theta_{closed})$ (kJ / mol)', fontsize=24, labelpad=10)
    # else: 
    cbar.set_label(r'$F(\xi_1, \xi_2)$ (kJ / mol)', fontsize=8, labelpad=3)
    cbar.ax.tick_params(labelsize=6)
    cbar.outline.set_linewidth(1)
    ax.clabel(contours, inline=1, fontsize=6)

    # Add both reference positions
    if angle_coord:
        ax.scatter(calc_theta(vec_open, vec_open), calc_theta(vec_open, vec_closed), 
                label="Raised ref.", marker="X", alpha=1, edgecolors="#404040", 
                s=550, lw=3, color="#EAAFCC")
        ax.scatter(calc_theta(vec_open, vec_closed), 0, label="Lowered ref.", 
                marker="X", alpha=1, edgecolors="#404040", s=550, lw=3, color="#A1DEA1")
    else:
        ax.scatter(np.dot(vec_open, vec_closed), np.dot(vec_closed, vec_closed), 
                label="Lowered ref.", marker="X", alpha=1, edgecolors="#ededed", 
                s=50, lw=1, color=config.closed_color)
        ax.scatter(np.dot(vec_open, vec_open), np.dot(vec_open, vec_closed), 
                label="Raised ref.", marker="X", alpha=1, edgecolors="#ededed", 
                s=50, lw=1, color=config.open_color)
    if "holo" in state:
        ax.scatter(np.dot(vec_open, vec_open), np.dot(vec_open, vec_closed), 
                label="Raised ref.", marker="X", alpha=1, edgecolors="#ededed", 
                s=50, lw=1, color=config.open_color)    

    # Plot parameters and axis labels
    ax.tick_params(axis='both', which='major', pad=3)
    plt.xticks([0, 2, 4, 6])
    plt.yticks([0, 2, 4, 6])
    if angle_coord:
        ax.set_xlabel(r"$\theta_{open}$ (rad)", labelpad=3)
        ax.set_ylabel(r"$\theta_{closed}$ (rad)", labelpad=3)
    else: 
        ax.set_xlabel(r"$\xi_1$ (nm$^2$)", labelpad=3)
        ax.set_ylabel(r"$\xi_2$ (nm$^2$)", labelpad=3)
    _, xmax = ax.get_xlim()
    _, ymax = ax.get_ylim()
    ax.set_xlim(0,xmax)
    ax.set_ylim(0,ymax)
    ax.set_aspect('equal', adjustable='box')
    plt.legend(fontsize=6)

    if angle_coord:
        utils.save_figure(fig, f"{ fig_path }/2dfes_angles_{ state }.png")
    elif not no_bs:
        utils.save_figure(fig, f"{ fig_path }/2dfes_bs_{ state }.png")
    else:
        utils.save_figure(fig, f"{ fig_path }/2dfes_{ state }.png")

    plt.close()

def plot_2derror(ferr, fes, vec_open, vec_closed, fig_path):
    """Makes a plot of the FES error.

    Parameters
    ----------
    fes : pd.DataFrame
        The table containing discretized free energy surface data. 
    vec_open : float
        The reference beta-vector for the open conformation.
    vec_closed : float
        The reference beta-vector for the closed conformation.
    fig_path : str
        Path for storing the figure. 
    
    """
    fig, ax = plt.subplots(constrained_layout=True, figsize=(2.5,2))

    # Get the relevant discretized arrays from table columns
    open_bins, closed_bins = fes.odot, fes.cdot 

    mask = ~np.isnan(ferr)
    mask[0] = 0
    x, y = open_bins[mask], closed_bins[mask]
    z = ferr[mask]

    d = ax.scatter(x, y, c=z, cmap=plt.cm.viridis, s=1, 
            norm=mcolors.Normalize(vmin=0, vmax=1))

    # Colormap settings
    cbar = plt.colorbar(d, ticks=np.arange(0, 1.2, 1/5))
    cbar.set_label(r'$F(\xi_1, \xi_2)$ (kJ / mol)', fontsize=8, labelpad=3)
    cbar.ax.tick_params(labelsize=6)
    cbar.outline.set_linewidth(1)

    ax.scatter(np.dot(vec_open, vec_open), np.dot(vec_open, vec_closed), 
                label="Raised ref.", marker="X", alpha=1, edgecolors="#ededed", 
                s=50, lw=1, color=config.open_color)
    ax.scatter(np.dot(vec_open, vec_closed), np.dot(vec_closed, vec_closed), 
                label="Lowered ref.", marker="X", alpha=1, edgecolors="#ededed", 
                s=50, lw=1, color=config.closed_color)

    # Plot parameters and axis labels
    ax.tick_params(axis='both', which='major', pad=3)
    plt.xticks([0, 2, 4, 6])
    plt.yticks([0, 2, 4, 6]) 
    ax.set_xlabel(r"$\xi_1$ (nm$^2$)", labelpad=3)
    ax.set_ylabel(r"$\xi_2$ (nm$^2$)", labelpad=3)
    ax.set_xlim(0,6)
    ax.set_ylim(0,6)
    ax.set_aspect('equal', adjustable='box')
    plt.legend(fontsize=6, loc='upper right')

    utils.save_figure(fig, f"{ fig_path }/2dfes_error_{ state }.png")

    plt.close()

    return None

def get_ref_vecs(struct_path, core, ref_state):
    """Calculates the beta-vectors for the reference structures.

    Parameters
    ----------
    struct_path : str
        Path to the structure directory, i.e. project_b3/structures
    core : str 
        The core residues used for alignment throughout this project.
    ref_state: MDAnalysis.Universe
        The reference structure object.

    Returns
    -------
    vec_open : np.ndarray
        The beta vector in Angstrom for the aligned open reference state.
    vec_closed : np.ndarray
        The beta vector in Angstrom for the aligned closed reference state.

    """
    # Load in relevant reference structures
    open_ref = mda.Universe(f"{ struct_path }/open_ref_state.pdb", length_unit="nm")
    closed_ref = mda.Universe(f"{ struct_path }/closed_ref_state.pdb", length_unit="nm")

    # Align the traj and ref states to one structure
    align.AlignTraj(open_ref, ref_state, select=core, in_memory=True).run()
    align.AlignTraj(closed_ref, ref_state, select=core, in_memory=True).run()

    # Determine open + closed reference beta flap vectors in units of Angstrom
    r1, r2 = 206, 215
    r1_open = open_ref.select_atoms(f"name CA and resnum { r1 }").positions[0]
    r2_open = open_ref.select_atoms(f"name CA and resnum { r2 }").positions[0]
    vec_open = r2_open/10 - r1_open/10
    r1_closed = closed_ref.select_atoms(f"name CA and resnum { r1 }").positions[0]
    r2_closed = closed_ref.select_atoms(f"name CA and resnum { r2 }").positions[0]
    vec_closed = r2_closed/10 - r1_closed/10

    return vec_open, vec_closed

def calc_theta(vec_ref, vec_sim):
    """Determine the angle between two 3D vectors.
    
    Solves the expression theta = cos^(-1)((A · B) / (|A| * |B|))

    Parameters
    ----------
    vec_ref : nd.array
        The referenece beta vector as a 3D array.

    vec_sim : nd.array
        An instance of the simulated beta as a 3D array. 

    Returns
    -------
    theta : float
        The angle formed by the two vectors, in radians.

    """
    # Calculate the dot product of A and B
    dot_product = np.dot(vec_ref, vec_sim)

    # Calculate the magnitudes of A and B
    magnitude_ref = np.linalg.norm(vec_ref)
    magnitude_sim = np.linalg.norm(vec_sim)

    # Calculate the angle (theta) between A and B using the formula
    theta = np.arccos(dot_product / (magnitude_ref * magnitude_sim))

    return theta

def plot_1dfes(bs_path, rxn_coord, fig_path):
    """Makes a plot against the 1D reaction coordinate.

    Parameters
    ----------
    rxn_coord : str
        Name of the reaction coordinate should match the column name.
    fig_path : str
        Path to figure directory for particular fes. 

    Returns
    -------
    None.

    """
    # Initialize the plot
    fig, ax = plt.subplots(constrained_layout=True, figsize=(2.5,2.5))

    # Load table containing discretized free energy surface data.
    fes = plumed.read_as_pandas(f"{ bs_path }/fes1d_{ rxn_coord }_fulldata.dat")
    fes = fes.replace([np.inf, -np.inf], np.nan).dropna()
    labels = {"apo" : "native,\napo state", "holo" : "native,\nholo state",
        "apo_K57G" : "K600G,\napo state", "holo_K57G" : "K600G,\nholo state"}

    # Labels for axes for the reaction coordinates
    rxn_coord_labs = {
        "open" : r"$\xi_1$",
        "closed" : r"$\xi_2$",
        "sb" : "K57--E200"
        }
        
    # Select reaction coordinate and lable axes accordingly
    if rxn_coord == "open":
        ax.plot(fes["odot"], fes["ffr1d_open"], label=labels[state], lw=2)
        ax.set_xlabel(f"{rxn_coord_labs[rxn_coord]} (nm$^2$)", labelpad=3,
            fontsize=8)
    elif rxn_coord == "closed":
        ax.plot(fes["cdot"], fes["ffr1d_closed"], label=labels[state], lw=2)
        ax.set_xlabel(f"{rxn_coord_labs[rxn_coord]} (nm$^2$)", labelpad=3,
            fontsize=8)
    elif rxn_coord == "sb":
        ax.plot(fes["sb"], fes["ffr1d_sb"], label=labels[state], lw=2,
            fontsize=8)
        ax.set_xlabel(f"{rxn_coord_labs[rxn_coord]} (nm)", labelpad=3)

    ax.set_ylabel(f"F({rxn_coord_labs[rxn_coord]}) (kJ / mol)", labelpad=3,
        fontsize=8)
    
    # Plot settings
    ax.tick_params(axis='both', which='major', pad=3)
    plt.legend(fontsize=6)
    _, xmax = ax.get_xlim()
    _, ymax = ax.get_ylim()
    ax.set_xlim(0,xmax)
    ax.set_ylim(0,ymax)

    # Save figure and close figure object
    utils.save_figure(fig, f"{ fig_path }/fes_{ rxn_coord }.png")
    # plt.show()
    plt.close()

    return None

def get_bias_data(home, recalc=False):
    """Get the 2D array of bias data from all windows.

    Parameters
    ----------
    recalc : bool
        Redetermine the collective variable from plumed files rather 
        than loading arrays from as numpy files.

    Returns
    -------
    bias : np.ndarray
        The bias applied to the full, concatenated trajectory for each 
        window, dim = (n-frames-concat, n-windows).
    frames : int
        The number of frames in the full concatenated trajectory.
    """
    bias_file = f"{ home }/bias_concat.npy"

    if not os.path.exists(bias_file) or recalc: 

        # Add all the COLVAR data to one large DataFrame 
        # NB requires a lot of memory!
        for i in range(nw):
            df = pd.read_csv(f"{ home }/COLVAR_" + str(i+1) + ".dat",
                       delim_whitespace=True, comment='#')
            if "bias" not in locals():
                frames = len(df.iloc[:,-2])
                fpw = frames // nw
                bias = np.zeros((frames, nw))
            # Reshape the bias array which is the second to last column
            print("Number of frames", frames)
            bias[:,i] = df.iloc[:,-2]

        utils.save_array(bias_file, bias)

    else:

        bias = np.load(bias_file, allow_pickle=True)
        fpw = len(bias[:,0]) // nw
    
    print(f"BIAS ARRAY {bias.shape}\n")

    return bias, fpw

def make_ith_histogram(df, bs_path, i, include_sb=False):
    """Use plumed HISTOGRAM function to obtain reweighted histograms for 
    the bootstrap.

    Parameters
    ----------
    df : pd.DataFrame
        Dataframe containing the collective variables and logweights.
    bs_path : str
        Path to directory for bootstraping data.
    i : int
        Integer for the bootstrap.
    include_sb : bool
        Whether to include a free energy profile related to the salt 
        bridge.

    Returns
    -------
    None.

    """
    print(df, df.head())
    plumed.write_pandas(df, f"{ bs_path }/colvar_weights_{ i }.dat")

    with open(f"{ bs_path }/colvar_histograms_{ i }.dat","w") as f:
        f.write(f"""
        # vim:ft=plumed
        odot: READ FILE={ bs_path }/colvar_weights_{ i }.dat VALUES=opendot IGNORE_TIME
        cdot: READ FILE={ bs_path }/colvar_weights_{ i }.dat VALUES=closeddot IGNORE_TIME
        theta1: READ FILE={ bs_path }/colvar_weights_{ i }.dat VALUES=theta1 IGNORE_TIME
        theta2: READ FILE={ bs_path }/colvar_weights_{ i }.dat VALUES=theta2 IGNORE_TIME

        lw: READ FILE={ bs_path }/colvar_weights_{ i }.dat VALUES=logweights IGNORE_TIME

        hhr: HISTOGRAM ARG=odot,cdot GRID_MIN=0,0 GRID_MAX=6,6 GRID_BIN={ nbins },{ nbins } BANDWIDTH=0.05,0.05 LOGWEIGHTS=lw
        DUMPGRID GRID=hhr FILE={ bs_path }/hist_catr_{ i }.dat
        ffr: CONVERT_TO_FES GRID=hhr 
        DUMPGRID GRID=ffr FILE={ bs_path }/fes_catr_{ i }.dat

        hhr1d_open: HISTOGRAM ARG=odot GRID_MIN=0 GRID_MAX=6 GRID_BIN=50 BANDWIDTH=0.05 LOGWEIGHTS=lw
        ffr1d_open: CONVERT_TO_FES GRID=hhr1d_open
        DUMPGRID GRID=ffr1d_open FILE={ bs_path }/fes1d_open_{ i }.dat

        hhr1d_closed: HISTOGRAM ARG=cdot GRID_MIN=0 GRID_MAX=6 GRID_BIN=50 BANDWIDTH=0.05 LOGWEIGHTS=lw
        ffr1d_closed: CONVERT_TO_FES GRID=hhr1d_closed
        DUMPGRID GRID=ffr1d_closed FILE={ bs_path }/fes1d_closed_{ i }.dat""")

        if include_sb:
            f.write(f"""
            sb: READ FILE={ bs_path }/colvar_weights_{ i }.dat VALUES=saltbridge IGNORE_TIME
            hhr1d_sb: HISTOGRAM ARG=sb GRID_MIN=0 GRID_MAX=2.5 GRID_BIN=50 BANDWIDTH=0.05 LOGWEIGHTS=lw
            ffr1d_sb: CONVERT_TO_FES GRID=hhr1d_sb
            DUMPGRID GRID=ffr1d_sb FILE={ bs_path }/fes1d_sb_{ i }.dat""")

    subprocess.run((f"plumed driver --noatoms --plumed { bs_path }/colvar" 
        f"_histograms_{ i }.dat --kt { kBT }"), shell=True)

    return None

def get_no_bs_fes(home, bs_path):
    """Estimate the free energy surface, without bootstrapping.

    Parameters
    ----------
    home : str
        Path to the primary data path, containing plumed-driver output.
    bs_path : str
        Path to bootstrapping calculations directory.

    Returns
    -------
    fes : pd.DataFrame
        Pandas table with collective variable data and free energy 
        estimates in kJ/mol.

    """
    fes_dat = f"{ bs_path }/fes_catr_fulldata.dat"

    if not os.path.exists(fes_dat) or recalc:

        # Get the biasing data from COLVAR files
        bias, fpw = get_bias_data(home, recalc=False)

        # Get collective variable data from any COLVAR file
        df = pd.read_csv(f"{ home }/COLVAR_1.dat", delim_whitespace=True,
                         comment='#', names=colvar_columns) 
        if state == "holo":
            df["saltbridge"] = np.minimum(df["d5"], df["d6"])

        # Reshape data for bootstrapping; this actually does nothing here
        # since the original column order is retained in "c", but the 
        # bootstrapping step is still performed for consistency
        bb = bias.reshape((nw, nb, fpw // nb, nw))
        c = np.arange(0, nb)
        colvar_weights = pd.DataFrame()
        for col in df.columns:
            colvar_weights[col] = bs_cols(df, col, fpw, nb , c) 

        # Get the log weights for reweighting with WHAM 
        w = wham.wham(bb[:,c,:,:].reshape((-1, nw)), T=kBT)
        print("wham shape : ", np.shape(w["logW"]), type(w["logW"]))
        print(w["logW"])
        
        # Write the logweights to the pandas table for reweighting
        colvar_weights["logweights"] = w["logW"]

        # Use plumed to make reweighted histograms
        if state == "holo":
            make_ith_histogram(colvar_weights, bs_path, "fulldata",
                include_sb=True)
        else:
            make_ith_histogram(colvar_weights, bs_path, "fulldata")

    # Load in FES table and remove NaN
    fes = plumed.read_as_pandas(fes_dat)
    fes = fes.replace([np.inf, -np.inf], np.nan).dropna()

    return fes

def get_bs_fes(home, bs_path, np_ave_files):
    """Estimate the free energy surface, with bootstrapping.

    Parameters
    ----------
    home : str
        Path to the primary data path, containing plumed-driver output.
    bs_path : str
        Path to bootstrapping calculations directory.
    np_ave_files : (str) list
        

    Returns
    -------
    pfes : pd.DataFrame
        Pandas table with collective variable data and free energy 
        estimates in kJ/mol.
    fsum : np.ndarray
        The sum of all bootstrap estimates to the FES for each bin.
    fsq_sum : np.ndarray
        The sum of the square of all bootstrap estimates to the FES for 
        each bin.
    count : np.ndarray
        An array for keeping track on the counts for estimates to each
        histogram bin.

    """
    # Get the biasing data from COLVAR files
    bias, fpw = get_bias_data(home, recalc=False)

    print(bias)
    print(f"SIZE OF BIAS { np.shape(bias) }")

    # Get collective variable data from any COLVAR file
    df = pd.read_csv(f"{ home }/COLVAR_1.dat", delim_whitespace=True,
                        comment='#', names=colvar_columns) 
    if state == "holo":
        df["saltbridge"] = np.minimum(df["d5"], df["d6"])

    # Reshape the bias into blocks that can be used for bootstrapping
    # bb = bias.reshape((nb, frames // nb, nw))
    bb = bias.reshape((nw, nb, fpw // nb, nw))

    # Initialize arrays for the histogram data
    fsum, fsq_sum = np.zeros((nbins + 1)**2), np.zeros((nbins + 1)**2)
    count = np.zeros((nbins + 1)**2)

    # Construct bootstraps and determine reweighted histograms
    for i in range(bs):

        # Choose random blocks for bootstap
        c = np.random.choice(nb, nb)

        pfes_file = f"{ bs_path }/fes_catr_{ i }.dat"
        hist_file = f"{ bs_path }/hist_catr_{ i }.dat"

        if not os.path.exists(pfes_file) or recalc:

            # Get the log weights for reweighting with WHAM 
            w = wham.wham(bb[:,c,:,:].reshape((-1, nw)), T=kBT)

            # Write the logweights to a pandas table for reweighting
            colvar_weights = pd.DataFrame()
            for col in df.columns:
                colvar_weights[col] = bs_cols(df, col, fpw, nb , c) 
            colvar_weights["logweights"] = w["logW"]

            # Use plumed to make reweighted histograms
            make_ith_histogram(colvar_weights, bs_path, i)

        # Load in free energy estimates determined via plumed
        pfes = plumed.read_as_pandas(pfes_file
                ).replace([np.inf, -np.inf], np.nan)

        # Use the ith histogram to estimate average probability densities
        mask = ~np.isnan(pfes.ffr) # some bins will be NAN
        frr = pfes.ffr.replace([np.inf, -np.inf], np.nan)
        fsum = np.nansum([fsum, frr], axis=0) 
        fsq_sum = np.nansum([fsq_sum, frr*frr], axis=0)
        # only bins with a bootstrap estimate should have the count 
        # increased
        count[mask] += 1 

    for file, arr in zip(np_ave_files, [fsum, fsq_sum, count]):
        utils.save_array(file, arr)

    return pfes, fsum, fsq_sum, count

def bs_cols(df, col, fpw, nb, c):
    """Reshape column arrays for bootstrapping.

    Parameters
    ----------
    df : pd.DataFrame
        Table of data from a COLVAR file.
    col : str
        Column name.
    fpw : int
        Number of frames in each window
    nb : int
        Number of bootstraps
    c : np.array
        Array for extracting columns for the bootstrap 

    Returns
    .......
    bs_flat : np.1darray
        Data is re-aranged into blocks according to "c" for 
        bootstrapping.

    """
    # Get the columns data and reshape into blocks
    arr = np.array(df[col].iloc[1:])
    arr = arr.reshape((nw, nb, fpw // nb))

    # Select blocks and flatten back into a 1d array with the 
    # original array length
    bs_flat = arr[:,c,:].flatten()

    return bs_flat

if __name__ == '__main__':
    main(sys.argv)
```
