## Supplementary material for "*Clostridioides difficile* Toxins Unhinged: Allosterically Switchable Network Orients *β*-flap": Scripts, Structures etc.: FesPaperPlot.py

```
import numpy as np
import matplotlib.pyplot as plt
import sys
import os
import MDAnalysis as mda
from MDAnalysis.analysis import align
from matplotlib.tri import Triangulation
from scipy.signal import find_peaks
import argparse
import pandas as pd
sys.path.insert(0, "/home/lf1071fu/project_b3/ProjectB3/biased")
sys.path.insert(0, "/home/lf1071fu/project_b3/ProjectB3")
import config.settings as config
from tools import utils, traj_funcs
from FES2D import get_ref_vecs
import plumed
import matplotlib.colors as mcolors
from collections import defaultdict, deque
import math
import heapq

def main(argv):

    try:
        parser = argparse.ArgumentParser()
        parser.add_argument("-f", "--figpath",
                    action = "store",
                    dest = "fig_path",
                    default = ("paper-figs"),
                    help = ("Set a relative path destination for"
                        " the figure."))
        parser.add_argument("-b", "--barrier",
                    action = "store_true",
                    dest = "barrier",
                    default = False,
                    help = ("Include the transition barrier position/"
                        "estimation."))
        parser.add_argument("-p", "--pathway",
                    action = "store_true",
                    dest = "pathway",
                    default = False,
                    help = ("Include the transition pathway"))                        
        args = parser.parse_args()

    except argparse.ArgumentError:
        print("Command line arguments are ill-defined, please check the "
            "arguments.")
        raise

    global fig_path, vec_open, vec_closed, pathway

    fig_path = f"{ config.figure_head }/{ args.fig_path }"
    struct_path = config.struct_head
    barrier = args.barrier
    pathway = args.pathway

    # Do structure alignment and obtain reference values
    core_res, core = traj_funcs.get_core_res()
    ref_state = mda.Universe(f"{ struct_path }/ref_all_atoms.pdb", 
                             length_unit="nm")
    vec_open, vec_closed = get_ref_vecs(struct_path, core, ref_state)

    # Make State objects for each FES
    states = {}
    for i, p in [("apo", "umbrella/apo_state"), 
              ("holo", "umbrella/holo_state"), 
              ("K57G-apo", "umbrella/apo_K57G"), 
              ("K57G-holo", "umbrella/holo_K57G")]:
        states[i] = State(i, p)

    plot_2dfes([states["apo"], states["holo"]], "fes_native", barrier=barrier)
    plot_2dfes([states["K57G-apo"], states["K57G-holo"]], "fes_mutant",
        barrier=barrier)

    if pathway:
        #plot_pathway([states["apo"], states["holo"]], "paths_native")
        #plot_pathway([states["K57G-apo"], states["K57G-holo"]], "paths_mutant")
        # plot_pathway([(states["apo"], states["K57G-apo"]), 
        #     (states["holo"], states["K57G-holo"])], "profile_combo", 
        #     compare=True)
        plot_comparison_pathway([(states["apo"], states["holo"]), 
            (states["K57G-apo"], states["K57G-holo"])], "profile_combo_2")

class State:
    def __init__(self, name, path):
        self.name = name
        top_path = f"{ config.data_head }/{ path }/nobackup/plumed_driver"
        self.path = f"{ top_path }/bootstrap_files"
        self.arrays = {}
        if "holo" in name:
            self.ligand = "holo"
        else:
            self.ligand = "apo"
        if "K57G" in name:
            self.mutant = True
        else:
            self.mutant = False

        # Get bin positions for the CVs
        self.pfes = plumed.read_as_pandas(f"{ self.path }/"
                    "fes_catr_0.dat").replace([np.inf, -np.inf], np.nan)

        # Load the sum of the free energy esimates from each bin
        self.fsum = np.load(f"{ top_path }/hist_ave.npy", allow_pickle=True)
        self.fsq_sum = np.load(f"{ top_path }/hist_sq_ave.npy", allow_pickle=True)
        self.count = np.load(f"{ top_path }/bin_counter.npy", allow_pickle=True)

        # Error quantities
        self.count[(self.count == 0) | (self.count == 1)] = np.nan
        ave = self.fsum / self.count
        ave2 = self.fsq_sum / self.count
        var = (self.count / (self.count - 1)) * ( ave2 - ave * ave ) 
        error = np.sqrt( var )
        self.ferr = error / (ave)

        # Transition barrier estimation
        saddle = {"apo" : (3.67, 3.79),
            "holo" : (3.85, 3.75),
            "K57G-apo" : (3.75,4.25),
            "K57G-holo" : (4.25,3.75)}
        self.barrier_x = saddle[name][0]
        self.barrier_y = saddle[name][1]

    def get_endpoints(self, fes):
        """
        """
        c_ref_x, c_ref_y = (np.dot(vec_open, vec_closed), 
            np.dot(vec_closed, vec_closed))
        distances_closed = np.sqrt((fes[0] - c_ref_x)**2 + (fes[1] - c_ref_y)**2)
        nci = np.argmin(distances_closed)
        self.start = (fes[0, nci], fes[1, nci], fes[2, nci])

        o_ref_x, o_ref_y = (np.dot(vec_open, vec_open), np.dot(vec_open, vec_closed))
        distances_open = np.sqrt((fes[0] - o_ref_x)**2 + (fes[1] - o_ref_y)**2)
        noi = np.argmin(distances_open)
        self.end = (fes[0, noi], fes[1, noi], fes[2, noi])

        return None

    def get_minima(self,fes):
        """
        """
        ind_closed = np.argmin(np.where(fes[1,:] > 3, fes[2,:], 100))
        self.closed_min = tuple(fes[:,ind_closed])

        ind_open = np.argmin(np.where(fes[1,:] < 3, fes[2,:], 100))
        self.open_min = tuple(fes[:,ind_open])
    
        return None

    def get_path(self, fes, reference=False):
        if reference:
            path_file = f"{ self.path }/pathway_ref_{ self.name }.npy"
            start, end = self.start, self.end
        else:
            path_file = f"{ self.path }/pathway_{ self.name }.npy"
            start, end = self.closed_min, self.open_min
        if os.path.exists(path_file):
            self.pathway = np.load(path_file, allow_pickle=True)
        else:
            pathway = a_star(fes, start, end)
            self.pathway = np.array(pathway)
            print(self.pathway.shape)
            utils.save_array(path_file, self.pathway)
        
        return None

def plot_2dfes(fes_states, fig_name, barrier=False):
    """Plots the 2D FES as a colorbar + contour map.

    fes : pd.DataFrame
        The table containing discretized free energy surface data. 
    fig_path : str
        Path for storing the figure. 
    ave : np.ndarray

    ave_sq : np.ndarray

    """
    # Figure with 2 columns
    fig, axes = plt.subplots(1, 2, constrained_layout=True, figsize=(5,2.33))

    for i, state in enumerate(fes_states):
        
        ax = axes[i]

        # Get the relevant discretized arrays from table columns
        open_bins, closed_bins = state.pfes.odot, state.pfes.cdot 
        energy = np.divide(state.fsum, state.count, where=state.count != 0)

        # Apply a mask to ignore very large values
        mask = ((-1e2 < energy) & (energy < 1e2) & (state.count != 0))
        mask[0] = False

        # Apply mask to data and print a summary of the local minima
        x, y = open_bins[mask], closed_bins[mask]
        z = energy[mask] - min(energy[mask])
        fes = np.stack((x, y, z), axis=0)
        state.fes = fes
        closed_mask = np.where(y > 3, z, 100)
        open_mask = np.where(y <= 3, z, 100)
        print(f"\nState { state.name }")
        print(f"MIN-closed: { np.min(closed_mask)}\nMIN-open: { np.min(open_mask)}")
        print(f"diff : {np.min(open_mask) - np.min(closed_mask)}")

        # Find a low energy path between the end points
        if pathway:
            # Get the endpoints translated onto the grid
            state.get_endpoints(fes)
            state.get_minima(fes)
            print(state.closed_min)
            print(state.open_min)

            # Find the shortest path between the endpoints and
            state.get_path(fes, reference=True)

        # import config.settings as cf
        # utils.save_array(f"{ cf.figure_head }/animations/strs_WT_path/cv_lowE_path.npy", 
        #         state.pathway)

        # Plot the primary surface data
        d = ax.scatter(x, y, c=z, cmap=plt.cm.viridis, s=1,
                norm=mcolors.Normalize(vmin=0, vmax=100))

        # Create contour lines on the XY plane using tricontour
        tri = Triangulation(x, y)
        if "K57G" in state.name:
            contours = ax.tricontour(tri, z, cmap="inferno", 
            levels=[0,15,30,45,60,75,101,102])
        else:
            contours = ax.tricontour(tri, z, cmap="inferno", levels=6)

        # Add both reference positions
        ax.scatter(np.dot(vec_open, vec_closed), np.dot(vec_closed, vec_closed), 
                    label="Lowered ref.", marker="X", alpha=1, edgecolors="#ededed", 
                    s=50, lw=1, color=config.closed_color)
        ax.scatter(np.dot(vec_open, vec_open), np.dot(vec_open, vec_closed), 
                    label="Raised ref.", marker="X", alpha=1, edgecolors="#ededed", 
                    s=50, lw=1, color=config.open_color)

        if "holo" in state.name:
            # Colormap settings
            cbar = fig.colorbar(d, ax=ax, ticks=np.arange(0, 101, 20))
            cbar.set_label(r'$F(\xi_1, \xi_2)$ (kJ / mol)', fontsize=8, labelpad=3)
            cbar.ax.tick_params(labelsize=6, length=3)
            cbar.outline.set_linewidth(1)

            # Add the shifted value
            shifted_index = np.argmin(np.where(y > 3, z, 100))
            ax.scatter(np.array(x)[shifted_index], 
                    np.array(closed_bins[mask])[shifted_index], 
                    label="Shifted lowered min.", marker="P", 
                    edgecolors="#ededed", 
                    s=50, lw=1, color=config.closed_color) 

        if barrier:
            # Plot the approximate transition barrier point
            ax.scatter(state.barrier_x, state.barrier_y, 
                    label="Approx. transition point", marker="*", alpha=1, 
                    edgecolors="#ededed", s=50, lw=1, color="#ebbd34")
            
            # Print out the barrier height
            distances_barrier = np.sqrt((x - state.barrier_x)**2 + (y - state.barrier_y)**2)
            nearest_barrier_index = np.argmin(distances_barrier)
            print(f"\nF(saddle_pt) = {z[nearest_barrier_index]}")

        if pathway:
            if "apo" in state.name:
                path_color = "#c4c2c2"
            else:
                path_color = "#E65100"
            ax.scatter(state.pathway[::5,0], state.pathway[::5,1], 
                color=path_color, label="Pathway", s=3)
        
        # Label the contours in both plots
        ax.clabel(contours, inline=1, fontsize=6)
                
        # Plot parameters and axis labels
        if "apo" in state.name:
            ax.set_ylabel(r"$\xi_2$ (nm$^2$)", labelpad=3)
        ax.tick_params(axis='both', which='major', pad=3)
        ax.set_yticks([0, 2, 4, 6])
        ax.set_xticks([0, 2, 4, 6])
        if "holo" in state.name:
            ax.set_yticklabels([])
        ax.set_xlabel(r"$\xi_1$ (nm$^2$)", labelpad=3)
        _, xmax = ax.get_xlim()
        _, ymax = ax.get_ylim()
        ax.set_xlim(0, xmax)
        ax.set_ylim(0, ymax)
        ax.set_aspect('equal', adjustable='box')
        ax.legend(fontsize=6)

    if barrier:
        print(f"{ fig_path }/{ fig_name }_barrier.png")
        utils.save_figure(fig, f"{ fig_path }/{ fig_name }_barrier.png")
    else:
        print(f"{ fig_path }/{ fig_name }.png")
        utils.save_figure(fig, f"{ fig_path }/{ fig_name }.png")

    plt.close()

    return None

def plot_pathway(fes_states, fig_name, compare=False):
    """Plots the pathway figure. 
    fes : pd.DataFrame
        The table containing discretized free energy surface data. 
    fig_path : str
        Path for storing the figure. 
    ave : np.ndarray

    ave_sq : np.ndarray

    """
    # Figure with 2 columns
    fig, axes = plt.subplots(1, 2, constrained_layout=True, figsize=(5,2.33))

    for i, state in enumerate(fes_states):

        if compare:
        
            state, mutant = state

        ax = axes[i]

        xvals = np.arange(0,len(state.pathway[:,0]))
        ax.plot(xvals, state.pathway[:, 2], lw=2, 
            color="#8f8f8f", label=state.name)

        if compare:

            ratio = len(state.pathway[:, 0])/len(mutant.pathway[:,0])
            xvals = np.arange(0,len(mutant.pathway[:,0]))
            ax.plot(xvals*ratio, mutant.pathway[:, 2], lw=2, label=mutant.name,
                color="#E65100")
            
        tick_labels = ["Lowered ref.","Lowered apo min.", "Barrier", "Raised min.",
            "Raised ref."]
        barrier = np.argmax((state.pathway[:,2]))
        print(f"Barrier height: { np.max(state.pathway[:,2]) }")

        if "mutant" in fig_name:
            closed_ind = np.argmin((state.pathway[:,1] - state.closed_min[1])**2) 
            open_ind = np.argmin((state.pathway[:,1] - state.open_min[1])**2) 
            tick_inds = [0, closed_ind, barrier, open_ind, -1]
        else: 
            closed_ind = np.argmin((state.pathway[:,0] - state.closed_min[0])**2) 
            open_ind = np.argmin((state.pathway[:,1] - state.open_min[1])**2) 
            tick_labels = tick_labels[:3] + ["Open min./\nRaised ref."]
            tick_inds = [0, closed_ind, barrier, -1]

        ax.set_xticks([xvals[j] for j in tick_inds])
        ax.set_xticklabels(tick_labels, rotation=45)

        if i == 0:
            ax.set_ylabel(r"$\Delta G$ (kJ / mol)", labelpad=3, fontsize=8)
        ax.set_xlabel("Reaction coordinate", labelpad=3, fontsize=8)

        ax.grid(False)
        ax.set_ylim([-9,50])

        if compare:
            ax.legend(fontsize=6)

    utils.save_figure(fig, f"{ fig_path }/{ fig_name }.png")
    plt.show()

    plt.close()

def plot_comparison_pathway(fes_states, fig_name):
    """Plots the pathway figure. 
    fes : pd.DataFrame
        The table containing discretized free energy surface data. 
    fig_path : str
        Path for storing the figure. 
    ave : np.ndarray

    ave_sq : np.ndarray

    """
    # Figure with 2 columns
    fig, axes = plt.subplots(1, 2, constrained_layout=True, figsize=(5,2.33))

    legend = {"apo" : "WT apo", "holo" : "WT holo",
        "K57G-apo" : "K600G apo", "K57G-holo" : "K600G holo"}

    for i, state in enumerate(fes_states):

        apo, holo = state

        ax = axes[i]

        # Plot the low free energy profiles
        xvals_apo = np.arange(0,len(apo.pathway[:,0]))
        ax.plot(xvals_apo, apo.pathway[:, 2] - np.min(apo.pathway[:90, 2]), 
            lw=2, color="#8f8f8f", label=legend[apo.name])
        ratio = len(apo.pathway[:, 0])/len(holo.pathway[:,0])
        xvals_holo = np.arange(0,len(holo.pathway[:,0])) * ratio
        ax.plot(xvals_holo, holo.pathway[:, 2] - np.min(holo.pathway[:90, 2]), 
            lw=2, label=legend[holo.name], color="#E65100")
        tick_labels = ["Lowered ref.", "Raised ref."]


        ax.set_xticks([xvals_apo[j] for j in [0,-1]])
        ax.set_xticklabels(tick_labels)

        if i == 0:
            ax.set_ylabel(r"$\Delta G$ (kJ / mol)", labelpad=3, fontsize=8)
        ax.set_xlabel(r"Conformation change progress", labelpad=3, fontsize=8)

        ax.grid(False)
        ax.set_ylim([-9,50])

        ax.legend(fontsize=6)

    utils.save_figure(fig, f"{ fig_path }/{ fig_name }.png")
    plt.show()

    plt.close()

def a_star(data, start, goal):
    # A* Algorithm
    def heuristic(a, b, data):
        # Combined heuristic: Euclidean distance and absolute z value difference
        euclidean_dist = np.sqrt((a[0] - b[0]) ** 2 + (a[1] - b[1]) ** 2)
        ind_a0 = np.where(data[0,:] == a[0])[0]
        ind_a1 = np.where(data[1,:] == a[1])[0]
        ind_b0 = np.where(data[0,:] == b[0])[0]
        ind_b1 = np.where(data[1,:] == b[1])[0]
        ind_a = list(set(ind_a0) & set(ind_a1))[0]
        ind_b = list(set(ind_b0) & set(ind_b1))[0]
        z_diff = abs(data[2,ind_a] - data[2,ind_b])
        return euclidean_dist + z_diff

    def euclidean_distance(node, point):
        return np.sqrt((node[0] - point[0]) ** 2 + (node[1] - point[1]) ** 2)

    def get_neighbors(node, data, distance=0.05):
        neighbors = []
        for i in range(data.shape[1]):
            if euclidean_distance(node, data[:,i]) <= distance:
                neighbors.append(tuple(data[:,i]))
        return neighbors

    open_set = []
    heapq.heappush(open_set, (0, start))
    came_from = {}
    g_score = {start: 0}
    f_score = {start: heuristic(start, goal, data)}

    while open_set:
        _, current = heapq.heappop(open_set)

        if current == goal:
            path = []
            while current in came_from:
                path.append(current)
                current = came_from[current]
            path.append(start)
            path.reverse()
            return path

        for neighbor in get_neighbors(current, data):
            tentative_g_score = g_score[current] + neighbor[2]

            if neighbor not in g_score or tentative_g_score < g_score[neighbor]:
                came_from[neighbor] = current
                g_score[neighbor] = tentative_g_score
                f_score[neighbor] = tentative_g_score + heuristic(neighbor, goal, data)
                if neighbor not in open_set:
                    heapq.heappush(open_set, (f_score[neighbor], neighbor))

    return None

if __name__ == '__main__':
    main(sys.argv)
```
