## Supplementary material for "*Clostridioides difficile* Toxins Unhinged: Allosterically Switchable Network Orients *β*-flap": Scripts, Structures etc.: NewWindowConfs.py

```
import numpy as np
import pandas as pd
import argparse
import os
import sys
import subprocess
import time

def main(argv):

    try:
        parser = argparse.ArgumentParser()

        parser.add_argument("-i", "--iteration",
                            action = "store",
                            dest = "iteration",
                            default = 1,
                            help = ("Set an integer value to obtain "
                                "the next iteration of initial conforms"
                                "for the umbrella windows. Counts from "
                                "1."))
        parser.add_argument("-r", "--recalc",
                            action = "store_true",
                            dest = "recalc",
                            default = False,
                            help = ("Overwrite the existing csv for the "
                                "selected iteration if it already "
                                "exists."))
        parser.add_argument("-e", "--extra_windows",
                            action = "store_true",
                            dest = "extra_windows",
                            default = False,
                            help = ("Includes additional windows in the "
                                "transition region."))
        args = parser.parse_args()

    except argparse.ArgumentError:
        print("Command line arguments are ill-defined, please check the "
            "arguments")
        raise

    # Assign command line arg
    iteration = int(args.iteration)
    recalc = args.recalc
    extra_windows = args.extra_windows

    # Get relevant restraint points
    df_pts = pd.read_csv(f"restraint_pts_iteration{ iteration }.csv")
    
    # Print the DataFrame of sampled reactions coords data
    pd.set_option("display.max_columns", None)
    pd.set_option("display.max_rows", None)

    # Check if the current iteration already exists
    new_pts_file = f"restraint_pts_iteration{ iteration + 1 }.csv"
    if os.path.exists(new_pts_file) and not recalc:
        print("Are you sure you want to overwrite the previous "
            "iteration? Add `-r` recalc flag to overrule, exiting now.")
        sys.exit(1)

    # Add additional windows?
    if extra_windows:
        df_pts = additional_windows(df_pts, typ=2)

    # Get sampled collective variables in one table
    df_cat = pd.DataFrame(columns=["time", "opendot", "closeddot"])
    df_pts["InitialConform"] = 0
    for w in range(1,171):
        for r in range(1,5):
            file = f"window{ w }/run{ r }/COLVAR_{ w }.dat"
            if os.path.exists(file):
                df_new = add_colvar_data(w, r, file)

                df_cat = pd.concat([df_cat, df_new])

            if os.path.exists(os.path.dirname(file)):
                df_pts.loc[df_pts["Window"] == w, "InitialConform"] = 1

    # Find all the window restraint points which have no COLVAR data.
    # These are the windows which still need an initial conform
    needs_conform = df_pts[(df_pts["InitialConform"] < 1)]
    print(f"NEEDS A CONFORM, { needs_conform.shape }\n", needs_conform, "\n")

    # Set up some variables
    restraint_grid = np.array((needs_conform.OpenPoints, 
                            needs_conform.ClosedPoints,
                            needs_conform.Window))
    df_pts = df_pts.assign(NearestWindow=0, NearestRun=0,
                       NearestFrame=0)

    # Iterate over all the restraints to find the closest conform
    # from the sampled windows
    for g in restraint_grid.T:
        d, d_ind = cv_min_dist(g[:2], (df_cat["opendot"], 
                                    df_cat["closeddot"]))
        
        # If a threshold is met, record the trajectory and time frame to
        # access an initial conformation for each missing window
        if d < 0.5:
            w, r, t = df_cat.iloc[d_ind][["window", "run", "time"]]
            df_pts.loc[df_pts["Window"] == g[2], "NearestWindow"] = w
            df_pts.loc[df_pts["Window"] == g[2], "NearestRun"] = r
            df_pts.loc[df_pts["Window"] == g[2], "NearestFrame"] = t

    # New DataFrame to use for extracting data
    print(f"DF PTS, { df_pts.shape }\n", df_pts, "\n")
    df_pts.to_csv(new_pts_file)

    # Initialize window dirs with new conformations
    df_filt = df_pts[(df_pts["InitialConform"] < 1)]
    print("DF FILTERED", df_filt)   

    # Get the plumed template file
    with open("plumed.dat", "r") as f:
        plumed_lines = f.readlines()  

    new_windows = []

    for i, row in df_pts.iterrows():
        
        w = int(row["Window"])

        # Get the nearby intial conformation from a biased simulation
        if row["NearestWindow"] > 0:

            # Substitute restraint values into the windows plumed file
            plumed_wlines = []
            for line in plumed_lines:
                if "RESTRAINT_OPEN" in line:
                    line = line.replace("RESTRAINT_OPEN", 
                                        str(row["OpenPoints"]))
                if "RESTRAINT_CLOSED" in line:
                    line = line.replace("RESTRAINT_CLOSED",
                                        str(row["ClosedPoints"]))
                if "COLVAR_WINDOW" in line:
                    line = line.replace("COLVAR_WINDOW", "COLVAR_" + str(w))
                plumed_wlines.append(line)

            # Prepare paths
            destination = f"window{ w }"
            os.makedirs(destination, exist_ok=True)
            plumed_wfile = f"{ destination }/plumed_{ w }.dat"
            initial_out = f"{ destination }/initial_conform.pdb"
            print("Initial out", initial_out)

            # Write out the plumed file for the window
            with open(plumed_wfile, "w") as f:
                f.writelines(plumed_wlines)
        
            # Get paths for extracting sampled conformation
            nw = str(int(row["NearestWindow"]))
            r = str(int(row["NearestRun"]))
            traj = f"window{ nw }/run{ r }/fitted_traj.xtc"
            top = f"window{ nw }/run{ r }/w{ nw }_r{ r }.tpr"
            time_frame = int(row["NearestFrame"])

            if not os.path.exists(traj):
                continue

            new_windows.append(w)

            # Define the gromacs command
            gmx = ["echo", "-e", "24", "|", "gmx", "trjconv", "-f", 
                traj, "-s", top, "-o", initial_out, "-b", 
                str(time_frame - 1000), "-dump", str(time_frame), "-n", 
                "index.ndx", "-nobackup"]
            print("\n", " ".join(gmx), "\n")
            
            # Use gromacs subprocess to extract the conformation at the 
            # desired time
            process = subprocess.Popen(" ".join(gmx), 
                                      stdin=subprocess.PIPE, 
                                      stdout=subprocess.PIPE,
                                      shell=True, text=True)

            # Pass input to the GROMACS command to use protein + ligand
            stdout, stderr = process.communicate("24\n")
            print("Output:", stdout)
            print("Error:", stderr)

            time.sleep(1)

    # Get the array sbatch script template file
    with open("us_array_template.sh", "r") as f:
        sbatch_lines = f.readlines()  

    # Add the array elements to the sbatch command, depending on which 
    # new windows will be sampled
    batch_arr = [str((val-1)*4 + i) for val in new_windows for i in range(1,5)]
    batch_str = ",".join(batch_arr)
    new_batch_lines = []
    for line in sbatch_lines:
        if "--array=test" in line:
            line = line.replace("--array=test", 
                                f"--array={ batch_str }")
        new_batch_lines.append(line)

    print(new_windows)
    print(batch_arr)

    # Write out the new batch script
    with open(f"us_array_{ iteration + 1 }.sh", "w") as f:
        f.writelines(new_batch_lines)

    return None

def additional_windows(df, typ=1):
    """Adds restraint values for more windows in the transition region. 

    Parameters 
    ----------
    df : pd.DataFrame
        DataFrame with the original restraint points.

    Returns
    -------
    df : pd.DataFrame
        DataFrame with the original and supplementary restraint points.

    """
    # determine the point positions
    if typ == 1:
        open_extras = np.linspace((1 + 5/6), (5 + 1/6), num=10, endpoint=True)
        closed_extras = np.linspace(1.5, (4 + 5/6), num=10, endpoint=True)
        windows = np.arange(151, 161)
    elif typ == 2:
        open_extras = np.linspace((2), (5 + 1/3), num=10, endpoint=False)
        closed_extras = np.linspace((1 + 2/3), (5), num=10, endpoint=False)
        windows = np.arange(161, 171)

    # Drop unnamed columns
    df = df.drop(columns=["Unnamed: 0", "Unnamed: 0.1"]) #"Unnamed: 0.1.1"])

    # Initialize new rows to a DataFrame
    df_new = pd.DataFrame({"OpenPoints" : open_extras, 
                "ClosedPoints" : closed_extras, 
                "Window": windows})

    # Combine to one table
    df = pd.concat([df, df_new], ignore_index=True).fillna(0)

    return df

def cv_min_dist(grid_pt, data):
    """Finds the nearest sampled data to the desired restraint point.

    Parameters
    ----------
    grid_pt : (float, float)
        The restraint point which needs an initial conformation. Give 
        point as (open-beta-vec, closed-beta-vec).
    data : np.ndarray
        A 2D array of the sampled open-beta-vec and closed-beta-vec. Note 
        that the data is transposed before combining with the restraint 
        point.

    Returns
    -------
    min_d : float
        The distance of the restraint point to the nearest sampled 
        conformation.
    min_ind : int
        The index of the point with minimum distance, which can be used 
        to find the nearest sampled conformation.
    """
    d = np.sqrt(np.sum(np.square(np.transpose(data) 
                                 - np.array(grid_pt)), axis=1))
    min_d = np.min(d)
    min_ind = np.argmin(d)
    return min_d, min_ind 

def add_colvar_data(window, run, file, stride=10):
    """Adds COLVAR data to a DataFrame from the sampled window.

    Parameters
    ----------
    window : int
        The window of the sampled data.
    run : int
        The run (1-4) of the sampled data. 
    file : str
        The COLVAR file which can be read into a DataFrame.

    Returns
    df : pd.DataFrame
        The DataFrame object with data for the given window + run.

    """
    df = pd.read_csv(file, comment="#", delim_whitespace=True, 
                       names=["time", "opendot", "closeddot",
                              "theta1", "theta2", "restraint.bias", 
                              "restraint.force2"])
    df["window"] = int(window)
    df["run"] = int(run)

    return df[::stride]

if __name__ == "__main__":
    main(sys.argv)
```
