## Supplementary material for "*Clostridioides difficile* Toxins Unhinged: Allosterically Switchable Network Orients *β*-flap": Scripts, Structures etc.: UpdateExploration.py

```
import os
import pandas as pd
import numpy as np
import matplotlib.pyplot as plt
import sys
import argparse
from datetime import datetime

def main(argv):

    global path_head 

    # Get relevant restraint points
    df_pts = pd.read_csv("restraint_pts.csv")

    # Get sampled collective variables in one table
    df_cat = pd.DataFrame(columns=["time", "dot-open", "dot-closed"])
    for w in range(1,151):
        for r in range(1,5):
            file = f"window{ w }/run{ r }/COLVAR_{ w }.dat"
            if os.path.exists(file):
                df_new = add_colvar_data(w, r, file)

                df_cat = pd.concat([df_cat, df_new])

    plot_sampling(df_pts, df_cat)

    return None

def add_colvar_data(window, run, file):
    """Adds COLVAR data from the sampled window.
    """
    df = pd.read_csv(file, comment="#", delim_whitespace=True,
                       names=["time", "dot-open", "dot-closed",
                              "res-bias", "res-force"])
    df["window"] = window
    df["run"] = run

    return df.iloc[10:]

def plot_sampling(df_pts, df_cat):
    # Plot the two products over the traj
    fig, ax = plt.subplots(constrained_layout=True, figsize=(12,8))

    # Get the date for record keeping
    date = datetime.now().strftime("%Y-%m-%d")

    # Plot the sampling for the 2D reaction coordinate with data from
    # the COLVAR outputs
    ax.scatter(df_cat["dot-open"], df_cat["dot-closed"],
                marker=".", color="#d86d57", 
                label=f"current sampling { date }")

    # Plot all the restraint points
    ax.scatter(df_pts["OpenPoints"], df_pts["ClosedPoints"], 
               marker="o", color="#e0e0e0", edgecolors="#404040",
               s=150, lw=3, label="restraint points")

    # Plot settings
    ax.tick_params(axis='y', labelsize=18, direction='in', width=2, \
                    length=5, pad=10)
    ax.tick_params(axis='x', labelsize=18, direction='in', width=2, \
                    length=5, pad=10)
    for i in ["top","bottom","left","right"]:
        ax.spines[i].set_linewidth(2)
    ax.grid(True)
    ax.set_xlabel(r"$\vec{\upsilon} \cdot \vec{\upsilon}_{open}$ (nm$^2$)", labelpad=5, fontsize=24)
    ax.set_ylabel(r"$\vec{\upsilon} \cdot \vec{\upsilon}_{closed}$ (nm$^2$)", labelpad=5, fontsize=24)
    plt.legend(fontsize=18, ncol=3)
    
    plt.savefig(f"{ path_head }/current_sampling_{ date }.png", dpi=300)
    plt.close()

    return None

if __name__ == '__main__':
    main(sys.argv)
```
