## Supplementary material for "*Clostridioides difficile* Toxins Unhinged: Allosterically Switchable Network Orients *β*-flap": Scripts, Structures etc.: wham.py

```
# This script was downloaded from https://github.com/plumed/masterclass-21-3.git

import numpy as np

def wham(bias,
        *,
        frame_weight=None,
        traj_weight=None,
        T: float = 1.0,
        maxiter: int = 1000,
        threshold: float = 1e-40,
        verbose: bool = False):

    nframes = bias.shape[0]
    ntraj = bias.shape[1]

    # default values
    if frame_weight is None:
        frame_weight = np.ones(nframes)
    if traj_weight is None:
        traj_weight = np.ones(ntraj)

    assert len(traj_weight) == ntraj
    assert len(frame_weight) == nframes

    # divide by T once for all
    shifted_bias = bias/T
    # track shifts
    shifts0 = np.min(shifted_bias, axis=0)
    shifted_bias -= shifts0[np.newaxis,:]
    shifts1 = np.min(shifted_bias, axis=1)
    shifted_bias -= shifts1[:,np.newaxis]

    # do exponentials only once
    expv = np.exp(-shifted_bias)

    Z = np.ones(ntraj)

    Zold = Z

    if verbose:
        sys.stderr.write("WHAM: start\n")
    for nit in range(maxiter):
        # find unnormalized weights
        weight = 1.0/np.matmul(expv, traj_weight/Z)*frame_weight
        # update partition functions
        Z = np.matmul(weight, expv)
        # normalize the partition functions
        Z /= np.sum(Z*traj_weight)
        # monitor change in partition functions
        eps = np.sum(np.log(Z/Zold)**2)
        Zold = Z
        if verbose:
            sys.stderr.write("WHAM: iteration "+str(nit)+" eps "+str(eps)+"\n")
        if eps < threshold:
            break
    nfev=nit
    logW = np.log(weight) + shifts1

    if verbose:
        sys.stderr.write("WHAM: end")

    return {"logW":logW, "logZ":np.log(Z)-shifts0, "nit":nit, "eps":eps}
```
