## Supplementary material for "*Clostridioides difficile* Toxins Unhinged: Allosterically Switchable Network Orients *β*-flap": Scripts, Structures etc.: WindowAves.py

```
import os
import pandas as pd
import numpy as np
import matplotlib.pyplot as plt
import sys
import argparse
import pandas as pd
import MDAnalysis as mda
from MDAnalysis.analysis import rms, align
from MDAnalysis.analysis.distances import distance_array
sys.path.insert(0, "/home/lf1071fu/project_b3/ProjectB3")
import config.settings as config
from tools import utils, traj_funcs
import matplotlib.colors as mcolors
import seaborn as sns
import re

def main(argv):

    try:
        parser = argparse.ArgumentParser()

        parser.add_argument("-p", "--path",
                            action = "store",
                            dest = "path",
                            default = ("umbrella/holo_state/nobackup"),
                            help = ("Set relative path to the data "
                                "directory."))
        parser.add_argument("-f", "--figpath",
                            action = "store",
                            dest = "fig_path",
                            default = ("umbrella/holo_state"),
                            help = ("Set a relative path destination for"
                                " the figure."))
        parser.add_argument("-s", "--state",
                            action = "store",
                            dest = "state",
                            default = "holo",
                            help = ("Select state as 'apo', 'holo' or "
                                "'apo-K57G' or 'holo-K57G'."))
        parser.add_argument("-r", "--recalc",
                            action = "store_true",
                            dest = "recalc",
                            default = False,
                            help = ("Chose whether the trajectory arrays"
                                " should  be recomputed."))
        parser.add_argument("-e", "--extra",
                            action = "store_true",
                            dest = "extra_contacts",
                            default = False,
                            help = ("Additional contacts to dataframe?"))
        args = parser.parse_args()

    except argparse.ArgumentError:
        print("Command line arguments are ill-defined, please check the arguments")
        raise

    global path_head, state, recalc

    recalc = args.recalc
    home = f"{ config.data_head }/{ args.path }"
    fig_path = f"{ config.figure_head }/{ args.fig_path }"
    state = args.state
    extra_contacts = args.extra_contacts
    utils.create_path(fig_path)

    # Get values for the restraint points
    points = pd.read_csv(f"{ home }/restraint_pts.csv")
    df_path = f"{ home }/windows_averages_k232_n740.csv"

    if recalc or not os.path.exists(df_path) or extra_contacts: 

        if extra_contacts:

            df = pd.read_csv(df_path)

        else: 

            df = points[["Window", "OpenPoints", "ClosedPoints"]]

            # Repeat each row in df four times for run replicates
            df = df.iloc[df.index.repeat(4)]
            df = df.reset_index(drop=True)
            df['Run'] = [1, 2, 3, 4] * (len(df) // 4)

        open_state = mda.Universe(f"{ config.struct_head }/open_ref_state.pdb")
        closed_state = mda.Universe(f"{ config.struct_head }/closed_ref_state.pdb")

        # Indicies of the inflexible residues
        core_res, core = traj_funcs.get_core_res()

        topol = f"{ home }/topol_protein.top"

        c = 0
        nw = {"apo" : 181, "holo" : 170, "apo-K57G" : 120, "holo-K57G" : 120}

        for window in np.arange(1, nw[state] + 1):
            print(f"Window { window }")
            for run in [1,2,3,4]:
                
                run_path = f"{ home }/window{ window }/run{ run }"

                # Load in universe objects for the simulation run
                u = mda.Universe(topol, f"{ run_path }/fitted_traj.xtc", 
                        topology_format='ITP')
                protein = u.select_atoms("protein")
                align.AlignTraj(u, protein, select=core, in_memory=True).run()

                if extra_contacts:

                    # Add columns for the desired additional contacts
                    ave_sasas = get_sasas(u, run_path, f"w{ window }_r{ run }",
                                        holo=("holo" in state))
                    for col, ave_sasa in ave_sasas.items():
                        df.loc[c, col] = ave_sasa

                else:    

                    # Write out some sample snapshots
                    with mda.Writer(f"{ run_path }/snap_shots.pdb", protein.n_atoms) as W:
                        for ts in u.trajectory:
                            if ((ts.time % 5000) == 0):
                                W.write(protein)

                    # Determine and store the radius of gyration
                    ave_r_gyr = get_rgyr(u)
                    df.loc[c, "ave_Rg"] = ave_r_gyr

                    # Determine and store the RMSD to ref structures
                    ave_open = get_rmsd(u, open_state, core, ["backbone and (resid 195-218)"], "open")
                    ave_closed = get_rmsd(u, closed_state, core, ["backbone and (resid 195-218)"], "closed")
                    ave_open_alpha = get_rmsd(u, open_state, core, ["backbone and (resid 219-233)"], "open")
                    ave_closed_alpha = get_rmsd(u, closed_state, core, ["backbone and (resid 219-233)"], "closed")
                    df.loc[c, "ave_RMSD_open"] = ave_open
                    df.loc[c, "ave_RMSD_closed"] = ave_closed
                    df.loc[c, "ave_RMSD_open_alpha"] = ave_open_alpha
                    df.loc[c, "ave_RMSD_closed_alpha"] = ave_closed_alpha

                    # Determine the distances for critical contacts
                    ave_ds = get_contact_dists(u)
                    for col, ave_d in ave_ds.items():
                        df.loc[c, col] = ave_d

                    # Determine SASA for residues of interest
                    # ave_sasas = get_sasas(u, run_path, f"w{ window }_r{ run }",
                    #                     holo=("holo" in state))
                    # for col, ave_sasa in ave_sasas.items():
                    #     df.loc[c, col] = ave_sasa

                c += 1
      
        if extra_contacts:          
            df.to_csv(f"{ home }/windows_averages_sasa.csv")
        else:
            df.to_csv(df_path)

    else: 

        df = pd.read_csv(df_path)

    print(df.columns)
    # print("Min:", np.min(df), "\nMax:", 
    #     np.max(df))

    # Set style for paper format 
    sns.set_context("paper") 

    #plot_Rgyr(df, fig_path)
    #plot_rmsd(df, "open", fig_path)
    #plot_rmsd(df, "closed", fig_path)
    for col in df.columns:
        if any(word in col for word in ["Points", "Unnamed", "Window", "Run"]):
            continue
        plot_averages(df, col, fig_path)
    plot_windows(df, fig_path)

def get_rgyr(u):
    """Determine the average radius of gyration.

    The radius of gyration is a measure of how compact the 
    structure is, such that an increase may indicate unfolding 
    or opening.

    Parameters
    ----------
    u : MDAnalysis.core.universe.Universe
        The universe object, with the trajectory loaded in.

    Returns
    -------
    r_gyr : float
        The averageradius of gyration in units of Angstrom.
    
    """
    r_gyr = []
    time_ser = []
    protein = u.select_atoms("protein")
    for ts in u.trajectory:
       r_gyr.append(protein.radius_of_gyration())
    r_gyr = np.array(r_gyr)

    ave_r_gyr = np.mean(r_gyr)

    return ave_r_gyr

def get_rmsd(system, reference, alignment, group, ref_state):
    """Determines the rmsd over the trajectory against a reference structure.

    The MDAnalysis.analysis.rms.results array is saved as a numpy array file,
    which can be loaded if it has alreay been determined.

    Parameters
    ----------
    system : MDAnalysis.core.universe
        The universe object, with the trajectory loaded in.
    reference : MDAnalysis.core.universe
        The topology of the reference structure.
    alignment : MDAnalysis.core.groups.AtomGroup
        The group for the alignment, i.e. all the alpha carbons.
    group : MDAnalysis.core.groups.AtomGroup
        The group for the RMSD, e.g. the beta flap residues.
    ref_state : str
        The reference state as a conformational description, i.e. "open" or "closed".

    Returns
    -------
    rmsd_arr : np.ndarray
        A timeseries of the RMSD against the given reference, for the given
        atom group.
    """
    if type(group) != list:
        group = [group]
    R = rms.RMSD(system,
                 reference,  # reference universe or atomgroup
                 select=alignment,  # group to superimpose and calculate RMSD
                 groupselections=group)  # groups for RMSD
    R.run()

    rmsd_ave = np.mean(R.results.rmsd[:,3])

    return rmsd_ave

def get_contact_dists(u):
    """Calculates the distance for critical contacts.

    Parameters
    ----------
    system : MDAnalysis.core.universe
        The universe object, with the trajectory loaded in.
    
    Returns
    -------
    ave_d : (float) dict
        The mean distance for tabulated critical contacts.
    """

    # A list of tuples for selection strings of the contacts of interest
    from config.settings import selections_k232

    averages = {}

    for key, select in selections_k232.items():

        if ("IP6" in key) & ("holo" not in state):
            continue
        elif ("K57" in key or "K600" in key) & ("K57G" in state):
            continue

        # Define the distance using the tuple of selection strings
        sel_a = u.select_atoms(select[0])
        sel_b = u.select_atoms(select[1])

        # Iterate over trajectory framse to get the time series
        distances = np.zeros(u.trajectory.n_frames)
        for ts in u.trajectory:
            d = distance_array(sel_a.positions, sel_b.positions)
            # Use the smallest pair distance
            distances[ts.frame] = np.min(d)

        ave_d = np.mean(distances, axis=0)
        averages[key] = ave_d

    return averages

def get_sasas(u, path, sim_name, holo=False):
    """Calculates the SASA for residues.

    Parameters
    ----------
    system : MDAnalysis.core.universe
        The universe object, with the trajectory loaded in.
    path : str
        Path to the individual window data
    sim_name : str
        Name used for the .tpr file for the window.
    
    Returns
    -------
    ave_sasas : (float) dict
        The mean SASA for tabulated residues.

    """
    import subprocess
    # A dictionary for the residues this interesting SASA data
    resids_water = {"W761 SASA" : 218, "E743 SASA" : 200, "H757 SASA" : 214, 
        "E749 SASA" : 206, "D771 SASA" : 228, "E765 SASA" : 222, 
        "D756 SASA" : 213, "R751 SASA" : 208}

    ave_sasas = {}

    for key, resid in resids_water.items():

        # Use a subprocess to get SASA for a residue using the gromacs tool
        sasa_path = f"{ path }/sasa_{ resid }.xvg"

        # Use gromacs subprocess to get SASA data
        if not os.path.exists(sasa_path) or recalc:

            # Setup variables for command-line arguements
            p = path
            if holo:
                surface = 24 
                output = 25
            else:
                surface = 1
                output = 19

            # command-line arguements
            gmx_ndx = (f"""echo "ri { resid }\nq" | gmx22 make_ndx -f """
                f"""{ p }/{ sim_name }.tpr -o { p }/index_{ resid }.ndx """
                """-nobackup""")
            gmx_sasa = (f"gmx22 sasa -f { p }/fitted_traj.xtc -s " 
                f"{ p }/{ sim_name }.tpr -o { p }/sasa_{ resid }.xvg"
                f" -or { p }/res{ resid }_sasa.xvg -surface { surface } "
                f"-output { output } -n { p }/index_{ resid }.ndx "
                f"-nobackup")

            # Calculate SASA with gmx sasa using a subprocess
            if holo:
                gmx_ndx += f" -n { p }/index.ndx"
            subprocess.run(gmx_ndx, shell=True)
            subprocess.run(gmx_sasa, shell=True)

        # In case something went wrong in the subprocess...
        utils.validate_path(sasa_path, warning="Use 'gmx sasa' to extract "
            "sasa data.\n")

        # Read in the gromacs analysis output
        gmx_sasa = np.loadtxt(sasa_path, comments=["#", "@"])
        sasa_data = gmx_sasa[:,2]

        # Get the window average into a dictionary
        ave_sasa = np.mean(sasa_data, axis=0)
        print(f"Average { key } for { sim_name }: \n\t{ ave_sasa }")
        ave_sasas[key] = ave_sasa

    return ave_sasas

def plot_Rgyr(df, path):
    fig, ax = plt.subplots(constrained_layout=True, figsize=(12,8))

    ave_Rg = df.groupby('Id')['ave_Rg'].min().values

    d = ax.scatter(df[df["Run"] == 1]["OpenPoint"], 
                    df[df["Run"] == 1]["ClosedPoint"], c=ave_Rg, 
                    cmap="cividis", marker="o", edgecolors="#404040", s=200, lw=2)

    # Colormap settings
    cbar = plt.colorbar(d)
    cbar.set_label(r'Radius of Gyration ($\AA$)', fontsize=32, labelpad=10)

    ax.set_xlabel(r"$\xi_1$ (nm$^2$)")
    ax.set_ylabel(r"$\xi_2$ (nm$^2$)")

    plt.savefig(f"{ path }/window_Rgyr.png", dpi=300)
    plt.close()

def plot_rmsd(df, ref_state, path):
    fig, ax = plt.subplots(constrained_layout=True, figsize=(12,8))

    rmsd = df.groupby('Id')[f"ave_RMSD_{ ref_state }"].min().values

    d = ax.scatter(df["OpenPoint"][df["Run"] == 1], 
                    df["ClosedPoint"][df["Run"] == 1], c=rmsd, 
                    cmap="cividis", marker="o", edgecolors="#404040", s=200, lw=2)

    # Colormap settings
    cbar = plt.colorbar(d)
    cbar.set_label(f'RMSD$_{ ref_state }$ ($\AA$)', fontsize=32, labelpad=10)

    ax.set_xlabel(r"$\xi_1$ (nm$^2$)")
    ax.set_ylabel(r"$\xi_2$ (nm$^2$)")

    plt.savefig(f"{ path }/window_RMSD_{ ref_state }.png", dpi=300)
    plt.close()

def plot_averages(df, col, path):
    fig, ax = plt.subplots(constrained_layout=True, figsize=(2.5,2))

    ave_val = df.groupby('Window')[col].mean().values
    #print(col, min(ave_val), max(ave_val))

    # Fix the col name to add 543 to resnums
    def add_543(match):
        num = int(match.group())
        return str(num + 543)
    #col = re.sub(r'(?<!IP6)(\d+)', add_543, col)
    #col = col.replace("--", "-")

    if "SASA" in col:
        print(col)
        d = ax.tricontourf(df[df["Run"] == 1]["OpenPoints"], 
                    df[df["Run"] == 1]["ClosedPoints"], ave_val, 10,
                    cmap="coolwarm")
        # Colormap settings
        cbar = plt.colorbar(d)
        cbar.set_label(f"{ col } " + r"(nm$^2$)", labelpad=3, fontsize=8)

    elif "K775--IP6" in col:
        print(col)
        d = ax.tricontourf(df[df["Run"] == 1]["OpenPoints"], 
            df[df["Run"] == 1]["ClosedPoints"], ave_val, 10, cmap="coolwarm", 
            norm=mcolors.Normalize(vmin=0, vmax=15))
        # Colormap settings
        cbar = plt.colorbar(d)
        cbar.set_label(f"{ col } " + r"(nm$^2$)", labelpad=3, fontsize=8)

    elif "K775--E692" in col:
        print(col)
        d = ax.tricontourf(df[df["Run"] == 1]["OpenPoints"], 
            df[df["Run"] == 1]["ClosedPoints"], ave_val, 10, cmap="coolwarm", 
            norm=mcolors.Normalize(vmin=2, vmax=12), extend="max")
        # Colormap settings
        cbar = plt.colorbar(d)
        cbar.set_label(f"{ col } " + r"(nm$^2$)", labelpad=3, fontsize=8)
    
    # elif "K775--E796" in col:
    #     print(col)
    #     d = ax.tricontourf(df[df["Run"] == 1]["OpenPoints"], 
    #         df[df["Run"] == 1]["ClosedPoints"], ave_val, 10, cmap="coolwarm", 
    #         norm=mcolors.Normalize(vmin=0, vmax=15))
    #     # Colormap settings
    #     cbar = plt.colorbar(d)
    #     cbar.set_label(f"{ col } " + r"(nm$^2$)", labelpad=3, fontsize=8)

    else: 
        d = ax.tricontourf(df[df["Run"] == 1]["OpenPoints"], 
                    df[df["Run"] == 1]["ClosedPoints"], ave_val, 10,
                    cmap="coolwarm")

        # Colormap settings
        cbar = plt.colorbar(d) # , ticks=np.arange(1, 25.1, 4), extend="both")
        cbar.set_label(f"{ col } " + r"($\AA$)", labelpad=3,
            fontsize=8)

    # Add axes labels, tick positions 
    plt.xticks([0, 2, 4, 6], fontsize=6)
    plt.yticks([0, 2, 4, 6], fontsize=6)
    ax.set_xlabel(r"$\xi_1$ (nm$^2$)", labelpad=3, fontsize=8)
    ax.set_ylabel(r"$\xi_2$ (nm$^2$)", labelpad=3, fontsize=8)
    ax.set_xlim(0,6)
    ax.set_ylim(0,6)
    ax.set_aspect('equal', adjustable='box')
    cbar.ax.tick_params(labelsize=6, color="#B0B0B0")
    cbar.outline.set_linewidth(1)

    utils.save_figure(fig, f"{ path }/window_ave_{ col }.png")
    plt.close()

def plot_windows(df, path):
    fig, ax = plt.subplots(constrained_layout=True, figsize=(8,8))

    window_labs = df.groupby('Window')["Window"].mean().values.astype(int)
    x = np.array(df[df["Run"] == 1]["OpenPoints"])
    y = np.array(df[df["Run"] == 1]["ClosedPoints"])

    d = ax.scatter(x,y, marker="o", edgecolors="#404040", s=200, lw=2)

    # Add labels to each point
    for i, label in enumerate(window_labs):
        ax.annotate(label, (x[i], y[i]), textcoords="offset points", xytext=(0, 10), ha='center')

    ax.set_xlabel(r"$\xi_1$ (nm$^2$)")
    ax.set_ylabel(r"$\xi_2$ (nm$^2$)")

    utils.save_figure(fig, f"{ path }/window_labels.png")
    plt.close()

if __name__ == '__main__':
    main(sys.argv)
```
