## Supplementary material for "*Clostridioides difficile* Toxins Unhinged: Allosterically Switchable Network Orients *β*-flap": Scripts, Structures etc.: ComboPlots.py

```
import sys
import numpy as np
import os
import matplotlib.pyplot as plt
import matplotlib.patheffects as pe
import pandas as pd
import argparse
import MDAnalysis as mda
sys.path.insert(0, "/home/lf1071fu/project_b3/ProjectB3")
import config.settings as cf
from tools import utils, traj_funcs
import subprocess
import seaborn as sns

def main(argv):

    try:
        parser = argparse.ArgumentParser()

        parser.add_argument("-s", "--state",
                            action = "store",
                            dest = "state",
                            default = "apo",
                            help = ("Chose the type of simulation i.e. 'unbiased'",
                                "or 'mutation' or 'tcda'."))
        parser.add_argument("-w", "--sasa",
                            action = "store_true",
                            dest = "sasa",
                            default = False,
                            help = ("Make combo plots for SASA?"))
        
        args = parser.parse_args()

    except argparse.ArgumentError:
        print("Command line arguments are ill-defined, please check the arguments")
        raise

    global styles, fig_path, state_name

    # Assign group selection from argparse
    state_name = args.state
    sasa = args.sasa
    
    # Set up some path variables for state 'unbiased' or 'mutation'
    if state_name == "unbiased":
        fig_path = f"{ cf.figure_head }/unbiased_sims/combo_{ state_name }"
        table_path = f"{ cf.data_head }/unbiased_sims/combo_{ state_name }"
        state_paths = {
            "apo-closed" : f"{ cf.data_head }/unbiased_sims/apo_closed/analysis",
            "holo-closed" : f"{ cf.data_head }/unbiased_sims/holo_closed/analysis",
            "holo-open" : f"{ cf.data_head }/unbiased_sims/holo_open/analysis",
            "apo-open" : f"{ cf.data_head }/unbiased_sims/apo_open/analysis"} 
    elif state_name == "mutation":
        fig_path = f"{ cf.figure_head }/mutation/combo_{ state_name }"
        table_path = f"{ cf.data_head }/unbiased_sims/mutation/combo_{ state_name }"
        state_paths = {
            "K600G" : f"{ cf.data_head }/unbiased_sims/mutation/K57G/analysis", 
            "E743G" : f"{ cf.data_head }/unbiased_sims/mutation/E200G/analysis",
            "K600G/E743G" : f"{ cf.data_head }/unbiased_sims/mutation/double_mut/analysis",}
    elif state_name == "tcda":
        fig_path = f"{ cf.figure_head }/tcda/combo_{ state_name }"
        table_path = f"{ cf.data_head }/tcda/combo_{ state_name }"
        state_paths = {
            "TcdA-apo-open" : f"{ cf.data_head }/tcda/open_apo/analysis", 
            "TcdA-apo-closed" : f"{ cf.data_head }/tcda/closed_apo/analysis",
            "TcdA-holo-open" : f"{ cf.data_head }/tcda/open_holo/analysis", 
            "TcdA-holo-closed" : f"{ cf.data_head }/tcda/closed_holo/analysis"
            } 
    utils.create_path(fig_path)

    states = [State(name, path) for name, path in state_paths.items()]

    # A list of tuples for selection strings of the contacts of interest
    # and a dictionary for styles related to consistent plotting
    if "tcda" in state_name:
        selections = cf.selections_tcda
    elif "mutations" in state_name:
        selections = {"C698--H757" : ("resid 155 and name SG", 
            "resid 214 and (name ND1 or name NE2)"),
            "E592--W761" : ("resid 218 and name NE1", "resid 49 and name OE*"),}
    else:
        selections = cf.selections
    styles = cf.styles

    # Add data into the State objects
    for state in states:

        # Load in np.array data produced by BasicMD.py 
        #load_arrs(state)

        # Get arrays for the contacts of interest
        get_dist_arrs(state, selections)

        # Load in SASA data computed with gmx sasa
        if sasa:
            if "tcda" in state_name:
                for i in [221,203,60,217,224,209,211,52,225,216,231]:
                    get_sasa(state, i)
            else:
                for i in [218, 200, 57, 214, 206, 208, 222, 228, 
                    213, 221, 155]:
                    get_sasa(state, i)

    # Make plots for time series
    # stride = 50
    # plot_rmsf(states)
    # plot_rmsd_time(states, stride=stride)
    # plot_rgyr(states, stride=stride)

    # plot_salt_bridges(states, stride=stride)
    # plot_hbonds(states, stride=stride)
    
    # Set style for paper format 
    sns.set_context("paper") 

    # Make histograms of the contact distances
    for contact in selections.keys():

        if "C698--H757" in contact or "C699--H758" in contact:
            plot_hist(states, contact, ylim=[0,.69])
            plot_hist(states, contact, legend=False)
        else:
            plot_hist(states, contact)
            plot_hist(states, contact, legend=False)

    # Make histograms of the SASA for W218 and E200
    if sasa:
        if "tcda" in state_name:
            sasa_res = [(221,"W"),(203,"E"),(60,"K"),(217,"H"),
                (224,"K"), (209,"E"), (52,"E"),(211,"R"),(225,"E"),
                (216,"A"), (231,"D")]
            for i,j in sasa_res:
                plot_sasa(states, i, j)
                plot_sasa(states, i , j, legend=False)
        else:
            sasa_res = [(218,"W"),(200,"E"),(57,"K"),(214,"H"),
                (206,"E"), (208,"R"), (222,"E"), (228,"D"), (213,"D"),
                (221,"K"), (155,"C") ]# , (49,"E")]
            for i,j in sasa_res:
                plot_sasa(states, i, j)
                plot_sasa(states, i , j, legend=False)

    # Make a table of the values
    for state in states:
        df = pd.DataFrame()
        for contact in selections.keys():
            if ("IP6" in contact) & ("holo" not in state.name):
                continue
            if ("K57" in contact or "K600" in contact) & ("K600G" in state.name):
                continue
            if ("E200" in contact or "E743" in contact) & ("E743G" in state.name):
                continue   
            d = state.get_array(contact)
            df[contact] = d
        if sasa:
            for resid, aa in sasa_res:
                if "tcda" in state_name:
                    toxinnum_resid = resid + 541
                else: 
                    toxinnum_resid = resid + 543
                arr = state.get_array(f"sasa_{ resid }")
                df[f"{ aa }{toxinnum_resid} SASA"] = arr
                print(f"{ aa }{toxinnum_resid} SASA")
        df.to_csv(f"{ table_path }_{ state.name }.csv")

    print(df.columns)

    return None

class State:
    def __init__(self, name, path):
        self.name = name
        self.path = path
        self.arrays = {}
        self.data_path = f"{ os.path.dirname(path) }/nobackup"
        if "holo" in name:
            self.ligand = "holo"
        else:
            self.ligand = "apo"
        if "open" in name:
            self.conform = "open"
            self.clabel = "raised"
        else:
            self.conform = "closed"
            self.clabel = "lowered"

    def add_array(self, name, array):
        self.arrays[name] = array

    def get_array(self, name):
        return self.arrays[name]

    def remove_array(self, name):
        del self.arrays[name]

def get_dist_arrs(state, selections):
    """Get distance arrays for contacts of interest.

    Parameters
    ----------
    s : State
        Each simulation state gets a state object in which the np
        arrays are stored.
    
    Returns
    -------
    None.

    """
    # Load in some reference structures
    open_state = mda.Universe(f"{ cf.struct_head }/open_ref_state.pdb")
    closed_state = mda.Universe(f"{ cf.struct_head }/closed_ref_state.pdb")
    ref_state = mda.Universe(f"{ cf.struct_head }/alignment_struct.pdb")

    # Indicies of the inflexible residues
    core_res, core = traj_funcs.get_core_res()

    get_contacts(state, selections)

    return None

def load_arrs(state):
    """Add pre-computed arrays to State object.

    Parameters
    ----------
    s : State
        Each simulation state gets a state object in which the np
        arrays are stored. 

    Returns
    -------
    None. 

    """
    analysis_path = state.path

    # Read calculated outputs from numpy arrays
    if "open" in state.path:
        np_files = { "RMSD" : f"{ analysis_path }/rmsd_open.npy"}
    else: 
        np_files = { "RMSD" : f"{ analysis_path }/rmsd_closed.npy"}
    np_files.update({"RMSF" : f"{ analysis_path }/rmsf.npy", 
                    "calphas" : f"{ analysis_path }/calphas.npy",
                    "rad_gyr" : f"{ analysis_path }/rad_gyration.npy", 
                    "salt" : f"{ analysis_path }/salt_dist.npy", 
                    "timeser" : f"{ analysis_path }/timeseries.npy",
                    "hbonds" : f"{ analysis_path }/hpairs.npy"
                    })

    # Load numpy arrays into the State object
    if all(list(map(lambda x : os.path.exists(x), np_files.values()))):

        for key, file in np_files.items(): 
            state.add_array(key, np.load(file, allow_pickle=True))

    else: 

        print("Missing Numpy files! Run BasicMD.py first.")
        exit(1)

def get_contacts(state, selections):
    """Add timeseries of critical contacts to the State object.

    Parameters
    ----------
    s : State
        Each simulation state gets a state object in which the np
        arrays are stored. 
    selections : ((str) tuple) dict
        Selection strings for critical contacts are given as tuples.    
    """
    data_path = state.data_path

    if "holo" in state.name:
        topol = f"{ data_path }/topol_Pro_Lig.top"
    else:
        topol = f"{ data_path }/topol_protein.top"
    xtc = f"{ data_path }/fitted_traj_100.xtc"

    u = mda.Universe(topol, xtc, topology_format='ITP')
    u.transfer_to_memory()
    u = traj_funcs.do_alignment(u)

    from MDAnalysis.analysis.distances import distance_array

    for key, select in selections.items():

        # Skip over inapplicable selections
        if ("IP6" in key) & ("holo" not in state.name):
            continue
        if ("K600" in key) & ("K600G" in state.name):
            continue
        if ("K600" in key) & ("K600G/E743G" in state.name):
            continue
        if ("E743" in key) & ("E743G" in state.name):
            continue   
        if ("E743" in key) & ("K600G/E743G" in state.name):
            continue

        # Define the distance using the tuple of selection strings
        sel_a = u.select_atoms(select[0])
        sel_b = u.select_atoms(select[1])
    
        # Iterate over trajectory framse to get the time series
        distances = np.zeros(u.trajectory.n_frames)
        for ts in u.trajectory:
            d = distance_array(sel_a.positions, sel_b.positions)
            # Use the smallest pair distance
            distances[ts.frame] = np.min(d)
        
        # Update data in the State object
        state.add_array(key, distances)
    
    # Also store array of the time series
    time_ser = np.zeros(u.trajectory.n_frames)
    for ts in u.trajectory:
        time_ser[ts.frame] = ts.time
    state.add_array("timeseries", time_ser)

    return None

def get_sasa(state, resid):
    """Adds SASA data into the State object.

    This loads or computes with a subprocess the SASA using gromacs.

    Parameters
    ----------
    state : State
        The State object is needed for array data to be included.

    Returns
    -------
    None.

    """
    sasa_path = f"{ state.path }/../analysis/sasa_{ resid }.xvg"

    # Use gromacs subprocess to get SASA data
    if not os.path.exists(sasa_path):

        # Setup variables for command-line arguements
        p = state.data_path
        sim_name = f"{ p }/{ state.name.replace('-','_') }"
        if "holo" in state.name:
            gmx_ndx_holo = (f"""echo "1 | 13\nq" | gmx22 make_ndx -f """
            f"""{ sim_name }.tpr -o { p }/index.ndx -nobackup""")
            surface = 24 
            output = 25
        else:
            surface = 1
            output = 19

        # command-line arguements
        gmx_ndx = (f"""echo "ri { resid }\nq" | gmx22 make_ndx -f """
            f"""{ sim_name }.tpr -n { p }/index.ndx -o { p }/index_{ resid }.ndx -nobackup""")
        gmx_sasa = (f"gmx22 sasa -f { p }/fitted_traj_100.xtc -s " 
            f"{ sim_name }.tpr -o { p }/../analysis/sasa_{ resid }.xvg"
            f" -or { p }/../analysis/res{ resid }_sasa.xvg "
            f"-surface { surface } -output { output }"
            f" -n { p }/index_{ resid }.ndx -nobackup")

        # Calculate SASA with gmx sasa using a subprocess
        if "holo" in state.name:
            subprocess.run(gmx_ndx_holo, shell=True)
        subprocess.run(gmx_ndx, shell=True)
        subprocess.run(gmx_sasa, shell=True)

    # In case something went wrong in the subprocess...
    utils.validate_path(sasa_path, warning="Use 'gmx sasa' to extract "
        "sasa data.\n")

    # Read in the gromacs analysis output
    gmx_sasa = np.loadtxt(sasa_path, comments=["#", "@"])
    sasa_data = gmx_sasa[:,2]

    # Print out the average for the state
    print(f"{ state.name } { resid } : { np.round(np.average(sasa_data),2) } "
          f"+/- { np.round(np.std(sasa_data), 2) }") 

    # Add SASA data to the object
    state.add_array(f"sasa_{ resid }", sasa_data)

    return None

def plot_rmsf(states):
    """Makes an RMSF plot.

    Parameters
    ----------
    states : (State object) list
        The list of State objects contains the relevant numpy arrays for each state, 
        accessed via the "get_array()" module.

    Returns
    -------
    None.

    """
    fig, ax = plt.subplots(constrained_layout=True, figsize=(2.5,1.25))

    for i, c in enumerate(states):

        # Get state data
        key = c.name
        calphas = c.get_array("calphas")
        rmsf = c.get_array("RMSF")

        ax.plot(calphas, rmsf, lw=6, color=styles[key][0], 
                label=key, alpha=0.8, dash_capstyle='round', 
                ls=styles[key][1])

    # Plot settings
    ax.tick_params(axis='y', labelsize=18, direction='in', width=2, \
                    length=5, pad=10)
    ax.tick_params(axis='x', labelsize=18, direction='in', width=2, \
                    length=5, pad=10)
    for i in ["top","bottom","left","right"]:
        ax.spines[i].set_linewidth(2)
    ax.grid(True)
    ax.set_xlabel(r"Residue number", labelpad=5, fontsize=24)
    ax.set_ylabel(r"RMSF ($\AA$)", labelpad=5, fontsize=24)
    bottom, top = ax.get_ylim()
    ax.vlines([195,217.5], bottom, top, linestyles="dashed", lw=3,
              colors="#FF1990")
    ax.vlines([219.5,231], bottom, top, linestyles="dashed", lw=3,
              colors="#dba61f")
    ax.set_ylim(-1,6)
    plt.legend(fontsize=20, loc=2)

    utils.save_figure(fig, f"{ fig_path }/rmsf.png")
    plt.close()

    return None

def plot_rmsd_time(states, stride=1):
    """Makes a timeseries RMSD plot, against the respective reference structure.

    Parameters
    ----------
    states : (State object) list
        The list of State objects contains the relevant numpy arrays for each state, 
        accessed via the "get_array()" module.
    stride : int
        Set the stride for the data, increasing if it is too much data in the plot

    Returns
    -------
    None. 

    """
    fig, ax = plt.subplots(constrained_layout=True, figsize=(2.5,1.25))

    colors = {"open" : "#EAAFCC", "closed" : "#A1DEA1"}
    linestyles = {"holo" : "-", "apo" : "--"}

    for i, c in enumerate(states):

        # Get state data
        key = c.name
        rmsd = c.get_array("RMSD")
        time = c.get_array("timeser")

        # Add plot for each state
        plt.plot(time[::stride], rmsd[::stride,3], lw=6, color=styles[key][0], 
                label=key, alpha=0.8, ls=styles[key][1])

    # Plot settings
    ax.tick_params(axis='y', labelsize=18, direction='in', width=2, \
                    length=5, pad=10)
    ax.tick_params(axis='x', labelsize=18, direction='in', width=2, \
                    length=5, pad=10)

    x_labels = list(map(lambda x : str(x/1e6).split(".")[0], ax.get_xticks()))
    ax.set_xticklabels(x_labels)
    for i in ["top","bottom","left","right"]:
        ax.spines[i].set_linewidth(2)
    ax.grid(True)
    ax.set_xlabel(r"Time ($\mu s$)", labelpad=5, fontsize=24)
    ax.set_ylabel(r"RMSD ($\AA$)", labelpad=5, fontsize=24)
    _, ymax = ax.get_ylim()
    ax.set_ylim(0,ymax)
    plt.legend(fontsize=20)

    utils.save_figure(fig, f"{ fig_path }/rmsd_time.png")
    plt.close()

    return None

def plot_salt_bridges(states, stride=1):
    """Makes a timeseries plot for the key salt bridge K57--E200.

    Parameters
    ----------
    states : (State object) list
        The list of State objects contains the relevant numpy arrays for each state, 
        accessed via the "get_array()" module.

    Returns
    -------
    None.

    """

    fig, ax = plt.subplots(constrained_layout=True, figsize=(2.5,1.25))

    for i, c in enumerate(states):
        
        # Get state data
        key = c.name
        sc = c.get_array("salt")
        time = c.get_array("timeser")

        # Add plot for each state
        plt.plot(time[::stride], sc[::stride,3], lw=6, color=styles[key][0], 
                label=key, alpha=0.8, ls=styles[key][1])

    # Plot settings
    ax.tick_params(axis='y', labelsize=18, direction='in', width=2, \
                    length=5, pad=10)
    ax.tick_params(axis='x', labelsize=18, direction='in', width=2, \
                    length=5, pad=10)
    x_labels = list(map(lambda x : str(x/1e6).split(".")[0], ax.get_xticks()))
    ax.set_xticklabels(x_labels)
    ax.set_xlabel(r"Time ($\mu s$)", labelpad=5, fontsize=24)
    for i in ["top","bottom","left","right"]:
        ax.spines[i].set_linewidth(2)
    ax.grid(True)
    ax.set_ylabel(r"Distance K57--E200 $(\AA)$", labelpad=5, fontsize=24)
    plt.legend(fontsize=20)

    utils.save_figure(fig, f"{ fig_path }/salt_bridge_K57-E200.png")
    plt.close()

    return None

def plot_hbonds(states, stride=1):
    """Makes a timeseries plot for the key hbond N53--E200.

    Parameters
    ----------
    states : (State object) list
        The list of State objects contains the relevant numpy arrays for each state, 
        accessed via the "get_array()" module.

    Returns
    -------
    None.

    """

    fig, ax = plt.subplots(constrained_layout=True, figsize=(2.5,1.25))

    for i, c in enumerate(states):
        
        # Get state data
        key = c.name
        hbonds = c.get_array("hbonds")
        time = c.get_array("timeser")

        # Add plot for each state
        plt.plot(time[::stride], hbonds[::stride,3], lw=6, color=styles[key][0], 
                label=key, alpha=0.8, ls=styles[key][1])

    # Plot settings
    ax.tick_params(axis='y', labelsize=18, direction='in', width=2, \
                    length=5, pad=10)
    ax.tick_params(axis='x', labelsize=18, direction='in', width=2, \
                    length=5, pad=10)
    x_labels = list(map(lambda x : str(x/1e6).split(".")[0], ax.get_xticks()))
    ax.set_xticklabels(x_labels)
    ax.set_xlabel(r"Time ($\mu s$)", labelpad=5, fontsize=24)
    for i in ["top","bottom","left","right"]:
        ax.spines[i].set_linewidth(2)
    ax.grid(True)
    ax.set_ylabel(r"Distance N53--E200 $(\AA)$", labelpad=5, fontsize=24)
    plt.legend(fontsize=20)

    utils.save_figure(fig, f"{ fig_path }/hbond_N53-E200.png")
    plt.close()

    return None

def plot_rgyr(states, stride=1):
    """Makes a Radius of Gyration plot.

    Parameters
    ----------
    states : (State) list
        The list of State objects contains the relevant numpy arrays for each state, 
        accessed via the "get_array()" module.

    Returns
    -------
    None. 

    """

    fig, ax = plt.subplots(constrained_layout=True, figsize=(6*cm,2*cm))

    for i, c in enumerate(states):

        # Get state data
        key = c.name
        r_gyr = c.get_array("rad_gyr")
        time = c.get_array("timeser")

        # Add plot for each state
        plt.plot(time[::stride], r_gyr[::stride], lw=6, color=styles[key][0], 
                label=key, alpha=0.8, ls=styles[key][1])

    # Plot settings
    ax.tick_params(axis='y', labelsize=18, direction='in', width=2, \
                    length=5, pad=10)
    ax.tick_params(axis='x', labelsize=18, direction='in', width=2, \
                    length=5, pad=10)
    x_labels = list(map(lambda x : str(x/1e6).split(".")[0], ax.get_xticks()))
    ax.set_xticklabels(x_labels)
    for i in ["top","bottom","left","right"]:
        ax.spines[i].set_linewidth(2)
    ax.grid(True)
    ax.set_xlabel(r"Time ($\mu s$)")
    ax.set_ylabel(r"$R_G$ ($\AA$)")
    plt.legend()
    y_min, ymax = ax.get_ylim()
    ax.set_ylim(y_min-0.2,ymax+0.5)

    utils.save_figure(fig, f"{ fig_path }/rad_gyration.png")
    plt.close()

    return None

def plot_hist(states, contact, bins=15, legend=True, **kwargs):
    """Makes a histogram plot of the contact all the State objects.

    Parameters
    ----------
    states : (State) list
        The list of State objects contains the relevant numpy arrays
        for each state, accessed via the "get_array()" module.
    contact : str
        Name of the contact which is visualized by the histogram. The
        string is used to access the array from the relevant
        dictionaries. 

    Returns
    -------
    None.

    """
    fig, ax = plt.subplots(figsize=(2.5,1.25))
    
    for state in states:

        key = state.name
        label = state.ligand + "-" + state.clabel
        print(contact, key)

        # Skip over inapplicable selections
        if ("IP6" in contact) & ("holo" not in key):
            continue
        if ("K57" in contact or "K600" in contact) & ("K600G" in key):
            continue
        if ("E200" in contact or "E743" in contact) & ("E743G" in key):
            continue   

        dist = state.get_array(contact)
        ax.hist(dist, bins=bins, density=True, color=styles[key][0], 
                ls=styles[key][1], histtype='step', lw=2, label=label,
                alpha=styles[key][2])
    
    if "E753--R745" in contact or "E754--R746" in contact: 
        xmin, _ = ax.get_xlim()
        ax.set_xlim(xmin,5)
    elif "N747--E753" in contact or "N748--E754" in contact:
        xmin, _ = ax.get_xlim()
        ax.set_xlim(xmin,6)

    # Plot settings
    ax.tick_params(axis='both', which='major', pad=3, labelsize=6)
    ax.set_xlabel(f"Distance { contact } " + r"($\AA$)", labelpad=3, 
        fontsize=8)
    ax.set_ylabel("Frequency", labelpad=3, fontsize=8)
    if "xlim" in kwargs:
        plt.xlim(kwargs["xlim"])
    if "ylim" in kwargs:
        plt.ylim(kwargs["ylim"])
    if legend:
        plt.legend(ncols=1, fontsize=6)
    for i in ["top","bottom","left","right"]:
        ax.spines[i].set_linewidth(1)
    ax.grid(False)

    if legend:
        utils.save_figure(fig, f"{ fig_path }/{ contact }.png")
    else: 
        no_legend_path = f"{ fig_path }/no_legend"
        utils.create_path(no_legend_path)
        utils.save_figure(fig, 
            f"{ no_legend_path }/{ contact }.png")
    plt.close()

    return None

def plot_sasa(states, resid, res_type, legend=True,):
    """Makes a histogram plot of the residue SASA data.

    Parameters
    ----------
    states : (State) list
        The list of State objects contains the relevant numpy arrays
        for each state, accessed via the "get_array()" module.
    resid : int
        The residue id for the sasa data.
    res_type : str
        The residues type, used in the plots x-axis.

    Returns
    -------
    None.

    """
    fig, ax = plt.subplots(figsize=(2.5,1.25))
    if "tcda" in state_name:
        toxinnum_resid = resid + 541
    else: 
        toxinnum_resid = resid + 543

    # Add reference data as vertical lines
    if resid == 218:
        ax.axvline(x=0.314, color="#424242", ls="solid", 
                    label="lowered-ref", lw=3)
        ax.axvline(x=0.876, color="#424242", ls="dashed", 
                    label="raised-ref", lw=3)
        ax.set_ylim(0,6)
        plt.yticks([0, 2, 4])
    ax.grid(False)

    for state in states:

        # Add a histogram for SASA of each state
        key = state.name
        label = state.ligand + "-" + state.clabel
        ax.hist(state.get_array(f"sasa_{ resid }"), bins=20, density=True, 
                color=styles[key][0], lw=2, ls=styles[key][1], 
                histtype='step', label=label, alpha=styles[key][2])
    
    # Plot settings
    ax.tick_params(axis='both', which='major', pad=3, labelsize=6)
    ax.set_xlabel(f"SASA of { res_type }{ toxinnum_resid } (nm²)", 
        labelpad=3, fontsize=8)
    ax.set_ylabel("Frequency", labelpad=3, fontsize=8)
    for i in ["top","bottom","left","right"]:
        ax.spines[i].set_linewidth(1)
    ax.set_xlim([-0.15,2])
    ax.grid(False)
    if legend:
        # if resid in [218,221]:
        #     plt.legend(ncols=2, fontsize=6)
        # #else:
        plt.legend(ncols=1, fontsize=6)
        utils.save_figure(fig, 
            f"{ fig_path }/SASA_{ toxinnum_resid }.png")
    else: 
        no_legend_path = f"{ fig_path }/no_legend"
        utils.create_path(no_legend_path)
        utils.save_figure(fig, 
            f"{ no_legend_path }/SASA_{ toxinnum_resid }.png")

    plt.close()

    return None

if __name__ == '__main__':
    main(sys.argv)
```
