## Supplementary material for "*Clostridioides difficile* Toxins Unhinged: Allosterically Switchable Network Orients *β*-flap": Scripts, Structures etc.: settings.py

```
# settings.py

# General settings
PROJECT_NAME = "ProjectB3"
DEBUG_MODE = True

# Path settings
path_head = "/home/lf1071fu/project_b3"
data_head = f"{ path_head }/simulate"
figure_head = f"{ path_head }/figures"
struct_head = f"{ path_head }/structures"

# Protein settings related to residue IDs
# Shift residue count for consistency with the full-toxin
toxin_resid = 544 
beta_flap_group = "backbone and resid 195-218"
alpha_flap_group = "backbone and resid 219-233"
active_res_group = "resid 110 or resid 155"

# Alpha carbons for the beta-flap vector 
r1, r2 = 206, 215

# Aesthetics
open_color = "#0069D3" #"#A1DEA1"
closed_color = "#a03232" #EAAFCC"
styles = {"apo-open" : ("#0069D3", "solid", 0.7), 
          "holo-open" : ("#0069D3", "dashdot", 1), 
          "apo-closed" : ("#a03232", "solid", 1),
          "holo-closed" : ("#a03232", "dashdot", 0.7),
          "K600G" : ("#EBBA34", "solid", 1),
          "E743G" : ("#8442F5", "solid", 1),
          "K600G/E743G" : ("#36B3CF", "solid", 1),
          "TcdA-apo-open" : ("#0069D3", "solid", 0.7), 
          "TcdA-apo-closed" : ("#a03232", "dashdot", 1),
          "TcdA-holo-open" : ("#0069D3", "solid", 1), 
          "TcdA-holo-closed" : ("#a03232", "dashdot", 0.7)}

# Critical contacts dictionary
selections_all = {
    "L755--E743" : ("resid 212 and name N","resid 200 and name O"),
    "K792--K775" : ("resid 249 and name NZ","resid 232 and name O"),
    "K792--S773a" : ("resid 249 and name N","resid 230 and name OG"),
    "K792--S773b" : ("resid 249 and name N","resid 230 and name O"),
    "K792--S773c" : ("resid 249 and name NZ","resid 230 and name O"),
    "S773--I769" : ("resid 230 and name N","resid 226 and name O"),
    "K792--E776" : ("resid 249 and name NZ","resid 233 and name OE*"),
    "R752--S774" : ("resid 209 and name NH*",
        "resid 231 and (name OG or name O)"),
    "R752--N740" : ("resid 209 and name NH*","resid 197 and name OD*"),
    "R752--E796" : ("resid 209 and name NH*","resid 253 and name OE*"),
    "R745--E753" : ("resid 202 and name NE","resid 210 and name OE*"),
    "K764--E766" : ("resid 221 and name NZ","resid 223 and name OE*"),
    "R751--E765" : ("resid 208 and name NH*","resid 222 and name OE*"),
    "K600--G750" : ("resid 57 and name NZ","resid 207 and name O"),
    "K600--V744" : ("resid 57 and name NZ","resid 201 and name O"),
    "R571--S748" : ("resid 28 and name NH*","resid 205 and name O"),
    # (old) beta-flap network
    "R751--N747" : ("resid 204 and (name OD1 or name ND2)",
        "resid 208 and (name N* or name O)"),
    "N747--E753" : ("resid 204 and name ND2", "resid 210 and name OE*"),
    "E753--R745" : ("resid 202 and (name NE or name NH*)",
        "resid 210 and name OE*"),
    "R745--W761" : ("resid 202 and (name NE or name NH*)",
        "resid 218 and name CZ*"),

    "I762--E766" : ("resid 219 and name CA", "resid 223 and name CA"),
    "Y742--D771" : ("resid 199 and name OH", "resid 228 and name OD*"),
    "I762--E766" : ("resid 219 and (name CD or name CG*)", 
        "resid 223 and name OE*"),
    "L755--W761" : ("resid 218 and name CA", "resid 212 and name CA"),
    "E749--N747" : ("resid 206 and name N", "resid 204 and name OD1"),
    "C698--H757" : ("resid 155 and name SG", 
        "resid 214 and (name ND1 or name NE2)"),
    "C698--H653" : ("resid 155 and name SG", 
        "resid 110 and (name ND1 or name NE2)"),
    "E592--W761" : ("resid 218 and name NE1", "resid 49 and name OE*"),
    "N596--E592" : ("resid 49 and name OE*", "resid 53 and name ND2"),
    # Proposed allosteric network
    "K600--E743" : ("resid 57 and name NZ*","resid 200 and name OE*"),
    "E743--N596" : ("resid 53 and name ND2","resid 200 and name OE*"),
    "C698--E743" : ("resid 200 and name OE*","resid 155 and name SG"),
    "C698--H757" : ("resid 155 and name HG", 
        "resid 214 and (name ND1 or name NE2)"),
    "K600--E749" : ("resid 206 and name O*", "resid 57 and name NZ"),
    # Binding pocket contacts
    "K764--IP6" : ("resid 221 and name NZ","resname IPL and name O*"),
    "R751--IP6" : ("resid 208 and name N*","resname IPL and name O*"),
    "R571--IP6" : ("resid 28 and name NH*","resname IPL and name O*"),
    "R575--IP6" : ("resid 32 and name NH*","resname IPL and name O*"),
    "Y577--IP6" : ("resid 34 and name OH","resname IPL and name O*"),
    "K792--IP6" : ("resid 249 and name NZ","resname IPL and name O*"),
    "R752--IP6" : ("resid 209 and name NH*","resname IPL and name O*"),
    "K647--IP6" : ("resid 104 and name NZ","resname IPL and name O*"),
    "K600--IP6" : ("resid 57 and name NZ","resname IPL and name O*"),
    "K775--IP6" : ("resid 232 and name NZ","resname IPL and name O*"),
    "Y777--IP6" : ("resid 234 and name OH","resname IPL and name O*"),
    "K645--IP6" : ("resid 102 and name NZ","resname IPL and name O*"),
    # Alpha-flap
    "N763--S767" : ("resid 220 and name O", "resid 224 and name N"),
    "K764--I768" : ("resid 221 and name O", "resid 225 and name N"),
    "E765--I769" : ("resid 222 and name O", "resid 226 and name N"),
    "E766--K770" : ("resid 223 and name O", "resid 227 and name N"),
    "S767--D771" : ("resid 224 and name O", "resid 228 and name N"),
    "I768--I772" : ("resid 225 and name O", "resid 229 and name N"),
    "I769--S773" : ("resid 226 and name O", "resid 230 and name N"),
    "K770--S774" : ("resid 227 and name O", "resid 231 and name N"),
    "D771--K775" : ("resid 228 and name O", "resid 232 and name N"),
    "I772--E776" : ("resid 229 and name O", "resid 233 and name N"),
    "S767--I772" : ("resid 224 and name O", "resid 229 and name N"),
    # Does this network change?
    "E564--K565" : ("resid 21 and name OE*","resid 22 and name NZ*"),
    "K565--E592" : ("resid 21 and name OE*","resid 49 and name OE*"),
    "R571--E592" : ("resid 49 and name OE*","resid 28 and name NH*"),
    "Q741--E776" : ("resid 198 and name NE2","resid 233 and name OE*"),
    "C698--Q741" : ("resid 155 and name O","resid 198 and name NE2"),
    "K764--D756" : ("resid 221 and name NZ*", "resid 213 and name OD*"),
    "R752--I768" : ("resid 209 and name NH*", "resid 225 and name O"),
    "D756--S758" : (" resid 215 and name OG", "resid 213 and name OD*"),
    "R752--E765" : ("resid 209 and name NH*", "resid 222 and name OE*")
    }

    # Just the important selections
selections = {
    # (old) beta-flap network
    "R751--N747" : ("resid 204 and (name OD1 or name ND2)",
        "resid 208 and (name N* or name O)"),
    "N747--E753" : ("resid 204 and name ND2","resid 210 and name OE*"),
    "E753--R745" : ("resid 202 and (name NE or name NH*)",
        "resid 210 and name OE*"),
    "R745--W761" : ("resid 202 and (name NH*)",
        "resid 218 and name CZ*"),
    # Core proposed salt-bridge network
    "K600--E743" : ("resid 57 and name NZ*","resid 200 and name OE*"),
    "E743--N596" : ("resid 53 and name ND2*","resid 200 and name OE*"),
    "C698--E743" : ("resid 200 and name OE*","resid 155 and name SG"),
    "C698--H757" : ("resid 155 and name SG", 
       "resid 214 and (name ND1 or name NE2)"),
    "K600--E749" : ("resid 206 and (name OE* or name O)", 
        "resid 57 and name NZ*"),
    "K600--IP6" : ("resid 57 and name NZ*","resname IPL and name O*"),
    # Other interactions in the salt-bridge network
    "E592--W761" : ("resid 218 and name NE1", "resid 49 and name OE*"),
    "N596--E592" : ("resid 53 and name ND2", "resid 49 and name OE*"),
    "K775--IP6" : ("resid 232 and name NZ*","resname IPL and name O*"),
    "K764--IP6" : ("resid 221 and name NZ*","resname IPL and name O*"),
    "R751--IP6" : ("resid 208 and name NH*","resname IPL and name O*"),
    "R752--IP6" : ("resid 209 and name NH*","resname IPL and name O*"),
    "D771--K775" : ("resid 228 and name O", "resid 232 and name H"),
    "K764--D756" : ("resid 221 and name NZ*", "resid 213 and name OD*"),
    "R752--E765" : ("resid 209 and name NH*", "resid 222 and name OE*"),
    "K775--E692" : ("resid 232 and name NZ*", "resid 149 and name OE*"),
    "K775--N793" : ("resid 232 and name NZ*", "resid 250 and name OD1"),
    "K775--E796" : ("resid 232 and name NZ*", "resid 253 and name OE*"),
    "K775--E743" : ("resid 232 and name NZ*", "resid 200 and name OE*"),
    "K775--IP6" : ("resid 232 and name NZ*", "resname IPL and name O*"),
    "K775--D771" : ("resid 232 and name NZ*", "resid 228 and name OD*"),
    "K775--E776" : ("resid 232 and name NZ*", "resid 233 and name OE*"),
    "K775--N740" : ("resid 232 and name NZ*", "resid 197 and name OD1"),}

selections_k232 = {
    #"K775--E692" : ("resid 232 and name NZ*", "resid 149 and name OE*"),
    # "K775--N793" : ("resid 232 and name NZ*", "resid 250 and name OD1"),
    # "K775--E796" : ("resid 232 and name NZ*", "resid 253 and name OE*"),
    "K775--D771" : ("resid 232 and name NZ*", "resid 228 and name OD*"),
    "K775--N740" : ("resid 232 and name NZ*", "resid 197 and name OD1"),
    #"K775--E743" : ("resid 232 and name NZ*", "resid 200 and name OE*"),
    #"K775--IP6" : ("resid 232 and name NZ*", "resname IPL and name O*"),
    #"K775--IP6_5" : ("resid 232 and name NZ*", "resname IPL and (name O19 or name O20 or name O21)"),
    }

selections_tcda = { #+541
    # Old allosteric network
    "R752--N748" : ("resid 207 and (name OD1 or name ND2)",
        "resid 211 and (name N* or name O)"),
    "N748--E754" : ("resid 207 and name ND2", "resid 213 and name OE*"),
    "E754--R746" : ("resid 205 and name NE","resid 213 and name OE*"),
    "R746--W762" : ("resid 205 and (name NE* or name NH*)",
        "resid 221 and name CZ*"),
    # Proposed allosteric network
    "K601--E744" : ("resid 60 and name NZ", "resid 203 and name OE*"),
    "E744--N597" : ("resid 56 and name ND2","resid 203 and name OE*"),
    "C699--E744" : ("resid 203 and name OE*","resid 158 and name SG"),
    "C699--H758" : ("resid 158 and name SG", 
        "resid 217 and (name ND1 or name NE2)"),
    "K601--E750" : ("resid 209 and name O*", "resid 60 and name NZ"),
    "K601--IP6" : ("resid 60 and name NZ","resname IPL and name O*"),
    # Other contacts
    "N597--E593" : ("resid 52 and name OE*", "resid 56 and name ND2"),
    "E593--W762" : ("resid 221 and name NE1", "resid 52 and name OE*"),
    "K753--E766" : ("resid 212 and name NZ*", "resid 225 and name OE*"),
    "K765--A757" : ("resid 224 and name NZ*", "resid 216 and name O"),
    "D772--K776" : ("resid 231 and name O", "resid 235 and name H"),
    "R752--IP6" : ("resid 211 and name NH*", "resname IPL and name O*"),
    "K753--IP6" : ("resid 212 and name NZ*", "resname IPL and name O*"),
    "K765--IP6" : ("resid 224 and name NZ*", "resname IPL and name O*"),
    "K776--IP6" : ("resid 235 and name NZ*", "resname IPL and name O*"),
    "K776--N741" : ("resid 235 and name NZ*", "resid 200 and name OD1"),
    "K776--E693" : ("resid 235 and name NZ*", "resid 152 and name OE*"),
    }
```
