## Supplementary material for "*Clostridioides difficile* Toxins Unhinged: Allosterically Switchable Network Orients *β*-flap": Scripts, Structures etc.: DSSP.py

```
import sys
import numpy as np
import os
import argparse
import matplotlib.pyplot as plt
import matplotlib.patheffects as pe
import matplotlib.patches as mpatches
import mdtraj as md
import config.settings as c
from tools import utils, traj_funcs

def main(argv):

    try:
        parser = argparse.ArgumentParser()

        parser.add_argument("-p", "--path",
                            action = "store",
                            dest = "path",
                            required = True,
                            help = "Set path to the data directory.")
        parser.add_argument("-s", "--subset",
                            action = "store_true",
                            dest = "subset",
                            default = False,
                            help = ("Make a DSSP plot for just a subset "
                                "of residues."))
        parser.add_argument("-u", "--umbrella",
                            action = "store_true",
                            dest = "umbrella",
                            default = False,
                            help = ("Make plots from umbrella sampling "
                                "data, with a separate DSSP plot for "
                                "each window."))
        parser.add_argument("-f", "--figpath",
                            action = "store",
                            dest = "fig_path",
                            default = False,
                            help = ("Set a path destination for the "
                                "figure."))
        parser.add_argument("-t", "--topol",
                            action = "store",
                            dest = "topol",
                            default = "protein.gro",
                            help = ("File name for topology, inside the "
                                "path directory."))
        parser.add_argument("-b", "--bridge",
                            action = "store_true",
                            dest = "salt_bridge",
                            default = False,
                            help = ("Set a path destination for the "
                                "figure."))                    
        args = parser.parse_args()

    except argparse.ArgumentError:
        print("Command line arguments are ill-defined, please check the "
        "arguments")
        raise

    # Assign group selection from argparse 
    subset = args.subset
    umbrella = args.umbrella
    salt_bridge = args.salt_bridge
    topol = args.topol
    path = f"{ c.data_head }/{ args.path }"
    fig_path = f"{ c.figure_head }/{ args.fig_path }"

    # convert path to a list
    if umbrella:
        # Get a list of paths to all the window directories
        ds = os.listdir(path)
        windows = [path + "/" for d in ds if d.startswith("window")]
    else: 
        data_paths = [path]  

    # iterate over all paths
    for w, p in enumerate(data_paths):

        xtc = "fitted_traj_100.xtc"

        if umbrella:
            
            # Concatenate the runs for a single umbrella to one
            # trajectory
            traj_files = []
            trajs = []
            prev_time = 0
            for run in np.arange(1,5):
                traj_files.append(f"{ p }/run{ run }/{ xtc }") 
            for f in traj_files:
                traj = md.load(f, top=f"{ path }/{ topol }")
                traj.time += prev_time
                prev_time = traj.time[-1]
                trajs.append(traj)
            t = md.join(trajs)

        else: 

            # Otherwise, just one set of traj data to load in
            print("Loading in trajectory...")
            t = md.load(f"{ p }/{ xtc }", #stride=100, 
                        top=f"{ path }/{ topol }")
            print(t)

        # Use a subset of the residues
        if salt_bridge:
            t = t.atom_slice(t.top.select(("resid 49 to 99 or "
                                           "resid 174 to 224")))
        elif subset:
            t = t.atom_slice(t.top.select('resid 179 to 253'))
        else:
            # All residues, excluding the ligand
            t = t.atom_slice(t.top.select('resid 1 to 254'))

        print(t.time)

        # Compute DSSP for the residue selection
        dssp = md.compute_dssp(t, simplified=False)

        plot_dssp(dssp, t, subset, salt_bridge, fig_path, umbrella, w+1)

    return None

def plot_dssp(dssp, t, subset, salt_bridge, path, umbrella, w):
    """Makes an image plot of the secondary structure from DSSP data.

    Parameters
    ----------
    dssp : np.ndarray
        A 2D array of the character codes of the secondary structure of 
        each residue at each time frame; shape=(n_frames, n_residues).
    time : np.ndarray
        The simulation time corresponding to each frame, in picoseconds.
    subset : boolean
        Use a subset of the residues related to the beta and alpha flaps 
        for constructing the DSSP plots?
    salt_bridge: boolean 
        Use a subset of the residues related to the salt bridge K57--E200 
        for constructing the DSSP plots?
    path : str
        Path to the primary working directory.
    umbrella : boolean
        DSSP plot is based on four 25 ns runs in an umbrella window?
    window : int
        Number of the umbrella window.

    Returns
    -------
    None.

    """
    codes = {"H" : 0, "B" : 1, "E" : 2, "G" : 3, "I" : 4, "T" : 5, 
             "S" : 6, " " : 7, "C" : 8}
    codename = {0 : "alpha-helix", 1 : "beta-bridge", 2: "beta-strand",
                3 : r"$3_{10}$-helix", 4 : "pi-helix", 5 : "turn", 
                6 : "bend", 7 : "loops", 8 : "coil"}
    dssp = dssp.T
    print(dssp.shape)
    dssp_colors = np.vectorize(codes.get)(dssp)

    if salt_bridge:
        fig, axs = plt.subplots(2,1, sharex=True, 
                    constrained_layout=True, figsize=(12,6))
        ax1, ax2 = axs
        im1 = ax1.imshow(dssp_colors[:50,:], aspect='auto') 
        im2 = ax2.imshow(dssp_colors[50:,:], aspect='auto') 
        values = np.unique(dssp_colors)
        time = t.time
        resids = [r.resSeq for r in t.topology.residues]
        # Get the colors of the values, according to the colormap used 
        # by imshow
        colors = [im1.cmap(im1.norm(value)) for value in values]
        ax = ax2
    else: 
        fig, ax = plt.subplots(1, 1, constrained_layout=True, 
                               figsize=(12,4))
        im = ax.imshow(dssp_colors, aspect='auto')
        values = np.unique(dssp_colors)
        time = t.time
        resids = [r.resSeq for r in t.topology.residues]
        # Get the colors of the values, according to the colormap used
        # by imshow
        colors = [im.cmap(im.norm(value)) for value in values]

    # Create a patch (proxy artist) for every color
    patches = [mpatches.Patch(color=colors[i], label=codename[i]) \
                for i in range(len(values))]

    if subset:
        left, right = ax.get_xlim()
        ax.hlines([195-180,218-180], left - 100, right + 100, 
            linestyles="dashed", lw=3, colors="#D90000",
            path_effects=[pe.Stroke(linewidth=5, foreground='#E8E9E8'),
            pe.Normal()])
        ax.hlines([219.5-180,231-180], left - 100, right + 100, 
            linestyles="dashed", lw=3, colors="#F93BFF",
            path_effects=[pe.Stroke(linewidth=5, foreground='#E8E9E8'), 
            pe.Normal()])
        ax.set_yticks(np.arange(1, 75, 20))
        ax.set_yticklabels(np.arange(180, 255, 20))
        ax.set_ylabel("Residue Number", labelpad=5, fontsize=24)
        ax.grid(False)
    elif salt_bridge:
        residues = t.topology.residues
        resids1 = [r.resSeq for r in residues if r.resSeq < 150]
        resids2 = [r.resSeq for r in residues if r.resSeq > 150]
        print(resids1,"\n",resids2)
        ax1.set_yticks(np.arange(0, len(resids1), 25))
        ax1.set_yticklabels(resids1[::25])
        ax2.set_yticks(np.arange(0, len(resids2), 25))
        ax2.set_yticklabels(resids2[::25])
        ax1.grid(False)
        ax2.grid(False)
        shared_ylabel = fig.add_subplot(111, frameon=False, 
                                        xticks=[], yticks=[])
        shared_ylabel.yaxis.tick_left()
        shared_ylabel.yaxis.set_label_coords(-0.5, 0)
        shared_ylabel.yaxis.set_label_position("left")
        shared_ylabel.set_ylabel("Residue Number", labelpad=75, 
                                    fontsize=24)
        ax1.tick_params(axis='y', labelsize=20, direction='inout', 
                        width=2, length=10, pad=10)
        ax1.tick_params(axis='x', labelsize=20, direction='inout', 
                        width=2, length=10)
    else:
        ax.set_ylabel("Residue Number", labelpad=5, fontsize=24)
        ax.grid(False)

    # Plot settings
    ax.tick_params(axis='y', labelsize=20, direction='inout', width=2, 
                    length=10, pad=10)
    ax.tick_params(axis='x', labelsize=20, direction='inout', width=2,
                    length=10, pad=10)
    plt.legend(handles=patches, bbox_to_anchor=(1.05, 1), loc=2,
               borderaxespad=0., fontsize=18)

    # X-axis labels
    if time[-1] > 1e6:
        frames = dssp.T.shape[0]
        xticks = np.arange(0,frames+1,frames/10)
        ax.set_xticks(xticks)
        ax.set_xticklabels(list(map(lambda x : str(np.round(x/1e6,0)),
                                    time[::len(time) // 10])))
        ax.set_xlabel("Time (µs)", labelpad=5, fontsize=28)
    elif umbrella:
        xticks = np.arange(0,100e2+1,25e2)
        ax.set_xticks(xticks)
        c = ax.set_xticklabels(list(map(lambda x : str(x/1e2).split(".")[0], 
                            xticks)))
        ax.set_xlabel("Time (ns)", labelpad=5, fontsize=28)
        bottom, top = ax.get_ylim()
        ax.vlines(np.arange(25e2,100e2,25e2), bottom, top, ls="dashed", 
            lw=3, colors="#194D33",
            path_effects=[pe.Stroke(linewidth=5, foreground='#E8E9E8'),
            pe.Normal()])
        plt.title(f"Window { w }", fontsize=24)
    else:
        frames = dssp.T.shape[1]
        xticks = np.arange(0,frames+1,frames/10)
        ax.set_xticks(xticks)
        ax.set_xticklabels(list(map(lambda x : str(x/1e3).split(".")[0],
                            time[::len(time) // 10])))
        ax.set_xlabel("Time (ns)", labelpad=5, fontsize=24)

    if umbrella:
        utils.save_figure(fig, f"{ path }/dssp_{ w }.png")
    elif subset:
        utils.save_figure(fig, f"{ path }/dssp_flaps.png")
    else:
        utils.save_figure(fig, f"{ path }/dssp.png")
    plt.show()
    plt.close()

    return None

if __name__ == '__main__':
    main(sys.argv)
```
