## Supplementary material for "*Clostridioides difficile* Toxins Unhinged: Allosterically Switchable Network Orients *β*-flap": Scripts, Structures etc.: LigandAnalysis.py

```
import sys
import numpy as np
import os
import shutil
import argparse
import matplotlib.pyplot as plt
import matplotlib.patheffects as pe
import MDAnalysis as mda
from MDAnalysis.analysis import rms, align
from MDAnalysis.analysis.distances import distance_array
import config.settings as c
from tools import utils, traj_funcs

def main(argv):

    try:
        parser = argparse.ArgumentParser()

        parser.add_argument("-p", "--path",
                            action = "store",
                            nargs='+',
                            dest = "paths",
                            default = [
                                "unbiased_sims/holo_open/nobackup",
                                ("unbiased_sims/holo_closed/nobackup")],
                            help = """Set path to the data directory.""")
        parser.add_argument("-r", "--recalc",
                            action = "store_true",
                            dest = "recalc",
                            default = False,
                            help = ("Chose whether the trajectory arrays"
                                "should  be recomputed."))
        parser.add_argument("-f", "--figpath",
                            action = "store",
                            dest = "fig_path",
                            default = None,
                            help = ("Set a path destination for the "
                                "figure."))
        args = parser.parse_args()

    except argparse.ArgumentError:
        print("Command line arguments are ill-defined, please check the "
            "arguments")
        raise

    # Assign group selection from argparse
    data_paths = [f"{ c.data_head }/{ p }" for p in args.paths]
    fig_path = f"{ c.figure_head }/{ args.fig_path }"
    recalc = args.recalc

    if not fig_path:
        fig_path = data_paths[0]

    # Determine arrays from traj data or load from file
    datas = get_datas(data_paths, recalc)

    # Make a plot of the center of mass of the ligand
    plot_coms(data_paths, datas, fig_path)

    return None

def get_datas(data_paths, recalc):
    """Gets numpy arrays from traj data or stored array.

    Determines the timeseries of time and ligand COM from trajectory 
    data if not stored in the analysis directory. 

    Parameters
    ----------
    data_paths : (str) list
        List of paths to simulation data.
    recalc : bool
        Redetermine numpy arrays and overwrite existing arrays. 

    Returns
    -------
    datas : dict
        Dictionary using data_paths as keys which stores the numpy 
        arrays.

    """
    datas = {}

    # Indicies of the inflexible residues
    core_res, core = traj_funcs.get_core_res()

    for path in data_paths:

        # Set up path variables
        utils.validate_path(path)
        print(path)
        analysis_path = f"{ os.path.dirname(path) }/analysis"
        utils.create_path(analysis_path)
        datas[path] = {}
        np_files = {"COM_dist" : f"{ analysis_path }/com_dist.npy", 
                    "time_ser" : f"{ analysis_path }/timeseries.npy"}

        # Load in pre-determined numpy files
        if all(list(map(lambda x : os.path.exists(x), \
                        np_files.values()))) and not recalc:

            print(
                "LOADING NUMPY ARRAYS"
            )

            for key, file in np_files.items(): 
                datas[path][key] = np.load(file, allow_pickle=True)

        # Determine arrays from traj data
        else:

            print(
                "EVALUATING WITH MDANALYSIS"
            )

            # Get topol file or edit base topology
            topol = "topol_Pro_Lig.top"
            utils.process_topol(path, topol)

            # Load in universe objects for the simulation and the 
            # reference structures
            u = mda.Universe(f"{ path }/{ topol }", 
                            f"{ path }/fitted_traj_100.xtc",
                            topology_format='ITP')

            # Load in the trajectory and do alignment
            u.transfer_to_memory()
            u = traj_funcs.do_alignment(u)

            # Make a np array of the trajectory time series
            time_ser = []
            for ts in u.trajectory:
                time_ser.append(ts.time)
            time_ser = np.array(time_ser)

            # Save numpy array to file
            datas[path]["time_ser"] = time_ser
            utils.save_array(np_files["time_ser"], time_ser)
            
            # Make a np array of the COM dist wrt the starting position
            ipl = u.select_atoms("resname IPL")
            com_init = ipl.center_of_mass()
            com_dist = np.zeros(u.trajectory.n_frames)
            for ts in u.trajectory:
                com_dist[ts.frame] = distance_array(com_init, 
                                            ipl.center_of_mass())

            # Save numpy array to file
            datas[path]["COM_dist"] = com_dist
            utils.save_array(np_files["COM_dist"], com_dist)

    return datas

def plot_coms(data_paths, datas, fig_path):
    """Makes a time series of the distance from the ligand's initial COM.

    Parameters
    ----------
    data_paths : (str) list
        List of paths to simulation data.
    datas : dict
        Dictionary using data_paths as keys which stores the numpy 
        arrays.
    fig_path : str
        Path for saving the figure. 

    Returns
    -------
    None.

    """
    fig, ax = plt.subplots(constrained_layout=True, figsize=(12,4))
    labels = ["open conf", "closed conf"]
    colors = ["#EAAFCC", "#A1DEA1"]
    stride = 5
    i = 0

    # Overlay plot for each simulation
    for p in data_paths:

        # Access arrays
        arrs = datas[p]
        time_ser = arrs["time_ser"]
        com_dist = arrs["COM_dist"]

        plt.plot(time_ser[::stride], com_dist[::stride], "-", lw=3,
            color=colors[i], label=labels[i], alpha=0.8, 
            path_effects=[pe.Stroke(linewidth=5, foreground='#595959'),
            pe.Normal()])

        # counter for labels and colors
        i += 1

    # Format the axes 
    num_ticks = 10
    xticks = time_ser[::(len(time_ser) // num_ticks)]
    ax.set_xticks(xticks)
    if time_ser[-1] > 1e6:
        # Use microseconds for time labels
        ax.set_xlabel(r'Time ($\mu$s)', fontsize=24, labelpad=5)
        ax.set_xticklabels(list(map(lambda x : str(np.round(x/1e6,1)), xticks)))
    else:
        # Use nanoseconds for time labels
        ax.set_xlabel(r'Time (ns)', fontsize=24, labelpad=5)
        ax.set_xticklabels(list(map(lambda x : str(np.round(x/1e3,1)), xticks)))
    ax.set_ylabel(r"COM Distance ($\AA$)", labelpad=5, fontsize=24)
    plt.legend(fontsize=20)

    # Save plot to file
    utils.save_figure(fig, f"{ fig_path }/com_dist.png")
    plt.show()
    plt.close()

    return None

if __name__ == '__main__':
    main(sys.argv)
```
