## Supplementary material for "*Clostridioides difficile* Toxins Unhinged: Allosterically Switchable Network Orients *β*-flap": Scripts, Structures etc.: MutInf.py

```
import sys
import numpy as np
import os
import argparse
import config.settings as c
from tools import utils, traj_funcs
import matplotlib.pyplot as plt
import matplotlib.patheffects as pe
import MDAnalysis as mda
from MDAnalysis.analysis import rms, align, dihedrals
from MDAnalysis.analysis.distances import distance_array
import pandas as pd
from sklearn import metrics
import networkx as nx
from sklearn.cluster import SpectralClustering
from Bio import PDB
import time
from joblib import Parallel, delayed
import multiprocessing

def main(argv):

    try:
        parser = argparse.ArgumentParser()

        parser.add_argument("-p", "--path",
                            action = "store",
                            dest = "path",
                            default = "unbiased_sims/apo_open/nobackup",
                            help = """Set path to the data directory.""")
        parser.add_argument("-r", "--recalc",
                            action = "store_true",
                            dest = "recalc",
                            default = False,
                            help = """Chose whether the trajectory arrays
                                should be recomputed.""")
        parser.add_argument("-w", "--windows",
                            action = "store_true",
                            dest = "windows",
                            default = False,
                            help = ("Calculate NMI matricies for windows"
                                "from umbrella sampling"))
        parser.add_argument("-f", "--figpath",
                            action = "store",
                            dest = "fig_path",
                            default = "",
                            help = "Set a path destination for the "
                                "figure.")
        parser.add_argument("-t", "--topol",
                            action = "store",
                            dest = "topol",
                            default = "topol_protein.top",
                            help = """File name for topology, inside the 
                                path directory.""")   
        parser.add_argument("-x", "--xtc",
                            action = "store",
                            dest = "xtc",
                            default = "fitted_traj_100.xtc",
                            help = """File name for trajectory, inside 
                                the path directory.""")                             
        args = parser.parse_args()

    except argparse.ArgumentError:
        print("""Command line arguments are ill-defined, please check the
              arguments""")
        raise

    # Assign group selection from argparse 
    data_path = f"{ c.data_head }/{ args.path }"
    fig_path = f"{ c.figure_head }/{ args.fig_path }"
    recalc = args.recalc
    topol = args.topol
    xtc = args.xtc
    windows = args.windows

    global analysis_path

    # Check for valid paths
    for p in [data_path, fig_path]:
        utils.validate_path(p)
    
    start_time = time.time()

    # Store calculated outputs as dataframes etc. 
    analysis_path = f"{ os.path.dirname(data_path) }/analysis"
    cluster_path = f"{ os.path.dirname(data_path) }/analysis/cluster"
    if windows:
        analysis_path = f"{ data_path }/analysis"
        cluster_path = f"{ analysis_path }/cluster"
    if not os.path.exists(analysis_path):
        os.makedirs(analysis_path)
    if not os.path.exists(cluster_path):
        os.makedirs(cluster_path)

    # Extract torsion data from the simulation or load from file
    df_tor, u = get_torsions(data_path, topol, xtc, recalc=recalc)

    # Determine the normalized mutual information for all torsion pairs
    df_nmi = get_nmis(data_path, df_tor, recalc=recalc)

    end_time = time.time()
    print(f"\nThat took { end_time - start_time } seconds.\n")

    if not (df_nmi.values == df_nmi.values.T).all().all():
        print("ERROR: Non-symmetric NMI matrix.")
    else:
        print("The NMI matrix is symmetric.")

    # Apply corrections to the NMI
    df_corr = apply_nmi_corrections(df_tor, df_nmi, data_path)

    # Check how sparse the matrix is
    empty = df_corr.isna().sum().sum()
    elements = df_corr.shape[0] ** 2
    print((f"\nTOTAL: { elements }, EMPTY: { empty },\n" 
          f"RATIO: { np.round(empty / elements *100, 1) } %\n"))

    # Analyze the NMI matrix by eigendecomposition
    analyze_eigs(df_nmi, fig_path, descript="_full_torsion")
    plot_mi_hist(df_nmi, fig_path, descript="_full_torsion")
    analyze_eigs(df_corr, fig_path, descript="_full_torsion_corr")
    plot_mi_hist(df_corr, fig_path, descript="_full_torsion_corr")

    # Cluster using spectral clustering
    clusters = get_clusters(df_nmi, 3, cluster_path, "_full_torsion")
    
    # Make a plot of the torsion NMI matrix
    plot_nmi(df_nmi, f"{ fig_path }/all_torsions_nmi.png")

    # Determine the NMI between residue pairs, using a 
    # summation over the torsions
    res_nmi = get_res_nmi(data_path, df_nmi, u, recalc=recalc, 
                          corrected=False)
    res_nmi_corr  = get_res_nmi(data_path, df_corr, u, recalc=False, 
                                corrected=True)
    plot_nmi(res_nmi, f"{ fig_path }/residues_nmi.png")
    plot_nmi(res_nmi_corr, f"{ fig_path }/residues_nmi_corr.png")

    # Analyze the residue NMI matrix by eigendecomposition
    analyze_eigs(res_nmi, fig_path, descript="_res")
    plot_mi_hist(res_nmi, fig_path, descript="_res")
    analyze_eigs(res_nmi_corr, fig_path, descript="_res_corr")
    plot_mi_hist(res_nmi_corr, fig_path, descript="_res_corr")

    # Use spectral clustering to put residues into clusters based on 
    # residue nmi values
    clusters = get_clusters(res_nmi, 3, cluster_path, "_res_nmi_corr")
    clusters_corr = get_clusters(res_nmi_corr, 3, cluster_path, 
                           "_res_nmi_corr")
    print(clusters)
    visualize_clusters(clusters, cluster_path, "open_ref", 
                        f"{ c.struct_head }/open_ref_state.pdb",
                        corrected=False)
    visualize_clusters(clusters, cluster_path, "closed_ref", 
                        f"{ c.struct_head }/closed_ref_state.pdb",
                        corrected=False)
    visualize_clusters(clusters_corr, cluster_path, "open_ref", 
                        f"{ c.struct_head }/open_ref_state.pdb",
                        corrected=True)
    visualize_clusters(clusters_corr, cluster_path, "closed_ref", 
                        f"{ c.struct_head }/closed_ref_state.pdb",
                        corrected=True)

    sys.exit(1)

    if not windows:

        # Makes a (boolean) matrix for the residue contacts
        contacts = identify_contacts(data_path, topol, xtc, 
                                     res_nmi, recalc=False)
        plot_nmi(contacts, f"{ fig_path }/residue_contacts.png")

        # Construct a network based on connected residues
        res_graph = make_graph(res_nmi, contacts, data_path)
        res_graph = make_graph(res_nmi_corr, contacts, data_path)

    return None

def visualize_clusters(clusters, cluster_path, ref_name, ref_path, 
        corrected=False):
    """Visualize clusters by adding labels to pdb.

    Parameters
    ----------
    clusters : pd.Series
        Cluster labels for each residue.
    cluster_path : str
        Path to the cluster analysis directory. 
    ref_name : str
        Name of the structure, used in the output file.

    Returns
    -------
    None.

    """
    # Load in reference structure
    pdb_parser = PDB.PDBParser(QUIET=True)
    ref_struct = pdb_parser.get_structure(ref_name, ref_path)

    # Make groups accessible by the residue id.
    resid_clusts = {}
    for key, group in clusters.items():
        resid = int(key)
        resid_clusts[resid] = group

    # Number of clusters
    n_clusts = len(set(clusters.values()))

    # Set the beta factor values to the cluster group
    for model in ref_struct:
        for chain in model:
            for residue in chain:
                for atom in residue:
                    atom.set_bfactor(0.0)
                r = residue.id[1]
                residue["CA"].set_bfactor(resid_clusts[r])

    # Save the modified structures for visualization
    io = PDB.PDBIO()
    io.set_structure(ref_struct)
    if corrected:
        io.save(f"{ cluster_path }/{ ref_struct.id }_{ n_clusts }clusters_corr.pdb")
    else:
        io.save(f"{ cluster_path }/{ ref_struct.id }_{ n_clusts }clusters.pdb")

    return None

def get_clusters(df, n_clust, path, descript=""):
    """Determine clusters of the NMI matrix with spectral clustering.
    Parameters
    ----------
    df : pd.DataFrame
        A symmetric matrix containing the normalized mutual information 
        for all torsion/residue pairs.
    n_clust: int
        Number of clusters to split data into. 
    path : str
        Path for saving pandas series of classification markers.
    descript : str
        A short descriptor for naming csv file. 

    Returns
    -------
    clusters : pd.Series
        Cluster labels for each node. 

    """
    # DataFrame stored as a csv file
    df_file = f"{ path }/{ n_clust }clust{ descript }.csv"

    # Use scikit-learn implemetation of spectral clustering
    # affinity="precomputed" means the adjacency matrix is precomputed
    clustering = SpectralClustering(
                    n_clusters=n_clust, assign_labels="discretize",
                    affinity="precomputed"
                 ).fit(df.fillna(0))

    # Make a dictionary for the clusters with torsions labeled
    clusters = {}
    for tor_lab, clust in zip(df.columns.to_list(), clustering.labels_): 
        clusters[tor_lab] = clust

    # Convert to pd.Series and save to csv
    series = pd.Series(clusters)
    utils.save_df(series, df_file)

    return clusters

def analyze_eigs(df, fig_path, descript=""):
    """Makes basic plots to understand NMI matrix eigendecomposition. 

    Parameters
    ----------
    df : pd.DataFrame
        A square DataFrame or any matrix object understandable by numpy,
        which should undergo an eigendecomposition. 
    fig_path : str
        Directory for storing the eigendecomposition figures. 
    descript : str
        A short descriptor for naming plots. 

    Returns
    -------
    eigvals : np.ndarray
        Array of ordered eigenvalues.
    eigvecs : np.ndarray
        2D array of orrdered eigenvectors.

    """
    def plot_eigvals(eigvals, n, fig_path):
        """Makes a scree plot using the first n eigenvals.

        """
        fig, ax = plt.subplots()
        ax.scatter(np.arange(1, n+1), eigvals[:n], s=20)
        utils.save_figure(fig, f"{ fig_path }/eigenvalues{ descript }.png")
        plt.close()

    eigvals, eigvecs = np.linalg.eig(df.fillna(0))
    inds = eigvals.argsort()[::-1]
    eigvals = eigvals[inds]
    eigvecs = eigvecs[:,inds]

    # Scree plot for the eigenvalues
    plot_eigvals(eigvals, len(eigvals), fig_path)

    # 2D scatter plots for major eigenvectors
    fig, axes = plt.subplots(2,2)
    axes = axes.flatten()
    count = 0 
    for i, j in [(0,1),(0,2),(1,2)]:

        ax = axes[count]
        ax.scatter(eigvecs[:,i], eigvecs[:,j], 
                    c=np.arange(1,len(eigvals)+1), marker="o")
        ax.set_xlabel(f"{i + 1}")
        ax.set_ylabel(f"{j + 1}")
        count += 1

    utils.save_figure(fig, f"{ fig_path }/2d_eigenvecs{ descript }.png")
    plt.close()

    # Make 3D plot of first 3 eigenvectors
    fig = plt.figure()
    ax = fig.add_subplot(projection='3d')

    c = ax.scatter(eigvecs[:,0], eigvecs[:,1], eigvecs[:,2])
    ax.set_xlabel('Eigenvector 1', labelpad=25)
    ax.set_ylabel('Eigenvector 2', labelpad=25)
    ax.set_zlabel('Eigenvector 3', labelpad=25)
    plt.close()

    return eigvals, eigvecs

def get_torsions(path, topol, xtc, recalc=False):
    """Gets trajectories of phi, psi and chi torsions in a table.
    
    Uses a MultiIndex'ed pandas DataFrame for storing the torsion 
    trajectories so individual residues or types of torsions can be 
    conveniently accessed.

    Parameters
    ----------
    path : str
        Path to the directory with trajectory data.
    topol : str
        Name of the topol file, excluding solvent and counterions. 
    xtc : str
        Name of the trajectory file for extracting dihedral data.
    recalc : bool
        Redetermine the table from simulation data, even if a DataFrame 
        is already saved to file.

    Returns
    -------
    df_tor : pd.DataFrame
        A DataFrame of the torsions with row indexing by trajectory 
        frame. A MultiIndex is used for the columns, with residue numbers
        at the highest level and torsion labels at the secondary level, 
        (phi, psi, chi1, ..., chin).
    u : mda.Universe
        The relevant universe object.
    
    """
    # DataFrame stored as a csv file
    df_file = f"{ analysis_path }/torsions_df.csv"
    
    # Load in universe objects for the simulation and the reference 
    # structures
    u = mda.Universe(f"{ path }/{ topol }", 
                     f"{ path }/{ xtc }", 
                     topology_format='ITP')    

    # Load in the hierarchical DataFrame if it exists
    if os.path.exists(df_file) and not recalc:

        print(
            "LOADING TORSIONS DataFrame FROM CSV..."
        )  

        # Uses .hdf for heirarchical indexing
        df_tor = pd.read_csv(df_file, header=0, index_col=0)

    else:

        print(
            "EVALUATING TORSIONS WITH MDANALYSIS..."
        )   

        # Use standard alignment procedure 
        # u = traj_funcs.do_alignment(u)

        # Initialize DataFrame for all torsions
        df_tor = pd.DataFrame()

        # store all of the torsion AtomGroups
        groups = []
        labels = []

        # Iterate over protein residues (exclude IPL)
        for res in u.residues[:254]:

            # Apply binning to each series
            bin_edges = np.arange(-180,181)

            # Convenience variables
            res_id = res.resid 
            resn = res.resname

            # Get AtomGroups of the residues torsions
            group = [res.phi_selection(), res.psi_selection(), 
                      res.chi1_selection()]
            group.extend(get_chi_groups(res))

            # Assign Phi and Psi and handle terminal residues
            labs = [(f"{ resn }-{ res_id }-Phi"),
                   (f"{ resn }-{ res_id }-Psi"),]

            # Assign any/all chi torsions for the residues
            for c, g in enumerate(group[2:]):
                ind = (f"{ resn }-{ res_id }-Chi-{ c + 1 }")
                labs.append(ind)

            # Filter our non-groups
            group_filt = [g for g in group if g is not None ]
            labs_filt = [l for l, g in zip(labs, group) if g is not None]
            groups.extend(group_filt)
            labels.extend(labs_filt)

        # Determine torsions for all the AtomGroups
        tors = dihedrals.Dihedral(groups).run()
        t = tors.results.angles

        # Convert torsions to dictionary object
        tor_dict = {}
        for lab, tor in zip(labels, zip(*t)):
            tor_discrete = np.digitize(tor, bin_edges, right=False)
            tor_dict[lab] = tor_discrete 

        # Convert the residues' torsions dictionary and concatenate 
        # with the DataFrame to combine with other residues
        df_tor = pd.DataFrame(tor_dict)

        # Format column names and save DataFrame to file
        utils.save_df(df_tor, df_file)

    return df_tor, u

def calc_MI(df_tor, tor1, tor2, norm_type="NMI"):
    """Determines the normalized mutual information between two torsions.

    See Scikit-learn documentation for the mathematical description: 
    https://scikit-learn.org/0.18/modules/clustering.html#mutual-info-score
    NMI https://scikit-learn.org/0.18/modules/generated/sklearn.metrics.normalized_mutual_info_score.html#sklearn.metrics.normalized_mutual_info_score
    Adjusted NMI https://scikit-learn.org/0.18/modules/generated/sklearn.metrics.adjusted_mutual_info_score.html#sklearn.metrics.adjusted_mutual_info_score

    Parameters
    ----------
    tor1 : pd.Series 
        Trajectory data in bins for torsion 1.
    tor2 : pd.Series 
        Trajectory data in bins for torsion 2.
    norm_type : str
        Use a selection key (i.e. 'NMI', 'Adjusted') for the normalized 
        scoring function.

    Returns
    -------
    nmi : float
        The normalized mutual information of the two torsions.

    """
    # Select algortithm for normalization
    if norm_type == "NMI":
        nmi = metrics.normalized_mutual_info_score(df_tor[tor1], 
                                                 df_tor[tor2])
    elif norm_type == "Adjusted":
        nmi = metrics.adjusted_mutual_info_score(df_tor[tor1], 
                                                 df_tor[tor2])
    else:
        print(f"Invalid 'norm_type' used : { norm_type }. "
                "Select a valid function for normalization (i.e. 'NMI', "
                "'Adjust'.)")
        sys.exit(1)

    return nmi, (tor1, tor2)

def get_nmis(path, df_tor, recalc=False):
    """Makes a matrix of the NMI for all torsion pairs.

    See Scikit-learn documentation for the mathematical description: 
    https://scikit-learn.org/0.18/modules/clustering.html#mutual-info-score
    NMI https://scikit-learn.org/0.18/modules/generated/sklearn.metrics.normalized_mutual_info_score.html#sklearn.metrics.normalized_mutual_info_score
    Adjusted NMI https://scikit-learn.org/0.18/modules/generated/sklearn.metrics.adjusted_mutual_info_score.html#sklearn.metrics.adjusted_mutual_info_score

    Parameters
    ----------
    path : str
        Path to the directory with trajectory data.
    df_tor : pd.DataFrame
        A DataFrame of the torsions with row indexing by trajectory 
        frame. A MultiIndex is used for the columns, with residue numbers 
        at the highest level and torsion labels at the secondary level, 
        (phi, psi, chi1, ..., chin).

    Returns
    -------
    df_nmi : pd.DataFrame
        A symmetric matrix containing the normalized mutual information 
        for all torsion pairs.

    """
    # DataFrame stored as a csv file
    df_file = f"{ analysis_path }/nmis_df.csv"

    if os.path.exists(df_file) and not recalc:

        print(
            "LOADING NMI DataFrame from CSV..."
        )  

        df_nmis = pd.read_csv(df_file, header=0, index_col=0)

    else:

        print(
            "EVALUATING NMIs with SCIKIT-LEARN..."
        )   

        # Determine the number of CPU cores
        total_cores = multiprocessing.cpu_count()   
        cores_to_use = int(0.8 * total_cores)

        # Make a (symmetric) table with all NMI for residue pairs
        df_nmis = pd.DataFrame(index=df_tor.columns, columns=df_tor.columns)

        # Calculate MI for each pair of torsion columns in parallel
        results = Parallel(n_jobs=cores_to_use)(delayed(calc_MI)(df_tor, col_i, col_j)
                                    for i, col_i in enumerate(df_tor.columns)
                                    for j, col_j in enumerate(df_tor.columns)
                                    if i <= j)

        # Update df_nmis with the calculated MI values
        for nmi, (col_i, col_j) in results:
            df_nmis.loc[col_i, col_j] = nmi
            df_nmis.loc[col_j, col_i] = nmi

        utils.save_df(df_nmis, df_file)

    return df_nmis

def apply_nmi_corrections(df_tor, df_nmi, path):
    """Currently applies no actual changes to the matrix.

    Parameters
    ----------
    df_tor : pd.DataFrame
        A DataFrame of the torsions with row indexing by trajectory frame. A MultiIndex 
        is used for the columns, with residue numbers at the highest level and torsion 
        labels at the secondary level, (phi, psi, chi1, ..., chin).
    df_nmi : pd.DataFrame
        A symmetric matrix containing the normalized mutual information for
        all torsion pairs.

    Returns
    -------
    df_corr : pd.DataFrame
        A symmetric matrix containing the normalized mutual information for
        all torsion pairs with statistical corrections applied.
    
    """
    # for i in df_nmi.columns.to_list():
    #     # min_nmi = calc_MI(df_tor[i], df_tor[i].sample(frac=1).reset_index(drop=True))
    #     min_nmi = calc_MI(df_tor[i], df_tor[i][::-1])
    #     nmi_threshold[i] = min_nmi

    # tor_features = {}
    # for i in df_nmi.columns.to_list():
    #     unique = len(np.unique(df_tor[i]))
    #     tor_features[i] = (nmi_threshold[i], unique)
    # with open(f"{ analysis_path }/tor_features.pickle", 'wb') as file:
    #     pickle.dump(tor_features, file)

    # print(nmi_threshold)
    # print(min(nmi_threshold.values()), max(nmi_threshold.values()))

    df_corr = df_nmi.mask(df_nmi < 0.1, np.nan)

    # if not (df_corr.values == df_corr.values.T).all().all():
    #     print("ERROR: Non-symmetric NMI matrix.")
    #     sys.exit(1)

    return df_corr

def get_res_nmi(path, df, u, recalc=False, corrected=False):
    """Determines the residue NMI from torsional NMIs.

    Parameters
    ----------
    path : str
        Path to the directory with trajectory data.
    df : pd.DataFrame
        A symmetric matrix containing the normalized mutual information 
        for all torsion pairs with or without statistical corrections 
        applied.
    u : mda.Universe
        The relevant universe object. 
    recalc : bool
        Redetermine the table from simulation data, even if a DataFrame 
        is already saved to file.
    corrected : bool
        DataFrame should use raw or corrected NMI values.

    Returns
    -------
    res_nmi : pd.DataFrame
        A smaller matrix of the NMI between entire residues. Consists of 
        the sum of torsional NMIs between the residues. 

    """
    # DataFrame stored as a csv file
    if corrected:
        df_file = f"{ analysis_path }/res_nmi.csv"
    else:
        df_file = f"{ analysis_path }/res_nmi_corr.csv"

    if os.path.exists(df_file) and not recalc:

        print(
            "LOADING RESIDUE NMI DataFrame from CSV..."
        )  

        res_nmi = pd.read_csv(df_file, index_col=0, header=0)

    else:

        print(
            "EVALUATING RESIDUE NMIs using TORSION PAIR NMIs..."
        )   

        n = 254
        res_nmi = pd.DataFrame(index=range(1, n + 1), 
                               columns=range(1, n + 1))

        for i in range(n):
        
            df_resi = df.filter(regex=f"-{ i + 1 }-")

            for j in range(254):

                df_pair = df_resi.loc[df.index.str.contains(f"-{ j + 1 }-")]

                count = df_pair.count().sum()
                nmi_sum = df_pair.values.sum()

                res_nmi.loc[i+1, j+1] = nmi_sum / count
                res_nmi.loc[j+1, i+1] = nmi_sum / count

        utils.save_df(res_nmi, df_file)

    return res_nmi

def identify_contacts(path, topol, xtc, res_nmi, recalc=False):
    """Identifies which residues are considered as contacts.

    Uses a contact threshhold of 5.5 AA for the heavy atoms for at
    least 75% of the simulation data. 
    Based on https://doi.org/10.1016/bs.mie.2016.05.027 .

    Parameters
    ----------
    path : str
        Path to the directory with trajectory data.
    topol : str
        Name of the topol file, excluding solvent and counterions. 
    xtc : str
        Name of the trajectory file for extracting dihedral data.
    res_nmi : pd.DataFrame
        A matrix of the NMI between entire residues. Consists of the 
        sum of torsional NMIs between the residues.
    recalc : bool
        Redetermine the table from simulation data, even if a DataFrame 
        is already saved to file.

    Returns
    -------
    df_contacts : pd.DataFrame
        Boolean values are used to construct contact matrix.

    """
    # DataFrame stored as a csv file
    df_file = f"{ analysis_path}/res_contacts.csv"

    if os.path.exists(df_file) and not recalc:

        print(
            "LOADING RESIDUE CONTACT DataFrame FROM CSV..."
        )  

        df_contacts = pd.read_csv(df_file, index_col=0, header=0)

    else:

        print(
            "EVALUATING RESIDUE-RESIDUE CONTACT PAIRS as BOOLEAN "
            "with MDANALYSIS..."
        )   

        n = res_nmi.columns
        df_contacts = pd.DataFrame(index=n, columns=n)

        core_res, core = traj_funcs.get_core_res() 
        # Not all data is needed to determine if residues are in contact
        stride = 100 

        # Load in universe objects for the simulation and the reference structures
        u = mda.Universe(f"{ path }/{ topol }", f"{ path }/{ xtc }", 
                         topology_format='ITP')    
        total_frames = int(len(u.trajectory) / stride) 
        print("\tTOTAL FRAMES ", total_frames)

        align.AlignTraj(u, u.select_atoms("protein"), select=core, 
                        in_memory=True).run()

         # Iterate over residues
        for resi in u.residues:

            resi_id = resi.resid 
            resin = resi.resname
            name_i = f"{ resin } { resi_id }"

            resi_heavy = u.select_atoms(f"resid { resi_id } and not name H*")

            contact_counts = {}

            for ts in u.trajectory[::stride]:

                # Calculate distances between the heavy atoms of the 
                # target residue and all atoms
                dists = distance_array(resi_heavy.positions, 
                                        u.atoms.positions)

                # Identify atoms in contact based on the distance 
                # threshold 5.5 AA
                # The method .any() qualifies the atom if it contacts any 
                # heavy atom in the target
                in_contact = (dists < 5.5).any(axis=0)

                # Update contact counts for each residue
                for resj in u.residues:

                    resj_id = resj.resid
                    resjn = resj.resname
                    name_j = f"{ resjn } { resj_id }"

                    # Add to count if the residues qualifies
                    if name_j not in contact_counts:
                        contact_counts[name_j] = 0
                    if in_contact[resj.atoms.indices].sum() > 0: 
                        contact_counts[name_j] += 1

            # Identify residues in contact for at least 75% of the frames
            contact_percentage = 0.75
            contact_residues = [res for res, count in contact_counts.items() 
                                if count >= total_frames * contact_percentage]

            print(f"CONTACTS FOR { name_i } : ", contact_residues)

            for resj in u.residues:

                resj_id = resj.resid
                resjn = resj.resname
                name_j = f"{ resjn } { resj_id }"

                df_contacts.loc[name_i, name_j] = (name_j in contact_residues)
                df_contacts.loc[name_j, name_i] = (name_j in contact_residues)

        utils.save_df(df_contacts, df_file)
        print(df_contacts)

    return df_contacts

def make_graph(res_nmi, contacts, path, corr=False):
    """Makes a network weighted by NMI.

    Each residue forms a node, while the contacts determine the graph edges.
    The weight for each edge is the NMI of the connected residues and 
    the network object is saved as a ".gexf". 

    Parameters
    ----------
    res_nmi : pd.DataFrame
        A smaller matrix of the NMI between entire residues. Consists of the 
        sum of torsional NMIs between the residues.
    contacts : pd.DataFrame
        Boolean values are used to construct contact matrix.
    path : str
        Path for storing the network object as a ".gexf" which can be 
        visualized with Gephi.

    Returns
    """
    g = nx.Graph()

    # Add all residues as nodes in the graph
    all_res = res_nmi.columns.to_list()
    g.add_nodes_from(all_res)

    count = 0
    for i, res1 in enumerate(all_res):
        for j, res2 in enumerate(all_res):
            if (i < j) & (contacts.loc[res1, res2]):
                count +=1
                g.add_edge(res1, res2, weight=-np.log(res_nmi.loc[res1,res2]))

    print(type(g))

    if corr:
        nx.write_gexf(g, f"{ analysis_path }/connected_residues_corr.gexf")
    else:
        nx.write_gexf(g, f"{ analysis_path }/connected_residues.gexf")

    return g

def get_chi_groups(res):
    """Get the groups involved in all the chi2+ dihedral groups. 

    See a list of the chi dihedrals at 
    http://www.mlb.co.jp/linux/science/garlic/doc/commands/dihedrals.html.
    The dihedrals beyond chi1 are considered here.

    Parameters
    ----------
    res : MDAnalysis.core.groups.Residue
        Residue object for selecting relevant AtomGroup.

    Returns
    -------
    chi_groups : ((AtomGroup) list) list
        A list of list of AtomGroups, where each entry correspond to the 
        chi dihedrals beyond chi1.

    """
    if res.resname in ["ALA", "GLY", "VAL", "CYS", "SER", "THR"]:
        return []  # No chi dihedral greater than 1 

    # Identify chi atoms based on the residue type
    if res.resname == "ARG":
        chi_atoms = ["CA", "CB", "CG", "CD", "NE", "CZ", "NH1"]
    elif res.resname == "LYS":
        chi_atoms = ["CA", "CB", "CG", "CD", "CE", "NZ"]
    elif res.resname == "LYN":
        chi_atoms = ["CA", "CB", "CG", "CD", "CE", "NZ"]
    elif res.resname == "MET":
        chi_atoms = ["CA", "CB", "CG", "SD", "CE"]
    elif res.resname == "GLN":
        chi_atoms = ["CA", "CB", "CG", "CD", "OE1"]
    elif res.resname == "GLU":
        chi_atoms = ["CA", "CB", "CG", "CD", "OE1"]
    elif res.resname == "ASN":
        chi_atoms = ["CA", "CB", "CG", "OD1"]
    elif res.resname == "ASP":
        chi_atoms = ["CA", "CB", "CG", "OD1"]
    elif res.resname == "HIS": 
        chi_atoms = ["CA", "CB", "CG", "ND1"]
    elif res.resname == "ILE":
        chi_atoms = ["CA", "CB", "CG1", "CD"]
    elif res.resname == "LEU":
        chi_atoms = ["CA", "CB", "CG", "CD1"]
    elif res.resname == "PRO":
        chi_atoms = ["CA", "CB", "CG", "CD"]
    elif res.resname == "PHE":
        chi_atoms = ["CA", "CB", "CG", "CD1"]
    elif res.resname == "TRP":
        chi_atoms = ["CA", "CB", "CG", "CD1"]
    elif res.resname == "TYR":
        chi_atoms = ["CA", "CB", "CG", "CD1"]

    chi_atom_lists = []
    for i in range(len(chi_atoms) - 3):
        chi_atom_lists.append(chi_atoms[i:i+4])

    chi_groups = [] 
    for c in chi_atom_lists:
        chi_groups.append([a for a in res.atoms if a.name in c])

    return chi_groups

def plot_nmi(df, fig_path):
    """Makes a plot depicting the NMI matrix.

    Parameters
    ----------
    df : pd.DataFrame
        A symmetric matrix containing the normalized mutual information for
        all torsion pairs.    
    fig_path : str
        Path to the figure image file. 
    
    Returns
    -------
    None.

    """
    fig, ax = plt.subplots(constrained_layout=True, figsize=(12,12))
    cax = ax.matshow(df.astype(np.float32), cmap='YlGnBu')
    cbar = plt.colorbar(cax, shrink=0.75)

    if len(df.columns) > 500:
        cols = list(df.columns)
        label_pos = [i for i, label in enumerate(cols) if i % 100 == 0]
        label_names = [label.split("-")[1] for i, label in enumerate(cols) if i % 100 == 0]
    else:
        cols = list(df.columns)
        label_pos = [i for i, label in enumerate(cols) if i % 20 == 0]
        label_names = [label for i, label in enumerate(cols) if i % 20 == 0]

    plt.xticks(label_pos, label_names, rotation=45)
    plt.yticks(label_pos, label_names)
    ax.set_xlabel("Residue ID")
    ax.set_ylabel("Residue ID")

    cbar.ax.tick_params(labelsize=20, direction='out', width=2, length=5)
    cbar.outline.set_linewidth(2)
    ax.grid(False)

    utils.save_figure(fig, fig_path)
    plt.close()

    return None

def plot_mi_hist(df, fig_path, descript=""):
    """Plots a histogram of all torsion pair NMIs. 
    
    Parameters
    ----------
    df : pd.DataFrame
        A symmetric matrix containing the normalized mutual information
        for all torsion pairs.    
    fig_path : str
        Path to the directory for saving the figure image.
    descript : str
        A short descriptor for naming plots. 
    
    Returns
    -------
    None.

    """
    fig, ax = plt.subplots()
    ax.hist(df.values.flatten(), bins=50, color='#cc4bb0', edgecolor='black',
            density=True)
    ax.grid(axis='y', linestyle='--', alpha=0.7)
    ax.set_xlabel("NMI")
    ax.set_ylabel("frequency")
    ax.set_xlim(0,1)

    utils.save_figure(fig, f"{ fig_path }/nmi_histogram{ descript }.png")
    plt.close()

    return None

if __name__ == '__main__':
    main(sys.argv)
```
