## Supplementary material for "*Clostridioides difficile* Toxins Unhinged: Allosterically Switchable Network Orients *β*-flap": Scripts, Structures etc.: AnimateApoFes1d.py

```
import numpy as np
import matplotlib.pyplot as plt
from mpl_toolkits.mplot3d import Axes3D
from matplotlib.animation import FuncAnimation
import mplcursors
import sys
sys.path.insert(0, "/home/lf1071fu/project_b3/ProjectB3")
from tools import utils, traj_funcs
import config.settings as cf

# Load in pathway data
ani_path = f"{ cf.figure_head }/animations/strs_WT_apo_path"
pathway = np.load(f"{ ani_path }/cv_lowE_path.npy")
xvals = np.arange(0,len(pathway[:,0]))

# Make plot
fig, ax = plt.subplots(constrained_layout=True, figsize=(8,8))
ax.plot(xvals, pathway[:,2], lw=2, color="#8f8f8f", label="apo WT path")
track_path = ax.scatter([], [], s=150, color="#E65100")

# Set aesthetics
ax.set_ylabel(r"$\Delta G$ (kJ / mol)", labelpad=3, fontsize=16)
ax.grid(False)
ax.set_ylim([-9,50])
ax.set_xticks([])
plt.yticks(fontsize=12)
ax.legend(fontsize=12)

# Number of frames in the animation
rotation_frames = pathway.shape[0]
pause_frames = 20 
total_frames = rotation_frames + 2 * pause_frames

# Update function for the animation
def update(num):
    if num < pause_frames:
        # Pause at the initial position
        current_depth = pathway[0,2]
        p = 0
    elif num < pause_frames + rotation_frames:
        # Rotate the vector
        current_depth = pathway[num - pause_frames, 2]
        p = num - pause_frames
    else:
        # Pause at the final position
        current_depth = pathway[-1,2]
        p = len(pathway[:,0]) -1

    track_path.set_offsets([(p, current_depth),])
    
    return track_path,


# Create the animation
ani = FuncAnimation(fig, update, frames=total_frames, interval=50, 
            blit=True)

# Save and view
ani.save(f"{ cf.figure_head }/animations/apo_wt_fes_path.mp4", 
            writer="ffmpeg")
plt.show()
plt.close()
```
