## Supplementary material for "*Clostridioides difficile* Toxins Unhinged: Allosterically Switchable Network Orients *β*-flap": Scripts, Structures etc.: AnimateBetaVec.py

```
import numpy as np
import matplotlib.pyplot as plt
from mpl_toolkits.mplot3d import Axes3D
from matplotlib.animation import FuncAnimation
import mplcursors
import sys
sys.path.insert(0, "/home/lf1071fu/project_b3/ProjectB3")
from tools import utils, traj_funcs
import config.settings as cf

def interpolate_vectors(vec_init, vec_final, num_steps):
    interpolated_vectors = np.zeros((num_steps,3))
    for i in range(num_steps):
        t = i / (num_steps - 1)  # Compute the interpolation parameter
        vector = (1 - t) * vec_init + t * vec_final
        interpolated_vectors[i, :] = vector
    return np.array(interpolated_vectors)

# Initial and final vectors
# TO DO
# - aesthetics
# - incorporate beta-vecs into the animation

from VectorCoordCombo import get_ref_vecs

ref_state, vec_open, vec_closed = get_ref_vecs(cf.struct_head)
initial_vector = vec_closed
final_vector = vec_open

ideal_rotation = True
if ideal_rotation:

    # Number of frames in the animation
    rotation_frames = 100
    pause_frames = 20 
    total_frames = rotation_frames + 2 * pause_frames

    # Generate the array of interpolated vectors
    vectors_array = interpolate_vectors(
                        initial_vector, 
                        final_vector, 
                        rotation_frames)

else: 

    # Use the sampled beta vecs from along the path
    ani_path = f"{ cf.figure_head }/animations/strs_WT_apo_path"
    vectors_array = np.load(f"{ ani_path }/beta_vecs_on_path.npy")

    # Number of frames in the animation
    rotation_frames = vectors_array.shape[0]
    pause_frames = 20 
    total_frames = rotation_frames + 2 * pause_frames

# Create a figure and a 3D axis
fig = plt.figure()
ax = fig.add_subplot(111, projection='3d')
ax.set_xlim([-3, 1])
ax.set_ylim([-3, 1])
ax.set_zlim([-1, 3])

# Initialize the vector plot
vector_plot, = ax.plot(
                    [0, initial_vector[0]], 
                    [0, initial_vector[1]], 
                    [0, initial_vector[2]], 
                    marker="o", lw=3)

# Add a label for the alpha-carbons
origin_label = ax.text(*np.array((0.5,0,0)), r"$C_{\alpha}^{206}$" )
label = ax.text(*(vec_open + np.array((-0.2,0,0))), r"$C_{\alpha}^{215}$")

# Aesthetics
ax.grid(False)
ax.set_xticklabels([])
ax.set_yticklabels([])
ax.set_zticklabels([])

# Update function for the animation
def update(num):
    if num < pause_frames:
        # Pause at the initial position
        current_vector = initial_vector
    elif num < pause_frames + rotation_frames:
        # Rotate the vector
        current_vector = vectors_array[num - pause_frames, :]
    else:
        # Pause at the final position
        current_vector = final_vector
    
    # Move the label
    label.set_position((current_vector[0] + 0.2, current_vector[1] - 0.2))
    label.set_z(current_vector[2] + 0.2)
    vector_plot.set_data([0, current_vector[0]], [0, current_vector[1]])
    vector_plot.set_3d_properties([0, current_vector[2]])

    return vector_plot, label

# Create the animation
ani = FuncAnimation(fig, update, frames=total_frames, interval=50, 
            blit=True)

# Set the view
ax.view_init(elev=90, azim=90)

# Add interactivity
mplcursors.cursor(ax, hover=True)

# Save the animation
if ideal_rotation:

    ani.save(f"{ cf.figure_head }/animations/beta_vec_ref_interpolate.mp4", 
                writer='ffmpeg')

else:

    ani.save(f"{ cf.figure_head }/animations/beta_vec_sampled_interpolate.mp4", 
                writer='ffmpeg')

# Display the plot
plt.show()
```
