## Supplementary material for "*Clostridioides difficile* Toxins Unhinged: Allosterically Switchable Network Orients *β*-flap": Scripts, Structures etc.: AveIPL_charges.py

```
# -*- coding: utf-8 -*-

import os
import sys
from numpy import mean, std
import matplotlib.pyplot as plt
import matplotlib.image as mpimg
from matplotlib.gridspec import GridSpec
sys.path.insert(0, "/home/lf1071fu/project_b3/ProjectB3")
import config.settings as cf
from tools import utils, traj_funcs
import re

def main(argv):
    """
    Finds the average charges from multiconformational RESP fits.
    
    Parameters
    ----------
    Subdirectories for the configurations are named 'config_xx'. The .pdb in 
    config_0 directory is used to define the atoms by number in the dictionary
    Residue.atoms. The RESP output file from the second iteration, 'resp2.out' is
    used to retrieve the fitted charge from each configuration.
    
    Returns
    -------
    Prints the average charges and makes a plot of the charges. 
    
    """
    # Set up path variables
    home = os.getcwd()
    
    # Read in a pdb file
    os.chdir(home + "/config_0")
    pdb_lines = read_file("conform.pdb")
    for line in pdb_lines:
        line = line.split()
        if "ATOM" in line:
            Residue.pdb.append(line)
    
    # Use the .pdb file to define atoms of the residue
    for line in Residue.pdb:
        Residue.atoms[int(line[1])-3996] = Residue(line[1], line[2])
    os.chdir(home)
    
    # Collect all charges from conformations for each atom in a list
    folders = [f for f in os.listdir() if "config" in f]
    for folder in folders:
        os.chdir(folder)
        print(folder)
        include_charges()
        os.chdir(home)
    
    # Print out info on atom, including its average charge
    print(*[str(item) for _, item in Residue.atoms.items()], sep="\n")    
    
    # Make plot 
    make_plot()
    
class Residue():
    """
    The atoms belonging to the residue are stored within the dictionary 
    Residue.atoms, where individual atoms are accessed using their integer 
    values from the .pdb file as the key. 
    
    """
    pdb = []
    atoms = dict()
    
    def __init__(self, num, name):
        self.num = int(num) - 3996
        self.name = name
        self.charges = []
        self.element = "unknown"

    def __str__(self):
        return "Number: {}, Name: {}, Element: {}, Configs: {}, Charge: {}00, STD: {}"\
            .format(self.num, self.name, self.element, \
                len(self.charges), round(mean(self.charges), 4),\
                    round(std(self.charges), 4))
        
def read_file(file_name):
    """
    Return the lines of a file in a list.
    
    Parameters
    ----------
    file_name : file
        A text file, such as a .pdb or .mol2 file.
    
    Returns
    -------
    result : string list
        An ordered, line-separated list of strings from the file.
    
    """
    try:     
        with open(file_name, mode='r') as file: 
            lines = file.readlines()
    except OSError:
        print("file", file_name, "not found.")
    file_lines = []
    for line in lines:
        file_lines.append(line)
    return file_lines
            
def include_charges():
    """
    Include RESP fit charges for a configuration in Residue.atoms[x].charges
    
    For each atom 'x' in the residue, the RESP fitted charge for the 
    conformation in the current working directory is added to the charges list.
    The second RESP fit should be present in the folder as "resp2.out".
    
    """
    resp_out = read_file("IPL-resp2.out")
    
    # find the relevant lines from output for charges
    for i, line in enumerate(resp_out):
        if "Point Charges Before & After Optimization" in line:
            begin = i + 3
        if "Sum over the calculated charges:" in line:
            end = i - 1
    charges = resp_out[begin : end ]
    
    # Append the charges to the matching atom's charge list under: 
    # Residue.atoms[x].charges
    check = False
    for line in charges:
        number = int(line.split()[0])
        charge = float(line.split()[3])
        if -1.5 > charge or charge > 1.5: 
            check = True
        Residue.atoms[number].charges.append(charge) 
        Residue.atoms[number].element = line.split()[1]
    if check:
        print("Check RESP output: ", os.getcwd())

def make_plot():
    """
    Make a plot of charges, including std error bar and picture of \
    structure.

    Parameters
    ----------
    name : str
        Name of the structure. e.g. "FY1", "FY2" etc.
    
    """
    names = [atom.name for _, atom in Residue.atoms.items()]
    charges = [round(mean(atom.charges), 4) for _, atom in Residue.atoms.items()]
    stdev = [round(std(atom.charges), 4) for _, atom in Residue.atoms.items()]

    # Zip and sort the data
    zipped = list(zip(names, charges, stdev))
    sorted_zipped = sorted(zipped, key=lambda x: (x[0][0], int(x[0][1:])))
    names, charges, stdev = zip(*sorted_zipped)
    
    fig, ax = plt.subplots(constrained_layout=True, figsize=(2.5,2.5))

    # Color by atom group
    ax.scatter(names[0:6], charges[0:6], s=10, color="#000000", label="atomic charges \n[H$_3$L]$^{9-}$")
    plt.errorbar(names[0:6], charges[0:6], yerr=stdev[0:6], fmt="none", elinewidth=1)
    ax.scatter(names[6:12], charges[6:12], s=10, color="#cccccc")
    plt.errorbar(names[6:12], charges[6:12], yerr=stdev[6:12], fmt="none", elinewidth=1)
    ax.scatter(names[12:15], charges[12:15], s=10, color="#757575")
    plt.errorbar(names[12:15], charges[12:15], yerr=stdev[12:15], fmt="none", elinewidth=1)
    ax.scatter(names[15:21], charges[15:21], s=10, color="#ff6e6e")
    plt.errorbar(names[15:21], charges[15:21], yerr=stdev[15:21], fmt="none", elinewidth=1)
    ax.scatter(names[21:39], charges[21:39], s=10, color="#e81e1e")
    plt.errorbar(names[21:39], charges[21:39], yerr=stdev[21:39], fmt="none", elinewidth=1)
    ax.scatter(names[39:46], charges[39:46], s=10, color="#bf6317")
    plt.errorbar(names[39:46], charges[39:46], yerr=stdev[39:46], fmt="none", elinewidth=1)

    ax.tick_params(axis='y', which='both', labelsize=8)
    ax.xaxis.set_visible(False)
    plt.xlabel("Atom Names", labelpad=3)
    plt.ylabel("Atomic Partial Charge (e)", labelpad=3)
    plt.legend(fontsize=6)
    ax.set_ylim(-2,2)

    ax.grid(False)

    for i in ["top","bottom","left","right"]:
        ax.spines[i].set_linewidth(1) 

    fig_path = f"{ cf.figure_head }/paper-figs/supplementary"

    utils.save_figure(fig, f"{ fig_path }/IPL_charges.svg")
    plt.show()
    plt.close()

    return None

if __name__ ==  '__main__':
    main(sys.argv)
    sys.exit(0)
```
