## Supplementary material for "*Clostridioides difficile* Toxins Unhinged: Allosterically Switchable Network Orients *β*-flap": Scripts, Structures etc.: ExtractPath_strs.py

```
import sys
sys.path.insert(0, "/home/lf1071fu/project_b3/ProjectB3")
from tools import utils, traj_funcs
import config.settings as cf
import numpy as np
import pandas as pd
import os
import subprocess
import time
import MDAnalysis as mda
from MDAnalysis.analysis import align

ani_path = f"{ cf.figure_head }/animations/strs_WT_apo_path"
apo_WT_path = f"{ cf.data_head }/umbrella/apo_state/nobackup"

# What are the 2d CV values along the low free energy path?
cv_along_path = np.load(f"{ ani_path }/cv_lowE_path.npy")

# grab closest sampled structure to CV value from the biased data
def nearby_struct_data(cv_point, df_cat):
    data = (df_cat["opendot"], df_cat["closeddot"])

    d = np.sqrt(np.sum(np.square(np.transpose(data) 
                                 - np.array(cv_point)), axis=1))
    min_d = np.min(d)
    min_ind = np.argmin(d)
    window, run, time = df_cat.iloc[min_ind][["window", "run", "time"]]

    return str(int(window)), str(int(run)), int(time) 

# Get sampled collective variables in one table
df_cat = pd.DataFrame(columns=["time", "opendot", "closeddot"])
from biased.NewWindowConfs import add_colvar_data
for w in range(1,181):
    for r in range(1,5):
        file = f"{ apo_WT_path }/window{ w }/run{ r }/COLVAR_{ w }.dat"
        if os.path.exists(file):
            df_new = add_colvar_data(w, r, file)

            df_cat = pd.concat([df_cat, df_new])

# Iterate over the points along the path and
# Extract structure associated with each cv point
n, _ = cv_along_path.shape
print(n)
for i in range(n):

    # Identify a nearby structure with window, run and time
    w, r, time_frame = nearby_struct_data(cv_along_path[i,:2], df_cat)

    # Paths for the simulation data
    traj = f"{ apo_WT_path }/window{ w }/run{ r }/fitted_traj.xtc"
    top = f"{ apo_WT_path }/window{ w }/run{ r }/w{ w }_r{ r }.tpr"
    struct_out = f"{ ani_path }/structure_{ i }.pdb"

    # Define the gromacs command
    gmx = ["echo", "1", "|", "gmx", "trjconv", "-f", 
        traj, "-s", top, "-o", struct_out, "-b", 
        str(time_frame - 1000), "-dump", str(time_frame), "-nobackup"]
    print("\n", " ".join(gmx), "\n")

    # Use gromacs subprocess to extract the conformation at the 
    # desired time
    process = subprocess.Popen(" ".join(gmx), 
                                stdin=subprocess.PIPE, 
                                stdout=subprocess.PIPE,
                                shell=True, text=True)

    # Pass input to the GROMACS command to use protein + ligand
    stdout, stderr = process.communicate("1\n")
    print("Output:", stdout)
    print("Error:", stderr)

    time.sleep(1)

# Grab ref for alignment
ref_state = mda.Universe(f"{ cf.struct_head }/ref_all_atoms.pdb", 
                            length_unit="nm")
core_res, core = traj_funcs.get_core_res()

# Translate structures into beta-vectors
beta_vecs = np.zeros((n,3))
print(beta_vecs.shape)

for i in range(n):
    # Load in str
    struct = mda.Universe(f"{ ani_path }/structure_{ i }.pdb", 
                            length_unit="nm")

    # Center structure on the reference
    align.alignto(struct, ref_state, select=core)

    # Get calpha positions and determine beta vec
    atom1 = struct.select_atoms(f"name CA and resnum { cf.r1 }"
                ).positions[0]
    atom2 = struct.select_atoms(f"name CA and resnum { cf.r2 }"
                ).positions[0]
    beta_vec = atom2/10 - atom1/10
    print(beta_vec)

    # Add beta_vec to array
    beta_vecs[i,:] = beta_vec

# Store beta-vec array
beta_vecs_path = f"{ ani_path }/beta_vecs_on_path.npy"
utils.save_array(beta_vecs_path, beta_vecs)
```
