## Supplementary material for "*Clostridioides difficile* Toxins Unhinged: Allosterically Switchable Network Orients *β*-flap": Scripts, Structures etc.: TablePlot_barrier.py

```
import os
import sys
import datetime
import matplotlib.pyplot as plt
import numpy as np
import pandas as pd
sys.path.insert(0, "/home/lf1071fu/project_b3/ProjectB3")
from tools import utils, traj_funcs
import config.settings as cf
from tools import utils, traj_funcs
import seaborn as sns

# Load data into tables
df_hists_apo_open = pd.read_csv(f"{ cf.data_head }/unbiased_sims/combo_unbiased_apo-open.csv")
df_hists_apo_closed = pd.read_csv(f"{ cf.data_head }/unbiased_sims/combo_unbiased_apo-closed.csv")
df_hists_holo_open = pd.read_csv(f"{ cf.data_head }/unbiased_sims/combo_unbiased_holo-open.csv")
df_hists_holo_closed = pd.read_csv(f"{ cf.data_head }/unbiased_sims/combo_unbiased_holo-closed.csv")
df_hists_tcda = pd.read_csv(f"{ cf.data_head }/tcda/combo_tcda.csv")
df_tcda_apo_open = pd.read_csv(f"{ cf.data_head }/tcda/combo_tcda_TcdA-apo-open.csv")
df_tcda_apo_closed = pd.read_csv(f"{ cf.data_head }/tcda/combo_tcda_TcdA-apo-closed.csv")
df_tcda_holo_open = pd.read_csv(f"{ cf.data_head }/tcda/combo_tcda_TcdA-holo-open.csv")
df_tcda_holo_closed = pd.read_csv(f"{ cf.data_head }/tcda/combo_tcda_TcdA-holo-closed.csv")
df_apo_ave = pd.read_csv(f"{ cf.data_head }/umbrella/apo_state/nobackup/windows_averages_merged.csv")
df_holo_ave = pd.read_csv(f"{ cf.data_head }/umbrella/holo_state/nobackup/window_data/windows_averages_merged.csv")

# Load in settings
selections_tcda = cf.selections_tcda
selections = cf.selections
styles = cf.styles

# Abreviated selections
selections = {
    "K775--E692" : ("resid 232 and name NZ*", "resid 149 and name OE*"),
    "K775--N740" : ("resid 232 and name NZ*", "resid 197 and name OD1"),
    }

# TcdA residue analogues
tcda_analogues = {
                "K775--N740" : "K776--N741",
                "K775--E692" : "K776--E693"
                }

# Other vars
dfs = {"apo-open" : df_hists_apo_open, "apo-closed" : df_hists_apo_closed, 
        "holo-open" : df_hists_holo_open, "holo-closed" : df_hists_holo_closed}
df_tcda = {"TcdA-apo-open" : df_tcda_apo_open, "TcdA-apo-closed" : df_tcda_apo_closed, 
        "TcdA-holo-open" : df_tcda_holo_open, "TcdA-holo-closed" : df_tcda_holo_closed}

# Set up subplot dimensions
# Columns - 1) Text with contact info 2) 4 state hists 3) apo window aves 
# 4) holo window aves 5) TcdA 4 state hists
plt.rcParams['figure.constrained_layout.use'] = False
n_rows = len(selections) + 2
n_cols = 6
sns.set_context("paper") 
import matplotlib.gridspec as gridspec
fig, axes = plt.subplots(figsize=(8,2.5), constrained_layout=True)
gs = gridspec.GridSpec(n_rows, n_cols, width_ratios=[5,8,4,5,1,8], 
    height_ratios=([1] + (n_rows-2)*[6] + [1]))

ax0 = plt.subplot(gs[0,0])
ax0.axis('off')

# Column labels
ax1 = plt.subplot(gs[0,1])
ax1.text(0.5, .9, "Contact histograms\nfor TcdB CPD", ha='center', va='bottom', fontsize=10)
ax1.axis('off')
ax2 = plt.subplot(gs[0,2])
ax2.text(0.5, .9, "Average\ndistance\n(apo state)", ha='center', va='bottom', fontsize=10)
ax2.axis('off')
ax3 = plt.subplot(gs[0,3])
ax3.text(0.5, .9, "Average\ndistance\n(holo state)", ha='center', va='bottom', fontsize=10)
ax3.axis('off')
ax4 = plt.subplot(gs[0,5])
ax4.text(0.5, .9, "Contact histograms\nfor TcdA CPD", ha='center', va="bottom", fontsize=10)
ax4.axis("off")

# Iterate over the contacts
for i, col in enumerate(selections.keys()):

    # First column is the description part
    row_descript = plt.subplot(gs[i+1,0])
    row_descript.text(0.7, 0.6, col, ha='center', va='bottom', fontsize=10)
    row_descript.text(0.7, 0.2, tcda_analogues[col] + "\nin TcdA", 
        ha='center', va='bottom', fontsize=8, color="#2e2e2e")
    row_descript.axis('off')

    # Second column is for the histograms
    hist_ax = plt.subplot(gs[i+1,1])
    for state, df in dfs.items():
        if "IP6" in col and "apo" in state:
            continue
        hist_ax.hist(df[col], bins=25, density=True, color=styles[state][0], 
                ls=styles[state][1], histtype='step', lw=2, 
                alpha=styles[state][2])
    hist_ax.set_xlim(0,25)
    for j in ["top","right","bottom","left"]:
        hist_ax.spines[j].set_visible(False)
    hist_ax.set_xticks([2, 5,25])
    if i == (len(selections) - 1):
        hist_ax.set_xticklabels([2,5,25], fontsize=8)
    else:
        hist_ax.set_xticklabels([])
    _, ymax = hist_ax.get_ylim()
    hist_ax.set_ylim(0,ymax)
    hist_ax.set_yticks([])
    _, ymax = plt.subplot(gs[i+1,1]).get_ylim()
    plt.subplot(gs[i+1,1]).set_ylim(0,ymax)

    # Determine range for colorbar plot
    from matplotlib.colors import Normalize
    ave_val_holo = df_holo_ave.groupby('Window')[col].mean().values
    if "IP6" not in col:
        ave_val_apo = df_apo_ave.groupby('Window')[col].mean().values
        combo = np.concatenate((ave_val_apo, ave_val_holo))
    else:
        combo = ave_val_holo
    if "K775--N740" in col:
        norm = Normalize(vmin=np.min(combo), vmax=12.5)
    elif "K775--E692" in col:
        norm = Normalize(vmin=np.min(combo), vmax=12.5)
    else:
        norm = Normalize(vmin=np.min(combo), vmax=np.max(combo))

    # Third colomn for apo window aves
    apo_ave = plt.subplot(gs[i+1,2])
    
    d_apo = apo_ave.tricontourf(
        df_apo_ave[df_apo_ave["Run"] == 1]["OpenPoints"], 
        df_apo_ave[df_apo_ave["Run"] == 1]["ClosedPoints"], 
        ave_val_apo, 10, cmap="coolwarm", levels=100, norm=norm, extend="max")
    apo_ave.set_xticks([])
    apo_ave.set_yticks([])
    apo_ave.set_aspect('equal', adjustable='box')
    apo_ave.axis("off")

    # Fourth colomn for holo window aves
    holo_ave = plt.subplot(gs[i+1,3])
    d_holo = holo_ave.tricontourf(
            df_holo_ave[df_holo_ave["Run"] == 1]["OpenPoints"], 
            df_holo_ave[df_holo_ave["Run"] == 1]["ClosedPoints"], 
            ave_val_holo, 10, cmap="coolwarm", levels=100, norm=norm,
            extend="max")
    holo_ave.set_xticks([])
    holo_ave.set_yticks([])
    holo_ave.axis("off")
    holo_ave.set_aspect('equal', adjustable='box')
    # Color bar settings
    from matplotlib.ticker import MaxNLocator, FuncFormatter
    if np.max(combo) in ave_val_apo:
        cbar = plt.colorbar(d_apo, ax=holo_ave)
    else:
        cbar = plt.colorbar(d_holo, ax=holo_ave)
    cbar.ax.set_yticks([4,6,8,10,12], 
        labels=["4.0","6.0","8.0","10.0","12.0"])
    cbar.ax.set_ylim([0,12.1])
    cbar.ax.tick_params(axis='both', length=1)
    for spine in cbar.ax.spines.values():
        spine.set_visible(False)

    # Spacer column
    space_ax = plt.subplot(gs[i+1,4])
    space_ax.axis('off')

    # Fifth column is for the TcdA histograms
    tcda_ax = plt.subplot(gs[i+1,5])
    tcda_col = tcda_analogues[col]
    for state, df in df_tcda.items():
        if "IP6" in col and "apo" in state:
            continue
        tcda_ax.hist(df[tcda_col], bins=25, density=True, color=styles[state][0], 
                ls=styles[state][1], histtype='step', lw=2, 
                alpha=styles[state][2])
    tcda_ax.set_xlim(0,25)
    for j in ["top","right","bottom","left"]:
        tcda_ax.spines[j].set_visible(False)
    tcda_ax.set_xticks([2, 5,25])
    if i == (len(selections) - 1):
        tcda_ax.set_xticklabels([2,5,25], fontsize=8)
    else:
        tcda_ax.set_xticklabels([])
    _, ymax = tcda_ax.get_ylim()
    tcda_ax.set_ylim(0, ymax)
    tcda_ax.set_yticks([])
    _, ymax = plt.subplot(gs[i+1,4]).get_ylim()
    plt.subplot(gs[i+1,4]).set_ylim(0,ymax)

# Column xaxis labels
ax1 = plt.subplot(gs[n_rows - 1,1])
ax1.text(0.5, .1, "Distance ($\AA$)", ha='center', va='center', 
    fontsize=10)
ax1.axis('off')
ax2 = plt.subplot(gs[n_rows - 1,2])
ax2.text(0.5, 0.2, r"($\xi_1 \times \xi_2$)", 
    ha='center', va='top', fontsize=10)
ax2.axis('off')
ax3 = plt.subplot(gs[n_rows - 1,3])
ax3.text(0.5, 0.2, r"($\xi_1 \times \xi_2$)", 
    ha='center', va='top', fontsize=10)
ax3.axis('off')
ax4 = plt.subplot(gs[n_rows - 1,5])
ax4.text(0.5, 0.1, "Distance ($\AA$)", ha='center', va="center", 
    fontsize=10)
ax4.axis("off")

from matplotlib.lines import Line2D
line = Line2D([0.71, 0.71], [0, 1], color='k', linestyle='--', 
    transform=fig.transFigure, figure=fig)
fig.add_artist(line)


#plt.tight_layout()
utils.save_figure(fig, f"{ cf.figure_head }/contacts_table_barrier.png")
plt.show()

plt.close()
```
