## Supplementary material for "*Clostridioides difficile* Toxins Unhinged: Allosterically Switchable Network Orients *β*-flap": Scripts, Structures etc.: TablePlot_network_suppl.py

```
import os
import sys
import datetime
import matplotlib.pyplot as plt
import numpy as np
import pandas as pd
sys.path.insert(0, "/home/lf1071fu/project_b3/ProjectB3")
from tools import utils, traj_funcs
import config.settings as cf
from tools import utils, traj_funcs
import seaborn as sns

# Load data into tables
df_hists_apo_open = pd.read_csv(f"{ cf.data_head }/unbiased_sims/combo_unbiased_apo-open.csv")
df_hists_apo_closed = pd.read_csv(f"{ cf.data_head }/unbiased_sims/combo_unbiased_apo-closed.csv")
df_hists_holo_open = pd.read_csv(f"{ cf.data_head }/unbiased_sims/combo_unbiased_holo-open.csv")
df_hists_holo_closed = pd.read_csv(f"{ cf.data_head }/unbiased_sims/combo_unbiased_holo-closed.csv")
df_hists_tcda = pd.read_csv(f"{ cf.data_head }/tcda/combo_tcda.csv")
df_tcda_apo_open = pd.read_csv(f"{ cf.data_head }/tcda/combo_tcda_TcdA-apo-open.csv")
df_tcda_apo_closed = pd.read_csv(f"{ cf.data_head }/tcda/combo_tcda_TcdA-apo-closed.csv")
df_tcda_holo_open = pd.read_csv(f"{ cf.data_head }/tcda/combo_tcda_TcdA-holo-open.csv")
df_tcda_holo_closed = pd.read_csv(f"{ cf.data_head }/tcda/combo_tcda_TcdA-holo-closed.csv")
df_apo_ave = pd.read_csv(f"{ cf.data_head }/umbrella/apo_state/nobackup/windows_averages.csv")
df_holo_ave = pd.read_csv(f"{ cf.data_head }/umbrella/holo_state/nobackup/window_data/windows_averages.csv")

# Load in settings
selections_tcda = cf.selections_tcda
selections = cf.selections
styles = cf.styles

# Abreviated selections
selections = {"E592--W761" : ("resid 218 and name NE1", "resid 49 and name OE*"),
    "N596--E592" : ("resid 49 and name OE*", "resid 53 and name ND2"),
    "H757 SASA" : (),
    "W761 SASA" : (),
    "E749 SASA" : (),
    "E743 SASA" : ()
    }

# TcdA residue analogues
tcda_analogues = {"R751--N747" : "R752--N748", "N747--E753" : "N748--E754", 
                "E753--R745" : "E754--R746", "R745--W761" : "R746--W762", 
                "K600--E743" : "K601--E744", "E743--N596" : "E744--N597",
                "C698--E743" : "C699--E744", "C698--H757" : "C699--H758",
                "K600--E749" : "K601--E750", "K600--IP6" : "K601--IP6",
                "E592--W761" : "E593--W762", "N596--E592" : "N597--E593",
                "K775--IP6" : "K776--IP6", "K764--IP6" : "K765--IP6",
                "R751--IP6" : "R752--IP6", "R752--IP6" : "K753--IP6",
                "D771--K775" : "D772--K776", "K764--D756" : "K765--A757", 
                "R752--E765" : "K753--E766", "H757 SASA" : "H758 SASA",
                "W761 SASA" : "W762 SASA", "E749 SASA" : "E750 SASA", 
                "E743 SASA" : "E744 SASA"}

# Other vars
dfs = {"apo-open" : df_hists_apo_open, "apo-closed" : df_hists_apo_closed, 
        "holo-open" : df_hists_holo_open, "holo-closed" : df_hists_holo_closed}
df_tcda = {"TcdA-apo-open" : df_tcda_apo_open, "TcdA-apo-closed" : df_tcda_apo_closed, 
        "TcdA-holo-open" : df_tcda_holo_open, "TcdA-holo-closed" : df_tcda_holo_closed}

# Set up subplot dimensions
# Columns - 1) Text with contact info 2) 4 state hists 3) apo window aves 
# 4) holo window aves 5) TcdA 4 state hists
plt.rcParams['figure.constrained_layout.use'] = False
n_rows = len(selections) + 3
n_cols = 6
sns.set_context("paper") 
import matplotlib.gridspec as gridspec
fig, axes = plt.subplots(figsize=(8, 7), constrained_layout=True)
gs = gridspec.GridSpec(n_rows, n_cols, width_ratios=[6,8,4,5,1,8], 
    height_ratios=([1,6,6,1,6,6,6,6,1]))

ax0 = plt.subplot(gs[0,0])
ax0.axis('off')

# Column labels
ax1 = plt.subplot(gs[0,1])
ax1.text(0.5, .9, "Contact histograms\nfor TcdB CPD", ha='center', 
    va='bottom', fontsize=10)
ax1.axis('off')
ax2 = plt.subplot(gs[0,2])
ax2.text(0.5, .9, "Average\ndist./SASA\n(apo state)", ha='center', 
    va='bottom', fontsize=10)
ax2.axis('off')
ax3 = plt.subplot(gs[0,3])
ax3.text(0.5, .9, "Average\ndist./SASA\n(holo state)", ha='center', 
    va='bottom', fontsize=10)
ax3.axis('off')
ax4 = plt.subplot(gs[0,5])
ax4.text(0.5, .9, "Contact histograms\nfor TcdA CPD", ha='center', 
    va="bottom", fontsize=10)
ax4.axis("off")

# Iterate over the contacts
for i, col in enumerate(selections.keys()):

    if i >= 2:
        i += 1

    # First column is the description part
    row_descript = plt.subplot(gs[i+1,0])
    row_descript.text(0.7, 0.6, col, ha='center', va='bottom', fontsize=10)
    row_descript.text(0.7, 0.2, tcda_analogues[col] + "\nin TcdA", 
        ha='center', va='bottom', fontsize=8, color="#2e2e2e")
    row_descript.axis('off')

    # Second column is for the histograms
    hist_ax = plt.subplot(gs[i+1,1])
    for state, df in dfs.items():
        if "IP6" in col and "apo" in state:
            continue
        hist_ax.hist(df[col], bins=25, density=True, color=styles[state][0], 
                ls=styles[state][1], histtype='step', lw=2, 
                alpha=styles[state][2])
    hist_ax.set_xlim(0,25)
    for j in ["top","right","bottom","left"]:
        hist_ax.spines[j].set_visible(False)
    if i < 2:
        hist_ax.set_xlim(0,25)
        hist_ax.set_xticks([2, 5, 25])
    else:
        hist_ax.set_xlim(0,3)
        hist_ax.set_xticks([0, 1, 2, 3])
    if col == "N596--E592":
        hist_ax.set_xticklabels([2,5,25], fontsize=8)
    elif col == "E743 SASA":
        hist_ax.set_xticklabels([0,1,2,3], fontsize=8)
    else:
        hist_ax.set_xticklabels([])
    _, ymax = hist_ax.get_ylim()
    hist_ax.set_ylim(0,ymax)
    hist_ax.set_yticks([])
    _, ymax = plt.subplot(gs[i+1,1]).get_ylim()
    plt.subplot(gs[i+1,1]).set_ylim(0,ymax)

    # Determine range for colorbar plot
    from matplotlib.colors import Normalize
    ave_val_apo = df_apo_ave.groupby('Window')[col].mean().values
    ave_val_holo = df_holo_ave.groupby('Window')[col].mean().values
    combo = np.concatenate((ave_val_apo, ave_val_holo))
    if col == "W761 SASA":
        norm = Normalize(vmin=np.min(combo), vmax=1.3)
    elif col == "E743 SASA":
        norm = Normalize(vmin=np.min(combo), vmax=.8)
    elif col == "E592--W761":
        norm = Normalize(vmin=np.min(combo), vmax=21)
    else:
        norm = Normalize(vmin=np.min(combo), vmax=np.max(combo))

    # Third colomn for apo window aves
    if "IP6" in col:
        print("hi")
    else:
        apo_ave = plt.subplot(gs[i+1,2])
        
        d_apo = apo_ave.tricontourf(
            df_apo_ave[df_apo_ave["Run"] == 1]["OpenPoints"], 
            df_apo_ave[df_apo_ave["Run"] == 1]["ClosedPoints"], 
            ave_val_apo, 10, cmap="coolwarm", levels=100, norm=norm)
        apo_ave.set_xticks([])
        apo_ave.set_yticks([])
        apo_ave.axis("off")

    # Fourth colomn for holo window aves
    holo_ave = plt.subplot(gs[i+1,3])
    d_holo = holo_ave.tricontourf(
            df_holo_ave[df_holo_ave["Run"] == 1]["OpenPoints"], 
            df_holo_ave[df_holo_ave["Run"] == 1]["ClosedPoints"], 
            ave_val_holo, 10, cmap="coolwarm", levels=100, norm=norm)
    holo_ave.set_xticks([])
    holo_ave.set_yticks([])
    holo_ave.axis("off")
    # Color bar settings
    from matplotlib.ticker import MaxNLocator, FuncFormatter
    if np.max(combo) in ave_val_apo:
        cbar = plt.colorbar(d_apo, ax=holo_ave)
    else:
        cbar = plt.colorbar(d_holo, ax=holo_ave)
    cbar.locator = MaxNLocator(nbins=5)  # Forces exactly 5 bins
    cbar.update_ticks()
    cbar.ax.tick_params(axis='both', length=1)
    formatter = FuncFormatter(lambda x, pos: f"{np.round(x,1)}")
    cbar.ax.yaxis.set_major_formatter(formatter)
    for spine in cbar.ax.spines.values():
        spine.set_visible(False)

    # Spacer column
    space_ax = plt.subplot(gs[i+1,4])
    space_ax.axis('off')

    # Fifth column is for the TcdA histograms
    tcda_ax = plt.subplot(gs[i+1,5])
    tcda_col = tcda_analogues[col]
    for state, df in df_tcda.items():
        if "IP6" in col and "apo" in state:
            continue
        tcda_ax.hist(df[tcda_col], bins=25, density=True, color=styles[state][0], 
                ls=styles[state][1], histtype='step', lw=2, 
                alpha=styles[state][2])
    for j in ["top","right","bottom","left"]:
        tcda_ax.spines[j].set_visible(False)
    if i < 2:
        tcda_ax.set_xlim(0,25)
        tcda_ax.set_xticks([2, 5, 25])
    else:
        tcda_ax.set_xlim(0,3)
        tcda_ax.set_xticks([0, 1, 2, 3])
    if col == "N596--E592":
        tcda_ax.set_xticklabels([2,5,25], fontsize=8)
    elif col == "E743 SASA":
        tcda_ax.set_xticklabels([0,1,2,3], fontsize=8)
    else:
        tcda_ax.set_xticklabels([])
    _, ymax = tcda_ax.get_ylim()
    tcda_ax.set_ylim(0, ymax)
    tcda_ax.set_yticks([])
    _, ymax = plt.subplot(gs[i+1,4]).get_ylim()
    plt.subplot(gs[i+1,4]).set_ylim(0,ymax)

# Column xaxis labels
dist_label = plt.subplot(gs[3,1])
dist_label.text(0.5, .1, "Distance ($\AA$)", ha='center', va='center', 
    fontsize=10)
# apo_coord = plt.subplot(gs[3,2])
# apo_coord.text(0.5, .1, "Distance ($\AA$)", 
#     ha='center', va='center', fontsize=10)
# holo_coord = plt.subplot(gs[3,3])
# holo_coord.text(0.5, .1, "Distance ($\AA$)", 
#     ha='center', va='center', fontsize=10) 
dist_label.axis('off')
ax2 = plt.subplot(gs[n_rows - 1,2])
ax2.text(0.5, .1, r"($\xi_1 \times \xi_2$)", 
    ha='center', va='top', fontsize=10)
ax2.axis('off')
ax3 = plt.subplot(gs[n_rows - 1,3])
ax3.text(0.5, 0.1, r"($\xi_1 \times \xi_2$)", 
    ha='center', va='top', fontsize=10)
ax3.axis('off')
dist_label_tcda = plt.subplot(gs[3,5])
dist_label_tcda.text(0.5, 0.1, "Distance ($\AA$)", ha='center', va="center", 
    fontsize=10)
dist_label_tcda.axis("off")
sasa_label = plt.subplot(gs[n_rows - 1,1])
sasa_label.text(0.5, 0.1, "SASA (nm²)", ha='center', va='top', 
    fontsize=10)
sasa_label.axis('off')
sasa_label_tcda = plt.subplot(gs[n_rows - 1,5])
sasa_label_tcda.text(0.5, 0.1, "SASA (nm²)", ha='center', va='top', 
    fontsize=10)
sasa_label_tcda.axis('off')

from matplotlib.lines import Line2D
line = Line2D([0.71, 0.71], [0, 1], color='k', linestyle='--', 
    transform=fig.transFigure, figure=fig)
fig.add_artist(line)

#plt.tight_layout()
utils.save_figure(fig, f"{ cf.figure_head }/contacts_table_network_suppl.png")
plt.show()

plt.close()
```
