## Supplementary material for "*Clostridioides difficile* Toxins Unhinged: Allosterically Switchable Network Orients *β*-flap": Scripts, Structures etc.: PCA.py

```
import sys
import numpy as np
import os
import argparse
import matplotlib.pyplot as plt
import MDAnalysis as mda
from MDAnalysis.analysis import align, pca
import pandas as pd
import seaborn as sns
import config.settings as c
from tools import utils, traj_funcs

def main(argv):

    # Add command line arg to control group selection
    try:
        parser = argparse.ArgumentParser()

        parser.add_argument("-g", "--group",
    						action = "store",
    						dest = "group",
    						default = "beta-flap",
    						help = "Choose the subgroup for PCA. Options = backbone, "\
                                    "alpha-flap, beta-flap, calphas, Default = beta-flap")
        parser.add_argument("-p", "--path",
                            action = "store",
                            dest = "path",
                            default = "/home/lf1071fu/project_b3/simulate/unbiased_sims/apo_open/nobackup",
                            help = """Set path to the data directory.""")
        parser.add_argument("-f", "--fig_path",
                            action = "store",
                            dest = "fig_path",
                            default = "/home/lf1071fu/project_b3/figures/unbiased_sims/apo_open",
                            help = """Set path to the data directory.""")
        parser.add_argument("-t", "--topol",
                            action = "store",
                            dest = "topol",
                            default = "topol_protein.top",
                            help = """File name for topology, inside the path directory.""")   
        parser.add_argument("-x", "--xtc",
                            action = "store",
                            dest = "xtc",
                            default = "fitted_traj.xtc",
                            help = """File name for trajectory, inside the path directory.""")     
        parser.add_argument("-c", "--conform",
                            action = "store",
                            dest = "conform",
                            default = "open",
                            help = """The reference conformational state, i.e. "open" or "closed".""")
        args = parser.parse_args()

    except argparse.ArgumentError:
    	print("Command line arguments are ill-defined, please check the arguments")
    	raise

    global path_head

    # Assign group selection from argparse
    data_path = args.path
    group = args.group
    conform = args.conform
    topol = args.topol
    xtc = args.xtc
    fig_path = args.fig_path

    # Set up paths
    path_head = "/home/lf1071fu/project_b3"
    struct_path = f"{ path_head }/structures"

    # Set up selection strings for atom groups, using all atoms in the selected residues
    # beta_flap = "backbone and (resnum 195-231 or resnum 740-776)"
    if group == "alpha-flap":
        select_group = "resid 219-231 or resid 763-775"
    elif group == "backbone":
        select_group = "resid 8-251 or resid 552-795"
    elif group == "calphas":
        select_group = "name CA and (resid 8-251 or resid 552-795)"
    else:
        # The beta-flap group (default)
        select_group = "backbone and (resid 195-218 or resid 739-762)"
    ref_backbone = "resid 8-251 or resid 552-795"

    # Load in relevant reference structures
    if conform == "open":
        ref_state = mda.Universe(f"{ struct_path }/open_ref_state.pdb")
    elif conform == "closed":
        ref_state = mda.Universe(f"{ struct_path }/closed_ref_state.pdb")

    # Load in and align the traj
    u = mda.Universe(f"{ data_path }/{ topol }", f"{ data_path }/{ xtc }", 
                     topology_format="ITP", dt=1000)
    core_res, core = get_core_res() 
    align.AlignTraj(u, u.select_atoms("protein"), select=core, in_memory=True, dt=1000).run()
    uf = u.select_atoms(select_group)

    # Store calculated outputs as numpy arrays, use analysis dir up one level
    analysis_path = f"{ os.path.dirname(data_path) }/analysis/pca"
    if not os.path.exists(analysis_path):
        os.makedirs(analysis_path)
    fig_path = f"{ fig_path }/pc_{ group }"
    if not os.path.exists(fig_path):
        os.makedirs(fig_path)

    arrs = get_pc_data(u, select_group, uf, group, analysis_path)

    print(arrs["cumulated_var"])

    n_pcs = np.where(arrs["cumulated_var"] > 0.75)[0][0]
    print(f"The first { n_pcs } principal components explain at least 75% of the "\
           "total variance.\n")

    plot_eigvals(arrs["cumulated_var"], group, fig_path)

    plot_3PC(arrs["transformed"], group, fig_path)

    for i in range(4):
        pc_min_max(arrs["transformed"], u, group, i, fig_path)

    for i in range(4):
        visualize_PC(arrs["p_components"], arrs["transformed"], arrs["mean"], 
        i, uf, group, fig_path)

    return None

def get_pc_data(u, select_group, atom_group, group, path):
    """Determines PCA using MDAnalysis. 

    Parameters
    ----------
    u : mda.Universe
        
    """
    pc_files = {"p_components" : f"{ path }/p_components_{ group }.npy", 
                "cumulated_var" : f"{ path }/cumul_vaviance_{ group }.npy",
                "transformed" : f"{ path }/transformed_{ group }.npy",
                "mean" : f"{ path }/mean_{ group }.npy"}

    arrs = {}

    if all(list(map(lambda x : os.path.exists(x), pc_files))):

        print(
            "LOADING NUMPY ARRAYS"
        )

        for key, file in pc_files.items(): 
            arrs[key] = np.load(file, allow_pickle=True)

    else:

        print(
            "EVALUATING WITH MDANALYSIS"
        )

        pc = pca.PCA(u, select=select_group, align=True, n_components=50).run()
        arrs["p_components"] = pc.results.p_components
        arrs["cumulated_var"] = pc.results.cumulated_variance

        # Transforms the atom group into weights over each principal component
        # Here, weights of the first 3 components; shape --> (n_frames, n_PCs) 
        arrs["transformed"] = pc.transform(atom_group, n_components=3)

        arrs["mean"] = pc.mean.flatten()

        for key, file in pc_files.items():
            utils.save_array(file, arrs[key])

    return arrs

def plot_eigvals(cumulated_variance, group, fig_path):
    """Make a plot of the cummulative variance.

    The relative contribution to the overall variance by each principle
    component is shown for the first 25 PCs. This should simply demonstrate the
    number of relevant PCs in the dimensionality reduction.

    Parameters
    ----------
    cumulated_variance : np.1darray
        The cumulative variance, calculated from PCA eigenvalues.
    group : str
        The name of the subgroup selected for PCA.
    path : str
        The path to the directory where the figure should be saved.

    Returns
    -------
    None.

    """
    fig, ax = plt.subplots(constrained_layout=True)
    filled_marker_style = dict(marker='o', markersize=10, linestyle="-", lw=3,
                               markeredgecolor='#A31130')
    ax.plot(cumulated_variance[:25], color="#FF6666",
            **filled_marker_style)

    # Plot settings
    ax.tick_params(axis='y', labelsize=16, direction='in', width=2, \
                    length=5, pad=10)
    ax.tick_params(axis='x', labelsize=16, direction='in', width=2, \
                    length=5, pad=10)
    for i in ["top","bottom","left","right"]:
        ax.spines[i].set_linewidth(2)
    ax.grid(True)
    ax.set_xlabel(r"Principle component", labelpad=5, fontsize=16)
    ax.set_ylabel(r"Cumulative variance", labelpad=5, fontsize=16)

    print("FIG PATH:", fig_path)

    # Save fig
    utils.save_figure(fig, f"{ fig_path }/pca_scree_{ group }.png")
    plt.close()

    return None

def plot_3PC(transformed, group, fig_path):
    """Plot the first 3 principle components against each other. 

    Parameters
    ----------
    transformed : np.ndarray
        Trajectory projected onto the first 3 principle components.
    group : str
        The name of the subgroup selected for PCA.
    path : str
        The path to the directory where the figure should be saved.

    Returns
    -------
    None. 

    """
    df = pd.DataFrame(transformed, columns=['PC{}'.format(i+1) for i in range(3)])

    df["Time (ns)"] = df.index

    fig, axes2d = plt.subplots(nrows=3, ncols=3, sharex=True, sharey=True, 
                               constrained_layout=True)

    for i, row in enumerate(axes2d):
        for j, cell in enumerate(row):
            ts = df["Time (ns)"]
            cb = cell.scatter(df[f"PC{i+1}"], df[f"PC{j+1}"], c= ts, 
                              cmap="YlGnBu")
            if i == len(axes2d) - 1:
                cell.set_xlabel(f"PC {j+1}", fontsize=16)
            if j == 0:
                cell.set_ylabel(f"PC {i+1}", fontsize=16)
            if j == 2:
                cbar = fig.colorbar(cb, ax=cell)
                cbar.set_label("Time ($\mu$s)", fontsize=16)
                # cticks = list(map(lambda x: str(x/1000).split(".")[0],
                #       np.arange(0,len(ts),5000)))
                cbar.set_ticks(np.arange(0,len(ts),5000))
                ticks_loc = cbar.get_ticks().tolist()
                cbar.ax.yaxis.set_ticklabels([str(x/1000).split(".")[0] \
                                              for x in ticks_loc])
                cbar.ax.tick_params(labelsize=16, direction='out', width=2, length=5)
                cbar.outline.set_linewidth(2)
            cell.tick_params(axis='y', labelsize=16, direction='in', width=2,
                            length=5, pad=10)
            cell.tick_params(axis='x', labelsize=16, direction='in', width=2,
                            length=5, pad=10)
            for n in ["top","bottom","left","right"]:
                cell.spines[n].set_linewidth(2)

    # Save fig
    utils.save_figure(fig, f"{ fig_path }/pca_first3_{ group }.png")
    plt.show()
    plt.close()

    return None

def pc_min_max(transformed, u, group, rank, fig_path):
    """
    """
    pc = transformed[:,rank - 1]
    protein = u.select_atoms("protein")

    min_ind = np.argmin(pc)
    u.trajectory[min_ind]
    protein.write(f"{ fig_path }/PC{ rank }_min_{ group }.pdb")

    max_ind = np.argmax(pc)
    u.trajectory[max_ind]
    protein.write(f"{ fig_path }/PC{ rank }_max_{ group }.pdb")

    return None

def visualize_PC(p_components, transformed, mean, rank, atom_group, group, fig_path):
    """
    """
    pc1 = p_components[:, rank - 1]
    print(f"pc1 shape : { np.shape(pc1)}")
    trans1 = transformed[:, rank - 1]
    print(f"trans1 shape : { np.shape(trans1)}")
    projected = np.outer(trans1, pc1) + mean
    coordinates = projected.reshape(len(trans1), -1, 3)

    proj1 = mda.Merge(atom_group)
    proj1.load_new(coordinates, order="fac")

    with mda.Writer(f"{ fig_path }/pc_{ group }{ rank }.xtc", atom_group.n_atoms) as W:
        for ts in proj1.trajectory:
            if ts.frame % 100 == 0:
                W.write(proj1)

    with mda.Writer(f"{ fig_path }/pc_{ group }{ rank }.pdb") as W:
        W.write(proj1.atoms)

    return None

def get_core_res(recalc=False):
    """Finds the core residues which are immobile across the conformational states.

    Uses data from the combined simulation of the apo states open and closed simulations,
    to get the calphas of the residues with an RMSF below 1.5.

    Parameters
    ----------
    recalc : boolean
        Indicates whether the core_res array should be redetermined.

    Returns
    -------
    core_res : nd.array
        Indicies for the less mobile residues across conformational states. 
    core : str
        Selection string for the core residues.

    """
    core_res_path = f"{ path_head }/simulate/apo_state/open/data"
    if not os.path.exists(f"{ core_res_path }/core_res.npy") or recalc:
        top = f"{ core_res_path }/topol.top"
        a = mda.Universe(top, f"{ core_res_path }/simulate/holo_conf/data/full_holo_apo.xtc",
                         topology_format="ITP")
        calphas, rmsf = get_rmsf(a, top, core_res_path)
        core_res = calphas[(rmsf < 1.5)]
        utils.save_array(f"{ core_res_path }/core_res.npy", core_res)
    else:
        core_res = np.load(f"{ core_res_path }/core_res.npy")

    aln_str = "protein and name CA and ("
    core_open = [f"resid {i} or " for i in core_res]
    core_closed = [f"resid {i + 544} or " for i in core_res]
    core = aln_str + "".join((core_open + core_closed))[:-4] + ")"

    return core_res, core

if __name__ == '__main__':
    main(sys.argv)
```
