## Supplementary material for "*Clostridioides difficile* Toxins Unhinged: Allosterically Switchable Network Orients *β*-flap": Scripts, Structures etc.: README.md

# *Clostridioides difficile* Toxins Unhinged: Allosterically Switchable Network Orients $\beta$-flap

## Description
Allosteric proteins exhibit a functional response upon ligand binding far from the active site. 
*Clostridioides difficile* toxins use allosteric binding by an endogenous co-factor toorchestrate self-cleavage from within the target cell. This binding event induces a conformational shift, primarily effecting a lever-like "$`\beta`$-flap" region, with two known orientations. 
We uncovered a mechanism for this allosteric transition using extensive atomistic MD simulations and computational and experimental mutagenesis.
The mechanism relies on a switchable interaction network. 
The most prominent interaction pair is  K600–E743, with K600 interactions explaining $`\sim`$70 \% of the allosteric effect.
Rather than gradually morphing between two end states, the interaction network adopts two mutually exclusive configurations in the active and inactive state.
Similar switchable networks may explain allostery more broadly.
This mechanism in particular could aid in drug development targeting the *Clostridioides difficile* toxins autoproteolysis.

### Conformation naming
The names given to conformations changed during the development 
of this project. Within the scripts, the conformation named "closed" corresponds to the lowered conformation, while the name "open" corresponds to the "raised" conformation.

### Residue numbering
Throughout the paper, residues are numbered based on full-toxin numbering. Residue numbering in the scripts is indexed from the first modelled residue. To convert to full-toxin numbering, add 543 for TcdB and add 541 for TcdA. In particular, K57 -> K600 and E200 -> E743.

## File tree

### Paths to notable files
- Reference TcdB CPD conformations: 
    - `reproduce/ref-structures/lowered_ref_state.pdb`
    - `reproduce/ref-structures/raised_ref_state.pdb`
- Alignment structure: `ref-structures/ref_all_atoms.pdb`
- Alignment residues: `ref-structures/core_res.npy`
- Modified Amber14sb force field: `reproduce/amber14sb_ip6.ff`
- Project configuration: `config/settings.py`

### Analysis scripts
Scripts related to the analysis of MD data and figure generation. 

```md
AllosteryCPD/
- README.md
- AlphaFold/
    - AF_PCA.py
    - CombindedRMSD.py
    - ... (other scripts for AlphaFold structure generation/analysis)
- Biased/
    - FES2D.py
    - WindowAves.py
    - ... (other scripts for analysis of umbrella sampling data)
- Config/
    - settings.py
- Other/ (paper-plots)
    - AveIPL_charges.py
    - TablePlot_network.py
    - ... (additional scripts for figure generation)
- Tools/ 
    - connect.py
    - traj_funcs.py
    - utils.py
- BasicMD.py
- VectorCoordCombo.py
- ... (other basic simulation analysis scripts)
```

### Unbiased simulations
Key files for reproduction of simulation data are within the reproduce/ directory. 
These include the core unbiased simulations (apo/holo + lowered/raised), unbiased mutants (K600G, E743G, K600G-E743G) and unbiased TcdA/ (apo/holo + lowered/raised)
In general, this looks like:

```md
AllosteryCPD/
- reproduce/
    - example-system/
        - example-system.pdb (initial structure)
        - set_up.sh (define box, solvate, add ions)
        - run_sim.sh (run simulation)
        - cat_traj.sh (post-process simulation)
    - TcdA/
        - example-system/
            - example-system.pdb (initial structure)
            - run_sim.sh (define box, solvate, add ions, run simulation)
            - cat_traj.sh (post-process simulation)
    - MDP-apo/ (molecular dynamics parameters for apo simulations)
        - em_steep.mdp (energy minimization)
        - NVT.mdp (thermal equilibration)
        - NPT.mdp (pressure equilibration)
        - Production (md run)
    - MDP-holo/ (molecular dynamics parameters for holo simulations)
        - em_steep.mdp (energy minimization)
        - NVT.mdp (thermal equilibration, restraining protein and ligand)
        - NPT.mdp (pressure equilibration, restraining protein and ligand)
        - NVT2.mdp (thermal equilibration, restraining only ligand)
        - NPT2.mdp (pressure equilibration, restraining only ligand)
        - Production (md run)
    - ... (various systems, force fields)
```

### Biased simulations
Key files for reproduction of biased sampling data using umbrella sampling are within the reproduce/ directory. 
In general, this looks like:

```md
AllosteryCPD/
- reproduce/
    - umbrella-apo/
        - window1/
            - initial_conform.pdb
            - plumed_1.dat
        - ... (window2/, ..., window181/)
        - ref2.pdb (used for alignment)
        - us_array.sh (set up structure, equilibrate, simulate)
        - post_process.sh 
        - restraint_pts.csv (table of 2D umbrella positions)
    - umbrella-holo/
        - ...
    - umbrella-apo-K600G/
        - window1/
            - initial_conform.pdb
            - mutated.pdb
            - plumed_1.dat
        - ... (window2/, ..., window120/)
        - ref.pdb (used for alignment)
        - us_array.sh (set up structure, equilibrate, simulate)
        - post_process.sh 
        - restraint_pts.csv (table of 2D umbrella positions)
    - umbrella-holo-K600G/
        - ...
```
