## Supplementary material for "*Clostridioides difficile* Toxins Unhinged: Allosterically Switchable Network Orients *β*-flap": Scripts, Structures etc.: cat_trj.sh

```
#!/bin/bash
#SBATCH --time=1:00:00
#SBATCH --ntasks=1
#SBATCH --cpus-per-task=1
#SBATCH --mem=500
#SBATCH --partition=agkeller
#SBATCH --qos=standard
#SBATCH --job-name=cattraj

module add GROMACS/2021.5-foss-2021b-CUDA-11.4.1-PLUMED-2.8.0
MDP="/home/lf1071fu/unbiased_sims/af_replicas/MDP"
home=$(pwd)

for i in {1..9}; do

	cd af${i}/sim_data
	
	echo "1 1" | gmx trjconv -f af${i}.xtc -s af${i}.tpr -center yes -o centered_traj.xtc -nobackup

	echo "4 1" | gmx trjconv -f centered_traj.xtc -s af${i}.tpr -fit rot+trans -o fitted_traj.xtc -nobackup

	rm -rf centered_traj.xtc amber14sb.ff

	cd ../..

done
```
