## Supplementary material for "*Clostridioides difficile* Toxins Unhinged: Allosterically Switchable Network Orients *β*-flap": Scripts, Structures etc.: run_sim.sh

```
#!/bin/bash
#SBATCH --time=2-00:00:00
#SBATCH --ntasks=1
#SBATCH --cpus-per-task=8
#SBATCH --mem=500
#SBATCH --partition=agkeller
#SBATCH --gres=gpu
#SBATCH --qos=standard
#SBATCH --job-name=af_sim

module add GROMACS/2021.5-foss-2021b-CUDA-11.4.1-PLUMED-2.8.0
export OMP_NUM_THREADS=$SLURM_CPUS_PER_TASK
export OMP_PROC_BIND=false
MDP="/home/lf1071fu/unbiased_sims/af_replicas/MDP"
home=$(pwd)

mkdir -p equilibrate md_run

cp -r ${home}/amber14sb.ff ${home}/equilibrate
cp -r ${home}/amber14sb.ff ${home}/md_run

cp ${home}/cpd_initial.gro ${home}/posre.itp ${home}/topol.top ${home}/equilibrate
	
cd ${home}/equilibrate

if [ -f ${home}/md_run/af_${1}.cpt ]; then

	cd ${home}/md_run
	gmx_mpi mdrun -cpi af${1} -deffnm af${1} -nb gpu -update gpu -pme gpu -pin off -ntomp $SLURM_CPUS_PER_TASK -nobackup
	
else

	gmx grompp -f $MDP/em_steep.mdp -c cpd_initial.gro -p topol.top -o em_steep.tpr -nobackup
	gmx_mpi mdrun -deffnm em_steep -ntomp $SLURM_CPUS_PER_TASK -nobackup

	gmx grompp -f $MDP/NVT.mdp -c em_steep.gro -r em_steep.gro -p topol.top -o nvt.tpr -nobackup
	gmx_mpi mdrun -deffnm nvt -pin off -nb gpu -update gpu -pme gpu -ntomp $SLURM_CPUS_PER_TASK -nobackup

	gmx grompp -f $MDP/NPT.mdp -c nvt.gro -r nvt.gro -p topol.top -o npt.tpr -nobackup
	gmx_mpi mdrun -deffnm npt -pin off -nb gpu -update gpu -pme gpu -ntomp $SLURM_CPUS_PER_TASK -nobackup

	cp topol.top npt.gro npt.cpt ${current}/md_run
	cd ${current}/md_run
	
	gmx grompp -f ${MDP}/Production.mdp -c npt.gro -r npt.gro -t npt.cpt -p topol.top -o af${1}.tpr -nobackup
	gmx_mpi mdrun -deffnm af${1} -nb gpu -update gpu -pme gpu -pin off --ntomp $SLURM_CPUS_PER_TASK -nobackup
fi

if [ -f af${1}.gro ]; then

        rm -rf amber14sb.ff

fi
```
