## Supplementary material for "*Clostridioides difficile* Toxins Unhinged: Allosterically Switchable Network Orients *β*-flap": Scripts, Structures etc.: cat_trj.sh

```
#!/bin/bash
#SBATCH --time=1:00:00
#SBATCH --ntasks=1
#SBATCH --cpus-per-task=1
#SBATCH --mem=500
#SBATCH --partition=agkeller
#SBATCH --qos=standard
#SBATCH --job-name=cattraj

module add GROMACS/2021.5-foss-2021b-CUDA-11.4.1-PLUMED-2.8.0
MDP="/home/lf1071fu/unbiased_sims/MDP"
home=$(pwd)

files=$(ls -v run*/md_run/apo_closed.xtc)

commands=$(printf 'c\n%.0s' {1..19})'\nc'

echo $files[@]

echo $commands

# Concatenate trajectories
echo -e "$commands" | gmx trjcat -f ${files[@]} -o cat_trj.xtc -nobackup -settime

echo "1 1" | gmx trjconv -f cat_trj.xtc -s run1/md_run/apo_closed.tpr -center yes -o centered_traj.xtc -nobackup

echo "4 1" | gmx trjconv -f centered_traj.xtc -s run1/md_run/apo_closed.tpr -fit rot+trans -o fitted_traj.xtc -nobackup

echo "1" | gmx trjconv -f fitted_traj.xtc -s run1/md_run/apo_closed.tpr -o fitted_traj_100.xtc -skip 100 -nobackup

rm cat_trj.xtc centered_traj.xtc
```
