## Supplementary material for "*Clostridioides difficile* Toxins Unhinged: Allosterically Switchable Network Orients *β*-flap": Scripts, Structures etc.: run_sim.sh

```
#!/bin/bash
#SBATCH --time=14-00:00:00
#SBATCH --ntasks=1
#SBATCH --cpus-per-task=8
#SBATCH --mem=500
#SBATCH --partition=agkeller
#SBATCH --gres=gpu
#SBATCH --qos=standard
#SBATCH --job-name=apo_open

module add GROMACS/2021.5-foss-2021b-CUDA-11.4.1-PLUMED-2.8.0
export OMP_NUM_THREADS=$SLURM_CPUS_PER_TASK
export OMP_PROC_BIND=false
MDP="/home/lf1071fu/unbiased_sims/MDP"
home=$(pwd)

for i in {1..20}
do

	mkdir -p run$i
	cd run$i
	current=$(pwd)
	mkdir -p ${current}/equilibrate ${current}/md_run

	cp -r ${home}/amber14sb.ff ${current}/equilibrate
	cp -r ${home}/amber14sb.ff ${current}/md_run


	if [ "$i" -eq 1 ]; then

        	cp ${home}/cpd_initial.gro ${home}/posre.itp ${home}/topol.top ${current}/equilibrate
	
	else

        	cd ${home}/run$((i - 1))
        	prev=$(pwd)
        	cd ${prev}/equilibrate
        	cp index.ndx posre.itp topol.top ${current}/equilibrate
		cp ${prev}/md_run/apo_open.gro ${current}/equilibrate/cpd_initial.gro

        fi
	
	cd ${current}/equilibrate

	if [ -f ${current}/md_run/apo_open.cpt ]; then

		cd ${current}/md_run
		gmx_mpi mdrun -cpi apo_open -deffnm apo_open -nb gpu -update gpu -pme gpu -pin off -ntomp $SLURM_CPUS_PER_TASK -nobackup
	
	else


		gmx grompp -f $MDP/em_steep.mdp -c cpd_initial.gro -p topol.top -o em_steep.tpr -nobackup
		gmx_mpi mdrun -deffnm em_steep -ntomp $SLURM_CPUS_PER_TASK -nobackup

		gmx grompp -f $MDP/NVT.mdp -c em_steep.gro -r em_steep.gro -p topol.top -o nvt.tpr -nobackup
		gmx_mpi mdrun -deffnm nvt -pin off -nb gpu -update gpu -pme gpu -ntomp $SLURM_CPUS_PER_TASK -nobackup
	        cp ${prev}/md_run/holo_open.gro ./holo_o_initial.gro

		gmx grompp -f $MDP/NPT.mdp -c nvt.gro -r nvt.gro -p topol.top -o npt.tpr -nobackup
		gmx_mpi mdrun -deffnm npt -pin off -nb gpu -update gpu -pme gpu -ntomp $SLURM_CPUS_PER_TASK -nobackup

		cp topol.top npt.gro npt.cpt ${current}/md_run
		cd ${current}/md_run
	
		gmx grompp -f ${MDP}/Production.mdp -c npt.gro -r npt.gro -t npt.cpt -p topol.top -o apo_open.tpr -nobackup
		gmx_mpi mdrun -deffnm apo_open -nb gpu -update gpu -pme gpu -pin off --ntomp $SLURM_CPUS_PER_TASK -nobackup

	fi

	if [ -f apo_open.gro ]; then

        	### Post processing
	        # Centering and fitting of trajectory
        	echo "1 1" | gmx trjconv -f apo_open.xtc -s apo_open.tpr -pbc mol -center yes -o centered_traj.xtc -nobackup
	        echo "4 1" | gmx trjconv -f centered_traj.xtc -s apo_open.tpr -fit rot+trans -o fitted_traj.xtc -nobackup

        	rm -rf amber14sb.ff centered_traj.xtc
	fi

	cd $home

done
```
