## Supplementary material for "*Clostridioides difficile* Toxins Unhinged: Allosterically Switchable Network Orients *β*-flap": Scripts, Structures etc.: run_sim.sh

```
#!/bin/bash
#SBATCH --time=7-00:00:00
#SBATCH --ntasks=1
#SBATCH --cpus-per-task=8
#SBATCH --mem=10G
#SBATCH --partition=gpu
#SBATCH --gres=gpu:a100:1
#SBATCH --job-name=mut200

module add bio/GROMACS/2021.5-foss-2021b-CUDA-11.4.1-PLUMED-2.8.0
export OMP_NUM_THREADS=$SLURM_CPUS_PER_TASK
export OMP_PROC_BIND=false
MDP="/scratch/hpc-prf-cpdallo/mutation/MDP"
home=$(pwd)

for i in {17..20}
do

        mkdir -p run$i
        cd run$i
        current=$(pwd)
        mkdir -p ${current}/equilibrate ${current}/md_run

        cp -r ${home}/amber14sb.ff ${current}/equilibrate
        cp -r ${home}/amber14sb.ff ${current}/md_run


        if [ "$i" -eq 1 ]; then

                cp ${home}/cpd_initial.gro ${home}/posre.itp ${home}/topol.top ${current}/equilibrate

        else

                cd ${home}/run$((i - 1))
                prev=$(pwd)
                cd ${prev}/equilibrate
                cp index.ndx posre.itp topol.top ${current}/equilibrate
                cp ${prev}/md_run/E200G.gro ${current}/equilibrate/cpd_initial.gro

        fi

        cd ${current}/equilibrate

        if [ -f ${current}/md_run/E200G.cpt ]; then

                cd ${current}/md_run
                gmx_mpi mdrun -cpi E200G -deffnm E200G -nb gpu -update gpu -pme gpu -pin off -ntomp $SLURM_CPUS_PER_TASK -nobackup

        else


                gmx grompp -f $MDP/em_steep.mdp -c cpd_initial.gro -p topol.top -o em_steep.tpr -nobackup
                gmx_mpi mdrun -deffnm em_steep -ntomp $SLURM_CPUS_PER_TASK -nobackup

                gmx grompp -f $MDP/NVT.mdp -c em_steep.gro -r em_steep.gro -p topol.top -o nvt.tpr -nobackup
                gmx_mpi mdrun -deffnm nvt -pin off -nb gpu -update gpu -pme gpu -ntomp $SLURM_CPUS_PER_TASK -nobackup

                gmx grompp -f $MDP/NPT.mdp -c nvt.gro -r nvt.gro -p topol.top -o npt.tpr -nobackup
                gmx_mpi mdrun -deffnm npt -pin off -nb gpu -update gpu -pme gpu -ntomp $SLURM_CPUS_PER_TASK -nobackup

                cp topol.top npt.gro npt.cpt ${current}/md_run
                cd ${current}/md_run

                gmx grompp -f ${MDP}/Production.mdp -c npt.gro -r npt.gro -t npt.cpt -p topol.top -o E200G.tpr -nobackup
                gmx_mpi mdrun -deffnm E200G -nb gpu -update gpu -pme gpu -pin off --ntomp $SLURM_CPUS_PER_TASK -nobackup

        fi

        if [ -f E200G.gro ]; then

                rm -rf amber14sb.ff
        fi

        cd $home

done
```
