## Supplementary material for "*Clostridioides difficile* Toxins Unhinged: Allosterically Switchable Network Orients *β*-flap": Scripts, Structures etc.: run_sim.sh

```
#!/bin/bash
#SBATCH --time=14-00:00:00
#SBATCH --ntasks=1
#SBATCH --cpus-per-task=8
#SBATCH --mem=500
#SBATCH --partition=agkeller
#SBATCH --gres=gpu
#SBATCH --qos=standard
#SBATCH --job-name=holo_closed

module add GROMACS/2021.5-foss-2021b-CUDA-11.4.1-PLUMED-2.8.0
export OMP_NUM_THREADS=$SLURM_CPUS_PER_TASK
export OMP_PROC_BIND=false
MDP="/home/lf1071fu/unbiased_sims/holo_closed/MDP"
home=$(pwd)

for i in {1..20}
do

	mkdir -p run$i
	cd run$i
	current=$(pwd)
	mkdir -p ${current}/equilibrate ${current}/md_run

	cp -r ${home}/amber14sb_ip6.ff ${current}/equilibrate
	cp -r ${home}/amber14sb_ip6.ff ${current}/md_run


	if [ "$i" -eq 1 ]; then

        	cp ${home}/cpd_initial.gro ${home}/posre* ${home}/topol* ${current}/equilibrate
		cd equilibrate
		echo -e "1 | 20\nq" | gmx make_ndx -f cpd_initial.gro -o index.ndx -nobackup
	
	else

        	cd ${home}/run$((i - 1))
        	prev=$(pwd)
        	cd ${prev}/equilibrate
        	cp index.ndx posre* topol* ${current}/equilibrate
		cp ${prev}/md_run/holo_closed.gro ${current}/equilibrate/cpd_initial.gro

		cd ${current}/equilibrate

	fi

	if [ -f ${current}/md_run/holo_closed.cpt ]; then

		cd ${current}/md_run
		gmx_mpi mdrun -cpi holo_closed -deffnm holo_closed -nb gpu -update gpu -pme gpu -pin off -ntomp $SLURM_CPUS_PER_TASK -nobackup
	
	else


		gmx grompp -f $MDP/em_steep.mdp -n index.ndx -c cpd_initial.gro -p topol.top -o em_steep.tpr -nobackup
		gmx_mpi mdrun -deffnm em_steep -ntomp $SLURM_CPUS_PER_TASK -nobackup

		gmx grompp -f $MDP/NVT.mdp -n index.ndx -c em_steep.gro -r em_steep.gro -p topol.top -o nvt.tpr -nobackup
		gmx_mpi mdrun -deffnm nvt -pin off -nb gpu -update gpu -pme gpu -ntomp $SLURM_CPUS_PER_TASK -nobackup

                if [ "${i}" -eq 1 ]; then
			
			# NPT restraining Protein and Ligand
			gmx grompp -f $MDP/NPT.mdp -n index.ndx -c nvt.gro -r nvt.gro -p topol.top -o npt1.tpr -nobackup
	                gmx_mpi mdrun -deffnm npt1 -pin off -nb gpu -update gpu -pme gpu -ntomp $SLURM_CPUS_PER_TASK -nobackup

			# NVT then NPT, restraining only the Ligand
                        gmx grompp -f $MDP/NVT2.mdp -n index.ndx -c npt1.gro -r npt1.gro -p topol.top -o nvt2.tpr -nobackup
                        gmx_mpi mdrun -deffnm nvt2 -pin off -nb gpu -update gpu -pme gpu -ntomp $SLURM_CPUS_PER_TASK -nobackup
                        gmx grompp -f $MDP/NPT2.mdp -n index.ndx -c nvt2.gro -r nvt2.gro -p topol.top -o npt.tpr -nobackup
			gmx_mpi mdrun -deffnm npt -pin off -nb gpu -update gpu -pme gpu -ntomp $SLURM_CPUS_PER_TASK -nobackup

		else

			gmx grompp -f $MDP/NPT.mdp -n index.ndx -c nvt.gro -r nvt.gro -p topol.top -o npt.tpr -nobackup
	                gmx_mpi mdrun -deffnm npt -pin off -nb gpu -update gpu -pme gpu -ntomp $SLURM_CPUS_PER_TASK -nobackup

		fi


		cp topol* npt.cpt npt.gro index.ndx posre* ${current}/md_run
		cd ${current}/md_run
	
		gmx grompp -f ${MDP}/Production.mdp -n index.ndx -c npt.gro -r npt.gro -t npt.cpt -p topol.top -o holo_closed.tpr -nobackup
		gmx_mpi mdrun -deffnm holo_closed -nb gpu -update gpu -pme gpu -pin off -ntomp $SLURM_CPUS_PER_TASK -nobackup


	fi

	if [ -f holo_closed.gro ]; then

        	### Post processing
	        # Centering and fitting of trajectory

        	rm -rf amber14sb_ip6.ff 
	fi

	cd $home

done
```
