## Supplementary material for "*Clostridioides difficile* Toxins Unhinged: Allosterically Switchable Network Orients *β*-flap": Scripts, Structures etc.: set_up.sh

```
#!/bin/bash
#SBATCH --time=1:00:00
#SBATCH --ntasks=1
#SBATCH --cpus-per-task=1
#SBATCH --mem=500
#SBATCH --partition=agkeller
#SBATCH --job-name=holo_closed
#SBATCH --qos=standard

module add GROMACS/2021.5-foss-2021b-CUDA-11.4.1-PLUMED-2.8.0
export OMP_NUM_THREADS=$SLURM_CPUS_PER_TASK
export OMP_PROC_BIND=false
MDP="/home/lf1071fu/unbiased_sims/holo_closed/MDP"
home=$(pwd)

# Generate topology using .pdb of capped residue
gmx pdb2gmx -ff amber14sb_ip6 -f holo_closed.pdb -o complex_only.gro -ignh -water tip3p -nobackup

# Define dodecahedron box with ligand at center, > 1.2 nm to edge
gmx editconf -f complex_only.gro -o complex_box.gro -c -d 1.2 -bt dodecahedron -nobackup

# Solvate ligand with TIP3P
gmx solvate -cp complex_box.gro -cs spc216 -o complex_tip3p.gro -p topol.top -nobackup

# Add ions if needed
gmx grompp -f ${MDP}/em_steep.mdp -c complex_tip3p.gro -p topol.top -o ions.tpr -maxwarn 1 -nobackup
echo "SOL" | gmx genion -s ions.tpr -o cpd_initial.gro -p topol.top -pname NA -pq 1 -np 42 -nname CL -nq -1 -nn 16 -nobackup
```
