## Supplementary material for "*Clostridioides difficile* Toxins Unhinged: Allosterically Switchable Network Orients *β*-flap": Scripts, Structures etc.: cat_trj.sh

```
#!/bin/bash
#SBATCH --time=3:00:00
#SBATCH --cpus-per-task=1
#SBATCH --mem=10G
#SBATCH --partition=normal
#SBATCH --job-name=cat_mut
#SBATCH --qos=express

module add bio/GROMACS/2021.5-foss-2021b-CUDA-11.4.1-PLUMED-2.8.0

MDP="/scratch/hpc-prf-cpdallo/mutation/MDP"

files=$(ls -v run*/md_run/double_mut.xtc)

commands=$(printf 'c\n%.0s' {1..32})'\nc'

echo $files[@]

echo $commands

# Concatenate trajectories
echo -e "$commands" | gmx trjcat -f ${files[@]} -o cat_trj.xtc -nobackup -settime

echo "1 1" | gmx trjconv -f cat_trj.xtc -s run1/md_run/double_mut.tpr -center yes -o centered_traj.xtc -nobackup

echo "4 1" | gmx trjconv -f centered_traj.xtc -s run1/md_run/double_mut.tpr -fit rot+trans -o fitted_traj.xtc -nobackup

echo "1" | gmx trjconv -f fitted_traj.xtc -o fitted_traj_100.xtc -nobackup -skip 100

rm cat_trj.xtc centered_traj.xtc
```
