## Supplementary material for "*Clostridioides difficile* Toxins Unhinged: Allosterically Switchable Network Orients *β*-flap": Scripts, Structures etc.: run_sim.sh

```
#!/bin/bash
#SBATCH --time=6-00:00:00
#SBATCH --ntasks=1
#SBATCH --cpus-per-task=8
#SBATCH --mem=20G
#SBATCH --gres=gpu:a100:1
#SBATCH --partition=gpu
#SBATCH --job-name=closed_apo


module add bio/GROMACS/2021.5-foss-2021b-CUDA-11.4.1-PLUMED-2.8.0
export OMP_NUM_THREADS=$SLURM_CPUS_PER_TASK
export OMP_PROC_BIND=false
MDP="/scratch/hpc-prf-cpdallo/tcda/closed_apo/MDP"
home=$(pwd)


for i in {1..4}
do

        mkdir -p run$i
        cd run$i
        current=$(pwd)
        mkdir -p ${current}/equilibrate ${current}/md_run

        cp -r ${home}/amber14sb.ff ${current}/equilibrate
        cp -r ${home}/amber14sb.ff ${current}/md_run

        if [ -f closed_apo.cpt ]; then

                gmx_mpi mdrun -cpi closed_apo -deffnm open_apo -nb gpu -update gpu -pme gpu -pin off -ntomp $SLURM_CPUS_PER_TASK -nobackup


        elif [ "$i" -eq 1 ]; then

		mkdir -p ${current}/setup

		cp -r ${home}/closed_apo.pdb ${home}/amber14sb.ff ${current}/setup
                cd ${current}/setup

                ### Set up initial system by generating box, solvating etc.
                # Generate topology using .pdb of capped residue
                echo "1 1 1" | gmx pdb2gmx -ff amber14sb -f closed_apo.pdb -o cpd_only.gro -water tip3p -nobackup -ignh -his
                sed -i "s/HISE/ HIS/" topol.top

                # Define dodecahedron box with ligand at center, > 1.2 nm to edge
                gmx editconf -f cpd_only.gro -o cpd_box.gro -c -d 1.2 -bt dodecahedron -nobackup

                # Solvate ligand with TIP3P
                gmx solvate -cp cpd_box.gro -cs spc216 -o cpd_tip3p.gro -p topol.top -nobackup

                # Add ions as needed
                gmx grompp -f ${MDP}/em_steep.mdp -c cpd_tip3p.gro -p topol.top -o ions.tpr -maxwarn 1 -nobackup
                echo "SOL" | gmx genion -s ions.tpr -o cpd_initial.gro -p topol.top -pname NA -pq 1 -np 36 -nname CL -nq -1 -nn 30 -nobackup

                cp ${current}/setup/cpd_initial.gro ${current}/setup/posre.itp ${current}/setup/topol.top ${current}/equilibrate
		cd ${current}/equilibrate

        else

                cd ${home}/run$((i - 1))
                prev=$(pwd)
                cd ${prev}/equilibrate
                cp posre.itp topol.top ${current}/equilibrate
                cp ${prev}/md_run/closed_apo.gro ${current}/equilibrate/cpd_initial.gro

                cd ${current}/equilibrate
        fi


        gmx grompp -f $MDP/em_steep.mdp -c cpd_initial.gro -p topol.top -o em_steep.tpr -nobackup
        gmx_mpi mdrun -deffnm em_steep -ntomp $SLURM_CPUS_PER_TASK -nobackup

        gmx grompp -f $MDP/NVT.mdp -c em_steep.gro -r em_steep.gro -p topol.top -o nvt.tpr -nobackup
        gmx_mpi mdrun -deffnm nvt -pin off -nb gpu -update gpu -pme gpu -ntomp $SLURM_CPUS_PER_TASK -nobackup

        gmx grompp -f $MDP/NPT.mdp -c nvt.gro -r nvt.gro -p topol.top -o npt.tpr -nobackup
        gmx_mpi mdrun -deffnm npt -pin off -nb gpu -update gpu -pme gpu -ntomp $SLURM_CPUS_PER_TASK -nobackup

        cp topol.top npt.gro npt.cpt ${current}/md_run
        cd ${current}/md_run
   
	gmx grompp -f ${MDP}/Production.mdp -c npt.gro -r npt.gro -t npt.cpt -p topol.top -o closed_apo.tpr -nobackup
        gmx_mpi mdrun -deffnm closed_apo -nb gpu -update gpu -pme gpu -pin off --ntomp $SLURM_CPUS_PER_TASK -nobackup

        if [ -f closed_apo.gro ]; then

                ### Post processing
                # Centering and fitting of trajectory
                echo "1 1" | gmx trjconv -f closed_apo.xtc -s closed_apo.tpr -pbc mol -center yes -o centered_traj.xtc -nobackup
                echo "4 1" | gmx trjconv -f centered_traj.xtc -s closed_apo.tpr -fit rot+trans -o fitted_traj.xtc -nobackup

                rm -rf amber14sb.ff centered_traj.xtc
        fi

        cd $home

done
```
