## Supplementary material for "*Clostridioides difficile* Toxins Unhinged: Allosterically Switchable Network Orients *β*-flap": Scripts, Structures etc.: run_sim.sh

```
#!/bin/bash
#SBATCH --time=6-00:00:00
#SBATCH --ntasks=1
#SBATCH --cpus-per-task=8
#SBATCH --mem=20G
#SBATCH --gres=gpu:a100:1
#SBATCH --partition=gpu
#SBATCH --job-name=closed_holo


module add bio/GROMACS/2021.5-foss-2021b-CUDA-11.4.1-PLUMED-2.8.0
export OMP_NUM_THREADS=$SLURM_CPUS_PER_TASK
export OMP_PROC_BIND=false
MDP="/scratch/hpc-prf-cpdallo/tcda/closed_holo/MDP"
home=$(pwd)


for i in {1..4}
do

        mkdir -p run$i
        cd run$i
        current=$(pwd)
        mkdir -p ${current}/equilibrate ${current}/md_run

        cp -r ${home}/amber14sb_ip6.ff ${current}/equilibrate
        cp -r ${home}/amber14sb_ip6.ff ${current}/md_run

        if [ -f closed_holo.cpt ]; then

                gmx_mpi mdrun -cpi closed_holo -deffnm closed_holo -nb gpu -update gpu -pme gpu -pin off -ntomp $SLURM_CPUS_PER_TASK -nobackup


        elif [ "$i" -eq 1 ]; then

		mkdir -p ${current}/setup

		cp -r ${home}/closed_holo.pdb ${home}/amber14sb_ip6.ff ${current}/setup
                cd ${current}/setup

                ### Set up initial system by generating box, solvating etc.
                # Generate topology using .pdb of capped residue
                echo "1 1 1" | gmx pdb2gmx -ff amber14sb_ip6 -f closed_holo.pdb -o complex_only.gro -water tip3p -nobackup -ignh -his
                sed -i "s/HISE/ HIS/" topol.*
	        echo '#ifdef RESLIG' >> topol_Other_chain_B.itp
        	echo '#include "posre_Other_chain_B.itp"' >> topol_Other_chain_B.itp
	        echo '#endif' >> topol_Other_chain_B.itp
        	echo >> topol_Other_chain_B.itp

                # Define dodecahedron box with ligand at center, > 1.2 nm to edge
                gmx editconf -f complex_only.gro -o complex_box.gro -c -d 1.2 -bt dodecahedron -nobackup

                # Solvate ligand with TIP3P
                gmx solvate -cp complex_box.gro -cs spc216 -o complex_tip3p.gro -p topol.top -nobackup

                # Add ions as needed
                gmx grompp -f ${MDP}/em_steep.mdp -c complex_tip3p.gro -p topol.top -o ions.tpr -maxwarn 1 -nobackup
                echo "SOL" | gmx genion -s ions.tpr -o complex_initial.gro -p topol.top -pname NA -pq 1 -np 42 -nname CL -nq -1 -nn 27 -nobackup
		echo -e "1 |20\nq" | gmx make_ndx -f complex_initial.gro -o index.ndx -nobackup

                cp ${current}/setup/complex_initial.gro ${current}/setup/posre* ${current}/setup/topol* ${current}/setup/index.ndx ${current}/equilibrate
		cd ${current}/equilibrate

        else

                cd ${home}/run$((i - 1))
                prev=$(pwd)
                cd ${prev}/equilibrate
                cp posre* topol* index.ndx ${current}/equilibrate
                cp ${prev}/md_run/closed_holo.gro ${current}/equilibrate/complex_initial.gro

                cd ${current}/equilibrate
        fi


		gmx grompp -f $MDP/em_steep.mdp -n index.ndx -c complex_initial.gro -p topol.top -o em_steep.tpr -nobackup
		gmx_mpi mdrun -deffnm em_steep -ntomp $SLURM_CPUS_PER_TASK -nobackup

		gmx grompp -f $MDP/NVT.mdp -n index.ndx -c em_steep.gro -r em_steep.gro -p topol.top -o nvt.tpr -nobackup
		gmx_mpi mdrun -deffnm nvt -pin off -nb gpu -update gpu -pme gpu -ntomp $SLURM_CPUS_PER_TASK -nobackup

	if [ "${i}" -eq 1 ]; then
			
		# NPT restraining Protein and Ligand
		gmx grompp -f $MDP/NPT.mdp -n index.ndx -c nvt.gro -r nvt.gro -p topol.top -o npt1.tpr -nobackup
	        gmx_mpi mdrun -deffnm npt1 -pin off -nb gpu -update gpu -pme gpu -ntomp $SLURM_CPUS_PER_TASK -nobackup

		# NVT then NPT, restraining only the Ligand
                gmx grompp -f $MDP/NVT2.mdp -n index.ndx -c npt1.gro -r npt1.gro -p topol.top -o nvt2.tpr -nobackup
                gmx_mpi mdrun -deffnm nvt2 -pin off -nb gpu -update gpu -pme gpu -ntomp $SLURM_CPUS_PER_TASK -nobackup
                gmx grompp -f $MDP/NPT2.mdp -n index.ndx -c nvt2.gro -r nvt2.gro -p topol.top -o npt.tpr -nobackup
		gmx_mpi mdrun -deffnm npt -pin off -nb gpu -update gpu -pme gpu -ntomp $SLURM_CPUS_PER_TASK -nobackup

	else

		gmx grompp -f $MDP/NPT.mdp -n index.ndx -c nvt.gro -r nvt.gro -p topol.top -o npt.tpr -nobackup
	        gmx_mpi mdrun -deffnm npt -pin off -nb gpu -update gpu -pme gpu -ntomp $SLURM_CPUS_PER_TASK -nobackup

	fi

        cp topol* npt.gro npt.cpt posre* index.ndx ${current}/md_run
        cd ${current}/md_run

	# Production run   
	gmx grompp -f ${MDP}/Production.mdp -c npt.gro -r npt.gro -t npt.cpt -p topol.top -o closed_holo.tpr -n index.ndx -nobackup
        gmx_mpi mdrun -deffnm closed_holo -nb gpu -update gpu -pme gpu -pin off --ntomp $SLURM_CPUS_PER_TASK -nobackup

        if [ -f closed_holo.gro ]; then

	        ### Post processing
        	# Centering and fitting of trajectory
	        echo "24 24 24" | gmx trjconv -f closed_holo.xtc -s closed_holo.tpr -center yes -pbc cluster -o centered_traj.xtc -n index.ndx -nobackup

	        echo "4 24" | gmx trjconv -f centered_traj.xtc -s closed_holo.tpr -fit rot+trans -o fitted_traj.xtc -n index.ndx -nobackup

	        rm -rf amber14sb_ip6.ff centered_traj.xtc
        
	fi

        cd $home

done
```
