## Supplementary material for "*Clostridioides difficile* Toxins Unhinged: Allosterically Switchable Network Orients *β*-flap": Scripts, Structures etc.: cat_trj.sh

```
#!/bin/bash

files=$(ls -v window*/run*/fitted_traj.xtc)

commands=$(printf 'c\n%.0s' {1..723})'\nc'

echo $commands

# Concatenate trajectories
echo -e "$commands" | gmx22 trjcat -f ${files[@]} -o cat_trj.xtc -nobackup -settime

echo "4 1" | gmx22 trjconv -f cat_trj.xtc -s window2/run2/w2_r2.tpr -fit rot+trans -o fitted_traj.xtc -nobackup

rm cat_trj.xtc
~               
```
