## Supplementary material for "*Clostridioides difficile* Toxins Unhinged: Allosterically Switchable Network Orients *β*-flap": Scripts, Structures etc.: us_array.sh

```
#!/bin/bash
#SBATCH --time=12:00:00
#SBATCH --ntasks=1
#SBATCH --cpus-per-task=16
#SBATCH --mem=10G
#SBATCH --gres=gpu:a100:1
#SBATCH --partition=gpu
#SBATCH --job-name=us_vec
#SBATCH --array=1-724

module add bio/GROMACS/2021.5-foss-2021b-CUDA-11.4.1-PLUMED-2.8.0
export OMP_NUM_THREADS=$SLURM_CPUS_PER_TASK
export OMP_PROC_BIND=false

window=$((($SLURM_ARRAY_TASK_ID - 1) / 4 + 1))
run=$((($SLURM_ARRAY_TASK_ID - 1) % 4 + 1))

# PATH VARS
MDP="/scratch/hpc-prf-cpdallo/2dfes/MDP"
cd /scratch/hpc-prf-cpdallo/2dfes/window${window}/run${run}

if [ -f w${window}_r${run}.cpt ]; then 

	gmx_mpi mdrun -plumed plumed_${window}.dat -cpi w${window}_${run} -deffnm w${window}_r${run} -nb gpu -update gpu -pme gpu -pin off -ntomp $SLURM_CPUS_PER_TASK -nobackup
	

else
	### Set up initial system by generating box, solvating etc.
	# Generate topology using .pdb of capped residue
	echo "1 1 1" | gmx pdb2gmx -ff amber14sb -f initial_conform.pdb -o cpd_only.gro -water tip3p -nobackup -ignh -his
	sed -i "s/HISE/ HIS/" topol.top

	# Define dodecahedron box with ligand at center, > 1.2 nm to edge
	gmx editconf -f cpd_only.gro -o cpd_box.gro -c -d 1.2 -bt dodecahedron -nobackup

	# Solvate ligand with TIP3P
	gmx solvate -cp cpd_box.gro -cs spc216 -o cpd_tip3p.gro -p topol.top -nobackup

	# Add ions as needed
	gmx grompp -f ${MDP}/em_steep.mdp -c cpd_tip3p.gro -p topol.top -o ions.tpr -maxwarn 1 -nobackup
	echo "SOL" | gmx genion -s ions.tpr -o cpd_initial.gro -p topol.top -pname NA -pq 1 -np 42 -nname CL -nq -1 -nn 23 -nobackup

	## Simulate the system
	# Find local minimum and equilibrate system
	gmx grompp -f ${MDP}/em_steep.mdp -c cpd_initial.gro -p topol.top -o em_steep.tpr -nobackup
	gmx_mpi mdrun -deffnm em_steep -nobackup -ntomp $SLURM_CPUS_PER_TASK -pin off

	gmx grompp -f ${MDP}/NVT.mdp -c em_steep.gro -r em_steep.gro -p topol.top -o nvt.tpr -nobackup
	gmx_mpi mdrun -deffnm nvt -nb gpu -update gpu -pme gpu -pin off -ntomp $SLURM_CPUS_PER_TASK -nobackup

	gmx grompp -f ${MDP}/NPT.mdp -c nvt.gro -r nvt.gro -p topol.top -o npt.tpr -nobackup
	gmx_mpi mdrun -deffnm npt -nb gpu -update gpu -pme gpu -pin off -ntomp $SLURM_CPUS_PER_TASK -nobackup

	# Production run
	gmx grompp -f ${MDP}/Production.mdp -c npt.gro -r npt.gro -t npt.cpt -p topol.top -o w${window}_r${run}.tpr -nobackup
	gmx_mpi mdrun -plumed plumed_${window}.dat -deffnm w${window}_r${run} -nb gpu -update gpu -pme gpu -pin off -ntomp $SLURM_CPUS_PER_TASK -nobackup

fi

if [ -f w${window}_r${run}.gro ]; then

	### Post processing
	# Centering and fitting of trajectory
	echo "1 1" | gmx trjconv -f w${window}_r${run}.xtc -s w${window}_r${run}.tpr -pbc mol -center yes -o centered_traj.xtc -nobackup
	echo "4 1" | gmx trjconv -f centered_traj.xtc -s w${window}_r${run}.tpr -fit rot+trans -o fitted_traj.xtc -nobackup

	rm -rf amber14sb.ff

	rm centered_traj.xtc

fi

cd $home
```
